# Fast and Luminous: CLIP-tag2

**DOI:** 10.64898/2026.03.05.709795

**Authors:** Veselin Nasufovic, Anamarija Pišpek, Stefanie Kühn, Manuel Bibrowski, Jonas Fischer, Jonas Wilhelm, Birgit Koch, Julian Kompa, Runyu Mao, Miroslaw Tarnawski, Julien Hiblot, Kai Johnsson

**Affiliations:** Department of Chemical Biology, Max Planck Institute for Medical Research, Heidelberg, Germany; Protein Expression and Characterization Facility, Max Planck Institute for Medical Research, Heidelberg, Germany; Institute of Chemical Sciences and Engineering, École Polytechnique Fédérale de Lausanne, Lausanne, Switzerland

## Abstract

CLIP-tag is a self-labeling protein tag used for the specific fluorescence labeling of proteins. However, its low labeling speed and the poor cell permeability of its substrates result in low labeling efficiencies in live-cell applications. Here, we introduce a substrate optimized for live-cell applications, as well as an engineered CLIP-tag variant, CLIP-tag2, which reacts with the new substrate almost 1000-fold faster than the original CLIP-tag-substrate pair. CLIP-tag2 fusion proteins can be specifically and efficiently fluorescently labeled in cells within minutes at nanomolar substrate concentrations, and can be multiplexed with other self-labeling tags such as SNAP-tag2 and HaloTag7. These advances establish CLIP-tag2 as a powerful tagging platform for high-performance live-cell bioimaging.

## Introduction

Live-cell fluorescence microscopy generally requires the selective labeling of molecules of interest with suitable fluorescent probes.^1-3^ Among the available labeling strategies for proteins, self-labeling protein (SLP) tags stand out by their ease of use and flexibility.^4,5^ These tags react irreversibly and selectively with synthetic probes, with the commonly used SLPs being SNAP-tag and HaloTag7.^6,7^ SLPs combine the specificity of genetic tagging with the superior spectral coverage, brightness and photostability of synthetic fluorophores.^4,8-11^ They are also applied in biosensor engineering, affinity-based pull-down assays^12-15^ and have been used in *ex vivo* imaging of tissues and organs, as well as in *in vivo* imaging.^16^

The performance of SLPs in live-cell fluorescence imaging is largely determined by the labeling rates and cell permeabilities of their substrates.^17^ HaloTag7 and SNAP-tag2 react with their respective rhodamine-conjugated substrates with second-order rate constants (*k*_app_) of around 10^7^ M^-1^ s^-1^, and the relatively good cell permeability of these fluorescent substrates in live cells allows for labeling at nanomolar concentrations within minutes.^18-20^

An SLP with orthogonal substrate specificity to HaloTag7 and SNAP-tag2 is CLIP-tag, which undergoes an irreversible labeling reaction with *O*^*6*^-benzylcytosine (BC) conjugates (Figure 1A).^21^ CLIP-tag has been developed from SNAP-tag through 8 amino acid substitutions that confer orthogonality between the two tags, and reacts with BC with a second-order rate constant (*k*_app_) of around 10^4^ M^-1^s^-1^.^22,23^ The low labeling speed of CLIP-tag with BC conjugates in combination with the relatively low cell permeability of its substrates limits the utility of CLIP-tag for live-cell fluorescence labeling.^24^ Applying a strategy we used for the improvement of SNAP-tag2,^19^ we report here the development of a CLIP-tag substrate that enables more efficient live-cell labeling, as well as the subsequent engineering of CLIP-tag protein for faster labeling kinetics with this new substrate.

**Figure 1.**
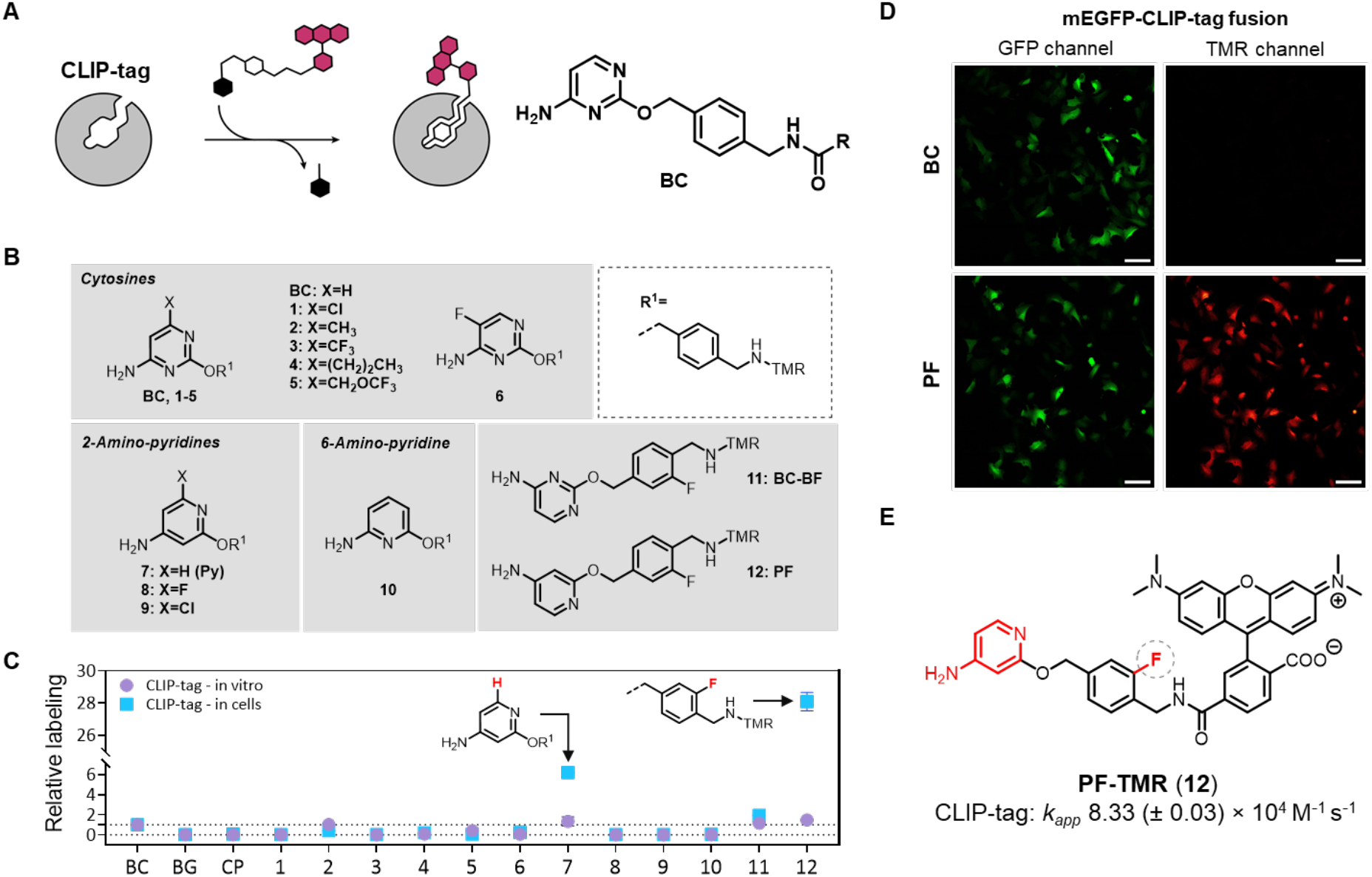
CLIP-tag substrate engineering: (**A**) Scheme of CLIP-tag labeling reaction with a generalized BC substrate. R represent the functional moiety to be conjugated to CLIP-tag. (**B**) Engineered CLIP-tag substrates. (**C**) Labeling kinetics of substrates synthesized in (**B**). Live cell assay was performed with the original CLIP-tag, whereas the biochemical assay here and in all following experiments was performed with CLIP-tag with the E30R mutation (i.e. CLIP_f_-tag), which results in a two-fold higher reactivity than parental CLIP-tag.^18,24^ In the biochemical assay, labeling kinetics of CLIP-tag with new substrates were measured by recording fluorescence polarization traces over time. Apparent second-order rate constants (*k*_*app*_) were calculated (Table S1) and normalized to the *k*_*app*_ of CLIP-tag with BC-TMR. Live-cell performance of new substrates was tested by labeling of U2OS cells that stably express a mEGFP-CLIP-tag fusion protein with TMR-substrates at 100 nM for 2 h. Cells were washed 2 times with imaging medium and analyzed *via* flow cytometry. Fluorescence intensity ratios of TMR/mEGFP were calculated (Table S1) and normalized to the ratio obtained for CLIP-tag with BC-TMR, as described previously.^19^ (**D**) Confocal laser scanning microscopy (CLSM) imaging of a mEGFP-CLIP-tag fusion expressed in U2OS cell line (described in (**C**)) labelled with 100 nM of BC-TMR and PF-TMR for 2 hours. Scale bar 100 µm. (**E**) Structure of PF-TMR (12) and the *k*_app_ for reaction with CLIP-tag.

## Results and Discussion

We first tested whether introduction of additional, mostly electron-withdrawing substituents to the cytosine leaving group of BC will increase the labeling efficiency of CLIP-tag (Figure 1B). These substrates were conjugated to tetramethylrhodamine (TMR) (Figure S1). However, all of these TMR derivatives (Figure 1B, **1**-**6**) were either unreactive or showed lower labeling efficiency than BC-TMR, both in vitro and in live cells (Figure 1C). We then applied the so-called nitrogen walk approach,^25^ and replaced the pyrimidine in BC with a pyridine (Figure 1B, **7-10**). For 4-amino pyridine **7**, this resulted in a two-fold increase in labeling kinetics in vitro and a 6-fold increase in labeling efficiency in live cells (Figure 1C). The fluoro-(**8**) and chloro-(**9**) analogs of **7**, which in principle should be better leaving groups, were unreactive. Next, we tested fluorination of the benzyl group, a derivatization that increased the labeling efficiency in the corresponding SNAP-tag2 substrate. Analog **11** showed a 1.16-fold increased labeling kinetics in comparison to BC-TMR, and a 1.9-fold higher fluorescence signal in live cells (Figure 1C). Substrate **11** did not show a significant increase in labeling efficiency with SNAP-tag or *O*^6^-alkylguanine-DNA alkyltransferase (hAGT) (Figure S2), which is important to maintain labeling specificity for CLIP-tag. Combination of the 4-amino pyridine leaving group with the fluorinated benzyl group resulted in **12** (in the following abbreviated as PF-TMR), which showed a 1.48-fold increase in labeling kinetics, and a nearly 30-fold higher labeling efficiency in live cells (Figure 1C). The improved performance of PF-TMR allowed detectable CLIP-tag labeling in live cells at 100 nM after 2 hours, while labelling with BC-TMR showed no detectable signal under identical conditions (Figure 1D). The reaction of PF-TMR with CLIP-tag showed a *k*_*app*_ of 8.3 × 10^4^ M^-1^ s^-1^ (Figure 1E, Table S5). The optimized PF substrates conjugated to various fluorophore probes also did not significantly affect cell viability, even at higher concentrations than those used for labeling (Figure S3).

In light of the efficient live-cell labeling of CLIP-tag with PF-TMR, we chose this probe as substrate for the further engineering of CLIP-tag. We believe an important feature of PF as CLIP-tag substrate is the pK_a_ value of the protonated pyridine nitrogen, calculated to be around 7.3 (calculated with Schrödinger; see SI section 3.10). As a consequence, PF is partially protonated at physiological pH, increasing its intrinsic reactivity in the labeling reaction. The importance of protonation for the reactivity of the pyridine substrates might be the reason why the introduction of electron-withdrawing fluoro- and chloro-substituents at C6 of the 4-amino pyridine scaffold in PF results in reduced reactivity (Figure 1B): the pK_a_ values of **8** and **9** were calculated to be 2.4 and 3.4, respectively (see section 3.10 in SI), suggesting that only a small fraction of these compounds is protonated at physiological pH. Furthermore, the protonation of PF at physiological pH should increase the solubility of PF derivatives, while the presence of the unprotonated form of PF should facilitate their cellular uptake. The dynamic equilibrium between a charged and uncharged form has also been shown to be important for cellular uptake of rhodamine-based probes.^26^

To increase the labeling speed of CLIP-tag with PF-TMR, we first introduced the eight CLIP-tag-specific substitutions (M60I, Y114E, A121V, K131N, S135D, L153S, G157P, and F159L) into SNAP-tag2, aiming to preserve orthogonality and increase labeling speed.^19,21,24^ This resulted in only a modest 2.7-fold improvement in labeling kinetics in comparison to the original CLIP-tag (Table S2). Next, we used an alanine scanning of residues proximal to the active site of CLIP-tag.^27^ This resulted in the identification of two mutations, L159A and G160A, that in combination led to a 10-fold improvement in labeling kinetics (Table S2). To systematically probe the contribution of individual residues in CLIP-tag, we then screened a synthetic deep mutational scanning library (sDMSL), in which each residue in CLIP-tag was individually replaced by all other 19 amino acids, excluding cysteine but including deletions.^19,28^ Screening of this library for mutants that exhibited an increased labeling rate with PF-TMR was performed via yeast surface display (YSD) and followed by fluorescence activated cell sorting (FACS) (Figure S4).^29^ In parallel, we tested the effect of several mutations previously shown to increase the reactivity of SNAP-tag2 and combined beneficial mutations identified in both screens.^19^ Furthermore, we replaced the unstructured loop between residues 37 and 54 by a computationally designed loop (^37^GQGEQGPP^54^).^19,30,31^ This designed loop was previously shown to increase labeling speed in SNAP-tag2.^19^ Together, these changes resulted in CLIP-tag2, which carries 15 amino acid substitutions and a redesigned loop (Figure 2A and B). The *k*_*app*_ for the reaction of CLIP-tag2 with PF-TMR (Figure 2C) was determined to be 1.4 × 10^7^ M^-1^ s^-1^ (Figure 2D, Figure S5 and Table S3). This represents an ∼1000-fold improvement over the reaction rate of CLIP-tag with BC-TMR, and is comparable to the reaction rates of HaloTag7 and SNAP-tag2 with their respective TMR substrates (Figure 2E, 1.9 × 10^7^ and 8.2 × 10^6^ M^-1^ s^-1^, respectively).^18,19^ Moreover, CLIP-tag2 maintains high substrate specificity, reacting ∼1000-fold slower with TF-TMR and ∼200-fold slower with CF-TMR (Figure 2E and Table S2), which should allow the specific and simultaneous labeling of SNAP-tag2 and CLIP-tag2. CLIP-tag2 (173 residues, 18.7 kDa) has about the same size as SNAP-tag2, but is smaller than HaloTag7 (297 residues, 33.7 kDa). Furthermore, CLIP-tag2 possesses a relatively high thermal stability with a melting temperature (T_m_) of ∼58 °C (Figure S7).

**Figure 2.**
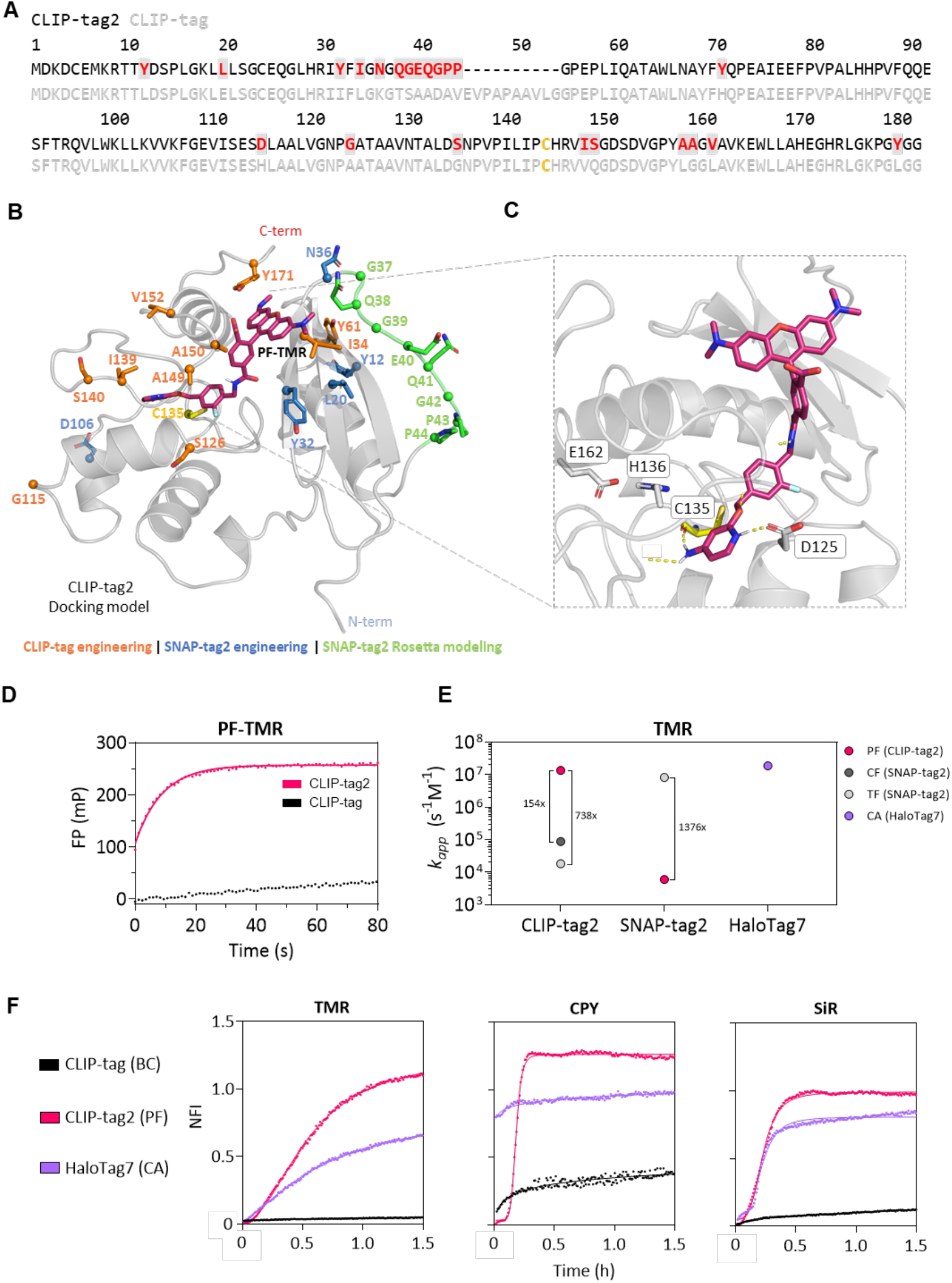
CLIP-tag2 engineering: (**A**) Amino acid sequence alignment of CLIP-tag2 (black) and CLIP-tag (grey). The CLIP-tag sequence is shown with the E30R mutation (i.e. CLIP_f_-tag).^24^ Amino acids changed during the course of engineering are highlighted in red, with active cysteine residue in yellow. (**B**) Docking model (Maestro Schrödinger) of CLIP-tag2 highlighting substituted residues in comparison to CLIP-tag sequence in (**A**). Amino acid residue coloring is based on the legend indicated below the sequence. N and C terminus are highlighted in blue and red, respectively. Active cysteine residue is highlighted in yellow. (**C**) Zoom-in into the active site of CLIP-tag2 based on the model shown in (**B**), showing interactions of protonated PF-TMR with D125. (**D**) In vitro labeling kinetics of CLIP-tag2 and CLIP-tag (with E30R mutation), with PF-TMR (4 nM), measured by fluorescence polarization at 10 nM protein concentration. Data was fitted to one-phase association model and are presented as the mean of triplicates ± 95% CI. (**E**) Comparison of labeling kinetics (*k*_app_) between CLIP-tag2, SNAP-tag2 and HaloTag7 with different TMR substrates (see Table S2 and S3). SNAP-tag2 reaction rates with TF-TMR and HaloTag7 reaction rates with CA-TMR were obtained previously.^18,19^ (**F**) Comparison of labeling kinetics between CLIP-tag2, CLIP-tag and HaloTag7 with their respective TMR (50 nM), CPY (50 nM) and SiR (100 nM) substrates in live U2OS cells. Cells stably co-express HaloTag7-CLIP-tag or HaloTag7-CLIP-tag2 fusion constructs in the nucleus, with mTurquoise2 as an expression marker. Labeling reactions were followed by confocal fluorescence microscopy and fluorescence intensity changes were normalized to the mTurquoise2 signal over time. The data was fitted to the sigmoidal curve model (97 ≥ *n* (TMR) ≥ 11 cells, 57 ≥ *n* (CPY) ≥ 19 cells, 46 ≥ *n* (SiR) ≥ 19 cells). Biological duplicate is shown in Figure S9. Calculated half-times (t_1/2_) of CLIP-tag2, CLIP-tag and HaloTag7 are shown in Table S4.

To visualize the position of the individual mutations in CLIP-tag2 relative to its substrate, we used AlphaFold2 to predict its structure and docked the PF-TMR substrate into its active site (Figure 2C and Figure S8, SI section 3.10). While further studies would be required to understand the role of individual mutations in increasing the reactivity of CLIP-tag2, the docking suggests that Asp125, which is also present in parental CLIP-tag, can interact with the proto-nated pyridine through a salt bridge (Figure 2C, and Figure S8A and B). This interaction is not possible for the protonated 6-amino pyridine derivative **10** (Figure S8C), providing a possible explanation why this PF isomer shows no significant protein labeling with CLIP-tag2.

We next measured the labeling kinetics of CLIP-tag2, CLIP-tag and HaloTag7 with their respective substrates in live cells (Figure 2F and Figure S9). U2OS cells stably expressing a nuclear localized HaloTag7-CLIP-tag2 or HaloTag7-CLIP-tag fusion proteins were incubated with low concentrations (50 nM or 100 nM) of their corresponding TMR, CPY and SiR sub-strates. The labeling kinetics were measured with confocal microscopy by recording the increase in nuclear fluorescence intensity over time. CLIP-tag2 showed rapid labeling with the three fluorophores tested, with t_1/2_ values (half maximum labeling time) below 20 minutes. In comparison, the labeling of parental CLIP-tag with BC derivatives was much slower and no saturation of the fluorescence signal was achieved within the 1.5-hour measurement period. The labeling speed of CLIP-tag2 with PF-SiR was comparable to the labeling of HaloTag7 with CA-SiR. For CPY, HaloTag7 labeling was faster than CLIP-tag2, while CLIP-tag2 labeling with PF-TMR was faster than labeling of HaloTag7 with CA-TMR. CLIP-tag2 labeling resulted in slightly higher fluorescence signals than HaloTag7 labeling with all three fluorophores (Figure S10). Furthermore, we previously measured the labeling kinetics of SNAP-tag2 using the same setup and in comparison, CLIP-tag2 showed an about two-fold faster labeling than SNAP-tag2 with CPY and SiR, and equivalent labeling speed with TMR.^19^

We then tested the fluorescence labeling of CLIP-tag2 fusion proteins in live U2OS cells (Figure 3A). CLIP-tag2 was targeted to various localizations via direct fusion to different localization sequences (H2B, LifeAct, CPE41, TOMM20) and could be efficiently and specifically labeled with PF coupled to TMR, MaP555, CPY and SiR at low substrate concentrations (100 nM) within 1.5 hours. To demonstrate the possibility for multiplex imaging of CLIP-tag2 with SNAP-tag2 and HaloTag, we simultaneously labeled CLIP-tag2, SNAP-tag2 and HaloTag9 in live U2OS cells. Specifically, cells expressing CLIP-tag2 localized to F-actin (LifeAct-CLIP-tag2), SNAP-tag2 localized to the plasma membrane (Lyn11-SNAP-tag2) and HaloTag localized to the endoplasmic reticulum (CalR-HaloTag9-KDEL)^32^ were simultaneously incubated with PF-MaP555, SiR-TF and CA-500R^33^ (200 nM each) for 30 min. Subsequent confocal laser scanning microscopy confirmed the specific labeling of each of the proteins (Figure 3B). This experiment demonstrates the possibility to perform three-color multiplexed live-cell imaging using SLPs with very similar, outstanding labeling efficiencies. Furthermore, CLIP-tag2 is compatible with live-cell STED microscopy (Figure S11).

**Figure 3.**
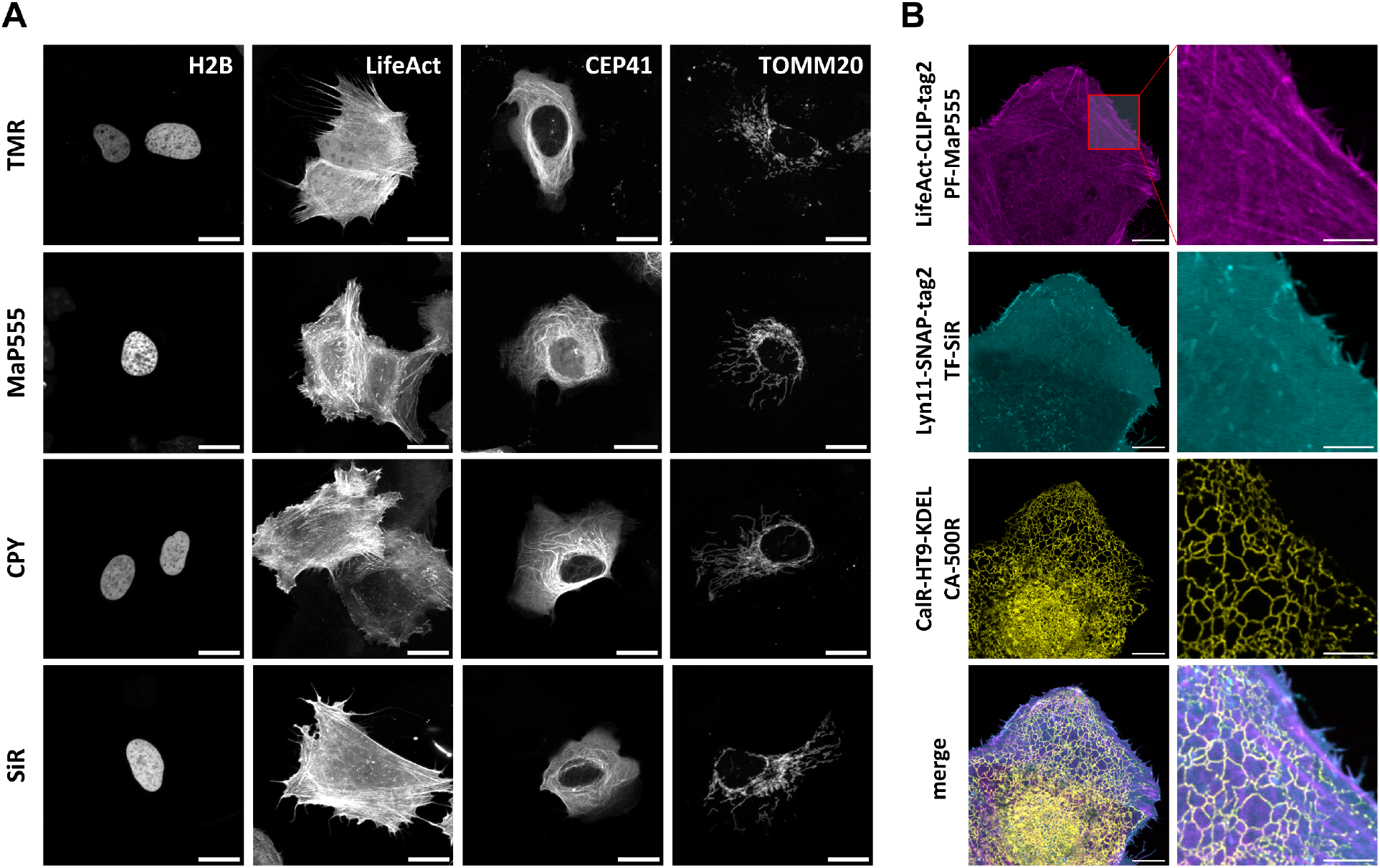
CLIP-tag2 labeling in live cells: (**A**) CLIP-tag2 labeling in U2OS cells transiently expressing CLIP-tag2 fusion constructs. Cells were incubated with 100 nM of PF-TMR/MaP555/CPY/SiR substrates for 1.5 hour and washed three times with imaging medium and subsequently imaged in the same medium. Maximum projections are shown. Pixel intensities were scaled for each fluorophore and/or sub-cellular localization. Scale bar 20 μm. (**B**) CLSM of CLIP-tag2, SNAP-tag2 and HaloTag9 expressed in U2OS cells via rAAVs. Cells were incubated with 200 nM of PF-MaP555 (CLIP-tag2), TF-SiR (SNAP-tag2) and CA-500R (HaloTag9) for 30 min, after which the cells were washed twice with imaging medium and imaged. Scale bar: 10 µm (overview) and 5 µm (magnification). The SLPs were fused to the following marker proteins: nucleus – H2B, actin – LifeAct, microtubules – CEP41, outer mitochondrial membrane – TOMM20, inner plasma membrane – Lyn11, calreticulin – CalR, KDEL – ER retention signal.

In summary, we introduce CLIP-tag2 and its labeling substrate PF for the efficient and rapid labeling of fusion proteins in live cells. The labeling kinetics and specificity of the system make it a powerful tool for applications in live-cell (super-resolution) microscopy, multiplexed imaging strategies and an attractive starting point for the development of chemogenetic biosensors.

## Methods

Detailed methods and supplementary materials are provided in the Supplementary Information.

## Data availability

The data supporting the findings of this study are provided within the Supplementary Information. A plasmid of CLIP-tag2 will be available on Addgene.

## Acknowledgement

This work was supported by the Max Planck Society and École Polytechnique Fédérale de Lausanne (EPFL). AP and JW were supported by the Max Planck School Matter to Life. We thank S. Fabritz, T. Rudi, and J. Kling from the mass spectrometry facility of MPIMR for their support. We thank Elisa D’Este (optical nanoscopy facility, MPIMR) for support and Jochen Reinstein (MPIMR) for help with the stopped-flow kinetics. The authors thank A. Bergner, J. Kress and P. Breuer for providing materials and support.

