## Supporting Information for "Fast and Luminous: CLIP-tag2"

### Table of contents


1 Supplementary figures ... 5

2 Supplementary tables ... 16

3 Methods ... 22

3.1 Molecular cloning and protein generation ... 22

3.2 Measurement of protein stability ... 23

3.3 Measurement of labeling kinetics ... 23

3.3.1 Measurement of labeling rates using multi-well reader at fixed protein concentrations ... 23

3.3.2 Measurement of labeling rates using multi-well plate reader at varying protein concentrations ... 24

3.3.3 Measurement of CLIP-tag2 kinetics using stopped-flow device ... 24

3.4 Yeast surface display ... 25

3.4.1 Synthetic deep mutational scanning library (sDMSL) ... 26

3.4.2 Electrocompetent cell preparation ... 26

3.4.3 Homologous recombination ... 26

3.4.4 Protein expression ... 27

3.4.5 Protein labelling ... 27

3.4.6 Fluorescence activated cell sorting ... 27

3.4.7 Next generation sequencing and data analysis ... 28

3.4.8 Data analysis ... 28

3.5 Generation of mammalian stable U2OS CLIP-tag2 cell lines ... 28

3.6 Transient transfection of mammalian cells ... 29

3.7 Preparation of rAAVs ... 29

3.8 Microscopy ... 29

3.9 Cell viability assay ... 32

3.10 Maestro Schrödinger and Boltz2 calculations ... 32

3.11 Important protein sequences ... 33

3.11.1 Sequences for bacterial expression ... 33

3.11.2 Sequences for expression on yeast surface ... 33

3.11.3 Sequences for expression in mammalian cells ... 34

3.12 Scripts ... 36

3.12.1 NGS analysis of libraries ... 36
3.12.2 Analysis of labelling kinetics obtained in multi-well assays ... 37
3.12.3 Analysis of stopped-flow labelling kinetics ... 39
4 Chemical Synthesis of CLIP-tag2 Substrates ... 40
General Remarks ... 40
4.1 Synthesized molecules ... 41
4.1.1 Building blocks – nucleobases ... 41
4.1.2 Building blocks – benzyl alcohol ... 48
4.1.3 Conjugate precursors ... 48
4.1.4 Fluorophore conjugates ... 56
5 References ... 62
6 NMR Spectra ... 64

### 1 Supplementary figures

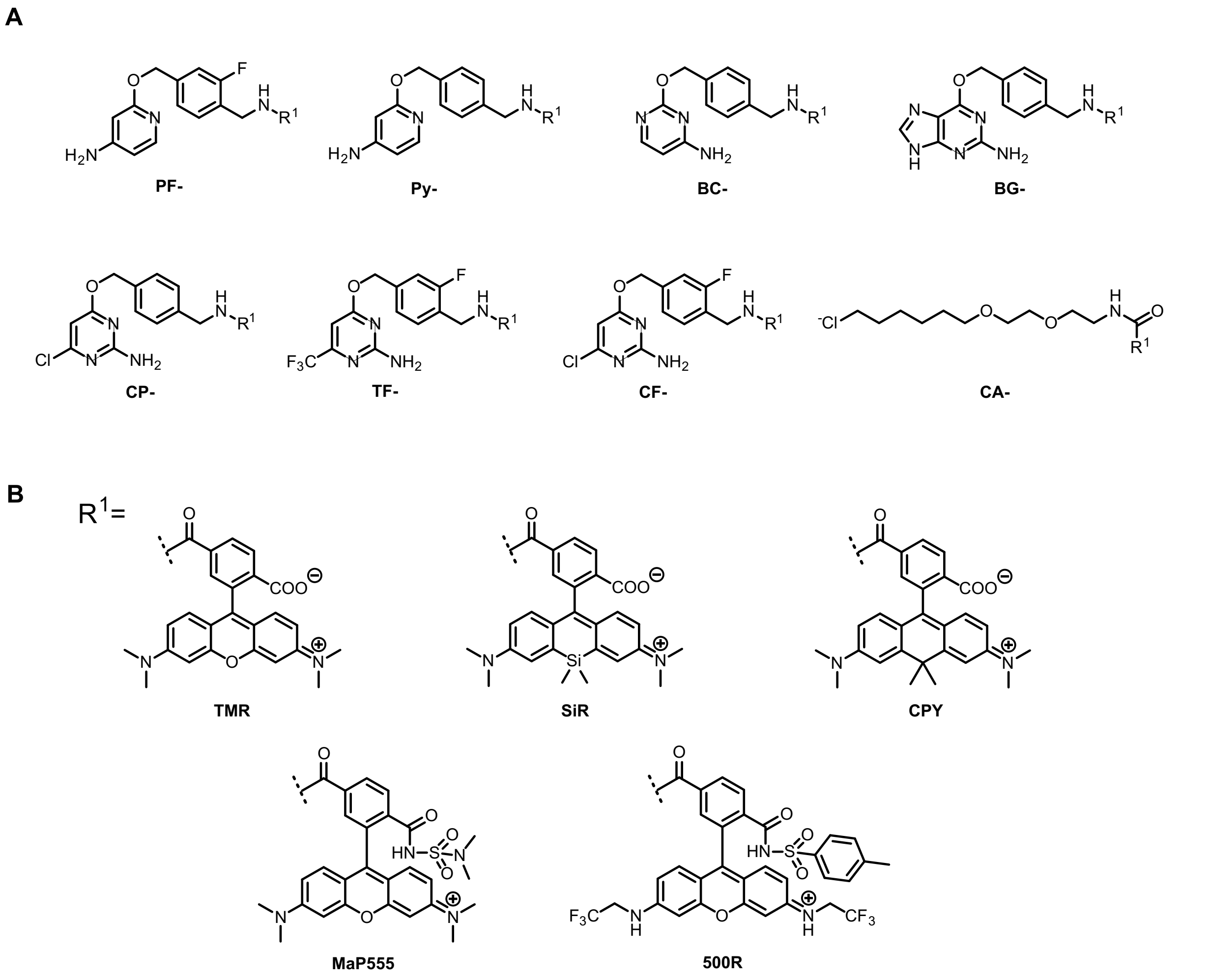


**Figure S1 Structures of SLP substrates used in this study.**
(**A**) Structures of CLIP-tag (PF, Py and BC), SNAP-tag (BG, CP, TF and CF) and HaloTag (CA) substrate cores.
(**B**) Structures of $R^1$ groups (fluorophores) attached to substrate cores shown in (**A**).[1-3]

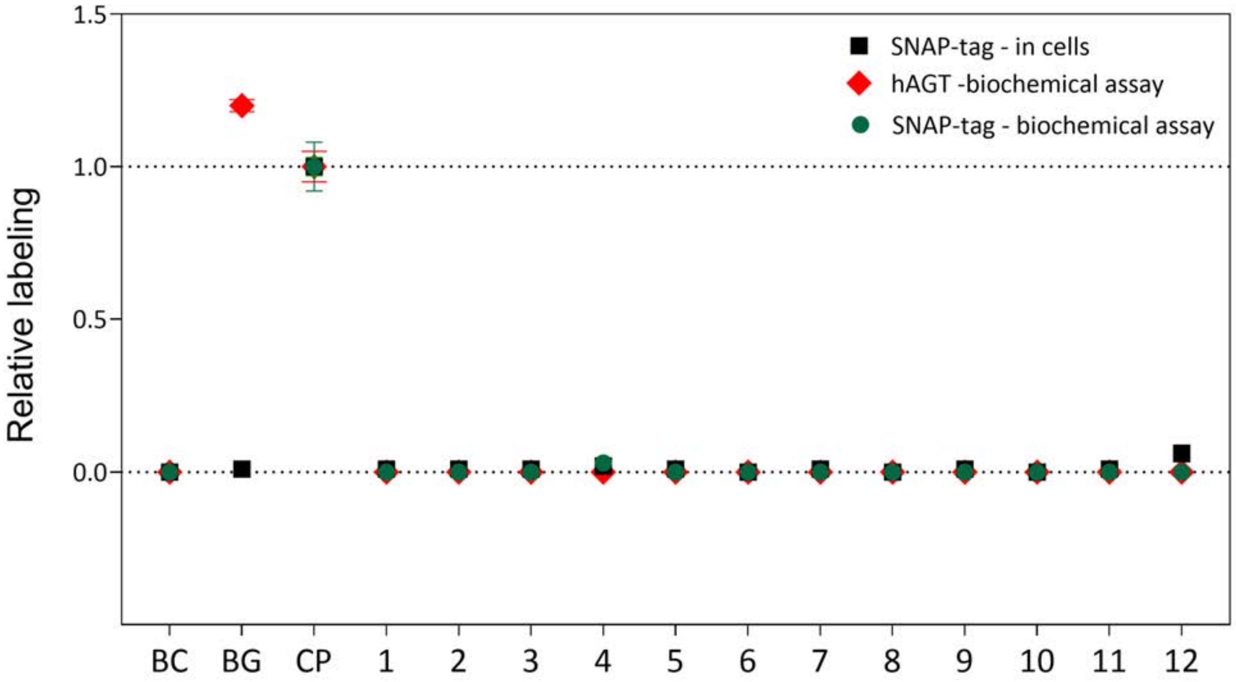


**Figure S2 Reactivity of CLIP-tag substrates with hAGT and SNAP-tag.**

Reactivity of CLIP-tag substrates (**1-12**) with hAGT, SNAP-tag (in cells) and $SNAP_f$-tag (SNAP-tag with E30R mutation,[4] in a biochemical assay) normalized to SNAP-tag/$SNAP_f$-tag labeling of CP-TMR. In a biochemical assay, labeling kinetics of $SNAP_f$-tag with new substrates were measured by recording fluorescence polarization traces over time. Apparent second-order rate constants ($k_{app}$) were calculated (Table S1) and normalized to the $k_{app}$ of CP-TMR in $SNAP_f$-tag labeling. Live-cell performance of new substrates was tested by labeling of U2OS cells stably expressing mEGFP-SNAP-tag or mEGFP-hAGT fusion protein with TMR-substrates at 100 nM for 2 h. Cells were washed 2 times with imaging medium and analyzed *via* flow cytometry. Fluorescence intensity ratios of TMR/mEGFP were calculated (Table S1) and normalized to the ratio obtained for SNAP-tag with CP-TMR. Bottom graph shows the zoom-in of the data shown on the left. Experiments were performed using exact experimental conditions like we described before.[2]

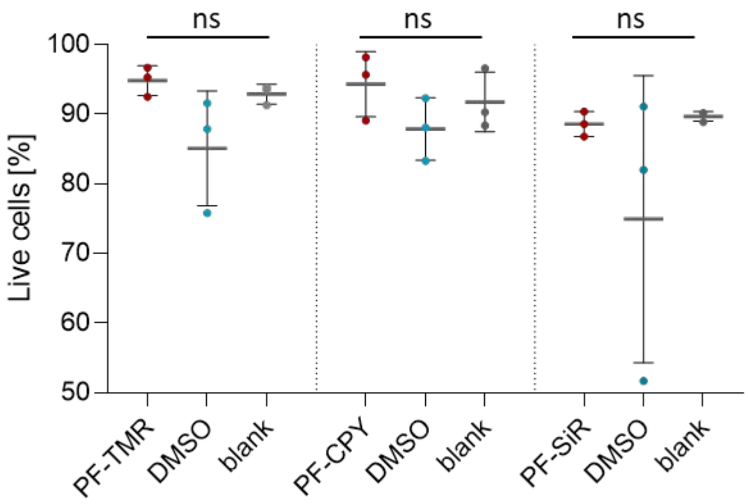


**Figure S3 Statistical analysis of cell viability treated with PF-TMR/CPY/SiR substrates.**
Dot plot graph showing the percentage of live cell events for each PF substrate in comparison to DMSO and untreated control samples. U2OS cells were incubates either with PF substrates (1 µM), DMSO (1% v/v) or remained untreated. Cells were incubated for 1 h at 37 °C, after which both dead cell (supernatant) and live cell (detached with trypsin) populations were collected and combined. Cells were then stained with SYTOX Blue (1 µM) and were analyzed using flow cytometry, in technical triplicates represented by dot plot graph. Black horizontal lines represent the mean percentage of live cells from each technical triplicate and the error bars represent standard deviation. The differences in cell viability between treated and untreated cells were statistically not significant based on two-tailed unpaired t-test including Welch's correction ($p > 0.05$, ns), indicating that under these conditions the PF substrates do not have a strong impact on cell viability.

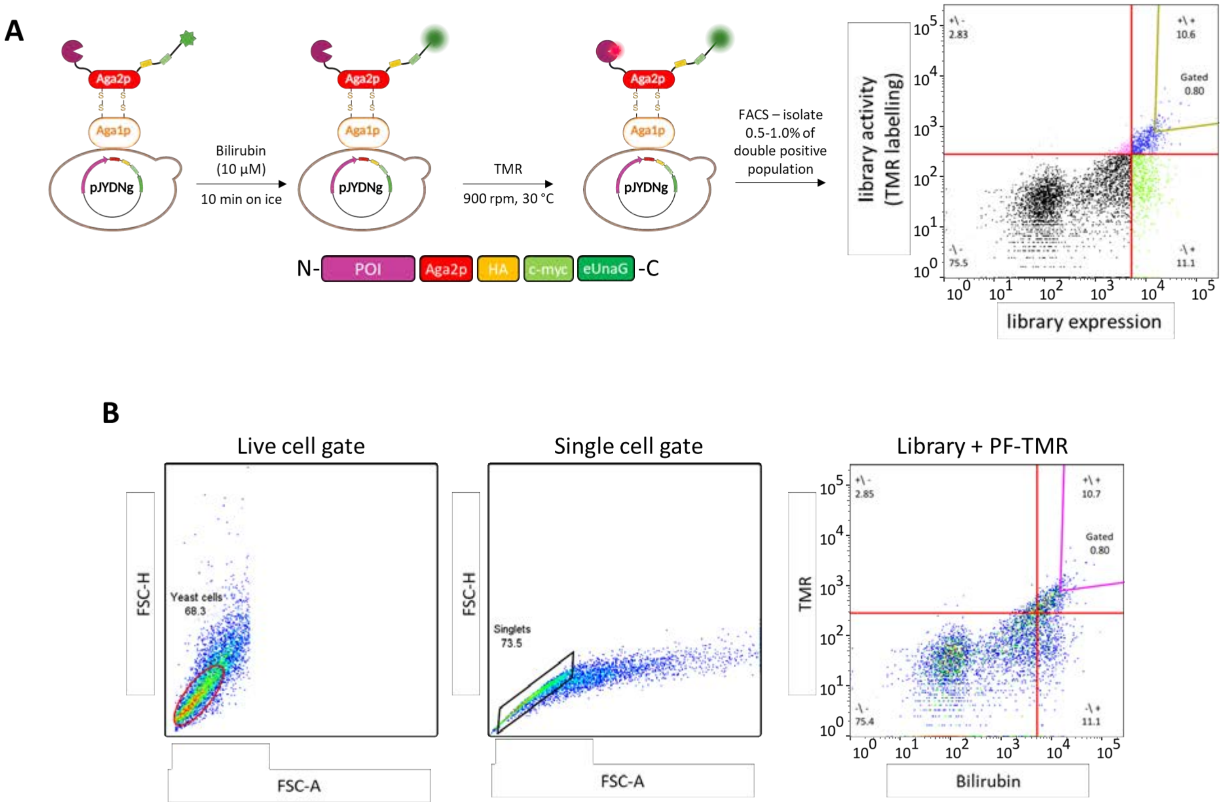


**Figure S4 Yeast surface display screening.**
(**A**) Outline of screening strategy in yeast surface display with a pJYDNg construct,[5] using a CLIP$_f$-tag (CLIP-tag with E30R mutation)[4] sDMSL (TWIST) library. Cells were initially labelled with expression marker bilirubin (10 µM), followed by incubation with a fluorescence marker PF-TMR at 5 or 3.5 nM concentrations (Table S6). Cells within the gated double positive population were isolated, cultured, and used for subsequent rounds of sorting.
(**B**) Gating strategy employed during yeast surface display. Cells were gated for live and single cell variants (singlets) and subsequently gated for double positive (TMR + bilirubin) labelling at the top 0.5 – 1.0 %.

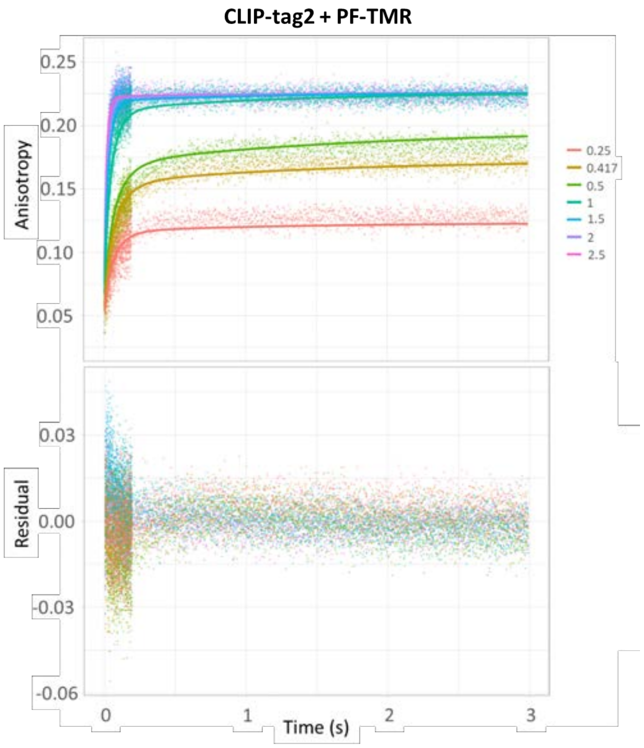


**Figure S5 *In vitro* labeling kinetics of CLIP-tag2 with PF-TMR determined *via* stopped-flow.**
Top graph represents full anisotropy traces (points) and predications of fits (lines), based on the model described in equations 7-14. Bottom graph shows residual from the fit. The PF-TMR concentration was fixed at 0.5 μM, while protein concentration was varied from 0.25 to 2.5 μM. Kinetic parameters derived from the fits are shown in Table S3.

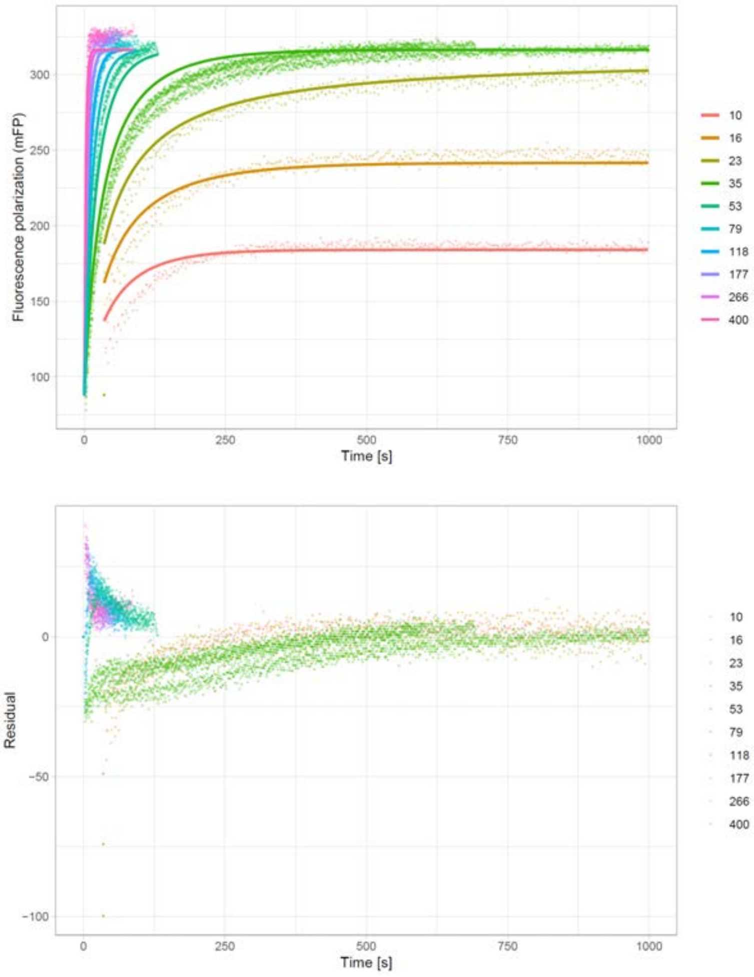


**Figure S6 *In vitro* labeling kinetics using fluorescence polarization at varying protein concentrations.**
A representative graph of *in vitro* labeling kinetics of $\text{CLIP}_f$-tag carrying four amino acid substitutions (I32Y, L34I, L159A and G160A) and PF-TMR using fluorescence polarization. Top graph represents full fluorescence polarization traces (points) and predications of fits (lines), based on the model described in equations 3-6, with bottom graphs showing residual from the fits. The PF-TMR concentration was fixed at 20 nM, while protein concentration was varied from 10 to 400 nM.

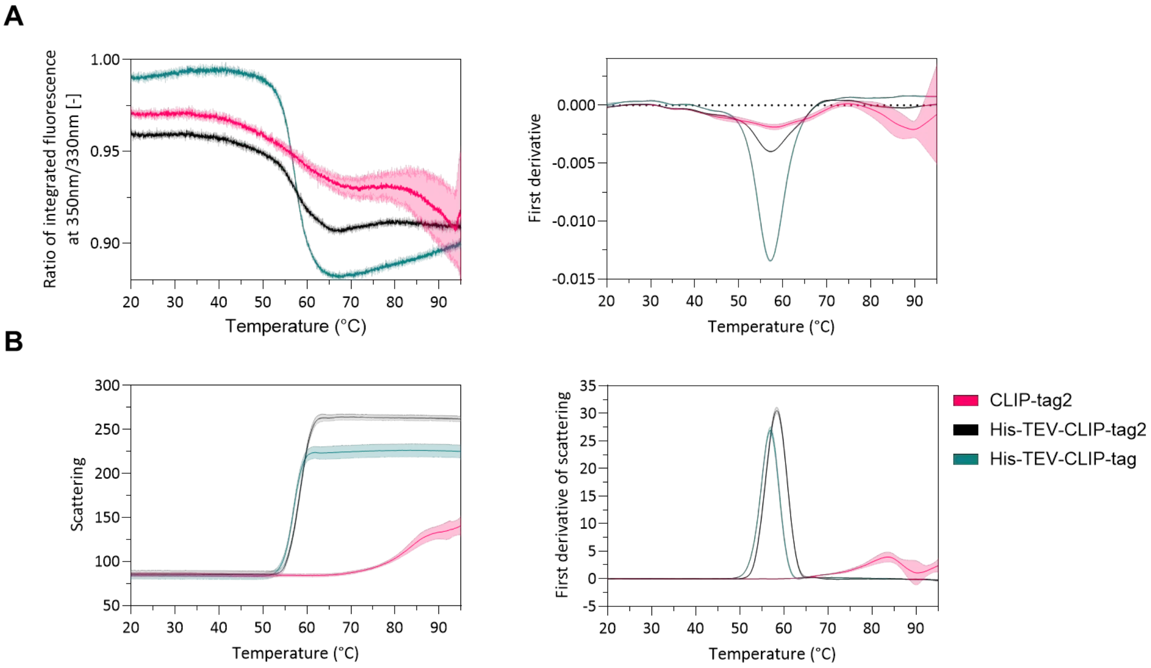


**Figure S7 Thermal stability measurements of CLIP-tag2, His-TEV-CLIP-tag2 and His-TEV-CLIP-tag using NanoDSF.**

(**A**) Ratio of integrated fluorescence at 350nm/330nm with its corresponding first derivative at increasing temperature from 20 to 95 °C. Ratio at 350nm/330n, is similar between the three proteins, with inflection points at 58.0, 57.3 and 57.2 °C for CLIP-tag2, His-TEV-CLIP-tag2 and His-TEV-CLIP-tag (CLIP-tag with E30 mutation termed $CLIP_f$-tag), respectively.

(**B**) Scattering plot with its corresponding first derivative. Without His-tag, CLIP-tag2 shows no sharp transition, indicating it is less prone to aggregation. All measurements were performed in technical triplicates.

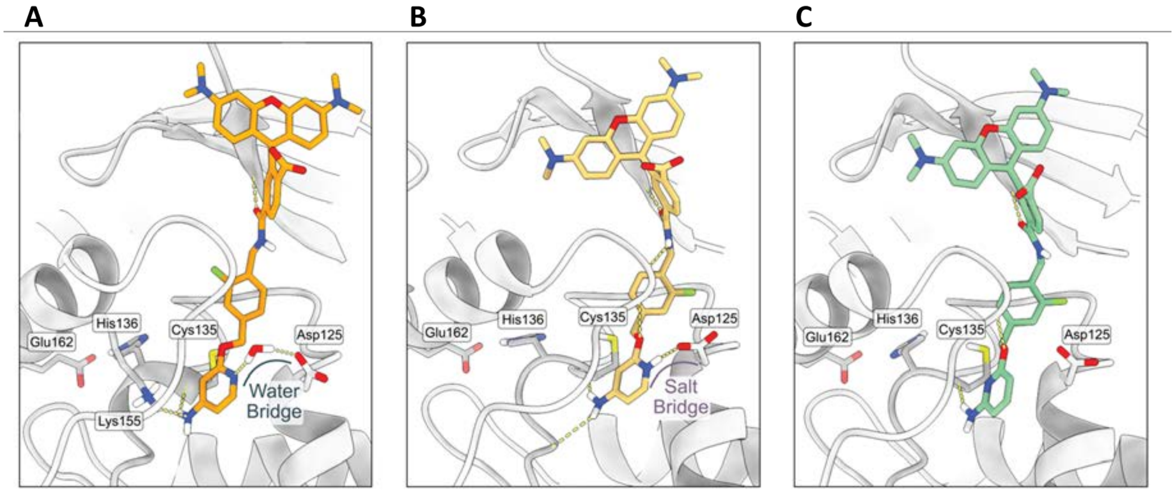


**Figure S8 Molecular dynamics simulation of substrates in CLIP-tag2 active site using Maestro Schrödinger.**
(**A**) Compound **12** (PF-TMR) modeled into the active site of CLIP-tag2. Pyridine nitrogen is oriented towards Asp125, which could enable formation of the water bridge.
(**B**) Protonated compound **12** (PF-TMR-H$^+$) modeled into the active site of CLIP-tag2. Pyridine nitrogen of the substrate is oriented towards Asp125, which could enable formation of the salt bridge.
(**C**) Compound **10** modeled into the active site of CLIP-tag2. Due to the placement of the amino group in the pyridine ring, pyridine nitrogen is not able to interact with Asp125.

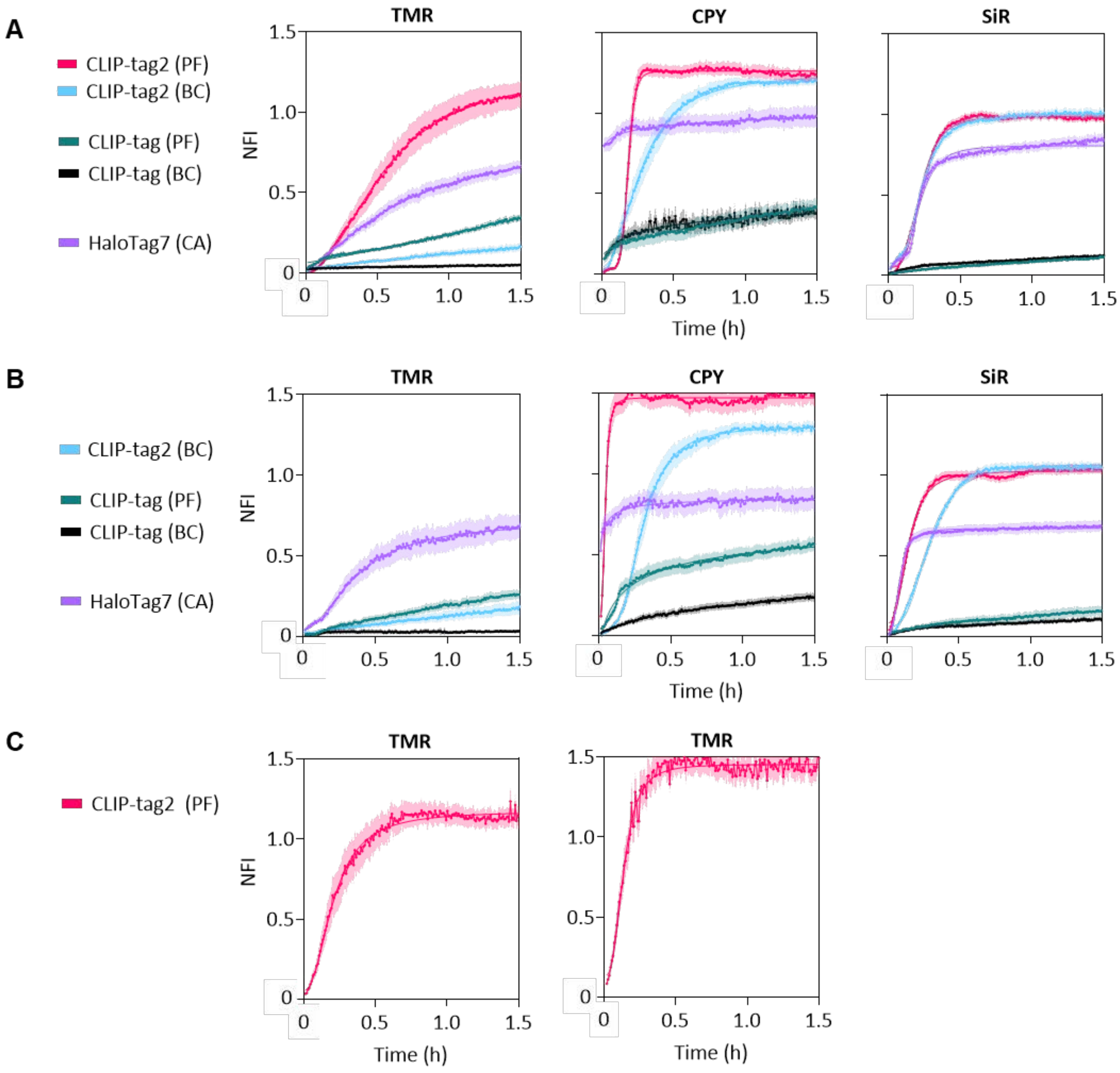


**Figure S9 Labelling kinetics of CLIP-tag2, CLIP-tag and HaloTag7 in live cells.**
(**A**, **B**) Kinetic traces of CLIP-tag2, CLIP-tag (CLIP-tag with E30R mutation termed $CLIP_f$-tag) and HaloTag7 from two biological replicates in live-cell fluorescent labeling with their respective TMR (50 nM), CPY (50 nM) and SiR (100 nM) substrates. Labeling of CLIP-tag2 with PF-TMR in (**B**) was performed separately as two technical replicates shown in (**C**). Cells stably co-express HaloTag7-$CLIP_f$-tag or HaloTag7-CLIP-tag2 fusion constructs in the nucleus, with mTurquoise2 as an expression marker. Labelling reactions were followed by confocal fluorescence microscopy and fluorescence intensity changes were normalized to the mTorquoise2 signal over time. The data was fitted to the sigmoidal curve model (97 ≥ *n* (TMR) ≥ 11 cells, 57 ≥ *n* (CPY) ≥ 19 cells, 46 ≥ *n* (SiR) ≥ 19 cells). Calculated half-times ($t_{1/2}$) of CLIP-tag2, $CLIP_f$-tag and HaloTag7 are shown in Table S4, and represent the mean ± 95% CI from the data obtained in (**A**), (**B**) and (**C**). NFI = normalized fluorescence intensity.

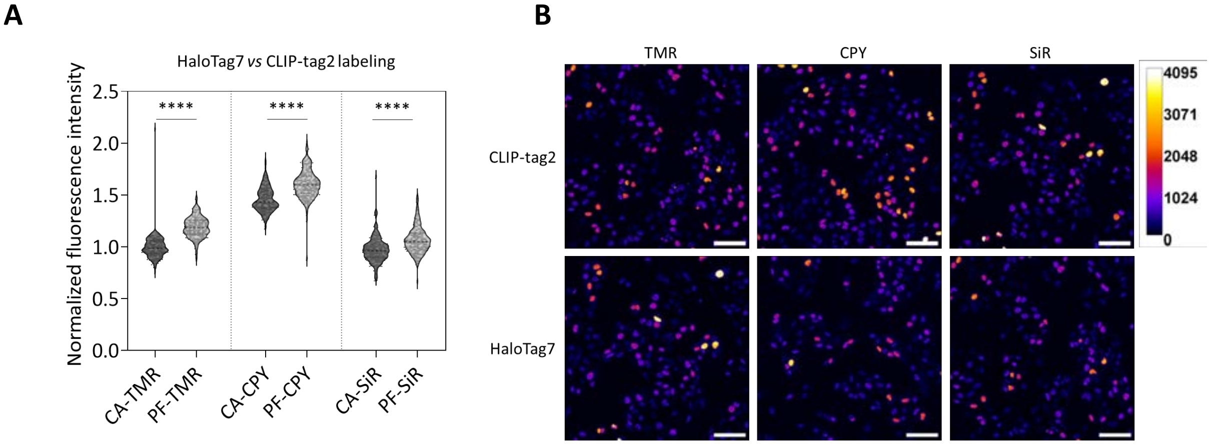


**Figure S10 Comparison of fluorescence brightness of CLIP-tag2 and HaloTag7 in live cells.**
(**A**) Comparison of CLIP-tag2 and HaloTag7 fluorescence brightness in live U2OS cells stably co-expressing HaloTag7-CLIPtag2 fusion with the mTorquoise2 as an expression marker in the nucleus. Cells were labelled with CA-/PF- TMR, CPY and SiR substrates (500 nM) for 2 hours and washed with imaging medium before imaging with confocal fluorescence microscopy. Statistical significance was assessed with a two-tailed unpaired t-test including Welch's correction ($P < 0.0001$).
(**B**) Representative confocal fluorescence images corresponding to the data in (**A**), shown in the substrate channel. Scale bar 100 µm.

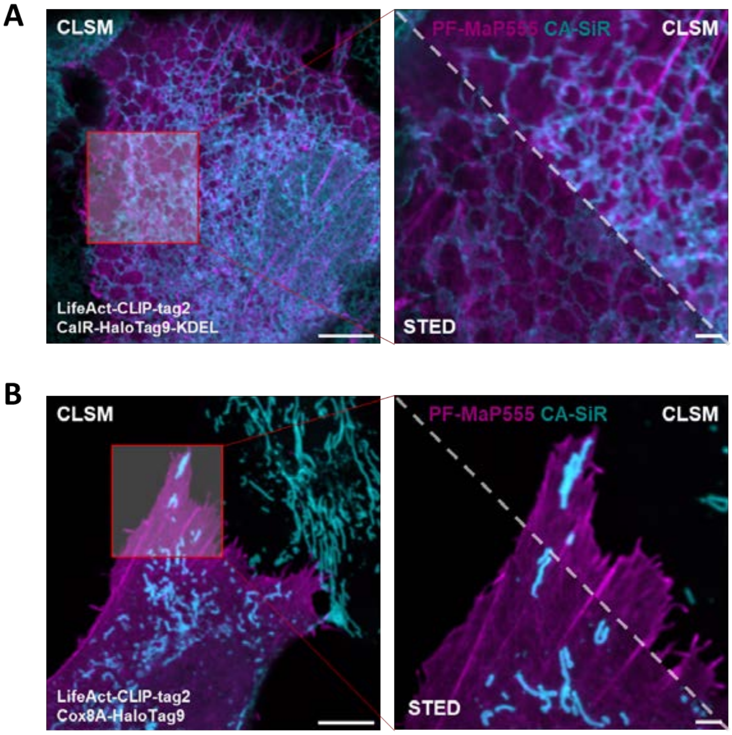


**Figure S11 Dual-color confocal and STED imaging with CLIP-tag2 and HaloTag9.**
Confocal laser scanning microscopy (CLSM) and stimulated emission depletion (STED) imaging of CLIP-tag2 and HaloTag9 expressed in U2OS cells via rAAVs. Cells were incubated with 200 nM of PF-MaP55 (CLIP-tag2) and CA-SiR (HaloTag9) for 30 min, after which the cells were washed twice with imaging medium and imaged. The SLPs were fused to the following marker proteins: actin – LifeAct, calreticulin – CalR, mitochondrial matrix – Cox8A localization sequence. Scale bar: 10 µm (overview) and 2 µm (magnification).

#### 2 Supplementary tables

**Table S1 Measurement of labelling efficiencies for reaction of SLPs with new substrates.**

| | Compound | SNAP$_f$-tag# | SNAP-tag | CLIP$_f$-tag# | CLIP-tag | hAGT |
|---|---|---|---|---|---|---|
| | | *In vitro* | In cells | *In vitro* | In cells | *In vitro* |
| | | $k_{app}$ (± s.d.) × $10^4$ [$M^{-1}$ $s^{-1}$] | Fluo. Int. Ratio TMR/GFP | $k_{app}$ (± s.d.) × $10^4$ [$M^{-1}$ $s^{-1}$] | Fluo. Int. Ratio TMR/GFP | $k_{app}$ (± s.d.) × $10^3$ [$M^{-1}$ $s^{-1}$] |
| Leaving group modifications | BC-TMR | n.d. | 0.003 ± 0.001 | 1.13 ± 0.02 | 0.038 ± 0.001 | n.d. |
| | BG-TMR | 34.9 ± 3.43 | 0.005 ± 0.001 | 0.02 ± 0.01 | 0.001 ± 0.001 | 0.97 ± 0.02 |
| | CP-TMR | 6.17 ± 0.50 | 0.898 ± 0.024 | n.d. | 0.003 ± 0.001 | 0.80 ± 0.04 |
| | 1 | n.r. | 0.008 ± 0.000 | 0.01 ± 0.00 | 0.001 ± 0.000 | n.r. |
| | 2 | n.r. | 0.009 ± 0.000 | 1.17 ± 0.00 | 0.016 ± 0.001 | n.r. |
| | 3 | n.r. | 0.006 ± 0.000 | n.r. | 0.001 ± 0.000 | n.r. |
| | 4 | 0.16 ± 0.02 | 0.017 ± 0.001 | 0.09 ± 0.00 | 0.008 ± 0.000 | n.r. |
| | 5 | n.r. | 0.010 ± 0.000 | 0.45 ± 0.00 | 0.002 ± 0.000 | n.r. |
| | 6 | n.r. | 0.003 ± 0.000 | 0.06 ± 0.00 | 0.009 ± 0.000 | n.r. |
| | 7 | n.r. | 0.012 ± 0.002 | 1.50 ± 0.52 | 0.236 ± 0.003 | n.r. |
| | 8 | n.r. | 0.001 ± 0.000 | n.r. | n.r. | n.r. |
| | 9 | n.r. | 0.007 ± 0.000 | n.r. | 0.001 ± 0.000 | n.r. |
| | 10 | n.r. | 0.002 ± 0.000 | n.r. | 0.001 ± 0.000 | n.r. |
| Linker modifications | 11 | n.r. | 0.008 ± 0.000 | 1.32 ± 0.11 | 0.074 ± 0.001 | n.r. |
| | 12 | n.r. | 0.051 ± 0.007 | 1.67 ± 0.10 | 1.075 ± 0.022 | n.r. |

$k_{app}$ = apparent second-order rate constant; n.r. = not reactive; s.d. = standard deviation; Fluo. Int. = fluorescence intensity. Experiment performed like previously described.[2]

#SNAP-tag and CLIP-tag with E30R mutation.

**Table S2 Rate constants for reaction of CLIP-tag2 and SNAP-tag2 with PF-TMR, TF-TMR and CF-TMR obtained *via* measurements in a multi-well plate reader.**

| Protein | Substrate | $k_{app}$ ± C.I. ($M^{-1}$ $s^{-1}$) |
|---|---|---|
| SNAP-tag2 | PF-TMR | (5.97 ± 0.03) × $10^3$ |
| SNAP-tag2 + M60I-Y114E-A121V-K131N-S135D-L153S-G157P-F159L | | (1.20 ± 0.04) × $10^5$ |
| CLIP-tag2 | TF-TMR | (1.83 ± 0.05) × $10^4$ |
| | CF-TMR | (8.79 ± 0.68) × $10^4$ |
| $CLIP_f$-tag# + L159A-G160A | PF-TMR | (4.20 ± 0.17) × $10^5$ |

$k_{app}$ = apparent second-order rate constant

#$CLIP_f$-tag – CLIP-tag with E30R mutation.

*$k_{app}$ values were measured by multi-well plate reader at varying protein concentrations (± 95% CI) to assess the orthogonality to SNAP-tag2. Values represent the mean of triplicates.

**Table S3 Rate constants for reaction of CLIP-tag2 with PF-TMR obtained *via* stopped-flow measurements.**

| Kinetic parameter | $k_1$ | $k_{app}$ | $k_{-1}$ | $k_2$ | $K_D$ |
|---|---|---|---|---|---|
| **Substrate** | **(± s.d.) × $10^7$ [$M^{-1}$ $s^{-1}$]** | | **(± s.d.) [$s^{-1}$]** | | **(± s.d.) [nM]** |
| PF-TMR | 2.70 (± 0.02) | 1.35 (± 0.06) | 0.93 (± 0.05) | 0.92 (± 0.07) | 34.0 (± 1.65) |

$k_1$ = reversible binding constant; $k_{app}$ = apparent second-order rate constant; $k_{-1}$ = reversible unbinding constant; $k_2$ = irreversible covalent constant; $K_D$ = binding affinity constant.

**Table S4 Calculated half-times ($t_{1/2}$) of CLIP-tag2, $CLIP_f$-tag and HaloTag7 from in-cell kinetic measurement.**

| Protein | Substrate | $t_{1/2}$ (95% CI) [min] TMR | CPY | SiR |
|---|---|---|---|---|
| CLIP-tag2 | PF | 17.9 ± 0.9 | 6.8 ± 0.1 | 11.2 ± 0.1 |
| | BC | >> 90 | 17.2 ± 0.3 | 15.6 ± 0.2 |
| $CLIP_f$-tag | PF | >> 90 | >> 90 | >> 90 |
| | BC | >> 90 | >> 90 | >> 90 |
| HaloTag7 | CA | 31.3 ± 2.9 | << 2 | 9.1 ± 0.2 |

**Table S5 Next-generation sequencing (NGS) primers used for amplification of libraries after FACS screen.**

| Primer Name | Adapter Sequence | Primer Bind Sequence | Adapter Ligation Sequence |
|---|---|---|---|
| Fwd1a | CTGAGAAGCG | GCTGCTTCTTCTGCTTTGG | CTGAGAAGCGGCTGCTTCTTCTGCTTTGG |
| Rev1 | AACCGGAAAT | TCCTCGATAGC | AACCGGAAATTCCTCGATAGC |
| Fwd2a | CCAGCTTAGT | GCTGAATGCGTACTTCCATC | CCAGCTTAGTGCTGAATGCGTACTTCCATC |
| Fwd3a | TCCTATGCTC | ATCCTGATTCCATGTCATCGC | TCCTATGCTCATCCTGATTCCATGTCATCGC |
| Rev2 | ATATGGGCCA | ACATCGGAATC | ATATGGGCCAACATCGGAATC |
| Rev3 | GACAACGTTA | TCCAACAAGTTGATG | GACAACGTTATCCAACAAGTTGATG |
| Fwd1b | ATAGGCTGAC | GCTGCTTCTTCTGCTTTGG | ATAGGCTGACGCTGCTTCTTCTGCTTTGG |
| Fwd2b | AGCCAGCTCT | GCTGAATGCGTACTTCCATC | AGCCAGCTCTGCTGAATGCGTACTTCCATC |
| Fwd3b | GATGTCAACT | ATCCTGATTCCATGTCATCGC | GATGTCAACTATCCTGATTCCATGTCATCGC |
| Fwd1c | AGTAGGAGGA | GCTGCTTCTTCTGCTTTGG | AGTAGGAGGAGCTGCTTCTTCTGCTTTGG |
| Fwd2c | AGCGCATGGA | GCTGAATGCGTACTTCCATC | AGCGCATGGAGCTGAATGCGTACTTCCATC |
| Fwd3c | TCACGAGCGT | ATCCTGATTCCATGTCATCGC | TCACGAGCGTATCCTGATTCCATGTCATCGC |
| Fwd1d | CCGTACGATG | GCTGCTTCTTCTGCTTTGG | CCGTACGATGGCTGCTTCTTCTGCTTTGG |
| Fwd2d | ATTATCGGAC | GCTGAATGCGTACTTCCATC | ATTATCGGACGCTGAATGCGTACTTCCATC |
| Fwd3d | ACGTAGGCAC | ATCCTGATTCCATGTCATCGC | ACGTAGGCACATCCTGATTCCATGTCATCGC |
| Fwd1e | AGTCATTGAG | GCTGCTTCTTCTGCTTTGG | AGTCATTGAGGCTGCTTCTTCTGCTTTGG |
| Fwd1f | GATCTCATTC | GCTGCTTCTTCTGCTTTGG | GATCTCATTCGCTGCTTCTTCTGCTTTGG |
| Fwd1g | CGCTTATCCT | GCTGCTTCTTCTGCTTTGG | CGCTTATCCTGCTGCTTCTTCTGCTTTGG |
| Fwd1h | TATCATGCAG | GCTGCTTCTTCTGCTTTGG | TATCATGCAGGCTGCTTCTTCTGCTTTGG |
| Fwd2e | ATCTGCGTAC | GCTGAATGCGTACTTCCATC | ATCTGCGTACGCTGAATGCGTACTTCCATC |
| Fwd2f | GATTGCACGC | GCTGAATGCGTACTTCCATC | GATTGCACGCGCTGAATGCGTACTTCCATC |
| Fwd2g | ATGCTTCCTA | GCTGAATGCGTACTTCCATC | ATGCTTCCTAGCTGAATGCGTACTTCCATC |
| Fwd2h | TGCTAACTTC | GCTGAATGCGTACTTCCATC | TGCTAACTTCGCTGAATGCGTACTTCCATC |
| Fwd3e | ATAGCAGTGC | ATCCTGATTCCATGTCATCGC | ATAGCAGTGCATCCTGATTCCATGTCATCGC |
| Fwd3f | GAGCGAGTCA | ATCCTGATTCCATGTCATCGC | GAGCGAGTCAATCCTGATTCCATGTCATCGC |
| Fwd3g | CAGGCGATCT | ATCCTGATTCCATGTCATCGC | CAGGCGATCTATCCTGATTCCATGTCATCGC |
| Fwd3h | TTCACGGAAG | ATCCTGATTCCATGTCATCGC | TTCACGGAAGATCCTGATTCCATGTCATCGC |

**Table S6 Experimental conditions used during different rounds of screening and selection.**

| FACS Round | Substrate (PF-TMR) | |
|---|---|---|
| | Labelling time (min) | Concentration (nM) |
| Round 1 | 15 | 5 |
| Round 2 | 7.5 | 5 |
| Round 3 | 15 | 3.5 |
| Round 4 | 7.5 | 3.5 |

**Table S7 Microscopy settings used for measuring labelling kinetics of CLIP-tag2, CLIPf-tag and HaloTag7 in live U2OS cells.**

| Fluorophore | $\lambda_{ex}$ (nm) | $\lambda_{em}$ (nm) | HyD detector gain | Laser power (%) |
|---|---|---|---|---|
| TMR | 552 | 562-623 | 60 | 0.3 |
| CPY | 615 | 625-715 | 100 | 0.3 |
| SiR | 652 | 662-777 | 100 | 0.3 |
| mTurquoise2 | 448 | 458-520 | 100 | 0.3 |

**Table S8 Microscopy settings used for measuring fluorescence brightness between CLIP-tag2 and HaloTag7 in live U2OS cells.**

| Fluorophore | $\lambda_{ex}$ (nm) | $\lambda_{em}$ (nm) | HyD detector gain | Laser power (%) |
|---|---|---|---|---|
| TMR | 552 | 562-623 | 100 | 1.5 |
| CPY | 615 | 625-715 | 100 | 0.8 |
| SiR | 652 | 662-777 | 100 | 0.8 |
| mTurquoise2 | 448 | 458-520 | 100 | 0.8 |

**Table S9 Microscopy settings used in labeling of CLIP-tag2 fused to H2B, LifeAct, CEP41 and TOMM20.**

| Fluorophore | $\lambda_{ex}$ (nm) | Laser power (%) | | | | $\lambda_{em}$ (nm) |
|---|---|---|---|---|---|---|
| | | H2B | LifeAct | CEP41 | TOMM20 | |
| TMR | 550 | 0.5 | 1.0 | 2.0 | 2.0 | 560-600 |
| MaP555 | 550 | 0.5 | 1.0 | 2.0 | 2.0 | 560-600 |
| CPY | 610 | 0.5 | 0.5 | 0.5 | 0.5 | 620-700 |
| SiR | 635 | 0.5 | 0.5 | 2.0 | 2.0 | 645-750 |

**Table S10 Filter settings used in FP measurements.**

| Fluorophore | Excitation filter (nm) | Emission filter (nm) |
|---|---|---|
| TMR | 535/25 | 595/35 |

**Table S11 Filter settings used in YSD screening using FACS.**

| Fluorophore | Excitation laser (nm) | Emission filter (nm) | Filter |
|---|---|---|---|
| TMR | 561 | 582/15 | PE |
| Bilirubin | 488 | 527/32 | FITC |

**Table S12 Buffer and media compositions used in this study.**

| Reagent | Composition |
|---|---|
| $LB^{Amp}$ | 5 g $L^{-1}$ yeast extract, 10 g $L^{-1}$ peptone, 0.1 g $L^{-1}$ ampicillin |
| Activity buffer | 50 mM HEPES, 50 mM NaCl, pH 7.3 |
| FP buffer | Activity buffer, 0.1 mg $mL^{-1}$ BSA, 1 mM DTT, pH = 7.3 |
| Gibson Assembly master mix (60 x) | 0.64 µL T5 exonuclease (10 U $µL^{-1}$), 20 µL Phusion polymerase (2 U $µL^{-1}$), 160 µL Taq ligase (40 U $µL^{-1}$), 700 µL H2O, 320 µL 5 × ISO buffer, aliquoted to 20 µL per reaction |
| His-tag extraction buffer | 50 mM $KH_2PO_4$, 300 mM NaCl, 5 mM imidazole, pH 8.0 |
| His-tag wash buffer | 50 mM $KH_2PO_4$, 300 mM NaCl, 10 mM imidazole, pH 7.5 |
| His-tag elution buffer | 50 mM $KH_2PO_4$, 300 mM NaCl, 500 mM imidazole, pH 7.5 |
| YPD medium | 20 g $L^{-1}$ glucose, 20 g $L^{-1}$ peptone, 10 g $L^{-1}$ yeast extract, autoclaved (prior to addition of glucose) |
| YPD agar plates | YPD medium, 15 g $L^{-1}$ agar (1.5 % w/v) |
| YPDS | 1:1 mixture of YPD and 1 M sorbitol |
| Tris buffer | 1 M Tris-HCl, pH = 8 |
| Tris-DTT buffer | 0.39 g DTT (2.5 M) dissolved in Tris buffer (800 µL), filtered |
| Tris-LiAc buffer | 1.02 g LiAc × 2 $H_2O$ dissolved in Tris buffer (2 mL), filtered |
| Electroporation buffer | 10 mM Tris-base, 270 mM Sucrose, 2.1 mM $MgCl_2$ × 6 $H_2O$, pH = 7.5, autoclaved |
| SDCAA drop out medium | 20 g $L^{-1}$ glucose, 6.7 g $L^{-1}$ Difco yeast nitrogen base, 5 g $L^{-1}$ Bacto casamino acids (without tryptophan), 38 mM $Na_2HPO_4$ × 12 $H_2O$, 62 mM $NaH_2PO_4$ × $H_2O$, autoclaved (prior to addition of glucose) |
| SDCAA drop out agar plates | SDCAA drop out medium, 3 g $L^{-1}$ agar, 1 M sorbitol |
| SGCAA drop out medium | 20 g $L^{-1}$ galactose, 6.7 g $L^{-1}$ Difco yeast nitrogen base, 5 g $L^{-1}$ Bacto casamino acids (without tryptophan), 38 mM $Na_2HPO_4$ × 12 $H_2O$, 62 mM $NaH_2PO_4$ × $H_2O$, autoclaved (prior to addition of galactose) |
| Imaging medium | DMEM without phenol red + GlutaMAX™ (1x), 4.5 g $L^{-1}$ glucose, sodium pyruvate (1x), 10 % FBS |
| Cell growth medium | DMEM with phenol red + GlutaMAX™ (1x), 4.5 g $L^{-1}$ glucose, sodium pyruvate (1x), 10 % FBS |

### 3 Methods

#### 3.1 Molecular cloning and protein generation

Experiments were done with the original CLIP-tag sequence as described or with a mutant carrying the additional point mutation E30R, referred to as $CLIP_f$-tag.[4, 6] $CLIP_f$-tag has been reported to possess an about 2-fold faster labeling rate with BC-TMR than CLIP-tag.[3] Mutations were introduced *via* site-directed-mutagenesis (SDM) or Gibson assembly.[7] Primers for SDM were designed using the NEBaseChanger (New England Biolabs, NEB) and for Gibson assembly using the Geneious Prime® software. For Gibson assembly, primers contained an overlap of ~24 bp between vector amplified DNA and insert. Q5 site-directed mutagenesis kit (NEB) was used for SDM following the manufacturer's protocol. All primers were synthesized by Merck KGaA or Eurofins. The melting temperatures were calculated using the NEB $T_m$ Calculator. For PCR reactions KOD polymerase (Merck KGaA) was used.

PCR products were purified using QIAquick PCR Purification Kit, as per manufacturer instruction. The resulting plasmid constructs contained pET51b(+) vector (Novagen) and were transformed into high-efficiency NEB 5-alpha competent E. coli, and cultured on lysogeny broth (LB) agar plates supplemented with ampicillin ($LB^{Amp}$). Sequences of the target plasmids were validated by Sanger sequencing (Eurofins or Microsynth).

Plasmids with pET51b(+) vector encoding the protein of interest (POI) were transformed into electrocompetent or chemically competent *E. coli* BL21(DE3)-pLysS cells (Novagen). Cultures were grown in $LB^{Amp}$ medium at 37 °C. Cultures were grown to an $OD_{600}$ of 0.6-0.8 and were subsequently induced for protein expression by addition of β-D-thiogalactopyranoside (IPTG, 0.5 M) and incubated at 16 °C overnight. Cells were harvested by centrifugation (4000 x g, 4 °C, 15 min), pellet was resuspended in 20-30 mL His-tag extraction buffer supplemented with 1 mM PMSF and 0.25 mg/mL lysozyme, and lysed by sonication (50 % duty cycle, 70 % power, 7 min) on wet ice.

Cell lysates were cleared by centrifugation (15 000 x g, 4 °C, 15 min). Ni-NTA resin (HisPur™ Ni-NTA Resin, ThermoFischer, 600 µL) was added to the soluble fraction and incubated on a rolling shaker (4 °C, 1.5 h). Beads were isolated by centrifugation (1200 x g, 4 °C, 2 min) and the supernatant was carefully discarded. The resin was resuspended in the His-tag wash buffer before it was loaded to a polypropylene column (1 mL, Qiagen) for gravity purification. The column was washed with His-tag wash buffer (3 x 4 mL) and proteins were eluted by gravity after addition of His-tag elution buffer (2 mL). The eluted proteins were desalted with activity buffer (3 x 5 mL) and concentrated using Amicon Ultra centrifugal filters (10 kDa MWCO, 0.5 mL, Merck KGaA) to an approximate final volume of 250 µL. Final protein concentrations were measured with a

NanoDrop 2000C photo/spectrometer (Thermo) and purity was verified via SDS-Page and ESI-MS. Purified proteins were aliquoted, flash-frozen in liquid nitrogen and stored at −80 °C.

#### 3.2 Measurement of protein stability

Thermal stabilities of proteins were measured on a Prometheus NT48 nanoscale differential scanning fluorimeter (NanoDSF). Protein samples were prepared in activity buffer (0.8 mg $mL^{-1}$). The measurements were performed by following changes in tryptophan fluorescence (ratio signal at 350/330nm) over a temperature range of 20-95 °C, with a temperature increase of 1 °C $min^{-1}$. The inflection point of the first derivative corresponds to the proteins melting temperature $T_m$. The measurements were performed in technical triplicates.

#### 3.3 Measurement of labeling kinetics

A similar procedure was followed as described in Wilhelm et al.[3] and Kühn et al.[2]

Labelling rates were measured in black non-binding flat bottom 96-well plates (200 µL final reaction volume, Corning) on a microplate reader (Spark 20M, Tecan), by recording fluorescence polarization (FP) over time at 37 °C. All measurements were performed in technical triplicates in FP buffer, by addition of substrate to the protein, using a multichannel pipette or the Tecan SPARK injector module. For TMR substrates, 535 nm excitation filter with a 25 nm bandwidth was used. For emission filter and bandwidth, 595 and 35 nm was used. The G-factors were calculated using a FP buffer only (blank), with free fluorophore substrate as a (reference) control. Substrate baseline was determined by recording FP of the free fluorophore substrates.

##### 3.3.1 Measurement of labeling kinetics using a multi-well reader at fixed protein concentrations

For screening purposes, rates were recorded at fixed concentrations of 50 nM of protein and 20 nM of substrate. For faster proteins, rates were measured at 10 nM of protein and 4 nM of substrate. A one-phase association equation was fitted to the data using a custom R script and second order rate constants ($k_{app}$) were calculated using the following equation:

$$Y = Y_0 + (Y_{max} - Y_0) \times \left(1 - e^{(-k \times x)}\right) \tag{1}$$

where $Y$ is fluorescence polarization in mFP, $Y_{max}$ is a fluorescence polarization plateau in mFP, $Y_0$ is fluorescence polarization at time = 0, $k$ is labelling rate constant in $s^{-1}$ and $x$ is time in s. The apparent second-order rate constant ($k_{app}$) was calculated using the following equation:

$$k_{app} = \frac{k}{[protein]} \tag{2}$$

##### 3.3.2 Measurement of labeling kinetics using a multi-well plate reader at varying protein concentrations

Measurement at different protein concentrations and a global fit approach was used to allow for a more accurate determination of $k_{app}$ for selected CLIP-tag variants, as published in Wilhelm et al.[3] Fluorescence polarization was measured at a fixed substrate concentration (20 nM) and at varying protein concentrations (3 – 400 nM). Kinetic data were pre-processed using a custom R script. A kinetic model (eq. 3-6) was fitted to the data using DynaFit,[8] where delay time, protein concentration and fluorescence polarization baseline were fixed parameters, while substrate concentration was an adjustable parameter, to account for concentration determination errors. The standard deviation and confidence interval were calculated using Monte-Carlo method ($n$ = 1000, 5% worst fits excluded).[9] A representative graph is shown in Figure S6**.**

$$P + S \xrightarrow{k_{app}} PS \tag{3}$$

where $P$ is CLIP-tag protein variant, $S$ is the fluorophore substrate and $PS$ is the fluorescently labelled CLIP-tag variant.

$$\frac{d[P]}{dt} = -k_{app}[P][S] \tag{4}$$

$$\frac{d[S]}{dt} = -k_{app}[P][S] \tag{5}$$

$$\frac{d[PS]}{dt} = k_{app}[P][S] \tag{6}$$

##### 3.3.3 Measurement of CLIP-tag2 kinetics using stopped-flow device

Labelling kinetics of CLIP-tag2 with PF-TMR were measured by recording time-resolved changes in fluorescence anisotropy. The measurements were performed on a BioLogic SFM-400 stopped-flow instrument (BioLogic Science Instruments) in a single-mixing configuration at 37 °C, following a procedure described previously.[3]

The PF substrate concentration was fixed (0.5 µM), while the protein concentration (0.25-2.5 µM) was varied. Anisotropy measurements of the free PF substrate were performed to establish a baseline. Sampling was varied from 200 µs for 0.2 s, followed by 2 ms to a total acquisition time of 3 s. Each protein concentration was recorded in 15 technical replicates. Kinetic traces were preprocessed using a custom R script, which averaged replicates, removed pre-trigger time points and shifted time axes to start at zero. The kinetic model (eq. 7-14), was fitted to the processed data using DynaFit.[3, 8] Fluorophore substrate concentration was allowed to vary, to account for potential quantification inaccuracy, while protein concentration, anisotropy baseline and delay time were kept as fixed parameters. The use of a limiting protein concentration condition enabled more accurate concentration fitting, which influences the final anisotropy plateau (decreasing plateau). Monte Carlo simulation (with standard settings, n = 1,000, 5% worst fits excluded) was used to determine standard deviation (s.d.) and confidence interval (CI).[9] Generated stopped-flow graphics were obtained with custom R-script (Figure S2).

$$P + S \underset{k_{-1}}{\overset{k_1}{\leftrightarrow}} PS^* \tag{7}$$

$$PS^* \overset{k_2}{\rightarrow} PS \tag{8}$$

where *P* is the CLIP-tag2 protein, *S* is the fluorophore substrate, *PS** is the substrate-bound protein complex and PS is the fluorescently labeled CLIP-tag2. The system is described by the following equations:

$$\frac{d[P]}{dt} = -k_1[P][S] + k_{-1}[PS^*] \tag{9}$$

$$\frac{d[S]}{dt} = -k_1[P][S] + k_{-1}[PS^*] \tag{10}$$

$$\frac{d[PS^*]}{dt} = k_1[P][S] - k_{-1}[PS^*] - k_2[PS^*] \tag{11}$$

$$\frac{d[PS]}{dt} = k_2[PS^*] \tag{12}$$

The dissociation constant ($K_d$) accounting for the affinity between protein and the substrate, and $k_{app}$ defining the overall labeling speed were derived as follows:

$$K_d = \frac{k_{-1}}{k_1} \tag{13}$$

$$k_{app} = k_1 \frac{k_2}{(k_2 + k_{-1})} \tag{14}$$

#### 3.4 Yeast surface display

A similar procedure was followed as described in Kühn et al.[2] CLIP-tag variants were encoded on pJYDNg backbone (Addgene, #162452) with a C-terminal fusion to Aga2p (CLIP-tag-Aga2p-HA-cMyc-eUnaG).[5] The fluorescent protein eUnaG was labeled with bilirubin to serve as an expression marker.

##### 3.4.1 Synthetic deep mutational scanning library (sDMSL)

An sDMSL designed on the $CLIP_f$-tag scaffold was obtained from Twist Bioscience. The library consisted of single-point variants, in which each amino acid in $CLIP_f$-tag, except the reactive cysteine residue, has been replaced by the 19 proteinogenic amino acids, excluding cysteines but including deletions. The library was designed with a 50-bp overlap of the flanking regions to the pJYDNg expression vector for direct homologous recombination into yeast cells.

##### 3.4.2 Electrocompetent cell preparation

YPD media was inoculated with a single colony of Saccharomyces cerevisiae (*S. cerevisiae*) EBY 100 (American Type Culture Collection). The cells were grown at 30 °C and 250 rpm overnight. The overnight culture was used to inoculate 100 mL YPD to an $OD_{600}$ of 0.1 in a 1 L Erlenmeyer flask. Cells were grown at 30 °C and 250 rpm for 6 – 8 h until an $OD_{600}$ of 1.3 – 1.5 was reached. Tris-DTT (800 uL) and Tris-LiAc (2 mL) buffers were added, and the culture was incubated for 15 min (250 rpm, 30 °C). Cells were split into two 50 mL Falcon tubes and pelleted by centrifugation (2500 x g, 4 °C, 3 min). The pellets were washed with ice-cold electroporation buffer (25 mL), followed by resuspension in ice-cold electroporation buffer (200 uL, 2x) following the same centrifugation procedures. For transformation, electrocompetent cells were aliquoted (50 uL) and directly used for electroporation.

##### 3.4.3 Homologous recombination

For homologous recombination, purified vector (1 µg) and insert DNA (2-3 µg) were mixed to a final volume of 100 µL. The DNA mix was precipitated by addition of sodium acetate (10 µL, 3 M), glycogen (0.5 µg $µL^{-1}$, ThermoFisher) and isopropanol (100 µL) and stored at -20 °C overnight. Afterwards, the mixture was pelleted by centrifugation (10000 x g, 20 min) and the supernatant was removed. The DNA pellet was washed with 70 % EtOH (200 µl) and pelleted by centrifugation (10000 x g, 20 min.). The supernatant was discarded, and the pellet was air-dried. Once the pellet was dry, freshly prepared electrocompetent EBY100 yeast cells (50 µL) were used to resuspend the DNA. The cell–DNA mixture was transferred to the precooled electroporation cuvette (GenePulser cuvette, 0.2-cm electrode gap, Bio-Rad Laboratories) to perform electroporation using a Bio-Rad GenePulser Xcell device (0.54 kV, 25 µF, infinite resistance with exponential decay). Immediately after electroporation, pre-warmed YPDS medium (1 ml) was added into the cuvette. The suspension was transferred to culture tubes (15 ml). The cuvette was washed with YPDS medium (1 mL) and transferred to the same culture tube. The culture was incubated for 1 h (250 rpm, 30 °C). Yeast cells were pelleted by centrifugation (2 500 x g, 5 min), followed by resuspension in SDCAA selective medium (1 ml). To determine transformation efficiency, serial dilutions were plated on SDCAA agar plates. Plates were incubated for 2-3 days (30 °C). The remaining cell suspension was transferred into SDCAA medium (100 ml) and incubated for 2 days

(30 °C). This culture was then used directly for yeast surface display screening. Aliquots were preserved by freezing in a 1:1 mixture of the culture and 50% glycerol in Tris buffer (pH 8).

##### 3.4.4 Protein expression

SDCAA medium (5 mL) was inoculated either with several colonies from an SDCAA plate or with an aliquot (200 µL) of fully grown yeast liquid culture obtained after transformation. The culture was incubated overnight (250 rpm, 30 °C). The overnight pre-culture was centrifuged (4000 g, 5 min), the supernatant was discarded and SGCAA medium (5 mL) was added to induce protein expression on the yeast surface. Protein expression was allowed to proceed for at least 20 h (250 rpm, 30 °C).

##### 3.4.5 Protein labelling

The density of the yeast culture expressing the proteins was determined by accounting that an $OD_{600}$ of 1 corresponds to about $10^7$ cells $mL^{-1}$.[10] Yeast cells ($10^7$ cells) were added to PBS (500 µL), gently vortexed and centrifuged (14 000 x g, 1 min). Supernatant was discarded by carefully pipetting. Yeast cells were incubated with bilirubin (50 µL, 10 µM, in PBS supplemented with 0.1 mg $mL^{-1}$ BSA (PBSB)), vortexed and placed on ice for 10 min. The cells were pelleted and washed with PBS (150 µL), followed by centrifugation (14 000 x g, 1 min). The wash and centrifugation procedure was repeated, after which yeast cells were incubated in PBSB (50 µL) and labelled with TMR substrate (50 µL, PF-TMR or biotin-PF-TMR) at a desired labelling concentration (Table S6). Each sample was vortexed and incubated at 30 °C and 900 rpm (ThermoMixer™, Eppendorf), either for 15 or 7.5 min. After incubation, the reaction was quenched through the addition of $CLIP_f$-tag (500 µL, 500 µM), and incubated for further 20 min (900 rpm, 30 °C). The samples were centrifuged (14 000 x g, 1 min), washed with PBS (150 µL), centrifuged and washed again. Supernatant was disposed, and cells were resuspended in PBS (1 mL), and filtered through 5 mL round bottom polystyrene tubes with cell strainer snap caps (352235, Falcon). Filtered cells were subsequently subjected to FACS analysis.

##### 3.4.6 Fluorescence activated cell sorting

Yeast display libraries were analysed and sorted on a BD FACSMelody™ Cell Sorter (BD Biosciences) using the appropriate filter settings for bilirubin (488 nm excitation, 527/32 nm emission/band width) and TMR (561 nm excitation, 582/15 nm emission/band width). Sorting was carried out with a 100 µm sorting nozzle and a 1.5 neutral density filter. Detector voltages were adjusted using a single and double labelled control sample, together with negative (unstained) control samples. The gating strategy is depicted in Figure S6. Cells were gated for live and single cell events based on forward scattered and side scattered signals. Top 0.5-1.0% of single cell population with double positive signal (10 000 events) was sorted.

Following the sorting, cells were incubated with SDCAA medium (5 mL) supplemented with penicillin-streptomycin (Gibco, 5000 U $mL^{-1}$, 1:100 dilution) for 2 days. The fully grown yeast culture (200 µL) was added to SDCAA medium (5 mL) for the next sorting round, which was grown for 2 days (250 rpm, 30 °C). Additional 1 mL of culture was taken for plasmid isolation and sequencing using a Zymoprep Yeast Plasmid Miniprep II kit (Zymo Research), as per manufacturer instructions.

##### 3.4.7 Next generation sequencing and data analysis

The NGS was performed as described in Kühn et al.[2]

Randomized regions from recovered plasmids were amplified with next generation sequencing (NGS) primers following standard PCR protocol (for sequences, see Table S5). The PCR products were subsequently purified using Qiagen PCR purification kit. NGS was carried out by Eurofins Genomics using an Illumina adaptor-ligation workflow. The NGS package for each sample yielded approximately 10 million total reads (2 x 150 bp paired-end reads, 5 million read pairs). Amplicons ranging from 150-300 bp were combined in equal proportions. Primer-specific adapter sequences later on enabled assignment of specific sorting round for data analysis. For NGS, 2 µg of DNA in 100 µL of $ddH_2O$ was supplied per sample.

##### 3.4.8 Data analysis

Illumina sequencing datasets were first separated into barcode-defined groups corresponding to individual selection rounds. For each barcode, paired reads (forward and reverse reads) were merged into a single dataset. Primer and adapter regions were removed to ensure the correct reading frame of the underlying template, after which the cleaned reads were aligned to the reference sequence. The alignments were analysed with a custom R script to find enriched sequences or substitutions, listed by their read frequency. Mutation frequencies were calculated relative to the number of reads covering each position. Due to palindromic sequence present within the first reverse primer (Table S5, Rev1), data was truncated from residues 34-71 to prevent false positives.

#### 3.5 Generation of mammalian stable U2OS CLIP-tag2 cell lines

As described in Kühn et al (Generation of mammalian stable cell line).[2]

Stable U2OS cell lines were generated using the Flp-In T-Rex system (ThermoFischer). Lipofectamine 3000 (8 µL, L3000008) was diluted in Opti-MEM I reduced serum medium supplemented with GlutaMAX (200 µL, 51985026) and mixed well. In a second reaction tube, the plasmid encoding the gene of interest on a pcDNA/FRT backbone (440 ng) the pOG44 plasmid

(3560 ng) encoding the Flp-In recombinase were diluted in Opti-MEM I + GlutaMAX medium (200 µL). The P3000 reagent (8 µL) was added to the DNA mixture and thoroughly mixed. The DNA-P3000 mixture was allowed to equilibrate at room temperature for 10 min. Afterwards, the DNA–P3000 mixture was combined with the previously diluted Lipofectamine 3000, mixed and incubated at room temperature for additional 15 min. The transfection mix was then applied to cells cultured in T-25 flasks (at 80% confluency) and left to incubate overnight.

#### 3.6 Transient transfection of mammalian cells

U2OS cells were seeded into 96-well glass-bottom plates (Ibidi). Cells reached approximately 80% confluency before transfection with Lipofectamine 3000 (Thermo Fisher Scientific) according to the manufacturer's instructions. Specifically, one tube contained 0.2 µL Lipofectamine 3000 diluted in 10 µL Opti-MEM, while a second tube contained 100 ng plasmid DNA mixed with 10 µL Opti-MEM and supplemented with 0.2 µL P3000 reagent. The contents of the two tubes were then combined and incubated at room temperature for 10–15 min before being added to the cells. Approximately 16 h after transfection, the cells were subjected to live-cell staining.

#### 3.7 Preparation of rAAVs

Recombinant AAVs (rAAVs) were produced as described elsewhere.[11] In brief, pGP-AAV-CAG plasmids encoding for the different self-labeling tags fused to a localization sequence (LifeAct-CLIP2, Lyn11-SNAP2, CalR-HaloTag9-KDEL, Cox8A-HaloTag7) were transfected into HEK293 cells using Polyethylenimine 25000 (Sigma-Aldrich). 5 days post transfection, the cells were harvested and lysed using TNT extraction buffer (20 mM Tris pH 7.5, 150 mM NaCl, 1% TX-100, 10 mM $MgCl_2$). The cell debris was removed by centrifugation, the supernatant was collected and treated with Benzonase (Turbo Nuclease, Jena Bioscience) and applied to a AVB Sepharose column using an ÄktaPure FPLC instrument (Cytiva). rAAVs were concentrated with Ultra 15 mL Centrifugal Filters (Amicon®, MWKO: 100 kDa) and the buffer was exchanged to PBS pH 7.3.

#### 3.8 Microscopy

##### 3.8.1 Confocal fluorescence microscopy of transiently transfected U2OS cells

Labelling was carried out 16 h after transient transfection by incubating cells with 100 nM fluorescent CLIP-tag2 substrates diluted in growth medium for 1.5 h. Following incubation, labelled cells were washed three times with imaging medium (growth medium without phenol red) and subsequently imaged in the same imaging medium.

Confocal fluorescence microscopy of U2OS cells transiently expressing H2B/LifeAct/CEP41/TOMM20-CLIP-tag2 constructs was carried out on a TCS SP8 X microscope (Leica microsystems). Live-cell experiments were performed µ-Plate 96 Well Square (Ibidi) in a humidity chamber, maintained at physiological temperature of 37 °C and 5% $CO_2$ level in imaging medium. Acquisition was carried out using a HC PL APO CS2 40x/1.10 water-immersion objective and the following imaging parameters: line average of three, pixel size of 101nm x 101nm, pixel dwell time of 1.69µs, 1AU pinhole, Z stack 4x1µM, Laser settings are described in the Table S9. Image analysis was performed using Fiji software.[12]

##### 3.8.2 Fluorescence intensity comparison between HaloTag7 and CLIP-tag2

Confocal fluorescence imaging of U2OS cells stably expressing HaloTag7–P30–CLIP-tag2/$CLIP_f$-tag–NLS–P2A–NLS–mTurquoise2 constructs was performed on a Stellaris 5 microscope (Leica Microsystems). Live-cell experiments were performed using 8-well glass-bottom plates (Ibidi) placed inside a humidity chamber, maintained at 37 °C and 5% $CO_2$. Data were collected with a HC PL APO CS2 20x/0.75 water-immersion objective. Images were acquired at 400 Hz with a line average of 2, using optical zoom of 1. Image size were 1024 × 1024 pixels with a 12-bit pixel depth. Three-dimensional stacks were collected using *Z*-increments of 2-5 µm. HaloTag7 and CLIP-tag2 were labeled with CA- and PF-fluorophores, respectively (500 nM for 2 hours). Cells were washed twice with PBS before imaging in imaging medium. Laser settings are described in the table below (Table S8). Image analysis was performed using Fiji software.[12] For visualization purposes, maximum intensity projections of *z*-stacked images were calculated. For quantification, intensities of multiplane images were summed and regions of interest (ROIs) were defined manually. Mean fluorescence intensities of the ROI's fluorescence channels were calculated (multi-ROI measurement) and ratios were calculated from fluorescent labels to respective mTurquoise2 expression signals.

##### 3.8.3 In-cell kinetic measurements of CLIP-tag2, $CLIP_f$-tag and HaloTag7

As described in Kühn et al (In cell kinetics measurements).[2]

Labeling of CLIP-tag2, $CLIP_f$-tag and HaloTag7 with PF-fluorophore, BC-fluorophore and CA-fluorophore substrates was performed in live U2OS cells with stable expression of HaloTag7-P30-CLIP-tag2/$CLIP_f$-tag-NLS-P2A-NLS-mTurquoise2. Measurements were performed on a Stellaris 5 microscope. After the addition of TMR (50 nM final), CPY (50 nM final) and SiR (100 nM final) to the cells, images were obtained in three z stacks which acquired every 30 s over imaging time. Fluorescence intensities from multiplane images were summed and the fluorophore substrate signal was normalized to the mTurquoise2 expression signal using CellProfiler.[13] Images were acquired with HC PL APO CS2 20x/0.75 water-immersion objective with optical zoom value of 1.28. Image size was 1024 × 1024 pixels with 32 bits per pixel. Image acquisition was performed with a scan speed of 400 Hz and a line average of 2. Laser settings are described in Table S7. Kinetic data were analyzed in GraphPad Prism by fitting a sigmoidal curve (eq. 15).

$$Y = Y_0 + \frac{(Y_{\max} - Y_0)}{\left(1 + \left(\frac{t_{1/2}}{x}\right)^H\right)} \tag{15}$$

where $Y$ is the ratio of signal/m Turquoise2 fluorescence intensity, $Y_{\max}$ is the plateau of the fluorescence intensity ratio, $Y_0$ is the $y$-intercept, $H$ is the Hill slope, and $x$ is time (min).

For each substrate, the experiment was performed in biological duplicates, and the mean $t_{1/2}$ value was calculated from the two replicates. Respective confidence intervals (CI) were propagated using equations 16-19 as follows:

1. Mean $t_{1/2}$ of both replicates was calculated:

$$\overline{t_{1/2}} = \frac{(t_{1/2})_1 + (t_{1/2})_2}{2} \tag{16}$$

2. Standard error (SE) was computed for each $t_{1/2}$ of a replicate:

$$SE_{(t_{1/2})_1} = \frac{CI_{1\,high} - CI_{1\,low}}{2 \times 1.96}$$
$$SE_{(t_{1/2})_2} = \frac{CI_{2\,high} - CI_{2\,low}}{2 \times 1.96} \tag{17}$$

3. Standard error of the mean was calculated.:

$$SE_{\overline{(t_{1/2})}} = \sqrt{\frac{SE^2_{(t_{1/2})_1} + SE^2_{(t_{1/2})_2}}{2}} \tag{18}$$

4. 95% confidence interval was computed for the mean $t_{1/2}$:

$$CI_{\overline{(t_{1/2})}} = \overline{t_{1/2}} \pm 1.96 \times SE_{\overline{(t_{1/2})}} \tag{19}$$

##### 3.8.4 Live-cell STED imaging

U2OS cells were transduced with ~ $10^9$ -$10^{10}$ rAAV particles for 16 hours and afterwards labeled with 200 nM dyes for 30 min at 37 °C in imaging medium. The medium was replaced twice and the cells were imaged live on a Abberior STED Expert Line 595/775/RESOLFT QUAD scanning microscope. The system was equipped with a UPlanSApo 100x/1.4 oil immersion objective lens (Abberior Instruments) and avalanche photodiodes (APD). Images were recorded in a 60 x 60 µm (overview) or 25 x 25 µm (zoom) frame with 80, 60 nm (confocal) or 30 nm (STED) pixel sizes. The pixel dwell time was set between 7 and 15 µs with up to 7 times line accumulation. The following channels were selected. 500R: Ext. 488 nm (20%); Em. 489 – 530 nm. MaP555: Ext. 561 nm (20%); Em. 587 – 630 nm. SiR: Ext. 640 nm (3%); Em. 670 – 746 nm; STED 775 nm (20%).

##### 3.9 Cell viability assay

As described in Kühn et al (Cell viability assay).[2]

U2OS cells were seeded on transparent 96-well culture plates and left to incubate (37 °C, 5% $CO_2$) for one day. Cells were then incubated with PF-fluorophore (1 µM), 1% DMSO (v/v), or left untreated. After 1 hour of incubation at 37 °C, the medium was transferred to a non-binding U-bottom 96-well plate (Flacon), and detached cells were pelleted by centrifugation (3000 x g, 5 min) before the supernatant was removed. Adherent cells were detached with trypsin (30 µL, 10 min, 37 °C) and resuspended in FACS buffer (2% FBS in PBS) to a final volume of 100 µL. These cells were combined with the detached cell fraction to ensure recovery of both dead and live populations. SYTOX Blue dead cell stain (100 µL, 2 µM, ThermoFischer) was added (1 µM final concentration). Samples were analyzed via flow cytometry (10 000 events, 405 nm excitation laser, 450/50 nm emission/band width filter). Technical triplicates were performed for each sample, and the data was analyzed with FlowJo software (BD Biosciences).

##### 3.10 Maestro Schrödinger and Boltz2 calculations

Ligand pKa values and the most probable ionization states were predicted with the Epik module in Maestro (Schrödinger Maestro 2024-4; Schrödinger Inc., New York) using default settings.

In-silico ligand–protein co-folding was performed with Boltz2,[14] and the resulting poses were aligned to reference crystal structures (PDB IDs: 3KZZ & 6Y8P). The most plausible binding mode from these comparisons was selected for molecular dynamics (MD) studies carried out in Maestro (Schrödinger Maestro 2024-4). Protein–ligand complexes were prepared with the Protein Preparation Wizard (default parameters), and each system was solvated in an orthorhombic periodic box with a 10 Å buffer using the Desmond System Builder. Sodium and chloride ions were added to neutralize the system and to achieve a final ionic strength of 0.15 M NaCl. The OPLS4 force field was applied and MD simulations were executed with the Desmond Molecular Dynamics module following the module's standard algorithms and protocols. After equilibration, simulations of 100 ns were performed at 300 K and 1.01325 bar. Trajectory analysis, including monitoring of protein–ligand interactions and extraction of representative frames, was conducted using the Simulation Interaction Diagram tool within Maestro.

All computations were executed on a Linux-based high-performance computing server featuring two Intel Xeon CPUs (4.1 GHz, 32 cores), 512 GB RAM, a 9 TB local RAID SSD storage system, and four Nvidia L40S GPUs (48 GB each). Molecular graphics and analyses were conducted using UCSF ChimeraX, developed by the Resource for Biocomputing, Visualization, and Informatics at the University of California, San Francisco, with support from NIH R01-GM129325 and the Office of Cyber Infrastructure and Computational Biology, NIAID.

#### 3.11 Important protein sequences

##### 3.11.1 Sequences for bacterial expression

| General color code | | |
|---|---|---|
| His-Tag<br>Catalytic residue | TEV-cleavage site<br>CLIP-tag sequence | Mutated amino acid sequences compared to original CLIP-tag |

CLIP$_f$-tag (CLIP-tag-E30R)

```
MHHHHHHHHHHENLYFQG|MDKDCEMKRTTLDSPLGKLELSGCEQGLHRIIFLGKGTSAADAVEVPAPAAVLGGPEP
LIQATAWLNAYFHQPEAIEEFPVPALHHPVFQQESFTRQVLWKLLKVVKFGEVISESHLAALVGNPAATAAVNTALD
GNPVPILIPCHR VVQGDSDVGPYLGGLAVKEWLLAHEGHRLGKPGLGG
```

CLIPf-tag-L159A-G160A

```
MHHHHHHHHHHENLYFQG|MDKDCEMKRTTLDSPLGKLELSGCEQGLHRIIFLGKGTSAADAVEVPAPAAVLGGPEP
LIQATAWLNAYFHQPEAIEEFPVPALHHPVFQQESFTRQVLWKLLKVVKFGEVISESHLAALVGNPAATAAVNTALD
GNPVPILIPCHR VVQGDSDVGPYAAGLAVKEWLLAHEGHRLGKPGLGG
```

CLIP-tag2

```
MHHHHHHHHHHENLYFQG|MDKDCEMKRTTYDSPLGKLLLSGCEQGLHRIYFIGNGQGEQGPPGPEPLIQATAWLNA
YFYQPEAIEEFPVPALHHPVFQQESFTRQVLWKLLKVVKFGEVISESDLAALVGNPGATAAVNTALDSNPVPILIPC
HRVISGDSDVGPYAAGVAVKEWLLAHEGHRLGKPGYGG
```

##### 3.11.2 Sequences for expression on yeast surface

| General color code | | |
|---|---|---|
| Gal1 promoter<br>Linker<br>Factor Xa site | Aga2p<br>Catalytic residue<br>Affinity tags | Mutated amino acid sequences compared to original CLIP-tag<br>CLIP-tag sequence<br>eUnaG2 |

CLIPf-tag-L159A-G160A
MYYFLFKCNKSINKKLLIYLYTLTSRRKNKNPGSNSLLHTFSIKMRFPSIFTAVVFAASSALAAPANGTM|MDKDCE
MKRTTLDSPLGKLELSGCEQGLHRIIFLGKGTSAADAVEVPAPAAVLGGPEPLIQATAWLNAYFHQPEAIEEFPVPA
LHHPVFQQESFTRQVLWKLLKVVKFGEVISESHLAALVGNPAATAAVNTALDGNPVPILIPCHRVVQGDSDVGPYAA
GLAVKEWLLAHEGHRLGKPGLGGAAAFSQKLDINLLDNVVNSSYHGEGVSGGSAQELTTICEQIPSPTLESTPYSLS
TTTILANGKAMQGVFEYYKSVTFVSNCGSHPSTTSKGSPINTQYVFKDNSSTIEGRYPYDVPDYALQASGGGGSGGG
GSGGGGSASHEQKLISEEDLMLEKFVGTWKIESSENFGEYLKAIGAPKELADAGDATTPVLYISQKDGDKMTVKIEN
GPPTFLDTQVSFKLGEEFDEFPSDRRKGVKSVVNLSGEKLVYVQKWDGKETTYVREIKDGKLVVTLTMGDVVAVRSY
RRASE

##### 3.11.3 Sequences for expression in mammalian cells

| General color code | | |
|---|---|---|
| Kozak sequence<br>Catalytic residue<br>Linker | Fluorescent protein (FP)<br>CLIP-tag sequence<br>HaloTag7 | Mutated amino acid sequences compared to original CLIP-tag<br>Localization/fusion protein sequence<br>Cleavage sequence |

>pcDNA5/FRT/TO_H2B-CLIP-tag2
MPEPAKSAPAPKKGSKKAVTKAQKKGGKKRKRSRKESYSIYVYKVLKQVHPDTGISSKAMGIMNSFVNDIFERIAGE
ASRLAHYNKRSTITSREIQTAVRLLLPGELAKHAVSEGTKAITKYTSAKGG|DKDCEMKRTTYDSPLGKLLLSGCEQ
GLHRIYFIGNGQGEQGPPGPEPLIQATAWLNAYFYQPEAIEEFPVPALHHPVFQQESFTRQVLWKLLKVVKFGEVIS
ESDLAALVGNPGATAAVNTALDSNPVPILIPCHRVISGDSDVGPYAAGVAVKEWLLAHEGHRLGKPGYGG

>pcDNA5/FRT/TO_LifeAct-CLIP-tag2
MGVADLIKKFESISKEEGDPPVAT|DKDCEMKRTTYDSPLGKLLLSGCEQGLHRIYFIGNGQGEQGPPGPEPLIQAT
AWLNAYFYQPEAIEEFPVPALHHPVFQQESFTRQVLWKLLKVVKFGEVISESDLAALVGNPGATAAVNTALDSNPVP
ILIPCHRVISGDSDVGPYAAGVAVKEWLLAHEGHRLGKPGYGG

>pcDNA5/FRT/TO_TOMM20-CLIP-tag2
MVGRNSAIAAGVCGALFIGYCIYFDRKRRSDPNFKNRLRERRKKQKLAKERAGLSKLPDLKDAEAVQKFFLEEIQLG
EELLAQGEYEKGVDHLTNAIAVCGQPQQLLQVLQQTLPPPVFQMLLTKLPTISQRIVSAQSLAEDDVEGGSGDPPVG
G|DKDCEMKRTTYDSPLGKLLLSGCEQGLHRIYFIGNGQGEQGPPGPEPLIQATAWLNAYFYQPEAIEEFPVPALHH
PVFQQESFTRQVLWKLLKVVKFGEVISESDLAALVGNPGATAAVNTALDSNPVPILIPCHRVISGDSDVGPYAAGVA
VKEWLLAHEGHRLGKPGYGG

>pcDNA5/FRT/TO_Cep41-CLIP-tag2
MSLRRHIGNPEYLMKRIPQNPRYQHIKSRLDTGNSMTKYTEKLEEIKKNYRYKKDELFKRLKVTTFAQLIIQVASLS
DQTLEVTAEEIQRLEDNDSAASDPDAETTARTNGKGNPGEQSPSPEQFINNAGAGDSSRSTLQSVISGVGELDLDKG
PVKKAEPHTKDKPYPDCPFLLLDVRDRDSYQQCHIVGAYSYPIATLSRTMNPYSNDILEYKNAHGKIIILYDDDERL
ASQAATTMCERGFENLFMLSGGLKVLAQKFPEGLITGSLPASCQQALPPGSARKRSSPKGPPLPAENKWRFTPEDLK
KIEYYLEEEQGPADHPSRLNQANSSGRESKVPGARSAQNLPGGGPASHSNPRSLSSGHLQGKPWKYLECGRNPAFLY
KVVMAASLEVLFQGPFE|DKDCEMKRTTYDSPLGKLLLSGCEQGLHRIYFIGNGQGEQGPPGPEPLIQATAWLNAYF
YQPEAIEEFPVPALHHPVFQQESFTRQVLWKLLKVVKFGEVISESDLAALVGNPGATAAVNTALDSNPVPILIPCHR
VISGDSDVGPYAAGVAVKEWLLAHEGHRLGKPGYGG

>pcDNA5/FRT-HaloTag7-CLIPf-tag-NLS-P2A-NLS-mTurquoise2
TT|MGSEIGTGFPFDPHYVEVLGERMHYVDVGPRDGTPVLFLHGNPTSSYVWRNIIPHVAPTHRCIAPDLIGMGKSD
KPDLGYFFDDHVRFMDAFIEALGLEEVVLVIHDWGSALGFHWAKRNPERVKGIAFMEFIRPIPTWDEWPEFARETFQ
AFRTTDVGRKLIIDQNVFIEGTLPMGVVRPLTEVEMDHYREPFLNPVDREPLWRFPNELPIAGEPANIVALVEEYMD
WLHQSPVPKLLFWGTPGVLIPPAEAARLAKSLPNCKAVDIGPGLNLLQEDNPDLIGSEIARWLSTLEISG|SGRPPP
PPPPPPPPPPPPPPPPPPPPPPPPPPPPGGRSRSLE|MDKDCEMKRTTLDSPLGKLELSGCEQGLHRIIFLGKGTSAA
DAVEVPAPAAVLGGPEPLIQATAWLNAYFHQPEAIEEFPVPALHHPVFQQESFTRQVLWKLLKVVKFGEVISESHLA
ALVGNPAATAAVNTALDGNPVPILIPCHRVVQGDSDVGPYLGGLAVKEWLLAHEGHRLGKPGLGG|GAPDPKKKRKV

DPKKKRKVDPKKKRKELRASPQATNFSLLKQAGDVEENPGPSRMAPKKKRKVMVSKGEELFTGVVPILVELDGDVNG
HKFSVSGEGEGDATYGKLTLKFICTTGKLPVPWPTLVTTLSWGVQCFARYPDHMKQHDFFKSAMPEGYVQERTIFFK
DDGNYKTRAEVKFEGDTLVNRIELKGIDFKEDGNILGHKLEYNYFSDNVYITADKQKNGIKANFKIRHNIEDGGVQL
ADHYQQNTPIGDGPVLLPDNHYLSTQSKLSKDPNEKRDHMVLLEFVTAAGITLGMDELYK

>pcDNA5/FRT-HaloTag7-CLIP-tag2-NLS-P2A-NLS-mTurquoise2
TT|MGSEIGTGFPFDPHYVEVLGERMHYVDVGPRDGTPVLFLHGNPTSSYVWRNIIPHVAPTHRCIAPDLIGMGKSD
KPDLGYFFDDHVRFMDAFIEALGLEEVVLVIHDWGSALGFHWAKRNPERVKGIAFMEFIRPIPTWDEWPEFARETFQ
AFRTTDVGRKLIIDQNVFIEGTLPMGVVRPLTEVEMDHYREPFLNPVDREPLWRFPNELPIAGEPANIVALVEEYMD
WLHQSPVPKLLFWGTPGVLIPPAEAARLAKSLPNCKAVDIGPGLNLLQEDNPDLIGSEIARWLSTLEISG|SGRPPP
PPPPPPPPPPPPPPPPPPPPPPPPPPPGGRSRSLE|MDKDCEMKRTTYDSPLGKLLLSGCEQGLHRIYFIGNGQGEQ
GPPGPEPLIQATAWLNAYFYQPEAIEEFPVPALHHPVFQQESFTRQVLWKLLKVVKFGEVISESDLAALVGNPGATA
AVNTALDSNPVPILIPCHRVISGDSDVGPYAAGVAVKEWLLAHEGHRLGKPGYGG|GAPDPKKKRKVDPKKKRKVDP
KKKRKELRASPQATNFSLLKQAGDVEENPGPSRMAPKKKRKVMVSKGEELFTGVVPILVELDGDVNGHKFSVSGEGE
GDATYGKLTLKFICTTGKLPVPWPTLVTTLSWGVQCFARYPDHMKQHDFFKSAMPEGYVQERTIFFKDDGNYKTRAE
VKFEGDTLVNRIELKGIDFKEDGNILGHKLEYNYFSDNVYITADKQKNGIKANFKIRHNIEDGGVQLADHYQQNTPI
GDGPVLLPDNHYLSTQSKLSKDPNEKRDHMVLLEFVTAAGITLGMDELYK

#### 3.12 Scripts

##### 3.12.1 NGS analysis of libraries

As described in Kühn et al (NGS analysis of libraries).[2]

Demultiplex command:

```
je demultiplex F1=[path/to/fwd_reads] \
   F2=[path/to/rev_reads] O=out_split_reads \
   BF=[path/to/barcodes_file] GZ=false UF1=unassigned_1 \
   UF2=unassigned_2 M=jemultiplexer_out_stats
```

Combining paired reads command:

```
cat [path/to/fwd_reads] [path/to/rev_reads] >> output
```

Trimming command:

```
cutadapt -g [primer_sequences_file] \
   -a [adapter_sequences_file] --times 3 --overlap 6 \
   -e 0.2 -m 16 -j 8 -o output [input_reads] > \
   trim.log
```

Align command:

```
bowtie2 -p 8 -N 1 -L 6 --n-ceil L,150,1 --mp 6,2 --np 0 \
   --rdg 50,6 --rfg 50,6 --score-min L,-1,-0.5 -x \
   [template_sequence_file] -U [input.reads] -S output 2> \
   align.log"
```

Conversion to bam file command:

```
samtools view -b [input_file] > output
```

##### 3.12.2 Example DynaFit scripts[8] for the analysis of labelling kinetics using multi-well plate reader

As described in Kühn et al (SNAP-tag2 kinetics fitted to model 1).[2]

```
[task]
   data       = progress
   task       = fit
   confidence = monte-carlo

[mechanism]
   P + S ----> P.S      :     kapp

[constants] ; units: nM, sec

   kapp  = 0.001 ?

[concentrations] ; units: nM
   S = 20?

[responses]
   P.S = 2?

[parameters]

   A = 1

[data]
   delay 0
   offset FP-value free fluorophore
   directory  path/to/data
   sheet      data.csv

   column   2 | conc P =     10 | label   10_1
   column   3 | conc P =     10 | label   10_2
   column   4 | conc P =     10 | label   10_3
   column   5 | conc P =    118 | label  118_1
   column   6 | conc P =    118 | label  118_2
   column   7 | conc P =    118 | label  118_3
   column   8 | conc P =     16 | label   16_1
   column   9 | conc P =     16 | label   16_2
   column  10 | conc P =     16 | label   16_3
   column  11 | conc P =    177 | label  177_1
   column  12 | conc P =    177 | label  177_2
   column  13 | conc P =    177 | label  177_3
   column  14 | conc P =     23 | label   23_1
   column  15 | conc P =     23 | label   23_2
   column  16 | conc P =     23 | label   23_3
   column  17 | conc P =    266 | label  266_1
   column  18 | conc P =    266 | label  266_2
   column  19 | conc P =    266 | label  266_3
   column  20 | conc P =     35 | label   35_1
   column  21 | conc P =     35 | label   35_2
   column  22 | conc P =     35 | label   35_3
   column  23 | conc P =    400 | label  400_1
   column  24 | conc P =    400 | label  400_2
   column  25 | conc P =    400 | label  400_3
   column  26 | conc P =     53 | label   53_1
   column  27 | conc P =     53 | label   53_2
   column  28 | conc P =     53 | label   53_3
   column  29 | conc P =     79 | label   79_1
   column  30 | conc P =     79 | label   79_2
   column  31 | conc P =     79 | label   79_3
```

```
[output]
   Directory path/to/output/folder

[settings]
{ConfidenceIntervals}
   LevelPercent = 95
{Constraints}
   Constants = 1000000000000000
{Output}
   XAxisLabel = time [s]
   YAxisLabel = polarization
[end]
```

##### 3.12.3 Analysis of stopped-flow labelling kinetics

As described in Kühn et al (SNAP-tag2 kinetics fitted to model 2).[2]

```
[task]
   data  = progress
   task  = fit
   confidence = monte-carlo
[mechanism]
   P + S <===> P.S     :     k1    k_m1
   P.S ----> Z       :     k2
[constants] ; units: uM, sec
   k1 = 15?
   k_m1 = 0.01?
   k2 = 3?
[concentrations] ; units: uM
  S = 0.5?
[responses]
intensive
   S = 0.071
   P.S = 0.2?
   Z   = 1.0 * P.S

[data]
   directory  path/to/data
   sheet      data.csv

   column   2 | conc P = 2.5 | label 2.5

   column   3 | conc P = 2.0 | label 2.0

   column   4 | conc P = 1.5 | label 1.5

   column   5 | conc P = 1.0 | label 1.0

   column   6 | conc P = 0.5 | label 0.5

   column   7 | conc P = 0.417 | label 0.417

   column   8 | conc P = 0.25 | label 0.25



[output]
  directory path/to/output/folder

[settings]
{ConfidenceIntervals}
   LevelPercent = 95

{Constraints}
   Constants = 1000000000000000

{Output}
   XAxisLabel = time [s]
   YAxisLabel = anisotropy

{Marquardt}
   EqualizeDatasets = y

[end]
```

### 4 Chemical Synthesis of CLIP-tag2 Substrates

##### General Remarks

All reagents were obtained from commercial suppliers (Acros Chemicals, Alfa Aesar, TCI Chemicals GmbH, ABCR, Sigma-Aldrich, Activate Scientific, Carl Roth GmbH + Co. KG, Merck KGaA, VWR International). All solvents used in reactions were anhydrous. When necessary, solvents were degassed either by three freeze–pump–thaw cycles or by purging with nitrogen for a minimum of 15 minutes. Anhydrous solvents were handled under an argon atmosphere, and all reactions were conducted in oven-dried glassware. Deionized water was used for all experiments.

Reaction progress was monitored by thin-layer chromatography (TLC) or liquid chromatography–mass spectrometry (LCMS-2020, Shimadzu) connected to a Nexera X-2 UHPLC system equipped with a Supelco C18 column (80 Å, 1.9 µm pore size, 2.1 × 50 mm). A solvent gradient of 10–95% $MeCN/H_2O$ containing 0.1% v/v formic acid was applied at a flow rate of 1 mL/min over 6 minutes. TLC was performed on pre-coated silica gel plates (60G F254, Merck KGaA) using appropriate solvent systems. Visualization of reaction spots was achieved under UV illumination (254 nm or 366 nm) and/or by staining (dip, dry, and heat development). The staining solution consisted of $KMnO_4$ (1 g), $K_2CO_3$ (6.6 g), and 5% NaOH (1.7 mL) in $H_2O$ (90 mL).

All synthesized products were purified either by normal-phase flash column chromatography (FCC) using an automated Biotage Isolera One system with pre-packed ultrapure silica gel columns (12 g or 25 g, SiliCycle Inc.) or by reverse-phase high-performance liquid chromatography (RP-HPLC) on a Thermo Fisher Scientific UltiMate 3000 system equipped with a Supelco column (21.1 × 250 mm, 5 µm pore size, 8 mL/min flow rate). The RP-HPLC purification employed a solvent gradient of 10–95% $MeCN/H_2O$ with 0.1% v/v trifluoroacetic acid additive. A standard purification run lasted 45 minutes. Fractions containing the desired product were combined and concentrated in vacuo using a rotary evaporator (Hei-Vap Value, Heidolph) with heating in a 40 °C water bath and/or dried on a lyophilizer (Christ) connected to a Vacuubrand vacuum pump. Final fluorophore substrates were stored as DMSO stocks at –20 °C.

##### Nuclear Magnetic Resonance (NMR) Spectroscopy

Samples for NMR spectroscopy were dissolved in deuterated solvents, and spectra were recorded at 298 K on a BRUKER Avance III HD 400 NMR spectrometer equipped with a CryoProbe™ ($^1H$: 400 MHz, $^{13}C$: 101 MHz). Spectra were analyzed using MestReNova 14.1.0 (MestreLab Research). Multiplicities are reported as s = singlet, d = doublet, t = triplet, q = quartet, m = multiplet. Chemical shifts (δ) were calibrated to the residual solvent signals ($CDCl_3$, MeOD-$d_4$, MeCN-$d_3$, DMSO-$d_6$, DMF-$d_7$). Coupling constants (J) are reported in Hz. NMR spectra are presented as recorded.

##### High-Resolution Mass Spectrometry (HRMS)

HRMS data were acquired on a Bruker maXis II™ ETD-HRMS system using electrospray ionization (ESI) in positive mode. Measurements were conducted by the MS Core Facility of Max Planck Institute for Medical Research, Heidelberg.

#### 4.1 Synthesized molecules

##### 4.1.1 Building blocks – nucleobases

**2-(methylthio)-6-propylpyrimidin-4-ol (22):**

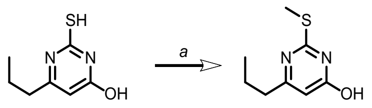


Thiopyrimidine (**21**) was methylated like previously described for similar substrate.[15] Thiopyrimidine (2.0 g, 11.7 mmol, 1 equiv.) was suspended in water (~50 mL) containing NaOH (2.34 g, 58 mmol, 5 equiv.) and dimethyl sulfate (1.63 g, 12.9 mmol, 1.1 equiv.) was added by slow dropping via addition funnel over 1 h. Reaction mixture was stirred for 3 hours after which it was heated to reflux and kept at this temperature for 10 min. Solution was filtered and pH was adjusted to 1-2 with 12 M HCl. After storage in the fridge at 4 °C, product was collected by filtration (filter with sintered frit glass filter, pore size 4) washed with additional water (~50 mL) and dried at 80 °C for 16 h. White solid product **22** (2.09 g, 97% yield) was obtained and used in the next step without further purification.

**TLC:** *Rf* = 0.53 (EtOAc/*n*-Hexane = 1:1).

**$^1$H NMR** (400 MHz, DMSO) *δ* = 12.45 (s, 1H), 5.92 (s, 1H), 2.47 (s, 3H), 2.39 (t, *J*=7.5, 2H), 1.61 (d, *J*=7.4, 2H), 0.89 (s, 3H) ppm.

**$^{13}$C{$^1$H} NMR** (101 MHz, DMSO) *δ* = 167.5, 163.5, 162.3, 106.5, 38.5, 20.6, 13.5, 12.7 ppm.

**HRMS** (EI$^+$) m/z: [M + H]$^+$, Calculated for $C_8H_{12}N_2OS^+$ 185.0743, found 185.0741.

**2-(methylthio)-6-propylpyrimidin-4-yl trifluoromethanesulfonate (23):**

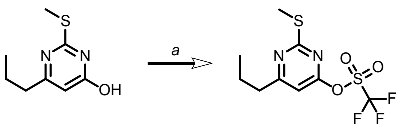


Methyl thiopyrimidine **22** (1 g, 5.43 mmol, 1 equiv.) and DIPEA (2.1 g, 16.3 mmol, 2.7 mL, 3 equiv.) was dissolved in DCM (~50 mL) and cooled to 0 °C. Triflic anhydride (1.83 g, 1.1 mL, 6.5 mmol, 1.2 equiv.) was added and reaction mixture was stirred at 0 °C for 4 h. After full conversion was confirmed by TLC, reaction was diluted with additional 50 mL of DCM and washed with saturated aqueous solution of NaHCO3 (2 x ~30 mL). Organic fraction was dried with solid $Na_2SO_4$ and purified by FCC to provide triflate 1.1 g **23** in 63% yield, as a thick yellow oil.

**TLC:** *Rf* = 0.75 (EtOAc/*n*-Hexane = 1:9).

**$^1$H NMR** (400 MHz, $CDCl_3$) *δ* = 6.63 (s, 1H), 2.75 (s, 2H), 2.57 (s, 3H), 1.78 (d, *J*=7.6, 2H), 1.00 (s, 3H) ppm.

**$^{13}C\{^1H\}$ NMR** (101 MHz, $CDCl_3$) $\delta$ = 176.3, 173.9, 162.8, 123.2, 120.1, 116.9, 113.7, 104.2, 39.8, 21.8, 14.2, 13.7 ppm.

**19F{$^1$H}** (376 MHz, $CDCl_3$) $\delta$ = -72.9 ppm.

**HRMS** ($EI^+$) m/z: $[M + H]^+$, Calculated for $C_9H_{12}N_2O_3F_3S_2^+$ 317.0236, found 317.0233.

***tert*-butyl (2-(methylthio)pyrimidin-4-yl)carbamate (20):**

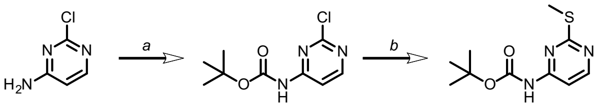

Chloropyrimidine (17) was protected with Boc2O like described before.[16] Carbamate 18 (1.34 g, 5.83 mmol, 1 equiv.) was dissolved in DMF (~100 mL) at room temperature and solid sodium methanethiolate (0.45 g, 6.42 mmol, 1.1 equiv.), was added in one portion. Reaction mixture was stirred for 16 h after which DMF was removed under reduced pressure. Solid residue was dissolved in DCM (~200 mL) and washed with water (3 x ~50 mL). Organic fraction was dried with solid $Na_2SO_4$ and purified by FCC.

TLC: Rf = 0.18 (EtOAc/n-Hexane = 1:9).

1H NMR (400 MHz, CDCl3) δ = 8.33 (d, J=5.8, 1H), 7.54 (d, J=5.7, 1H), 7.35 (s, 1H), 2.48 (s, 3H), 1.50 (s, 9H) ppm.

13C{1H} NMR (101 MHz, CDCl3) δ = 171.7, 159.9, 157.9, 157.7, 151.5, 106.8, 103.6, 82.3, 28.1, 28.0, 13.9 ppm.

HRMS (EI+) m/z: [M + H]+, Calculated for C10H16N3O2S+ 242.0958, found 242.0959.

***tert*-butyl (2-(methylsulfonyl)pyrimidin-4-yl)carbamate (19):**

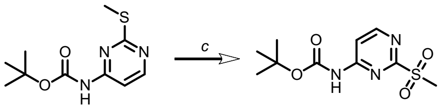

Oxidation to methyl sulfone was adapted from previously reported procedure.[17] Methylthiopyrimidine **20** (1.2 g, 4.9 mmol) was dissolved in mixture of water and DCM (~50 mL, 1:1) and sodium tungstate dihydrate (82 mg, 0.23 mmol, 0.05 mmol) was added followed by one drop of acetic acid. Hydrogen peroxide (2 mL, 2.25 g, 4 equiv., 30% in $H_2O$) was added dropwise at room temperature and mixture was heated to 50 °C under reflux condenser for 16 h. After cooling to 0 °C, remaining hydrogen peroxide was quenched by addition of aqueous solution of $Na_2SO_3$. Organic fraction was separated and aqueous fraction was extracted with additional DCM (3 x ~150 mL). Combined organic fractions were dried with solid $Na_2SO_4$ and purified by FCC to obtain 330 mg (24% yield) of methyl sulfone (**19**) as white amorphous solid.

**TLC:** *Rf* = 0.45 (EtOAc/*n*-Hexane = 1:1).

**$^1$H NMR** (400 MHz, $CDCl_3$) $\delta$ = 8.75 – 8.59 (m, 1H), 8.13 (d, *J*=5.8, 1H), 7.70 (s, 1H), 3.32 (s, 3H), 1.55 (s, 9H) ppm.

**$^{13}$C{$^1$H} NMR** (101 MHz, $CDCl_3$) $\delta$ = 165.2, 159.3, 158.7, 151.0, 110.8, 83.4, 39.1, 28.0 ppm.

**HRMS** ($EI^+$) m/z: $[M + H]^+$, Calculated for $C_{10}H_{15}N_3O_4NaS^+$ 296.0675, found 296.0679.

##### *tert*-butyl (2-(methylthio)-6-propylpyrimidin-4-yl)carbamate (24):

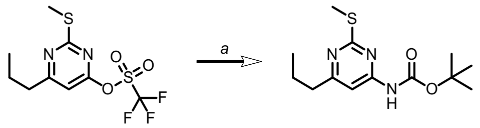


Triflate **23** (1.7 g, 5.42 mmol, 1 equiv.) was dissolved in anhydrous 1,4-dioxane (~50 mL) followed by addition of $Pd(OAc)_2$ (40 mg, 0.18 mmol, 0.033 equiv.), *tert*-Butyl carbamate (3.17 g, 27 mmol, 5 equiv.), XPhos (310 mg, 0.65 mmol, 0.12 equiv.) and $CsCO_3$ (5.3 g, 16.3 mmol, 3 equiv.). Reaction mixture was refluxed under inert nitrogen atmosphere for 4 h, let to cool down to room temperature and diluted with EtOAc (~200 mL) and washed with saturated solution of $NaHCO_3$, ammonium chloride and water (each ~50 mL). Organic fraction was dried with solid $Na_2SO_4$ and purified by FCC to provide 1.3 g of carbamate **24** in 85.5% yield, as a white solid.

**TLC:** *Rf* = 0.55 (EtOAc/*n*-Hexane = 1:9).

**$^1$H NMR** (400 MHz, $CDCl_3$) $\delta$ = 7.14 (s, 1H), 2.63 – 2.49 (m, 2H), 2.42 (s, 3H), 1.67 (d, *J*=7.7, 3H), 1.45 (s, 9H), 0.89 (s, 3H ppm.

**$^{13}$C{$^1$H} NMR** (101 MHz, $CDCl_3$) $\delta$ = 172.3, 170.9, 157.7, 151.6, 102.1, 82.0, 40.0, 28.2, 22.1, 14.0, 13.9 ppm.

**HRMS** ($EI^+$) m/z: $[M + H]^+$, Calculated for $C_{13}H_{22}N_3O_2S^+$ 284.14.27, found 284.1427.

##### *tert*-butyl (2-(methylsulfonyl)-6-propylpyrimidin-4-yl)carbamate (25):

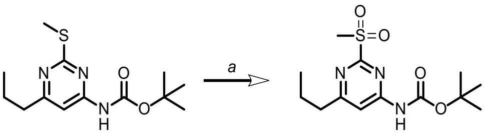


Carbamate **24** (0.4 g, 1.4 mmol, 1 equiv.) was dissolved in DCM (~50 mL) and cooled to 0 °C. mCPBA (1.94 g, 5.6 mmol, 4 equiv., 50% purity) was added in portions. Reaction mixture was let to reach room temperature over 16 h. After cooling of reaction mixture to 0 °C, remaining mCPBA was quenched by addition of aqueous solution of $Na_2SO_3$ (careful addition!). Mixture was diluted with additional DCM (~50 mL) and washed with saturated solution of $NaHCO_3$ (2x ~30 mL). Organic fraction was dried with solid $Na_2SO_4$ and purified by FCC to obtain white solid product (149 mg, 49.5% yield).

**TLC:** *Rf* = 0.90 (EtOAc/*n*-Hexane = 1:1).

**$^1$H NMR** (400 MHz, $CDCl_3$) $\delta$ = 7.26 (s, 1H), 6.92 (s, 1H), 2.61 (s, 3H), 2.17 – 2.02 (m, 2H), 1.10 (d, *J*=7.6, 2H), 0.85 (s, 9H), 0.30 (s, 3H) ppm.

**$^{13}$C{$^1$H} NMR** (101 MHz, $CDCl_3$) $\delta$ = 174.2, 164.8, 159.1, 151.2, 109.0, 83.1, 39.9, 39.0, 28.1, 22.0, 13.7 ppm.

**HRMS** ($EI^+$) m/z: $[M + Na]^+$, Calculated for $C_{13}H_{21}N_3O_4NaS^+$ 338.1145, found 338.1146.

***tert*-butyl (6-chloro-2-(methylthio)pyrimidin-4-yl)carbamate (27):**

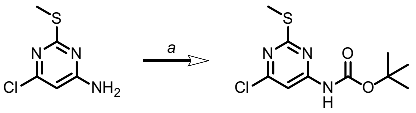


A solution of methylthiopyrimidine **26** (1.0 g, 5.69 mmol, 1 equiv.), $Et_3N$ (1.15 g, 1.58 mL, 2 equiv.) and $Boc_2O$ (1.24 g, 5.69 mmol, 1 equiv.) in THF (~30 mL) was refluxed for 2 days. The reaction mixture was cooled to room temperature, volatiles evaporated and the residue purified by automated FCC using prepacked 20 g column (isocratic elution) with 10% EtOAc in hexane to yield the protected amine pyrimidine **27** (1.57 g, 37% yield) as yellowish oil, which solidified upon storage at 4 °C.

**TLC:** *Rf* = 0.91 (EtOAc/*n*-Hexane = 2:8).

**$^1$H NMR** (400 MHz, $CDCl_3$) $\delta$ = 7.63 (s, 1H), 2.50 (s, 3H), 1.52 (s, 9H) ppm.

**$^{13}$C{$^1$H} NMR** (101 MHz, $CDCl_3$) $\delta$ = 172.5, 161.8, 158.5, 151.3, 103.1, 82.9, 28.2, 14.3 ppm.

**HRMS** ($EI^+$) m/z: $[M + H]^+$, Calculated for $C_{10}H_{15}ClN_3O_2S^+$ 276.0568, found 276.0569.

***tert*-butyl (6-chloro-2-(methylsulfonyl)pyrimidin-4-yl)carbamate (28):**

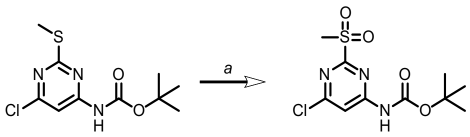


Carbamate **27** (0.56 g, 2.03 mmol, 1 equiv.) was dissolved in DCM (~80 mL) and cooled to 0 °C. mCPBA (3.50 g, 10.154 mmol, 5 equiv., 50% purity) was added in portions. Reaction mixture was let to reach room temperature over 16 h. After cooling of reaction mixture to 0 °C, remaining mCPBA was quenched by addition of aqueous solution of $Na_2SO_3$ (careful addition!). Mixture was diluted with additional DCM (~100 mL) and washed with saturated solution of $NaHCO_3$ (2x ~30 mL). Organic fraction was dried with solid $Na_2SO_4$ and purified by FCC to obtain white solid product (343 mg, 55% yield).

**TLC:** *Rf* = 0.85 (EtOAc/*n*-Hexane = 1:1).

**$^1$H NMR** (400 MHz, $CDCl_3$) $\delta$ = 8.19 (s, 1H), 7.71 (s, 1H), 3.34 (s, 2H), 1.57 (d, *J*=6.4, 9H) ppm.

**$^{13}$C{$^1$H} NMR** (101 MHz, $CDCl_3$) $\delta$ = 110.3, 77.2, 39.0, 28.0 ppm.

**HRMS** ($EI^+$) m/z: $[M + Na]^+$, Calculated for $C_{10}H_{14}N_3O_4ClSNa^+$ 330.0286, found 330.0286.

**6-methyl-2-(methylthio)pyrimidin-4-yl trifluoromethanesulfonate (30):**

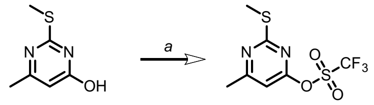


Methyl thiopyrimidine **29** (1 g, 6.4 mmol, 1 equiv.) and DIPEA (1.98 g, 15.36 mmol, 2.5 mL, 2.4 equiv.) was dissolved in DCM (~50 mL) and cooled to 0 °C. Triflic anhydride (2.1 g, 7.7 mmol, 1.2 equiv.) was added and reaction mixture was allowed to reach room temperature for 4 h. After full conversion was confirmed by TLC, reaction was diluted with additional 50 mL of DCM and washed with saturated aqueous solution of $NaHCO_3$ (2 x ~30 mL). Organic fraction was dried with solid $Na_2SO_4$ and purified by FCC to provide triflate 1.8 g **30** in 97% yield, as a yellow oil.

**TLC:** *Rf* = 0.45 (EtOAc/*n*-Hexane = 5:95).

**$^1$H NMR** (400 MHz, $CDCl_3$) $\delta$ = 6.56 (s, 1H), 2.46 (s, 6H) ppm.

**$^{13}$C{$^1$H} NMR** (101 MHz, $CDCl_3$) $\delta$ = 174.0, 173.9, 172.5, 172.5, 162.7, 123.2, 120.0, 116.8, 113.7, 104.8, 24.2, 24.2, 14.1 pm.

**$^{19}$F{$^1$H}** (376 MHz, $CDCl_3$) $\delta$ = -73.0 ppm.

**HRMS** ($EI^+$) m/z: $[M + H]^+$, Calculated for $C_7H_8F_3N_2O_3S_2^+$ 288.9923, found 288.9925.

***tert*-butyl (6-methyl-2-(methylthio)pyrimidin-4-yl)carbamate (31):**

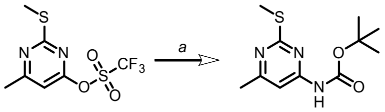


Triflate **30** (2 g, 7.16 mmol, 1 equiv.) was dissolved in anhydrous 1,4-dioxane (~60 mL) followed by addition of $Pd(OAc)_2$ (56 mg, 0.25 mmol, 0.035 equiv.), *tert*-Butyl carbamate (1.67 g, 13.3 mmol, 2 equiv.), XPhos (409 mg, 0.85 mmol, 0.12 equiv.) and $CsCO_3$ (3.5 g, 10.74 mmol, 1.5 equiv.).

Reaction mixture was refluxed under inert nitrogen atmosphere for 4 h, let to cool down to room temperature and diluted with EtOAc (~200 mL) and washed with saturated solution of $NaHCO_3$, ammonium chloride and water (each ~50 mL). Organic fraction was dried with solid $Na_2SO_4$ and purified by FCC to provide 1.18 g of carbamate **31** in 66% yield, as a white to yellowish solid.

**TLC:** *Rf* = 0.42 (EtOAc/*n*-Hexane = 10:90).

**$^1$H NMR** (400 MHz, $CDCl_3$) δ = 7.37 (s, 1H), 7.09 (s, 1H), 2.43 (s, 3H), 2.35 (s, 3H), 1.45 (s, 9H) ppm.

**$^{13}$C{$^1$H} NMR** (101 MHz, $CDCl_3$) *δ* = 171.0, 168.4, 157.7, 151.6, 102.6, 82.2, 28.1, 24.2, 14.0 ppm.

**HRMS** (EI$^+$) m/z: $[M + H]^+$, Calculated for $C_{11}H_{18}N_3O_2S^+$ 256.1114, found 256.1115.

**_tert_-butyl (6-methyl-2-(methylsulfonyl)pyrimidin-4-yl)carbamate (32):**

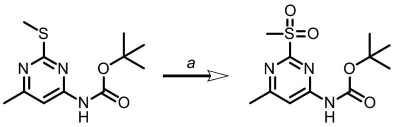


Carbamate **31** (0.44 g, 1.72 mmol, 1 equiv.) was dissolved in DCM (~50 mL) and cooled to 0 °C. mCPBA (2.2 g, 6.89 mmol, 4 equiv., 50% purity) was added in portions. Reaction mixture was let to reach room temperature over 16 h. After cooling of reaction mixture to 0 °C, remaining mCPBA was quenched by addition of aqueous solution of $Na_2SO_3$ (careful addition!). Mixture was diluted with additional DCM (~150 mL) and washed with saturated solution of $NaHCO_3$ (3x ~30 mL). Organic fraction was dried with solid $Na_2SO_4$ and purified by FCC to obtain white solid product **32** (149 mg, 41% yield).

**TLC:** *Rf* = 0.72 (EtOAc/*n*-Hexane = 1:1).

**$^1$H NMR** (400 MHz, DMSO) *δ* = 10.88 (s, 1H), 7.89 (d, *J*=0.7, 1H), 3.32 (s, 3H), 2.52 (s, 3H), 1.48 (s, 9H) ppm.

**$^{13}$C{$^1$H} NMR** (101 MHz, DMSO) *δ* = 169.5, 164.6, 159.7, 152.1, 109.4, 81.3, 27.8, 23.9 ppm.

**HRMS** (EI$^+$) m/z: $[M + H]^+$, Calculated for $C_{11}H_{17}N_3NaO_4S^+$ 310.0832, found 310.0834.

**6-methyl-2-(methylthio)pyrimidin-4-ol (34):**

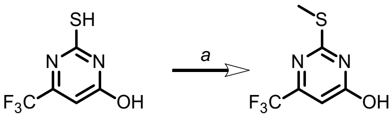


Methylthio-pyrimidine (**34**) was synthesized from thio-pyrimidine (**33**) following previously described procedure. Spectral data corresponds to the previously published.[18]

**$^1$H NMR** (400 MHz, $CDCl_3$) *δ* = 12.65 (d, *J*=18.6, 1H), 6.60 (s, 1H), 2.66 (s, 3H) ppm.

$^{13}$**C{$^1$H} NMR** (101 MHz, $CDCl_3$) $\delta$ = 164.7, 164.2, 153.7, 153.4, 121.6, 118.8, 108.3, 108.3, 108.3, 108.2, 13.5 ppm.

**2-(methylthio)-6-(trifluoromethyl)pyrimidin-4-yl trifluoromethanesulfonate (35):**

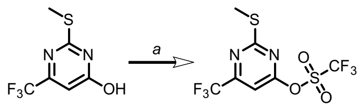


Methyl thiopyrimidine **34** (0.21 g, 1.9 mmol, 1 equiv.) and DIPEA (0.3 g, 2.4 mmol, 0.4 mL, 2.4 equiv.) was dissolved in DCM (~15 mL) and cooled to 0 °C. Triflic anhydride (0.33 g, 1.2 mmol, 1.2 equiv.) was added and reaction mixture was allowed to reach room temperature for 4 h. After full conversion was confirmed by TLC, reaction was diluted with additional 50 mL of DCM and washed with saturated aqueous solution of NaHCO3 (2 x ~15 mL). Organic fraction was dried with solid $Na_2SO_4$ and purified by FCC to provide triflate **35** in 64% yield, as colorless oil (220 mg).

**TLC:** *Rf* = 0.45 (EtOAc/*n*-Hexane = 5:95).

$^1$**H NMR** (400 MHz, $CDCl_3$) $\delta$ = 7.07 (s, 1H), 2.64 (s, 3H) ppm.

$^{13}$**C{$^1$H} NMR** (101 MHz, $CDCl_3$) $\delta$ = 176.5, 163.3, 160.0, 159.6, 123.5, 123.2, 120.8, 120.0, 118.0, 116.8, 115.3, 113.6, 101.8, 101.8, 14.5 ppm.

**19F{$^1$H}** (376 MHz, $CDCl_3$) $\delta$ = -70.2, -72.6 ppm.

**HRMS** ($EI^+$) m/z: $[M + H]^+$, Calculated for $C_7H_5N_2O_3F_6S_2^+$ 342.9640, found 342.9641.

***tert*-butyl (2-(methylthio)-6-(trifluoromethyl)pyrimidin-4-yl)carbamate (36):**

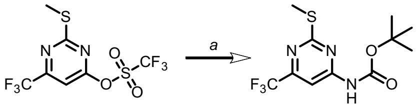


Triflate **35** (0.06 g, 0.175 mmol, 1 equiv.) was dissolved in anhydrous 1,4-dioxane (~5 mL) followed by addition of Tris(dibenzylideneacetone)dipalladium(0) (32 mg, 0.035 mmol, 0.2 equiv.), *tert*-Butyl carbamate (30 mg, 0.263 mmol, 1.5 equiv.), XPhos (33 mg, 0.07 mmol, 0.4 equiv.) and $CsCO_3$ (0.228 g, 0.7 mmol, 4 equiv.). Reaction mixture was refluxed under inert nitrogen atmosphere for 4 h, let to cool down to room temperature and diluted with EtOAc (~50 mL) and washed with saturated solution of $NaHCO_3$, ammonium chloride and water (each ~15 mL). Organic fraction was dried with solid $Na_2SO_4$ and purified by FCC to provide 43 mg of carbamate **36** in 80% yield, as a white to brownish solid.

$^1$**H NMR** (400 MHz, $CDCl_3$) $\delta$ = 7.95 (s, 1H), 2.56 (s, 3H), 1.56 (s, 8H) ppm.

**$^{13}C\{^1H\}$ NMR** (101 MHz, $CDCl_3$) *δ* = 173.3, 159.1, 156.8, 156.4, 151.1, 121.8, 119.1, 100.0, 99.9, 99.9, 99.9, 83.1, 28.1, 14.1 ppm.

**$^{19}F$ NMR** (376 MHz, $CDCl_3$) δ = -70.4 ppm.

**HRMS** ($EI^+$) m/z: $[M + H]^+$, Calculated for $C_{11}H_{15}F_3N_3O_2S^+$ 310.0832, found 310.0833.

***tert*-butyl (2-(methylsulfonyl)-6-(trifluoromethyl)pyrimidin-4-yl)carbamate (37):**

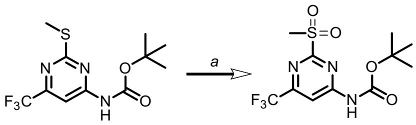


Carbamate **36** (0.1 g, 0.32 mmol, 1 equiv.) was dissolved in DCM (~30 mL) and cooled to 0 °C. mCPBA (0.24 g, 0.711 mmol, 2.2 equiv., 50% purity) was added in portions. Reaction mixture was let to reach room temperature over 16 h. After cooling of reaction mixture to 0 °C, remaining *m*CPBA was quenched by addition of aqueous solution of $Na_2SO_3$ (careful addition!). Mixture was diluted with additional DCM (~100 mL) and washed with saturated solution of $NaHCO_3$ (3x ~30 mL). Organic fraction was dried with solid $Na_2SO_4$ and purified by FCC to obtain white solid product **37** (110 mg, quant.).

**TLC:** *Rf* = 0.45 (EtOAc/*n*-Hexane = 3:7).

**$^1H$ NMR** (400 MHz, $CDCl_3$) *δ* = 8.51 (s, 1H), 7.95 (s, 1H), 3.39 (s, 3H), 1.58 (s, 9H) ppm.

**$^{13}C\{^1H\}$ NMR** (101 MHz, $CDCl_3$:MeOD = 9:1) *δ* = 167.4, 165.4, 162.2, 157.2, 156.9, 156.5, 156.1, 151.7, 134.3, 132.8, 132.3, 129.8, 129.6, 127.9, 121.3, 118.5, 107.5, 107.5, 107.4, 107.4, 83.4, 83.4, 38.7, 27.8 ppm.

**$^{19}F$ NMR** (376 MHz, $CDCl_3$) δ = -66.17 ppm.

**HRMS** ($EI^+$) m/z: $[M + H]^+$, Calculated for $C_{11}H_{15}N_3O_4F_3S^+$ 342.0730, found 342.0731.

##### 4.1.2 Building blocks – benzyl alcohol

##### 4.1.3 Conjugate precursors

***tert*-butyl (4-(((6-aminopyridin-2-yl)oxy)methyl)benzyl)carbamate (39):**

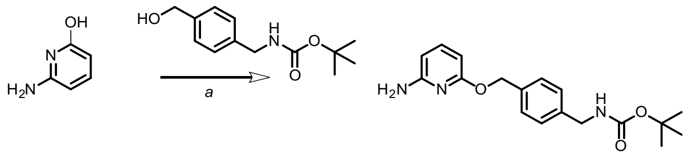

Amino pyridine **38** (0.05 g, 0.45 mmol, 1 equiv.) was dissolved in THF (5 mL) followed by addition of triphenylphosphine (238 mg, 908 µmol, 2 equiv.), **43** (140 mg, 590 µmol, 1.3 equiv.) and diisopropyl azodicarboxylate (180 µL, 908 µmol, 2 equiv.). The reaction mixture was stirred for 16 h at room temperature. Reaction mixture was dilluted with EtOAc (50 mL), and washed with saturated solution of $NaHCO_3$ (3 x ~20 mL), organic fraction was dried with solid anhydrous $Na_2SO_4$, volatiles removed under reduced pressure and purified by with preparative HPLC to provide 19 mg of carbamate **39** as yellowish oil (13% yield).

**TLC:** *Rf* = 0.80 (EtOAc/*n*-Hexane = 7:3).

**$^1$H NMR** (400 MHz, $CD_3OD$) *δ* = 7.8 (t, J = 8.3 Hz, 1H), 7.5 – 7.2 (m, 4H), 6.4 (dd, J = 38.0, 8.3 Hz, 2H), 5.3 (s, 2H), 4.2 (s, 2H), 1.5 (s, 9H) ppm.

**$^{13}$C NMR** (101 MHz, $CD_3OD$) *δ* = 157.2, 154.7, 145.8, 140.6, 133.3, 128.0, 127.1, 103.3, 93.1, 78.9, 71.2, 43.3, 27.4, 20.9 ppm.

**HRMS** ($EI^+$) m/z: $[M + H]^+$, Calculated for $C_{18}H_{24}N_3O_3^+$ 330.1812, found 330.1813.

***tert*-butyl (4-(((6-aminopyridin-2-yl)oxy)methyl)benzyl)carbamate (41):**

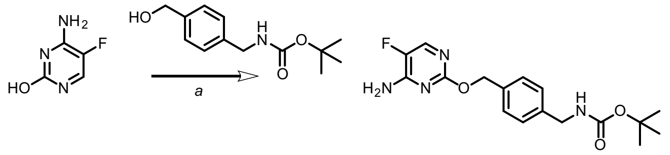

Carbamate (**41**) was synthesized like described for (**39**), purified by automated FCC using 20 g prepacked column (gradient 20% EtOAc in *n*-Hexane to 100% EtOAc, over 20 volumes of column) and 52 mg as white amorphous solid was obtained (23% yield). Unwanted *N*-alkylation side product of **40** was removed during purification.

**TLC:** *Rf* = 0.83 (EtOAc/*n*-Hexane = 7:3).

**$^1$H NMR** (400 MHz, $CDCl_3$) *δ* = 7.9 (d, J = 2.6 Hz, 1H), 7.5 – 7.4 (m, 2H), 7.3 (d, J = 3.5 Hz, 3H), 5.4 – 5.3 (m, 2H), 4.3 (d, J = 5.8 Hz, 2H), 1.5 (s, 8H) ppm.

**$^{13}$C NMR** (101 MHz, $CDCl_3$) *δ* = 160.1, 155.9, 154.5 (d, *J* = 13.4 Hz), 142.5 (d, *J* = 246.9 Hz), 140.8, 140.6, 138.6, 135.8, 128.1, 127.5, 79.6, 70.1, 68.9, 44.5, 28.4 ppm.

**$^{19}$F NMR** (376 MHz, $CDCl_3$) $\delta$ = -165.7 ppm.

**HRMS** ($EI^+$) m/z: $[M + H]^+$, Calculated for $C_{17}H_{22}FN_4O_3^+$ 349.1670, found 349.1674.

***tert*-butyl (4-(((4-aminopyridin-2-yl)oxy)methyl)benzyl)carbamate (52):**

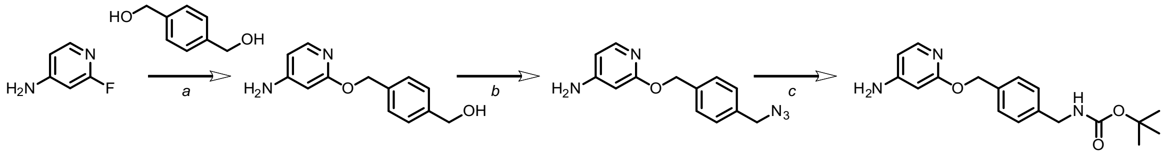

a) Bis-alcohol (**42**, 0.145 g, 1.07 mmol, 1.1 equiv.) was dissolved in round bottom flask in mixture of 1,4-dioxane and THF (1:1, 20 mL), cooled to 10 °C and solid NaH (0.045 g, 60%, 1.2 equiv.) was added. After stirring for 30 min at the same temperature under $N_2$ atmosphere, fluoropyridine **49** (0.107 g, 0.95 mmol, 1 equiv.) was added and reaction mixture was heated to reflux (~100 °C, oil bath) and stirred for 16 h. After cooling down to room temperature, reaction was quenched with solid NH4Cl (~100 mg) and volatiles were removed under reduced pressure. Crude residue was dissolved in DCM (~100 mL) and washed with water (2 x ~30 mL) and organic fraction was dried with solid anhydrous $Na_2$SO4. After removing the volatiles under reduced pressure, alcohol **50** was used in the next step without further purification.

b) Alcohol **50** was dissolved in mixture of toluene and THF (4:1, 20 mL) at room temperature (~25 °C) in the round bottom flask. To the reaction mixture DPPA (0.525 g, 1.9 mmol, 2 equiv.) was added, followed by DBU (0.29 g, 1.9 mmol, 2 equiv.) and it was stirred under $N_2$ atmosphere for 16 h. Reaction mixture was diluted with EtOAc (~50 mL) and washed with saturated solution of $NaHCO_3$ (3 x ~20 mL), organic fraction was dried with solid anhydrous $Na_2$SO4, volatiles removed under reduced pressure and azide (**51**) was used in the next step without further purification.

c) Crude azide (**51**) was dissolved in THF (20 mL) and tributylphosphine (0.256 g, 1.3 mmol, 1.33 equiv.) was added in one portion at room temperature, while keeping round bottom flask open to reduce chance of over pressure in the flask caused by released gas. The reaction mixture was stirred for 2 h and water (3 mL) was added in one portion. After stirring for 16 h, NaOH (1M, 5 mL) was added at 0 °C followed by $Boc_2O$ (0.43 g, 2.0 mmol, 2.0 equiv.) and stirring was continued for another 12 h, letting the reaction mixture to reach room temperature. Reaction mixture was extracted with EtOAc (3 x 30 mL), organic fractions were combined and dried with solid anhydrous $Na_2SO_4$. After removing the volatiles under reduced pressure, automated FCC using 20 g prepacked column (gradient 20% EtOAc in *n*-Hexane to 100% EtOAc, over 20 volumes of column) provided 95 mg of carbamate **52** as white amorphous solid (29.5% yield, over 3 steps).

**TLC:** *Rf* = 0.74 (EtOAc/*n*-Hexane = 7:3).

**$^1$H NMR** (400 MHz, $CDCl_3$) $\delta$ = 7.68 (d, *J*=5.7, 1H), 7.26 (d, *J*=8.1, 2H), 7.14 (d, *J*=7.9, 2H), 6.07 (dd, *J*=5.8, 2.0, 1H), 5.86 (d, *J*=2.0, 1H), 5.17 (s, 2H), 4.17 (d, *J*=6.2, 4H), 1.35 (s, 9H) ppm.

**$^{13}$C NMR** (101 MHz, $CDCl_3$) δ = 165.0, 156.0, 155.5, 147.0, 138.4, 136.7, 128.1, 127.4, 105.8, 93.9, 79.4, 67.1, 44.4, 28.4.

**HRMS** ($EI^+$) m/z: $[M + H]^+$, Calculated for $C_{18}H_{24}N_3O_3^+$ 330.1812, found 330.1811.

***tert*-butyl (4-(((4-amino-6-fluoropyridin-2-yl)oxy)methyl)benzyl)carbamate (48):**

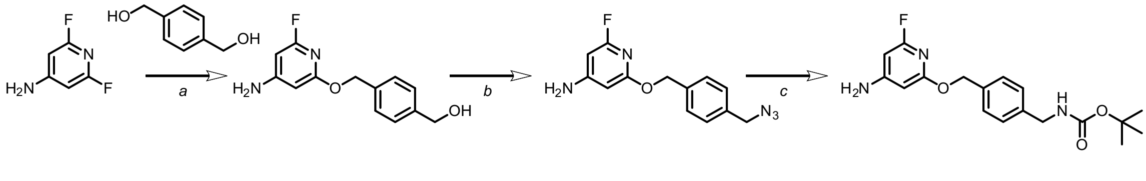

Carbamate (**48**) was synthesized following the procedure described for **52** and as a product white amorphous solid was obtained (99 mg, 29%, over 3 steps). For FCC, 20 g prepacked column was used, gradient 20% EtOAc in *n*-Hexane to 100% EtOAc, over 20 volumes of column. Additionally, product was purified with preparative HPLC.

**TLC:** *Rf* = 0.38 (EtOAc/*n*-Hexane = 3:7).

**$^{1}$H NMR** (400 MHz, $CDCl_3$) $\delta$ = 7.39 (d, *J*=7.9, 2H), 7.33 – 7.21 (m, 2H), 5.84 (t, *J*=1.3, 1H), 5.77 (d, *J*=1.6, 1H), 5.25 (s, 2H), 4.31 (d, *J*=4.5, 2H), 1.47 (s, 9H) ppm.

**$^{13}$C{$^{1}$H} NMR** (101 MHz, $CDCl_3$) $\delta$ = 164.8, 163.7, 163.5, 162.5, 158.7, 158.5, 156.1, 138.6, 136.2, 128.3, 127.5, 91.1, 91.0, 87.5, 87.1, 79.7, 67.7, 44.4, 28.4 ppm.

**$^{19}$F NMR** (376 MHz, $CDCl_3$) $\delta$ = -72.86 ppm.

**HRMS** ($EI^+$) m/z: $[M + H]^+$, Calculated for $C_{18}H_{23}N_3O_3F^+$ 348.1718, found 348.1715.

***tert*-butyl (4-(((4-amino-6-chloropyridin-2-yl)oxy)methyl)benzyl)carbamate (56):**

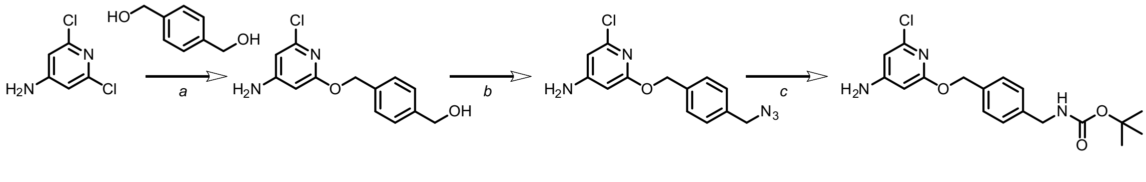

Carbamate (**56**) was synthesized following the procedure described for **52**, and as a product white amorphous solid was obtained (136 mg, 26%, over 3 steps). For FCC, 20 g prepacked column was used, gradient 20% EtOAc in *n*-Hexane to 100% EtOAc, over 20 volumes of column.

**TLC:** *Rf* = 0.45 (EtOAc/*n*-Hexane = 3:7).

**$^{1}$H NMR** (400 MHz, $CDCl_3$) $\delta$ = 7.45 – 7.38 (m, 2H), 7.30 (s, 1H), 6.26 (d, $J$=1.7, 1H), 5.91 (d, $J$=1.7, 1H), 5.29 (s, 2H), 4.45 – 4.21 (m, 2H), 1.48 (s, 9H) ppm.

**$^{13}$C{$^{1}$H} NMR** (101 MHz, $CDCl_3$) $\delta$ = 164.2, 156.7, 156.0, 148.8, 138.6, 136.1, 128.4, 127.6, 104.1, 92.6, 79.7, 67.9, 44.5, 28.4 ppm.

**HRMS** ($EI^+$) m/z: $[M + H]^+$, Calculated for $C_{18}H_{23}N_3O_3Cl^+$ 364.1422, found 364.1420.

***tert*-butyl (4-(((4-amino-6-(trifluoromethyl)pyrimidin-2-yl)oxy)methyl)benzyl)carbamate (57):**

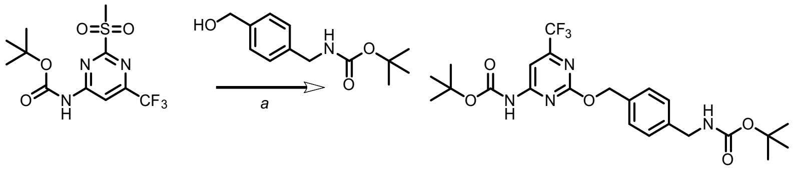

Benzyl alcohol **43** (52 mg, 0.219 mmol, 1.5 equiv.) was dissolved in THF (5 mL) and cooled down to 0 °C. Sodium hydride (9 mg, 0.234 mmol, 1.6 equiv.) was added and suspension was stirred at the constant temperature for 30 min. Methyl sulfone **37** (50 mg, 0.146 mmol, 1 equiv.) was added in one portion and reaction mixture was let to reach room temperature over 16 h. Remaining unreacted hydride and alkoxide was quenched by addition of solid $NH_4Ac$ (~30 mg), volatiles were removed under reduced pressure and mixture was purified by automated FCC, 20 g prepacked column was used, gradient 10% EtOAc in *n*-Hexane to 50% EtOAc, over 20 volumes of column. Solid white product was obtained (70 mg, 95% yield).

**TLC:** *Rf* = 0.66 (EtOAc/*n*-Hexane = 3:7).

**$^{1}$H NMR** (400 MHz, $CDCl_3$) $\delta$ = 7.96 (s, 1H), 7.62 (s, 1H), 7.43 (d, $J$=7.8, 2H), 7.28 (d, $J$=4.5, 2H), 5.40 (s, 2H), 4.32 (s, 2H), 1.55 (s, 8H), 1.47 (s, 9H) ppm.

**$^{13}$C{$^{1}$H} NMR** (101 MHz, $CDCl_3$) $\delta$ = 164.7, 161.6, 158.5, 158.1, 155.9, 151.1, 139.1, 134.8, 128.7, 127.5, 124.5, 121.7, 119.0, 116.3, 99.0, 83.1, 69.4, 44.4, 28.4, 28.1 ppm.

**$^{19}$F NMR** (376 MHz, $CDCl_3$) $\delta$ = -70.32 ppm.

**HRMS** ($EI^+$) m/z: $[M + H]^+$, Calculated for $C_{23}H_{30}F_3N_4O_5^+$ 499.2163, found 499.2166.

***tert*-butyl (4-(((4-amino-6-propylpyrimidin-2-yl)oxy)methyl)benzyl)carbamate (58):**

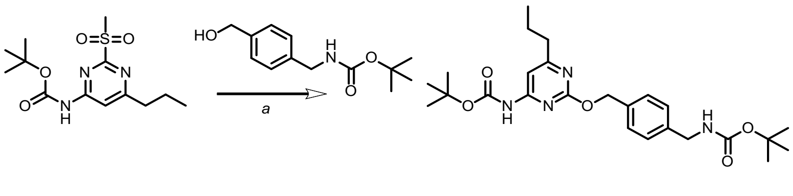

Benzyl alcohol **43** (75 mg, 0.317 mmol, 2.5 equiv.) was dissolved in THF (5 mL) and cooled down to 0 °C. Sodium hydride (13 mg, 0.317 mmol, 2.5 equiv.) was added and suspension was stirred at the constant temperature for 30 min. Methyl sulfone **25** (40 mg, 0.126 mmol, 1 equiv.) was added in one portion and reaction mixture was let to reach room temperature over 16 h. Remaining unreacted hydride and alkoxide was quenched by addition of solid $NH_4Ac$ (~30 mg), volatiles were removed under reduced pressure and mixture was purified by automated FCC, 20 g prepacked column was used, gradient 10% EtOAc in *n*-Hexane to 50% EtOAc, over 20 volumes of column. Solid white product was obtained (52 mg, 87% yield).

**TLC:** *Rf* = 0.80 (EtOAc/*n*-Hexane = 3:7).

$^{1}$**H NMR** (400 MHz, $CDCl_3$) $\delta$ = 7.50 (s, 1H), 7.39 (s, 1H), 7.31 (s, 2H), 7.19 (s, 2H), 5.27 (s, 2H), 4.22 (d, *J*=4.6, 2H), 2.58 (s, 2H), 1.77 – 1.59 (m, 2H), 1.45 (s, 9H), 1.38 (s, 9H), 0.89 (t, *J*=7.4, 3H) ppm.

$^{13}$**C{$^{1}$H} NMR** (101 MHz, $CDCl_3$) $\delta$ = 174.1, 163.4, 159.9, 156.0, 151.4, 138.8, 135.4, 129.0, 128.2, 127.5, 125.3, 100.8, 82.4, 79.6, 68.7, 44.4, 39.4, 28.4, 28.1, 22.0, 13.8 ppm.

**HRMS** ($EI^+$) m/z: $[M + H]^+$, Calculated for $C_{25}H_{37}N_4O_5^+$ 473.2758, found 473.2760.

***tert*-butyl (4-(((4-amino-6-chloropyrimidin-2-yl)oxy)methyl)benzyl)carbamate (59):**

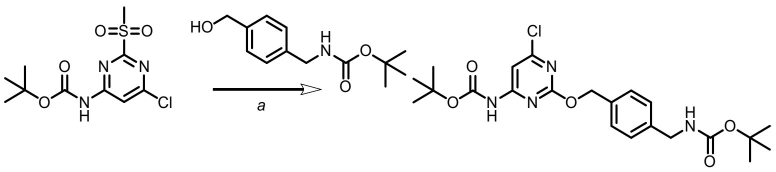


Benzyl alcohol **43** (100 mg, 0.422 mmol, 1.3 equiv.) was dissolved in THF (5 mL) and cooled down to 0 °C. Sodium hydride (17 mg, 0.422 mmol, 1.3 equiv.) was added and suspension was stirred at the constant temperature for 30 min. Methyl sulfone **28** (100 mg, 0.325 mmol, 1 equiv.) was added in one portion and reaction mixture was let to reach room temperature over 16 h. Remaining unreacted hydride and alkoxide was quenched by addition of solid $NH_4Ac$ (~50 mg), volatiles were removed under reduced pressure and mixture was purified by automated FCC, 20 g prepacked column was used, gradient 10% EtOAc in *n*-Hexane to 50% EtOAc, over 20 volumes of column. Solid white product was obtained (140 mg, 93% yield).

**TLC:** *Rf* = 0.65 (EtOAc/*n*-Hexane = 3:7).

$^{1}$**H NMR** (400 MHz, $CDCl_3$) $\delta$ = 7.65 (s, 1H), 7.41 (s, 2H), 7.30 (s, 1H), 5.37 (s, 2H), 4.33 (d, *J*=5.9, 2H), 1.55 (s, 9H), 1.48 (s, 9H) ppm.

$^{13}$**C{$^{1}$H} NMR** (101 MHz, $CDCl_3$) $\delta$ = 163.8, 163.0, 160.3, 155.9, 151.1, 139.0, 135.0, 128.4, 127.6, 101.7, 82.8, 79.6, 69.3, 44.4, 28.4, 28.1 ppm.

**HRMS** ($EI^+$) m/z: $[M + H]^+$, Calculated for $C_{22}H_{30}N_4O_5Cl^+$ 465.1899, found 465.1901.

***tert*-butyl (4-(((4-amino-6-chloropyrimidin-2-yl)oxy)methyl)benzyl)carbamate (60):**

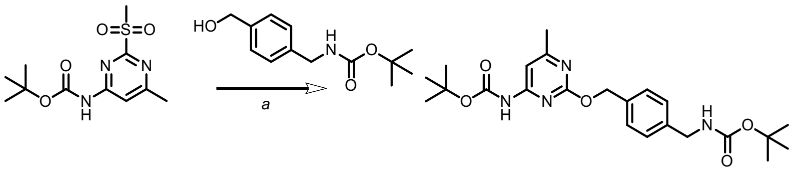

Benzyl alcohol **43** (21 mg, 0.1 mmol, 1.3 equiv.) was dissolved in THF (5 mL) and cooled down to 0 °C. Sodium hydride (4 mg, 0.09 mmol, 1.3 equiv.) was added and suspension was stirred at the constant temperature for 30 min. Methyl sulfone **32** (20 mg, 0.069 mmol, 1 equiv.) was added in one portion and reaction mixture was let to reach room temperature over 16 h. Remaining unreacted hydride and alkoxide was quenched by addition of solid $NH_4Ac$ (~10 mg), volatiles were removed under reduced pressure and mixture was purified by automated FCC, 20 g prepacked column was used, gradient 10% EtOAc in *n*-Hexane to 50% EtOAc, over 20 volumes of column. Solid white product was obtained (19 mg, 63% yield).

**TLC:** *Rf* = 0.52 (EtOAc/*n*-Hexane = 3:7).

**$^1$H NMR** (400 MHz, $CDCl_3$) $\delta$ = 7.58 (s, 1H), 7.39 (s, 2H), 7.30 (s, 1H), 7.28 (s, 1H), 5.40 (s, 2H), 4.32 (s, 2H), 2.53 (s, 3H), 1.56 (s, 9H), 1.48 (s, 9H) ppm.

**$^{13}$C{$^1$H} NMR** (101 MHz, $CDCl_3$) $\delta$ = 168.3, 161.7, 160.7, 156.1, 150.9, 139.1, 134.4, 128.2, 127.6, 101.4, 83.3, 69.6, 44.4, 28.4, 28.0, 27.9, 22.3 ppm.

**HRMS** ($EI^+$) m/z: $[M + H]^+$, Calculated for $C_{23}H_{33}N_4O_5^+$ 445.2445, found 445.2441.

***tert*-butyl (4-(((4-((tert-butoxycarbonyl)amino)-6-(2,2,2-trifluoroethoxy)pyrimidin-2-yl)oxy)-methyl)-benzyl)carbamate (61):**

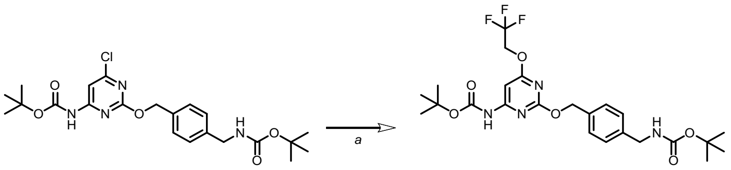

Trifluoroethanol (6 mg, 4 µL, 0.055 mmol, 1.3 equiv.) was added to anhydrous THF (~5 mL) and cooled to 0 °C. Solid NaH (2.2 mg, 0.055 mmol, 1.3 equiv., 60%) was added and stirred at the same temperature for 30 min. Chloride **59** (20 mg, 0.04 mmol, 1 equiv.) was added and reaction mixture was heated at 70 °C in a sealed vial over 16 h. After letting to cool to room temperature, unreacted alkoxide was quenched by addition of solid $NH_4Ac$ (~5 mg), volatiles were removed under reduced pressure and purified by automated FCC to give 8 mg of white solid product **61** (35% yield).

**TLC:** *Rf* = 0.63 (EtOAc/*n*-Hexane = 3:7).

**$^1$H NMR** (400 MHz, $CDCl_3$) $\delta$ = 7.41 – 7.27 (m, 2H), 7.22 (s, 1H), 7.14 (s, 1H), 7.00 (s, 1H), 5.24 (s, 2H), 4.66 (q, *J*=8.4, 2H), 4.24 (d, *J*=5.8, 2H), 1.45 (s, 9H), 1.39 (s, 9H) ppm.

**$^{13}C\{^{1}H\}$ NMR** (101 MHz, $CDCl_3$) *δ* = 170.9, 163.5, 160.3, 155.9, 151.3, 139.0, 135.3, 128.2, 127.6, 124.5, 121.8, 87.3, 82.3, 79.6, 68.9, 62.6, 62.3, 44.4, 28.4, 28.1 ppm.

**$^{19}F$ NMR** (376 MHz, $CDCl_3$) *δ* = -73.9 ppm.

**HRMS** ($EI^+$) m/z: $[M + H]^+$, Calculated for $C_{24}H_{32}N_4O_6F_3^+$ 529.2268, found 529.2273.

***tert*-butyl (4-(((4-((tert-butoxycarbonyl)amino)pyrimidin-2-yl)oxy)methyl)-2-fluorobenzyl)carbamate (62):**

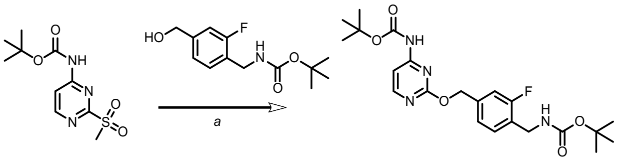

Carbamate **62** was synthesized following procedure described above for **57** from alcohol **44** and methylsulfone cytosine **19**. White solid product (10 mg, 20.3% yield) was obtained after purification by automated FCC.

**TLC:** *Rf* = 0.52 (EtOAc/*n*-Hexane = 3:7).

**$^{1}H$ NMR** (400 MHz, $CDCl_3$) *δ* = 8.41 (d, *J*=6.0, 1H), 7.64 (d, *J*=5.9, 1H), 7.33 (t, *J*=7.7, 1H), 7.22 – 7.09 (m, 2H), 5.37 (s, 2H), 4.36 (s, 2H), 1.55 (s, 9H), 1.46 (s, 9H) ppm.

**$^{13}C\{^{1}H\}$ NMR** (101 MHz, $CDCl_3$) *δ* = (101 MHz, $CDCl_3$) *δ* = 163.3, 162.1, 160.9, 160.5, 160.3, 160.3, 159.6, 158.6, 155.9, 151.1, 137.7, 137.6, 129.9, 125.8, 125.6, 123.3, 123.3, 114.7, 114.5, 102.6, 82.8, 79.8, 68.1, 68.1, 38.5, 28.4, 28.3, 28.1 ppm.

**19F NMR** (376 MHz, CDCl3) *δ* = -118.85 (t, *J*=9.2) ppm.

**HRMS** ($EI^+$) m/z: $[M + H]^+$, Calculated for $C_{22}H_{30}FN_4O_5^+$ 449.2195, found 449.2195.

***tert*-butyl (2-fluoro-4-(((4-iodopyridin-2-yl)oxy)methyl)benzyl)carbamate (64):**

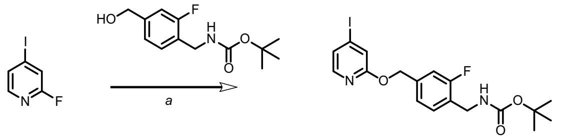

Iodide 64 was synthesized following procedure described above for **57** from alcohol **44** and fluoro-iodo-pyridine **63**. Colorless to yellowish oily product was obtained, which solidified upon storage in fridge (320 mg, 78% yield) was obtained after purification by automated FCC.

**TLC:** *Rf* = 0.80 (EtOAc/*n*-Hexane = 3:7).

$^1$H NMR (400 MHz, $CDCl_3$) $\delta$ = 7.75 (dd, $J$=5.3, 0.7, 1H), 7.39 – 7.13 (m, 4H), 7.13 – 6.98 (m, 2H), 5.25 (d, $J$=2.2, 2H), 4.28 (d, $J$=6.0, 2H), 1.37 (s, 9H) ppm.

**$^{13}$C{$^1$H} NMR** (101 MHz, $CDCl_3$) $\delta$ = 163.6, 163.3, 162.1, 159.7, 155.8, 147.0, 138.7, 138.6, 136.0, 130.0, 129.9, 128.3, 127.6, 126.3, 126.1, 125.6, 125.4, 123.4, 120.7, 120.6, 114.8, 114.6, 106.5, 106.4, 79.7, 67.6, 66.8, 66.7, 38.5, 28.4, 28.4 ppm.

**$^{19}$F NMR** (376 MHz, $CDCl_3$) $\delta$ = -119.03 (d, $J$=18.9), -119.03 ppm.

**HRMS** ($EI^+$) m/z: $[M + H]^+$, Calculated for $C_{18}H_{21}FIN_2O_3^+$ 459.0575, found 459.0571.

***tert*-butyl (4-(((4-((tert-butoxycarbonyl)amino)pyridin-2-yl)oxy)methyl)-2-fluorobenzyl)carbamate (65):**

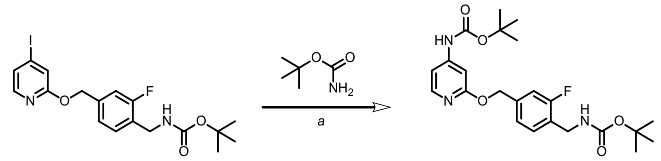


Iodide **64** (0.3 g, 0.654 mmol, 1 equiv.) was dissolved in anhydrous 1,4-dioxane (~25 mL) followed by addition of Tris(dibenzylideneacetone)dipalladium(0) (89 mg, 0.098 mmol, 0.15 equiv.), *tert*-Butyl carbamate (115 mg, 0.98 mmol, 1.5 equiv.), Xantphos (170 mg, 0.294 mmol, 0.45 equiv.) and $CsCO_3$ (853 g, 2.6 mmol, 4 equiv.). Reaction mixture was refluxed under inert nitrogen atmosphere for 3 h, let to cool down to room temperature and diluted with EtOAc (~100 mL) and washed with saturated solution of $NaHCO_3$, ammonium chloride and water (each ~30 mL). Organic fraction was dried with solid $Na_2SO_4$ and purified by FCC to provide 256 mg of bis-carbamate 65 in 87% yield, as a brownish solid.

**TLC:** $Rf$ = 0.65 (EtOAc/$n$-Hexane = 7:3).

**$^1$H NMR** (400 MHz, $CDCl_3$) $\delta$ = 7.97 (d, $J$=5.8, 1H), 7.29 (d, $J$=7.3, 2H), 7.13 (td, $J$=8.7, 1.6, 3H), 7.01 – 6.81 (m, 2H), 5.33 (d, $J$=2.0, 2H), 4.35 (d, $J$=6.0, 2H), 1.51 (s, 9H), 1.45 (s, 9H) ppm.

**$^{13}$C{$^1$H} NMR** (101 MHz, $CDCl_3$) $\delta$ = 164.5, 155.9, 152.0, 148.1, 148.0, 147.2, 147.2, 139.3, 139.3, 129.8, 129.7, 128.0, 123.1, 123.1, 114.5, 114.3, 107.6, 107.5, 98.4, 98.4, 81.5, 81.4, 79.6, 67.3, 66.4, 66.4, 38.5, 28.4, 28.4, 28.2, 28.2 ppm.

**19F NMR** (376 MHz, CDCl3) $\delta$ = -119.20 (t, $J$=9.2) ppm.

**HRMS** ($EI^+$) m/z: $[M + H]^+$, Calculated for $C_{23}H_{31}N_3O_5F^+$ 448.2242, found 448.2243.

##### 4.1.4 Fluorophore conjugates

**Conjugate 1:**

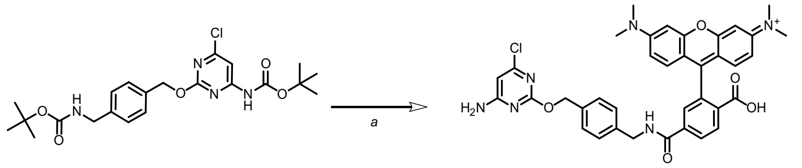


Active ester of TMR-6-COOH: TMR-6-COOH (14 mg, 0.033 mmol, 1.2 equiv.) was dissolved in DMF (~3 mL) and DIPEA (14 mg, 0.11 mmol, 4 equiv.) was added followed by addition of PyAOP (17 mg, 0.033 mmol, 1.2 equiv.). Reaction mixture was stirred for 10 min.

Carbamate **59** (13 mg, 0.027 mmol, 1 equiv.) was dissolved in DCM (~5 mL) and cooled to 0 °C. TFA (2 mL) was added and reaction mixture stirred for 1 h at the constant temperature. After removing volatiles (at 25 °C), active ester of TMR was added at the room temperature to the crude ammonium residue in DMF (~ 3 mL) and reaction was stirred for 4 h. Unreacted components were quenched by addition of 0.5 mL of 10% AcOH in water, volatiles removed under reduced pressure and purified by preparative HPLC to provide 7 mg (42% yield) of fluorophore conjugate **1** as a red solid.

**HRMS** ($EI^+$) m/z: $[2M]^+$, Calculated for $C_{37}H_{35}ClN_6O_5^{2+}$ 339.1173, found 339.1175.

**Conjugate 2:**

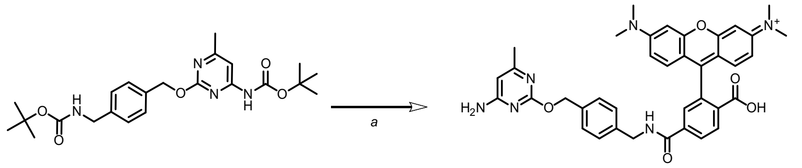


Conjugate (**2**) was synthesized as described above for **1**. Solid red product was isolated by preparative HPLC (8 mg, 54% yield).

**HRMS** ($EI^+$) m/z: $[M + 2H]^{2+}$, Calculated for $C_{38}H_{38}N_6O_5^{2+}$ 329.1447, found 329.1445.

**Conjugate 3:**

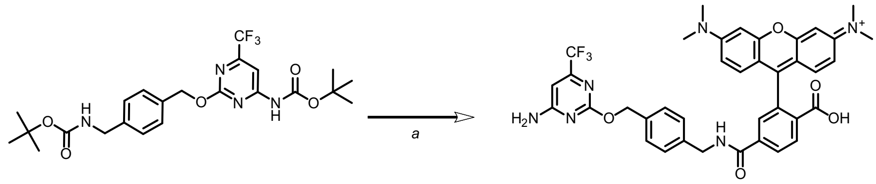


Conjugate (**3**) was synthesized like described above for **1**. Solid red product was isolated by preparative HPLC (7 mg, 41% yield).

**HRMS** ($EI^+$) m/z: $[M + H]^+$, Calculated for $C_{38}H_{34}F_3N_6O_5^+$ 711.2537, found 711.2541.

**Conjugate 4:**

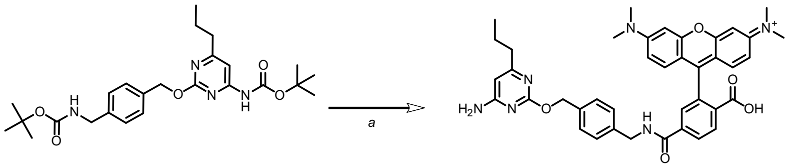

Conjugate (**4**) was synthesized as described above for **1**. Solid red product was isolated by preparative HPLC (8 mg, 46% yield).

**HRMS** ($EI^+$) m/z: $[M + 2H]^{2+}$, Calculated for $C_{40}H_{42}N_6O_5^{2+}$ 343.1603, found 343.1599.

**Conjugate 5:**

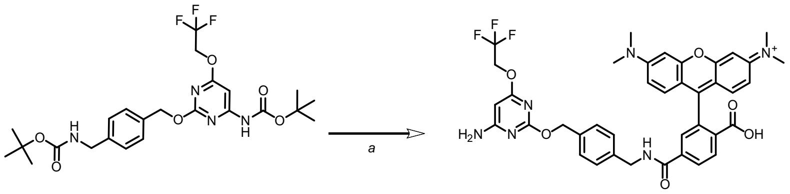

Conjugate (**5**) was synthesized as described above for **1**. Solid red product was isolated by preparative HPLC (5 mg, 44% yield).

**HRMS** ($EI^+$) m/z: $[M + H]^+$, Calculated for $C_{39}H_{36}F_3N_6O_6^+$ 741.2643, found 741.2640.

**Conjugate 6:**

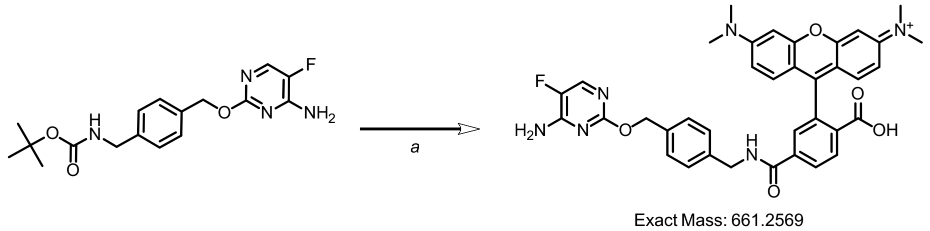


Conjugate (**5**) was synthesized as described above for **1**. Solid red product was isolated by preparative HPLC (9 mg, 24% yield).

**HRMS** ($EI^+$) m/z: $[M + H]^+$, Calculated for $C_{37}H_{34}FN_6O_5^+$ 661.2569, found 661.2573.

**Conjugate 7:**

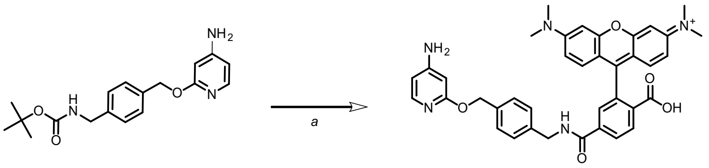

Conjugate (**7**) was synthesized as described above for **1**. Solid red product was isolated by preparative HPLC (10 mg, 51% yield).

**HRMS** ($EI^+$) m/z: $[M + H]^{2+}$, Calculated for $C_{38}H_{37}N_5O_5^{2+}$ 321.6392, found 321.6388.

**Conjugate 8:**

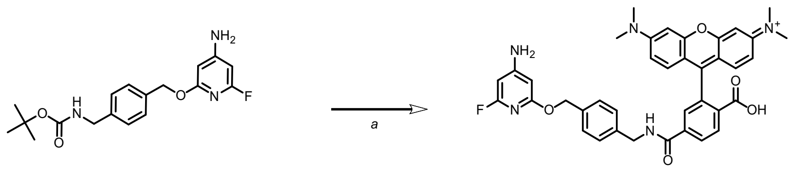

Conjugate (**8**) was synthesized as described above for **1**. Solid red product was isolated by preparative HPLC (8 mg, 42% yield).

**HRMS** ($EI^+$) m/z: $[M + 2H]^{2+}$, Calculated for $C_{38}H_{36}FN_5O_5^{2+}$ 330.6345, found 330.6344.

**Conjugate 9:**

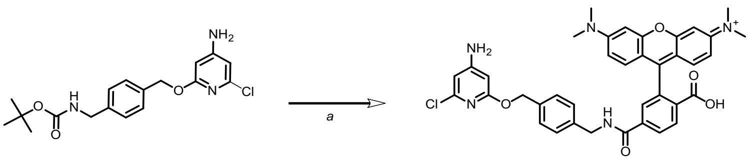

Conjugate (**9**) was synthesized as described above for **1**. Solid red product was isolated by preparative HPLC (5 mg, 26% yield).

**HRMS** ($EI^+$) m/z: $[M + 2H]^{2+}$, Calculated for $C_{38}H_{36}FN_5O_5^{2+}$ 330.6345, found 330.6344.

**Conjugate 10:**

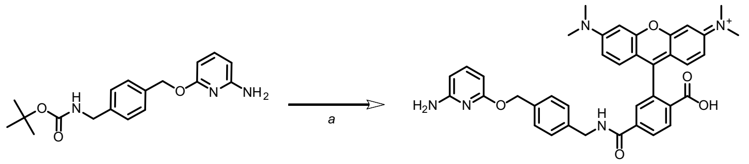

Conjugate (**10**) was synthesized as described above for **1**. Solid red product was isolated by preparative HPLC (7 mg, 36% yield).

**HRMS** ($EI^+$) m/z: $[M + H]^+$, Calculated for $C_{38}H_{36}N_5O_5^+$642.2711, found 642.2712.

**Conjugate 11:**

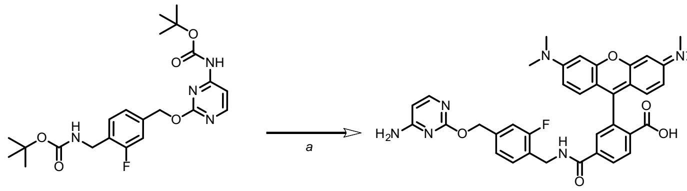

Conjugate (**11**) was synthesized as described above for **1**. Solid red product was isolated by preparative HPLC (4 mg, 67% yield).

**HRMS** ($EI^+$) m/z: $[M + H]^+$, Calculated for $C_{37}H_{34}FN_6O_5^+$ 661.2569, found 661.2575.

**Conjugate 12:**

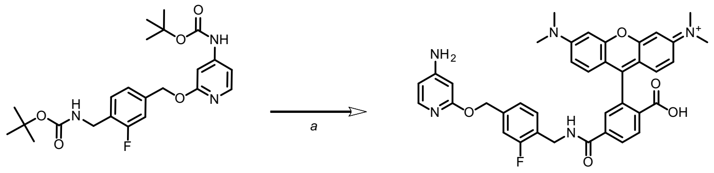

Conjugate (12) was synthesized as described above for **1**. Solid red product was isolated by preparative HPLC (7 mg, 92% yield).

**HRMS** ($EI^+$) m/z: $[M + H]^+$, Calculated for $C_{38}H_{35}FN_5O_5^+$ 660.2617, found 660.2623.

**Conjugate 13:**

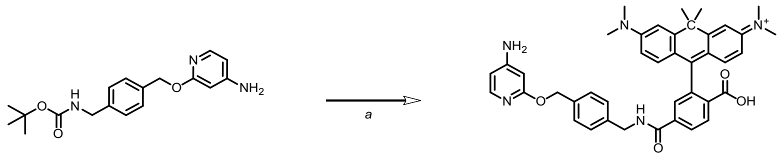

Conjugate **13** was synthesized as described above for **1** (CPY-6-COOH, 7 mg, 0.015 mmol was used). Dark blue to violet product was isolated by preparative HPLC (7 mg, 69% yield).

**HRMS** ($EI^+$) m/z: $[M]^+$, Calculated for $C_{41}H_{42}N_5O_4^+$ 668.3231, found 668.3226.

**Conjugate 14:**

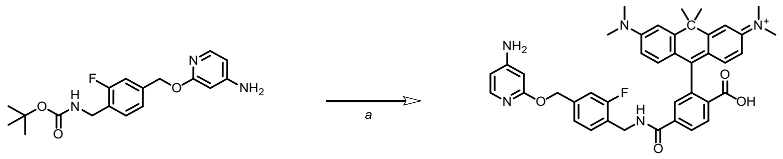

Conjugate **14** was synthesized as described above for **1** (CPY-6-COOH, 5 mg, 0.011 mmol was used). Dark blue to violet product was isolated by preparative HPLC (6 mg, 78% yield).

**HRMS** ($EI^+$) m/z: $[M]^+$, Calculated for $C_{40}H_{42}FN_5O_4^+$ 686.3137, found 686.3139.

**Conjugate 15:**

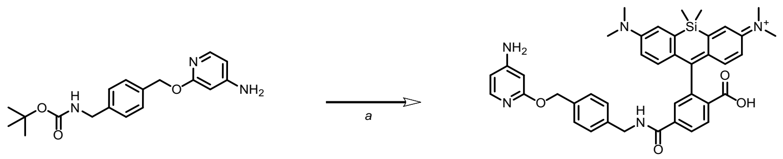


Conjugate **15** was synthesized as described above for **1** (SiR-6-COOH, 5.2 mg, 0.011 mmol was used). Dark blue product was isolated by preparative HPLC (13 mg, 86% yield).

**HRMS** ($EI^+$) m/z: $[M]^+$, Calculated for $C_{40}H_{42}N_5O_4Si^+$ 684.3001, found 684.3004.

**Conjugate 16:**

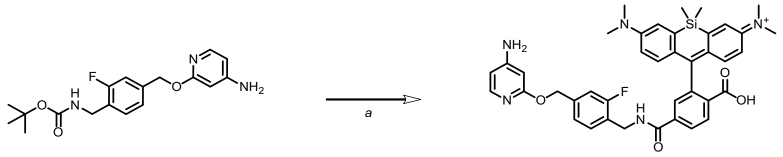


Conjugate **16** was synthesized as described above for **1** (SiR-6-COOH, 11 mg, 0.024 mmol was used). Dark blue product was isolated by preparative HPLC (7 mg, 89% yield).

**HRMS** ($EI^+$) m/z: $[M]^+$, Calculated for $C_{40}H_{41}N_5FO_4Si^+$ 702.2906, found 702.2909.

#### 6 NMR Spectra

##### 1H spectrum of compound 20:

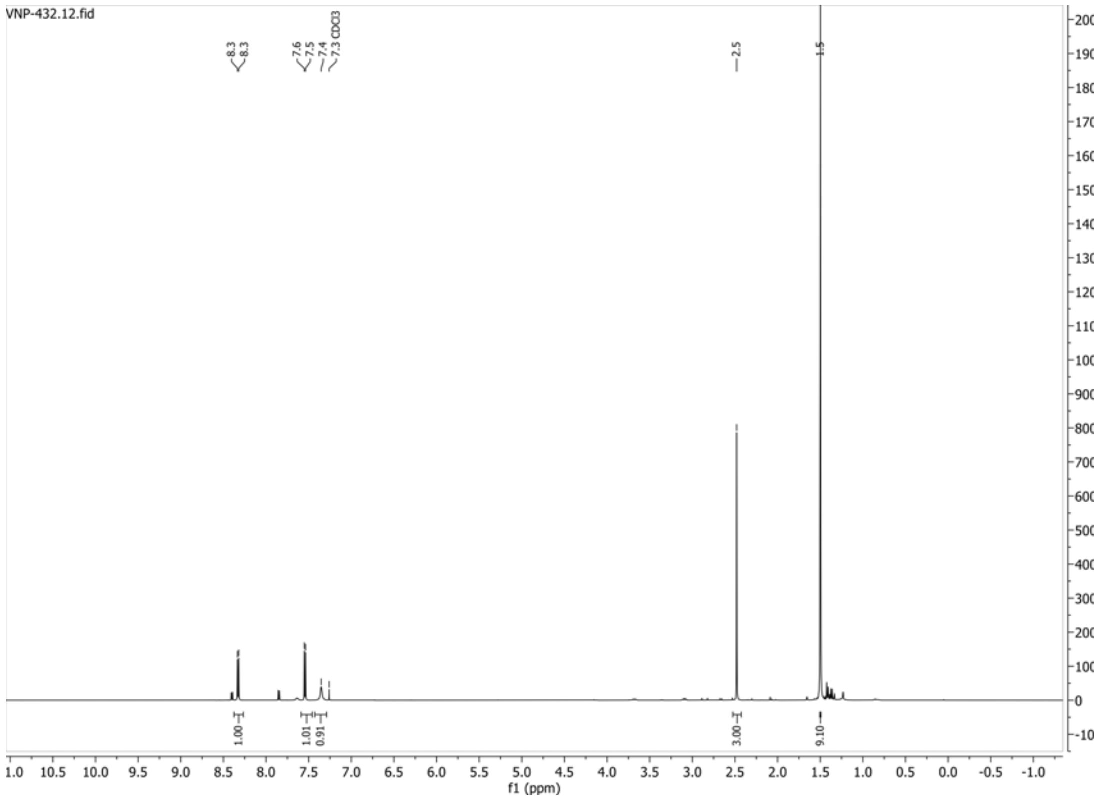


##### 13C{1H} spectrum of compound 20:

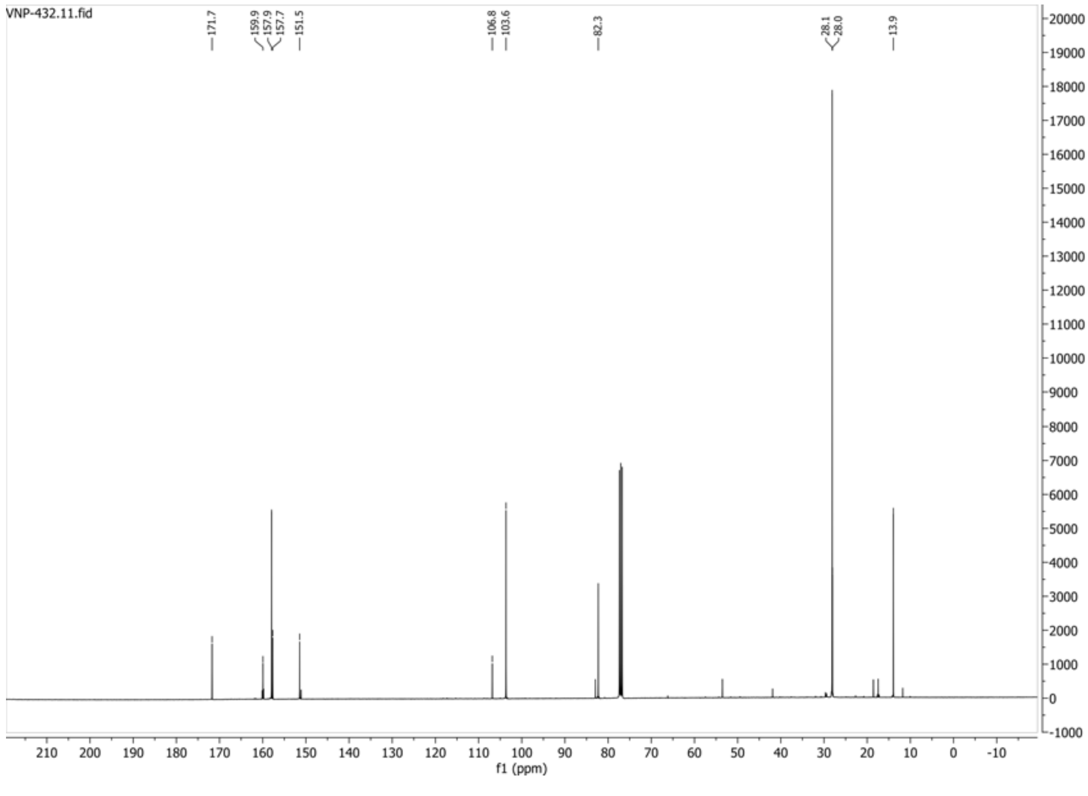

**HSQC-DEPT135 spectrum of compound 20:**

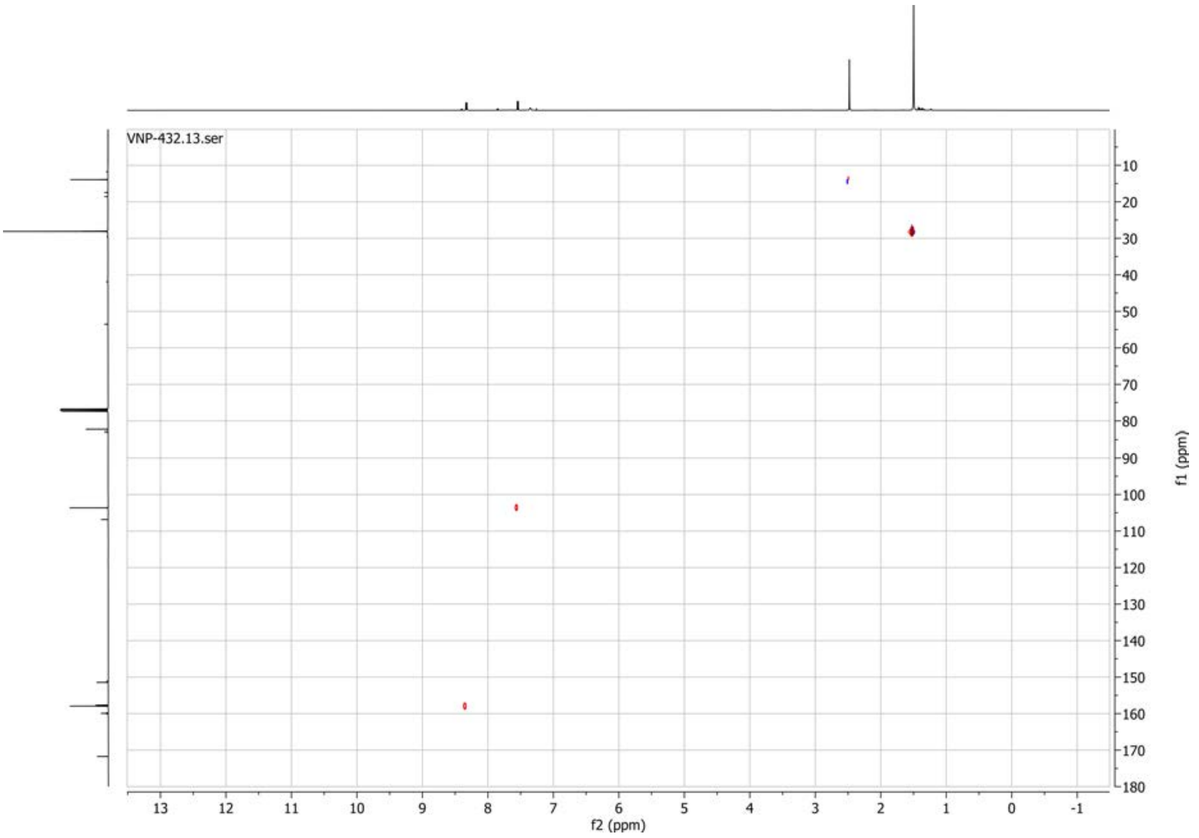

**1H spectrum of compound 19:**

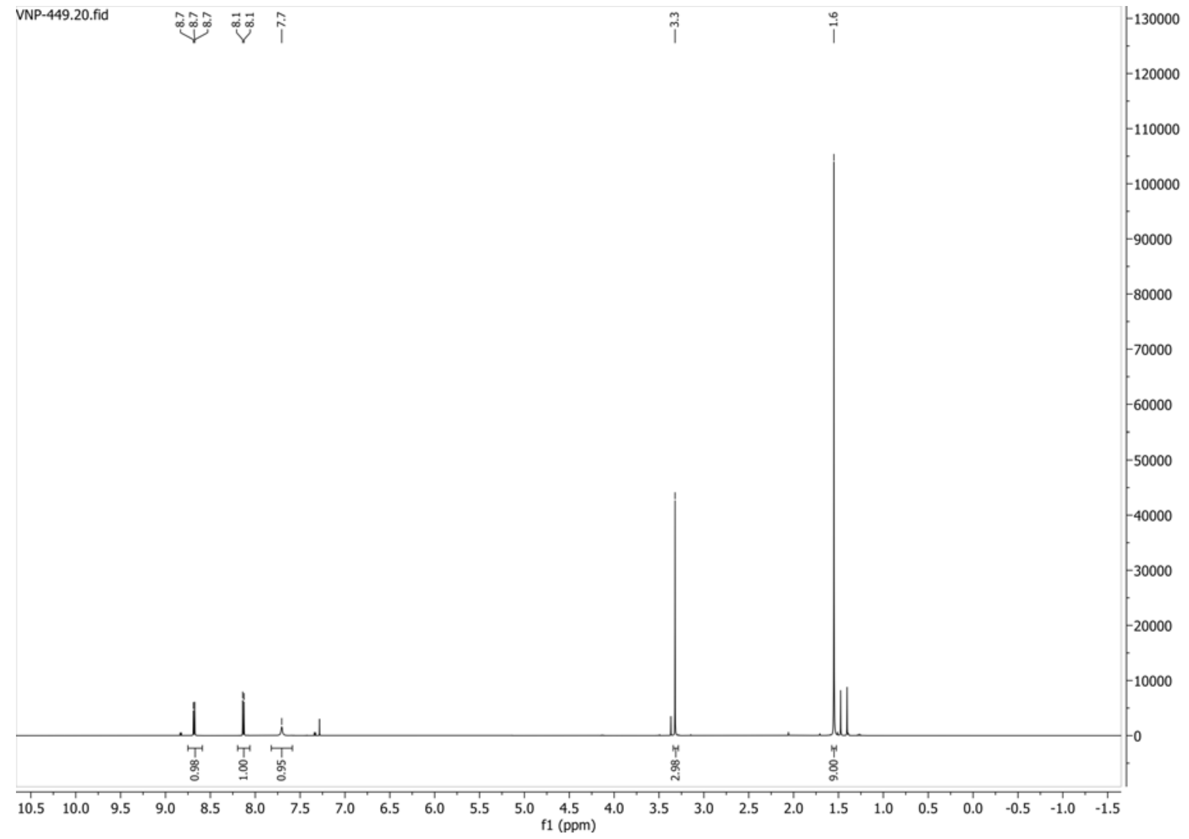


**13C{1H} spectrum of compound 19:**

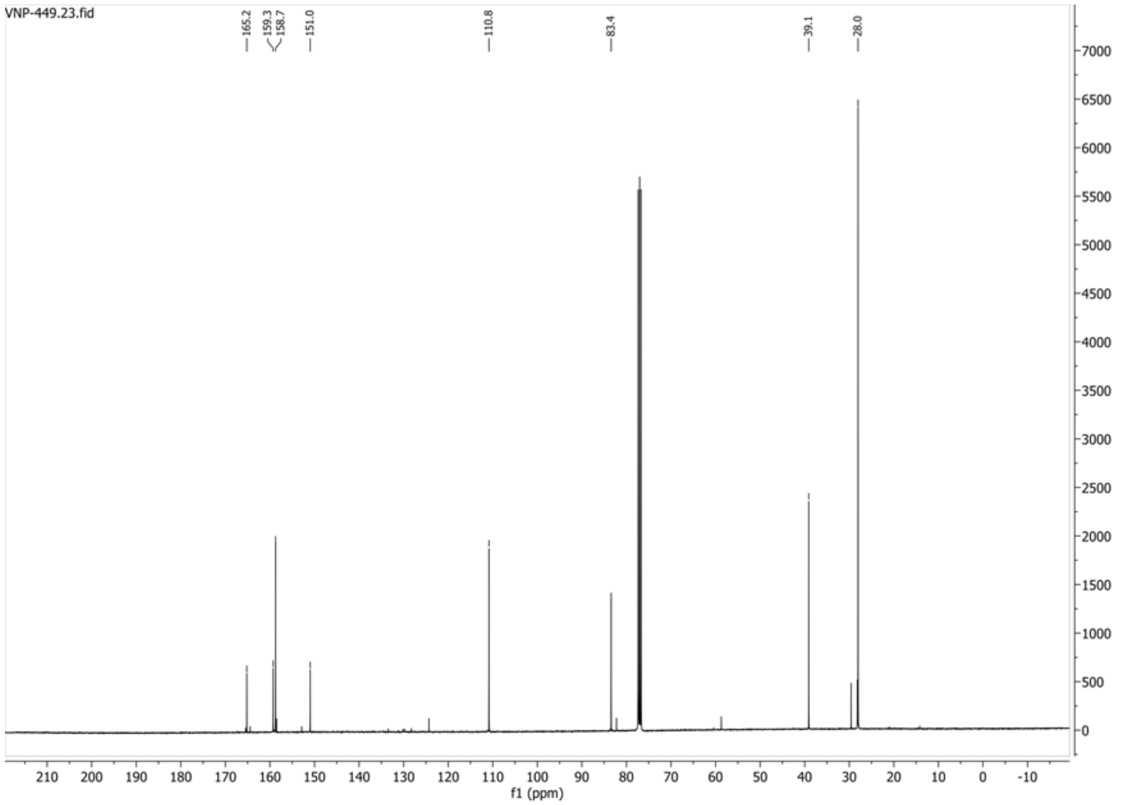

**HSQC-DEPT135 spectrum of compound 19:**

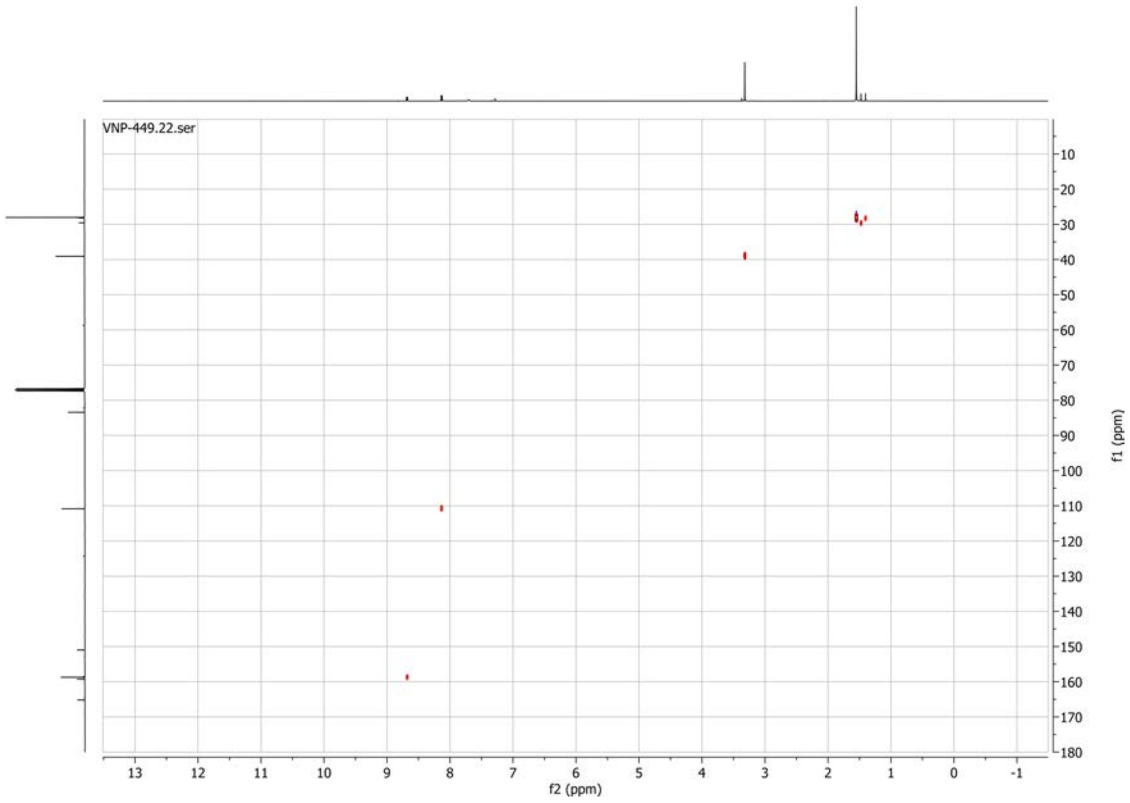

**1H spectrum of compound 22:**

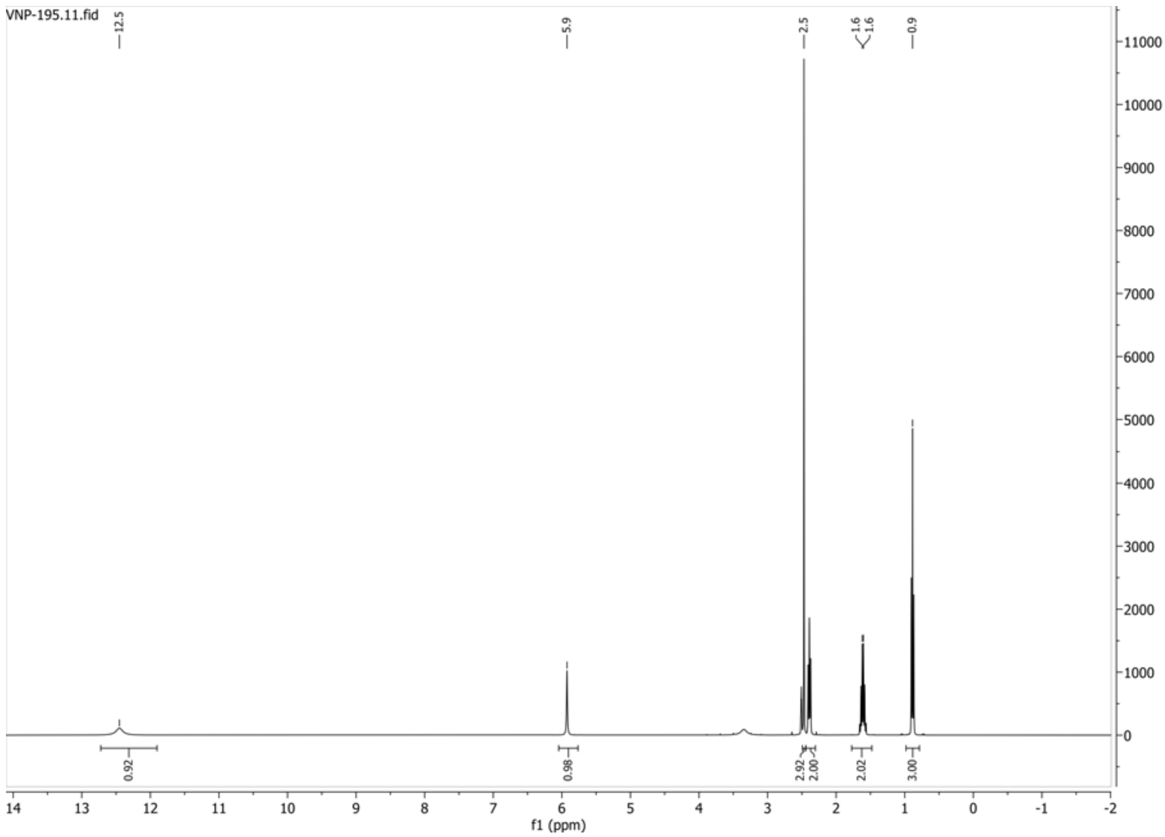


**13C{1H} spectrum of compound 22:**

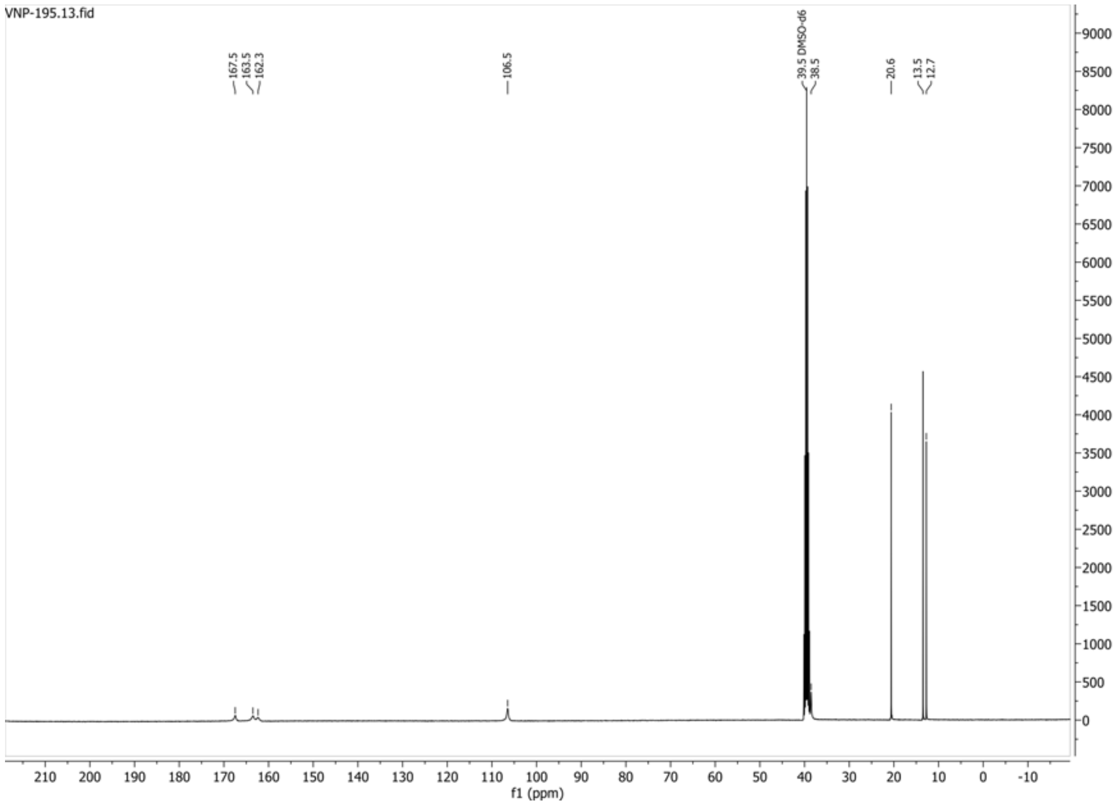

**HSQC-DEPT135 spectrum of compound 22:**

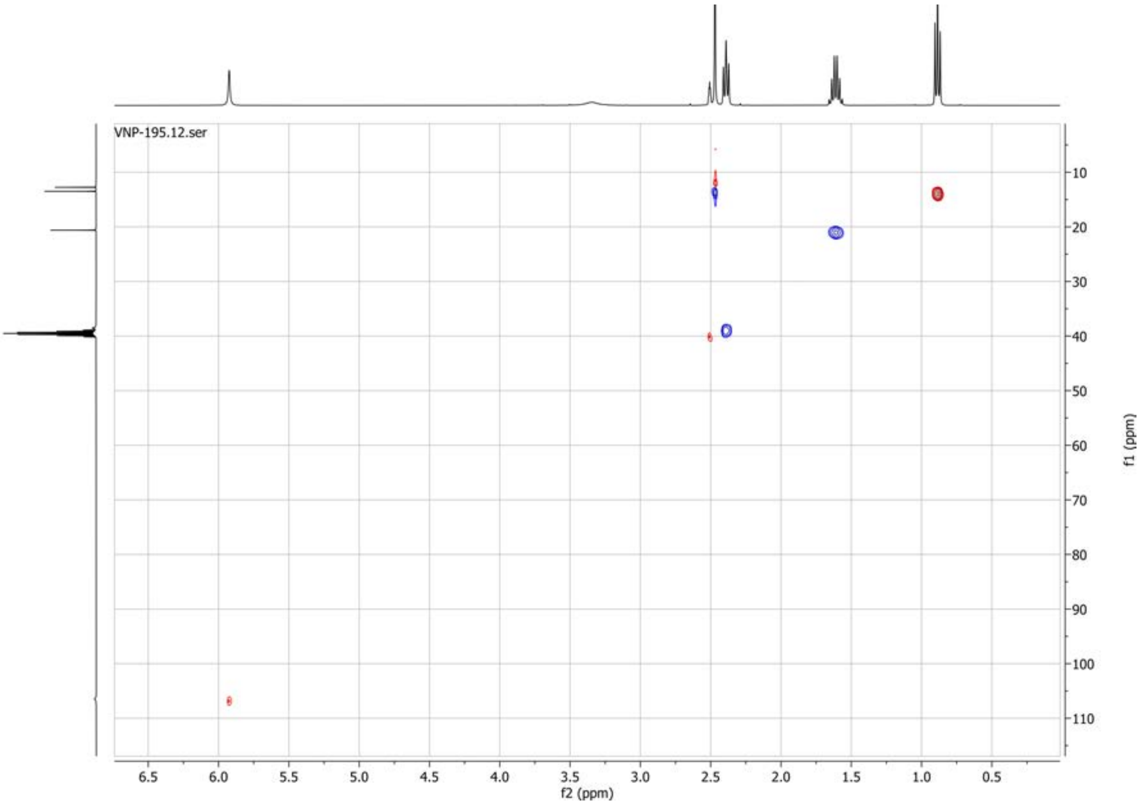

**1H spectrum of compound 23:**

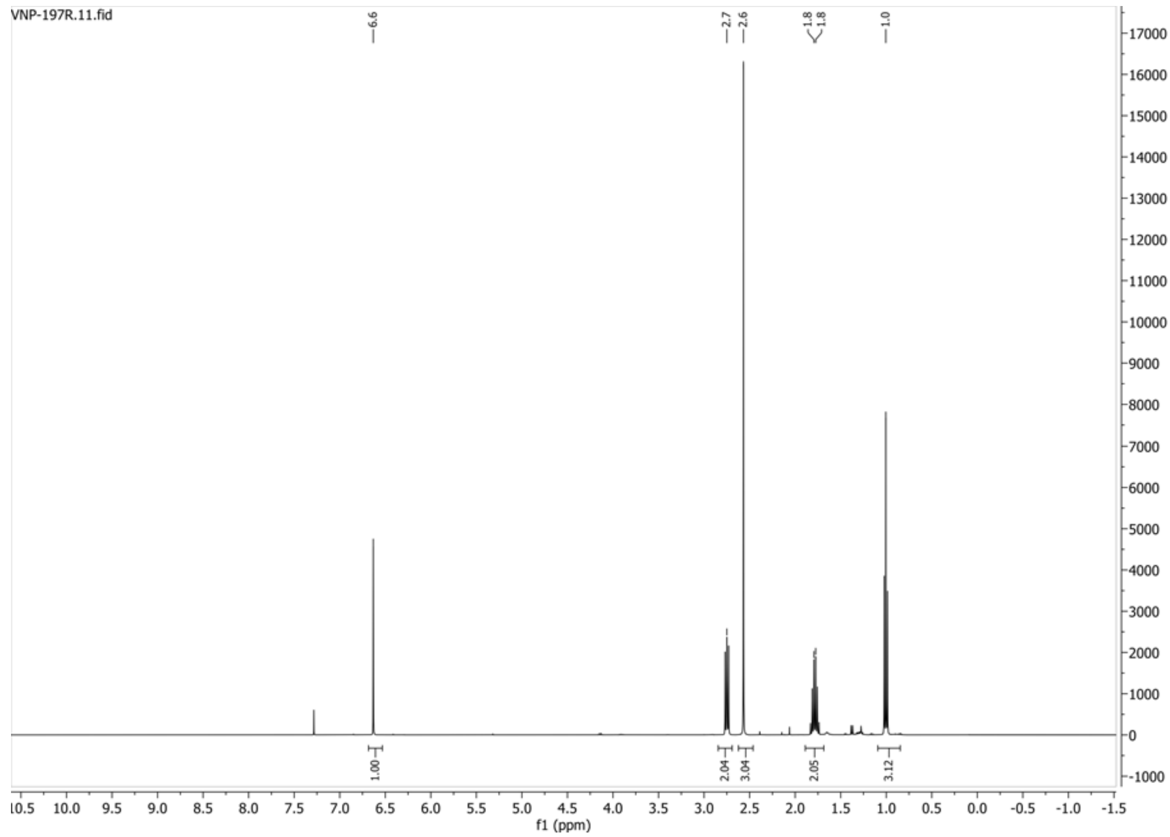


**13C{1H} spectrum of compound 23:**

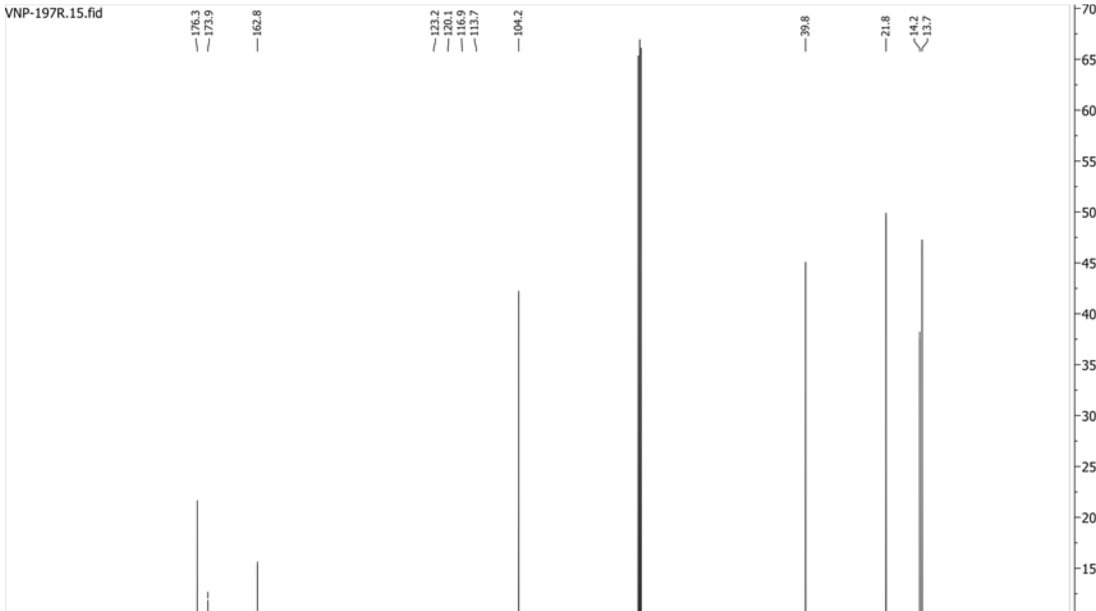

**19F{1H} spectrum of compound 23:**

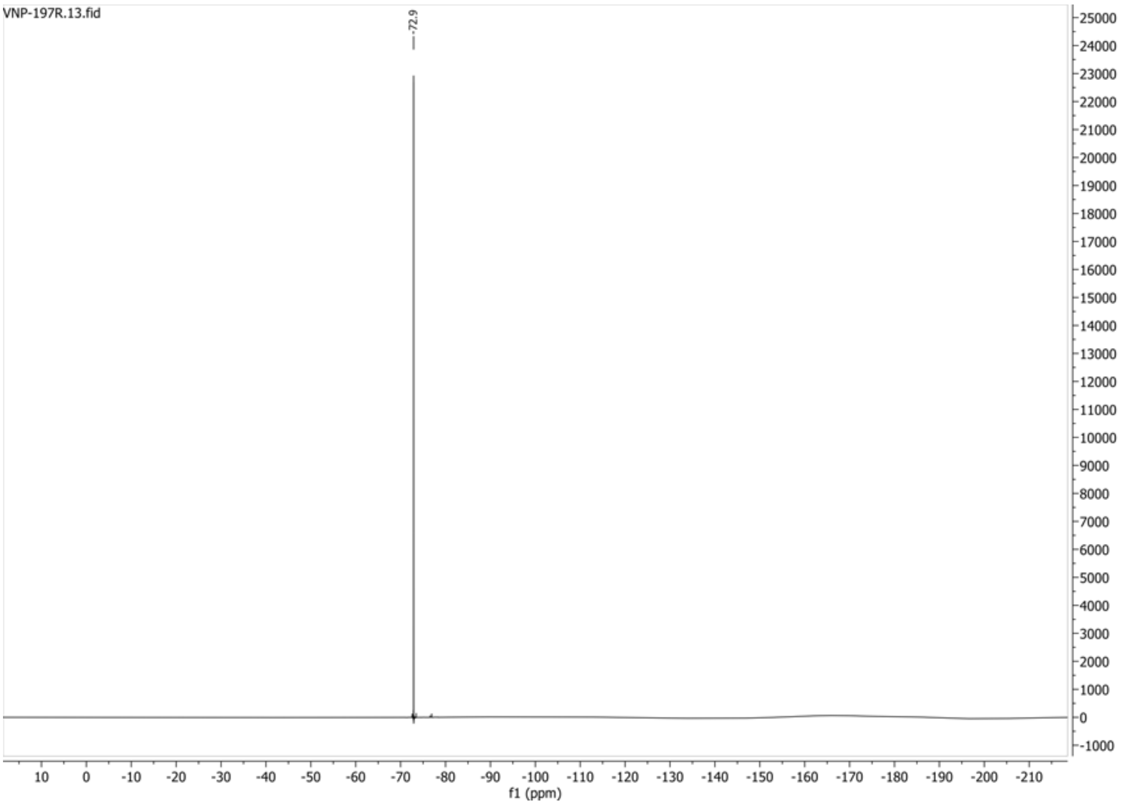


**HSQC-DEPT135 spectrum of compound 23:**

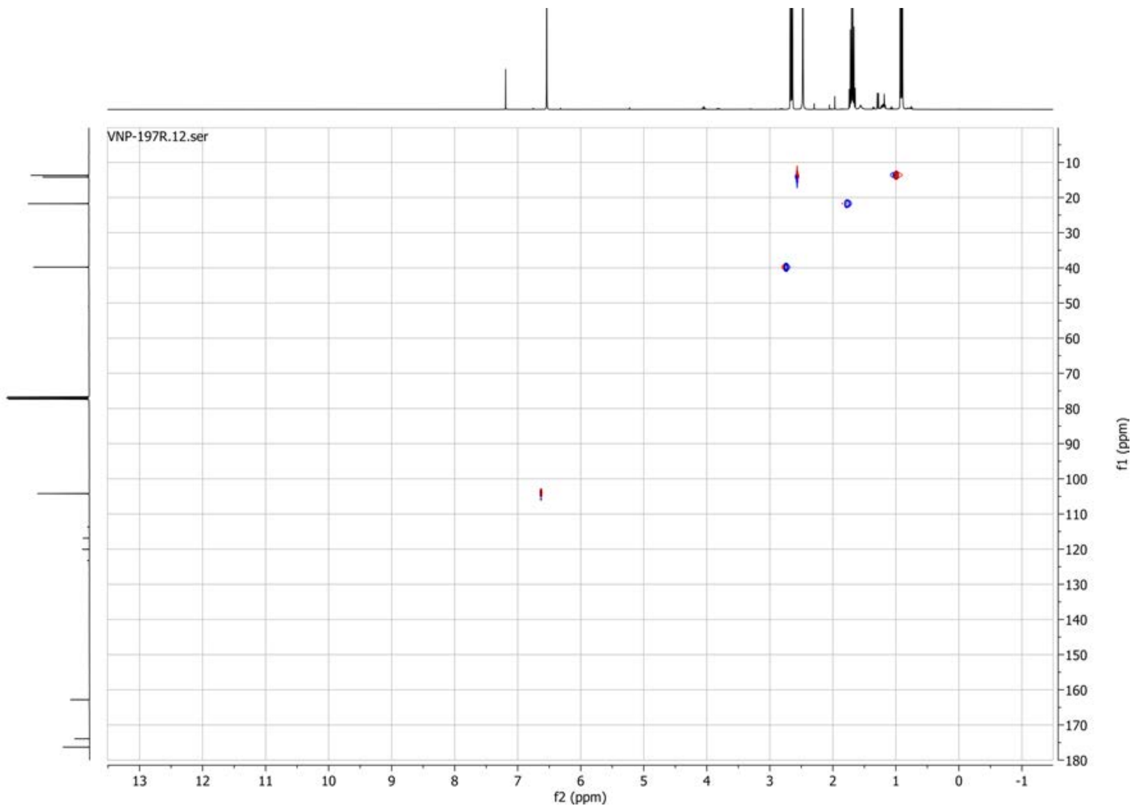

**1H spectrum of compound 24:**

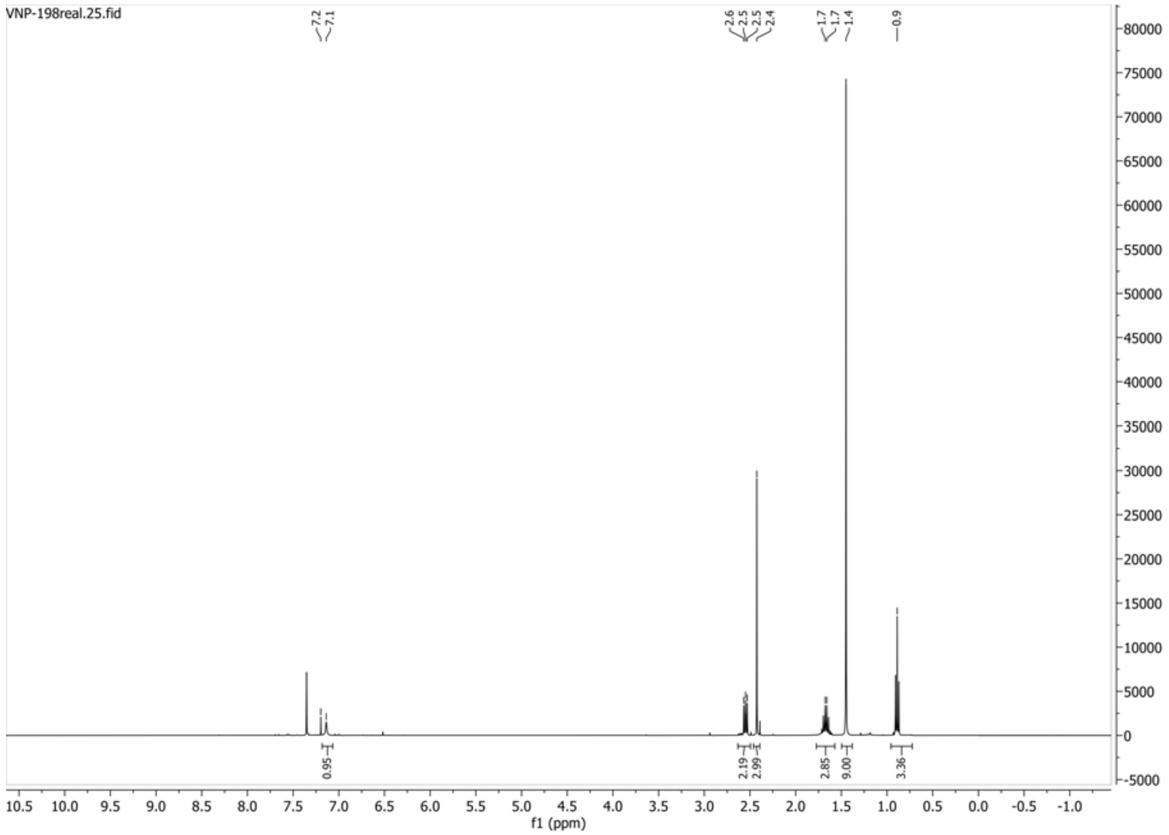


**13C{1H} spectrum of compound 24:**

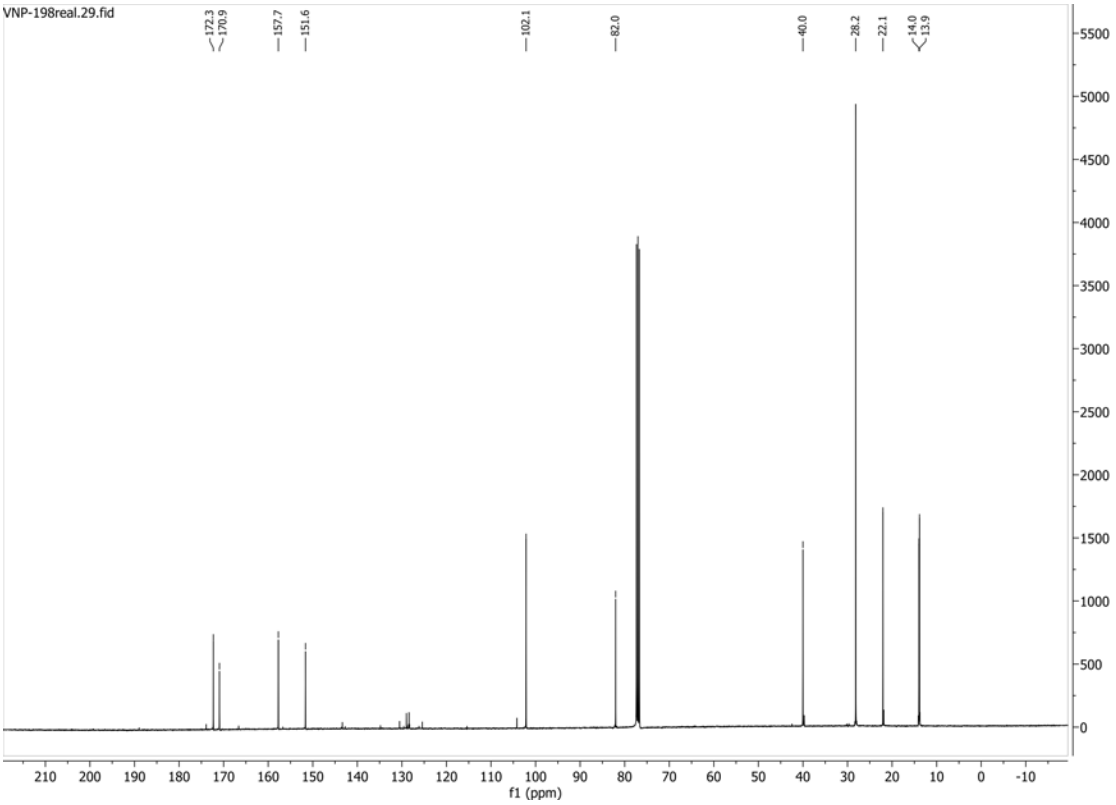

**HSQC-DEPT135 spectrum of compound 24:**

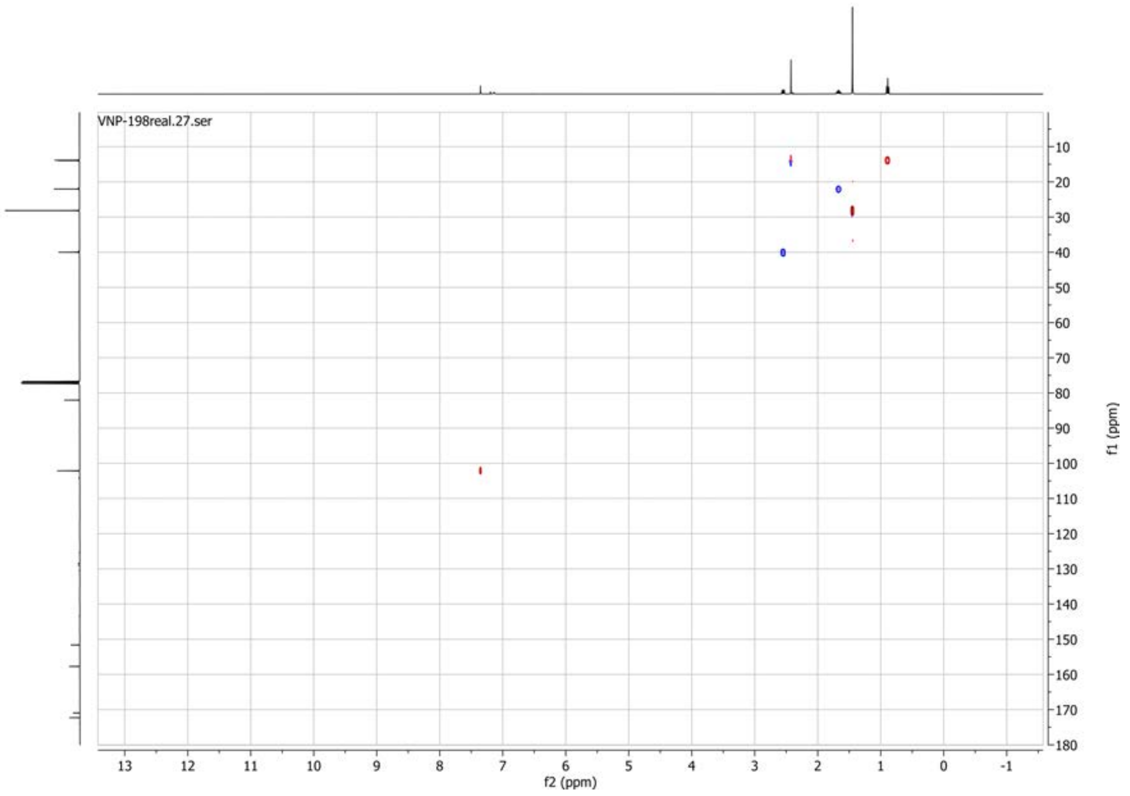

**1H spectrum of compound 25:**

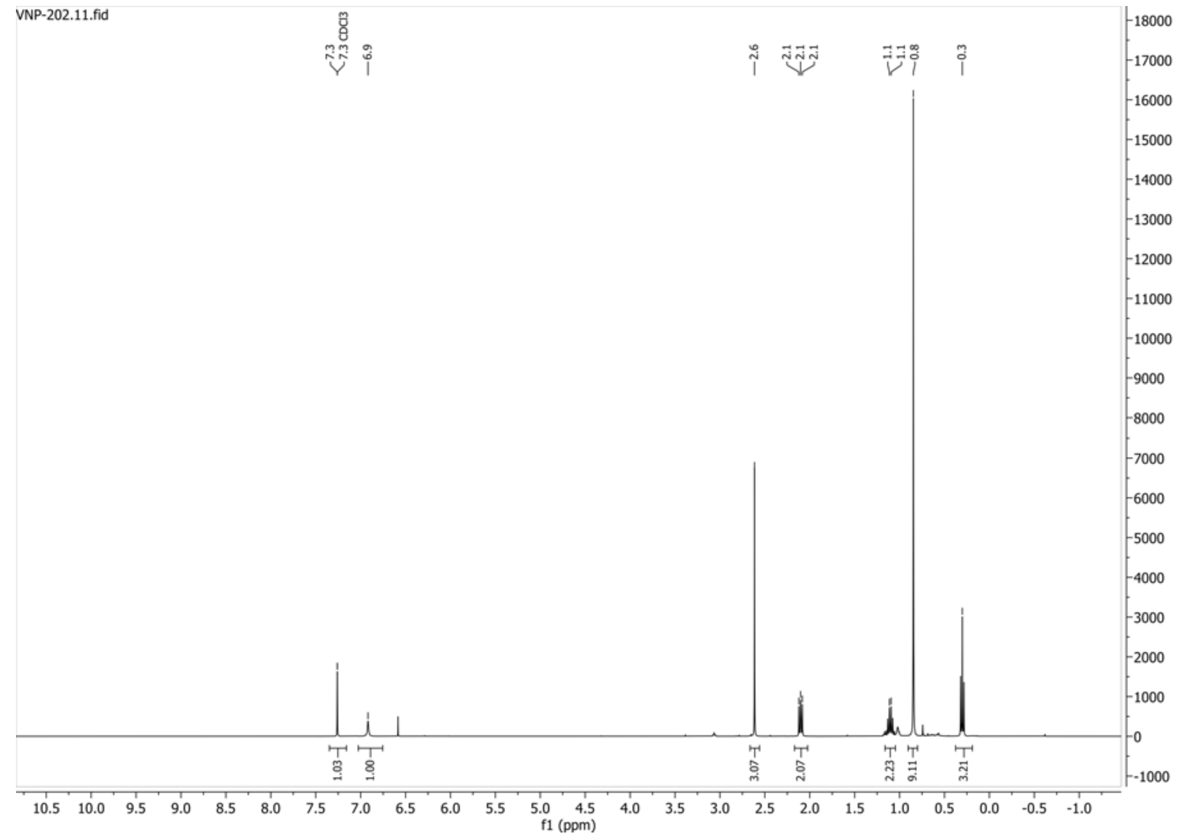


**13C{1H} spectrum of compound 25:**

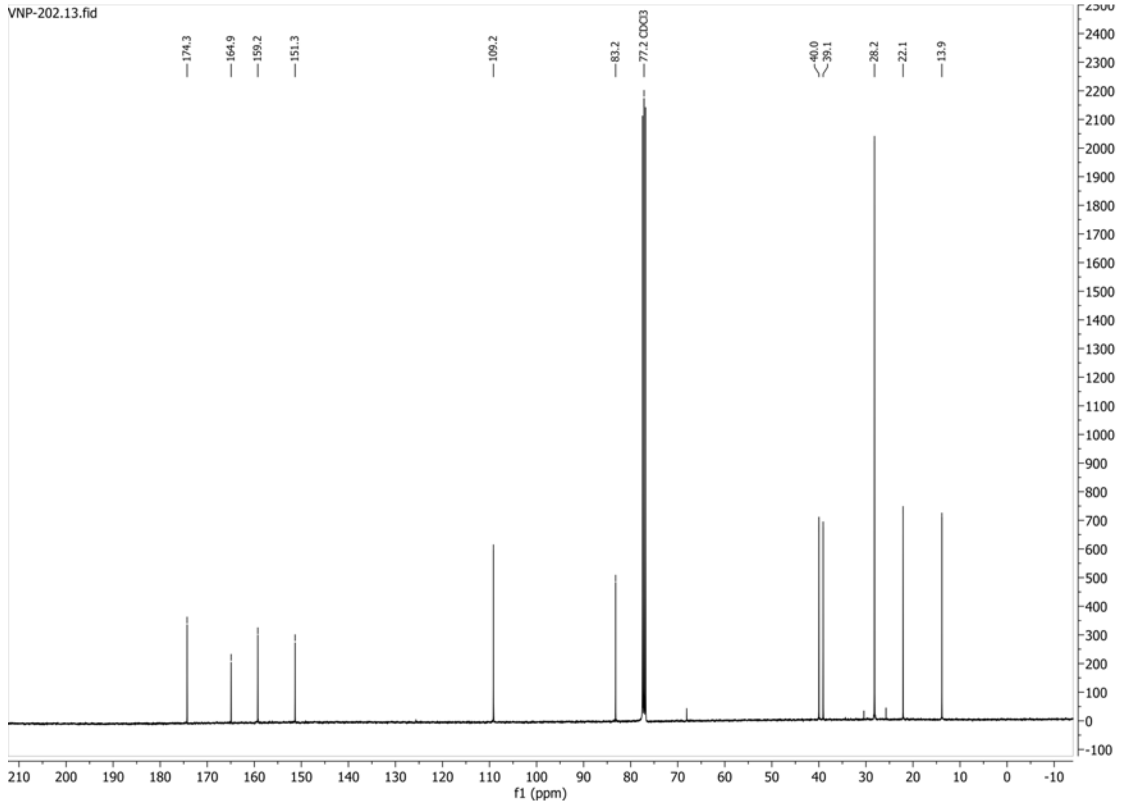

**HSQC-DEPT135 spectrum of compound 25:**

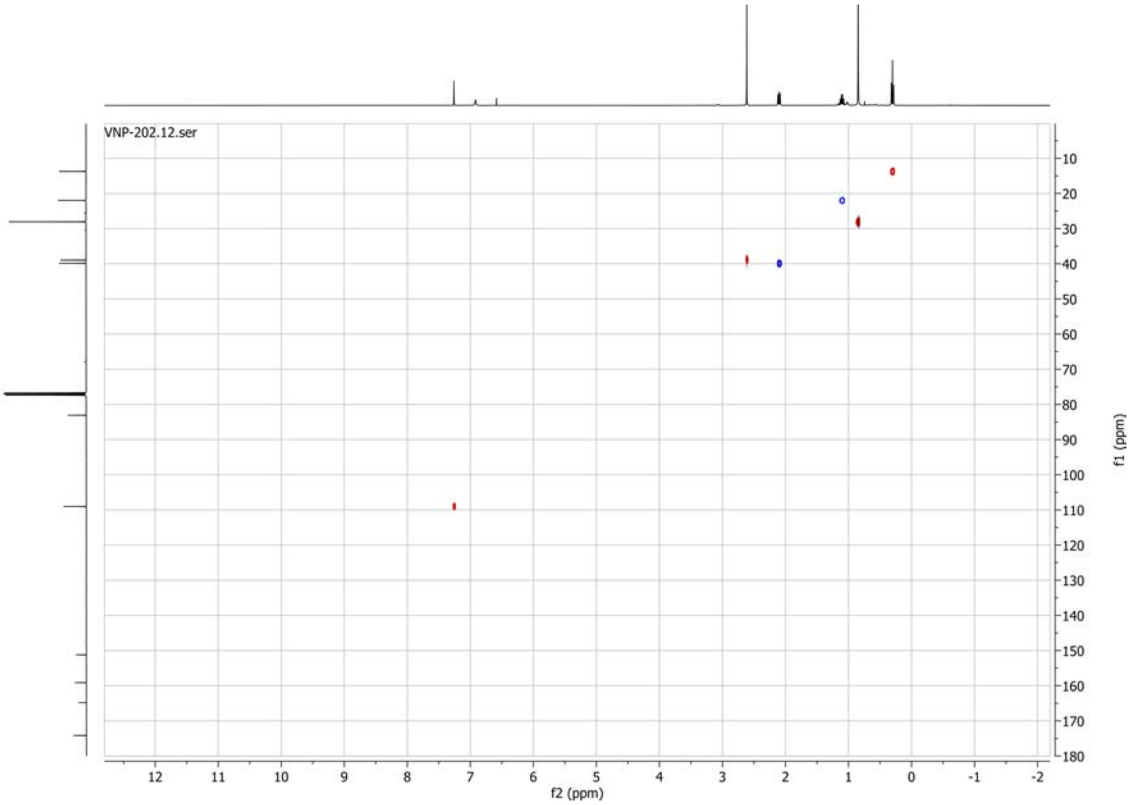

**1H spectrum of compound 27:**

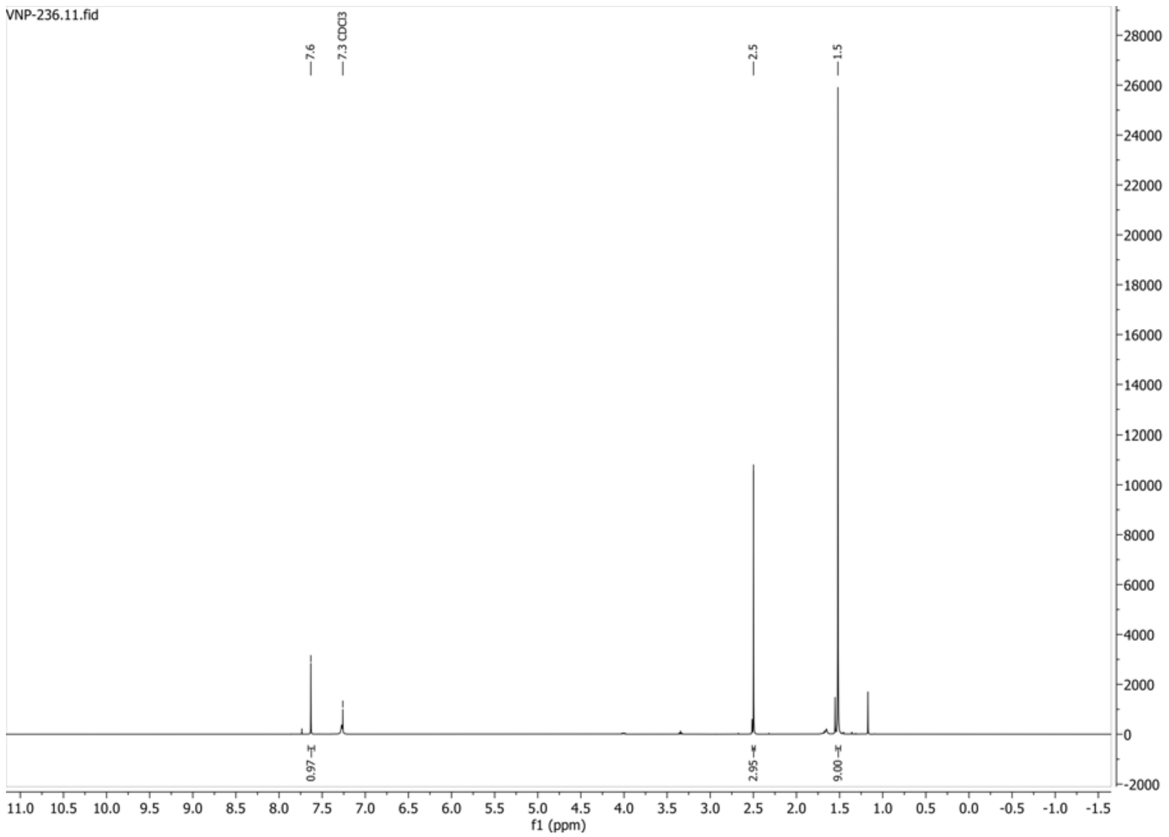


**13C{1H} spectrum of compound 27:**

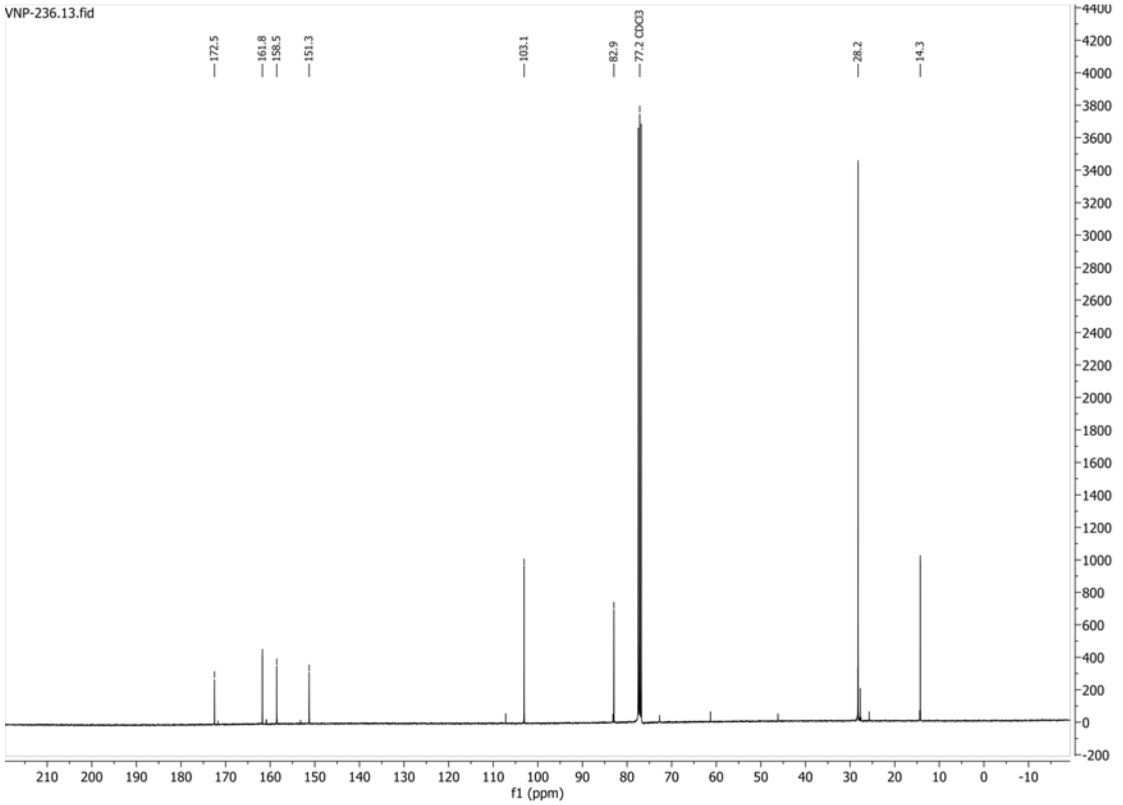

**HSQC-DEPT135 spectrum of compound 27:**

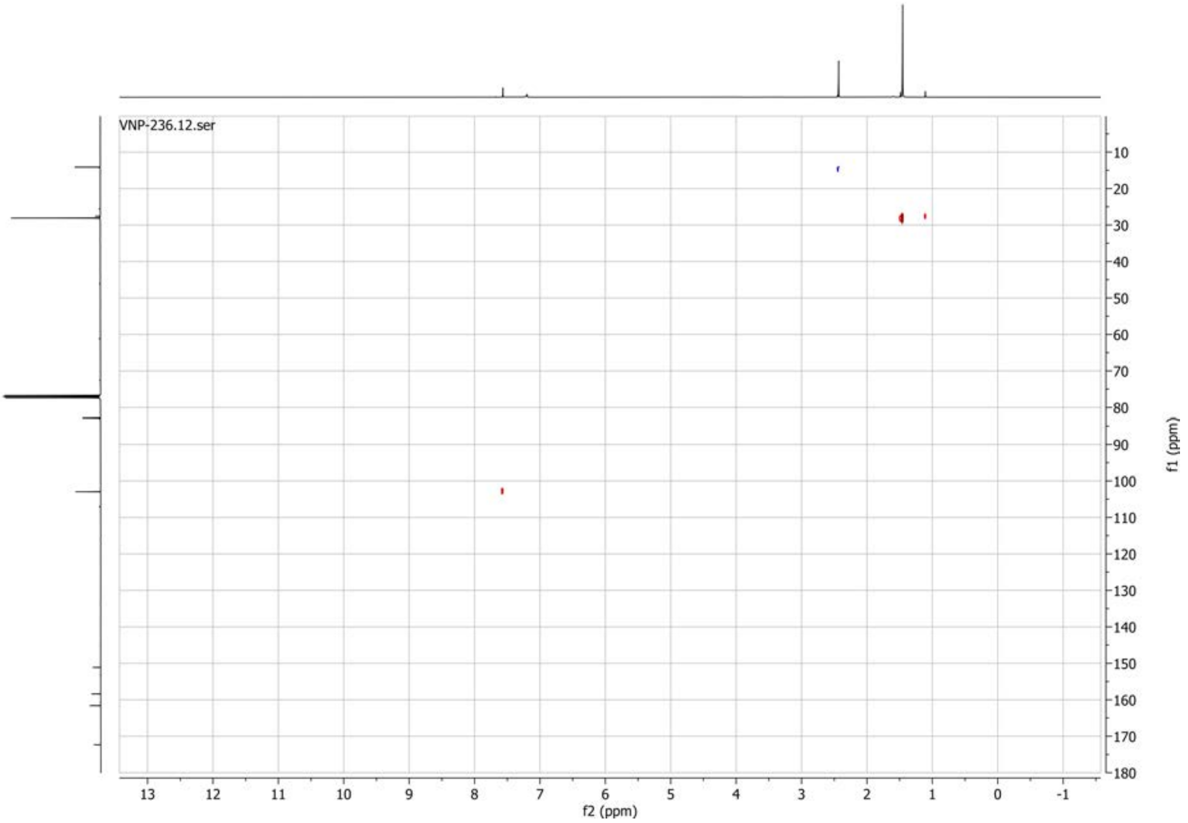

**1H spectrum of compound 28:**

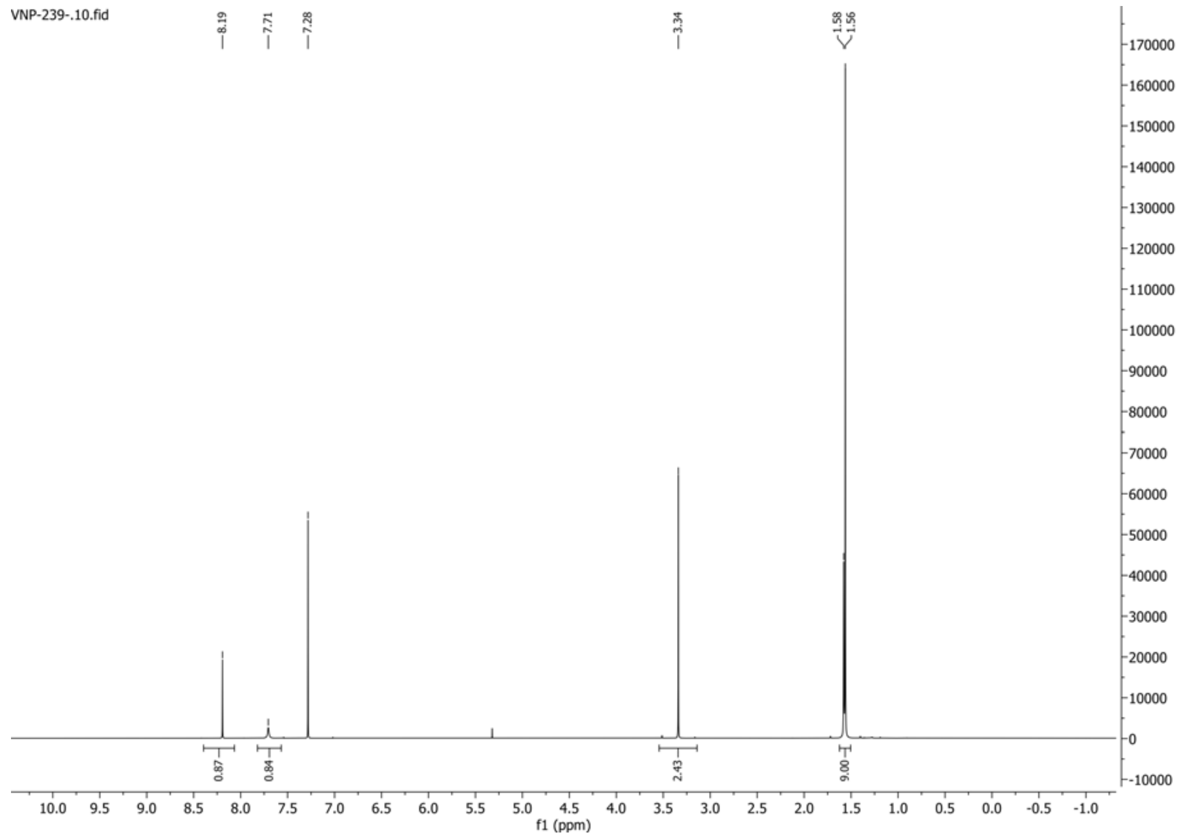


**13C{1H} spectrum of compound 28:**

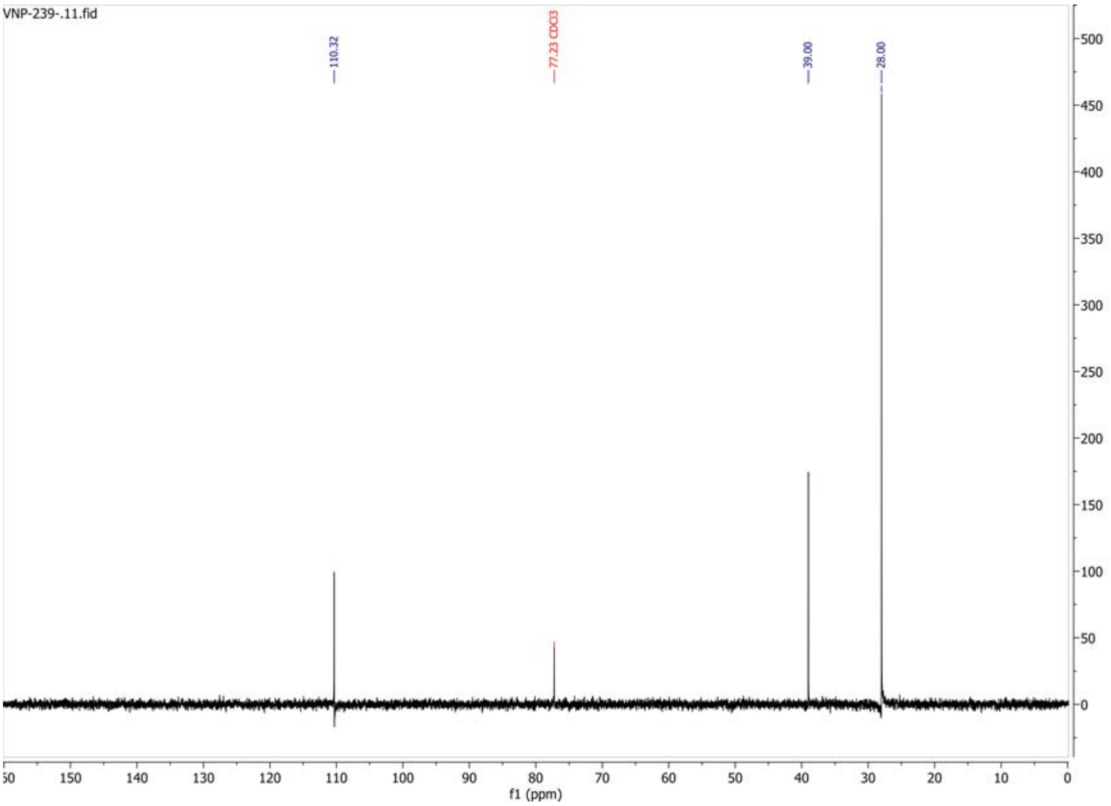

#### 1H spectrum of compound 30:

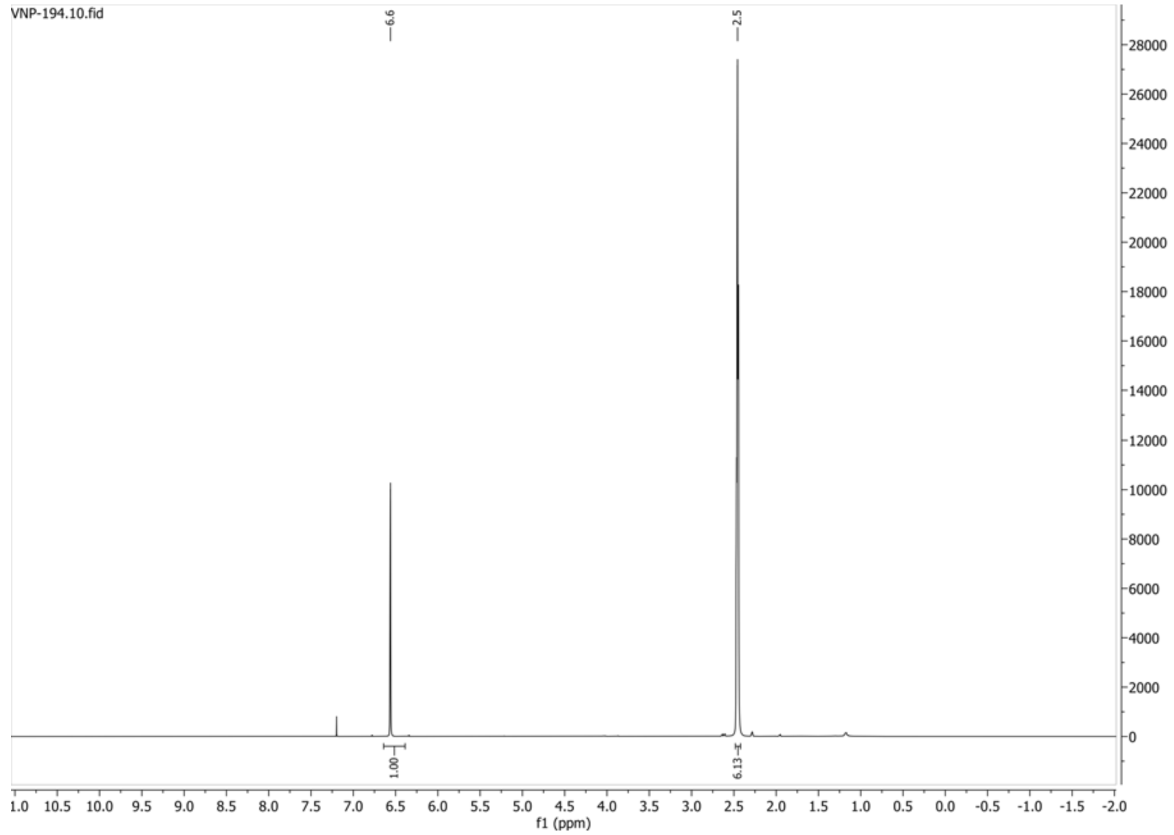


#### 13C{1H} spectrum of compound 30:

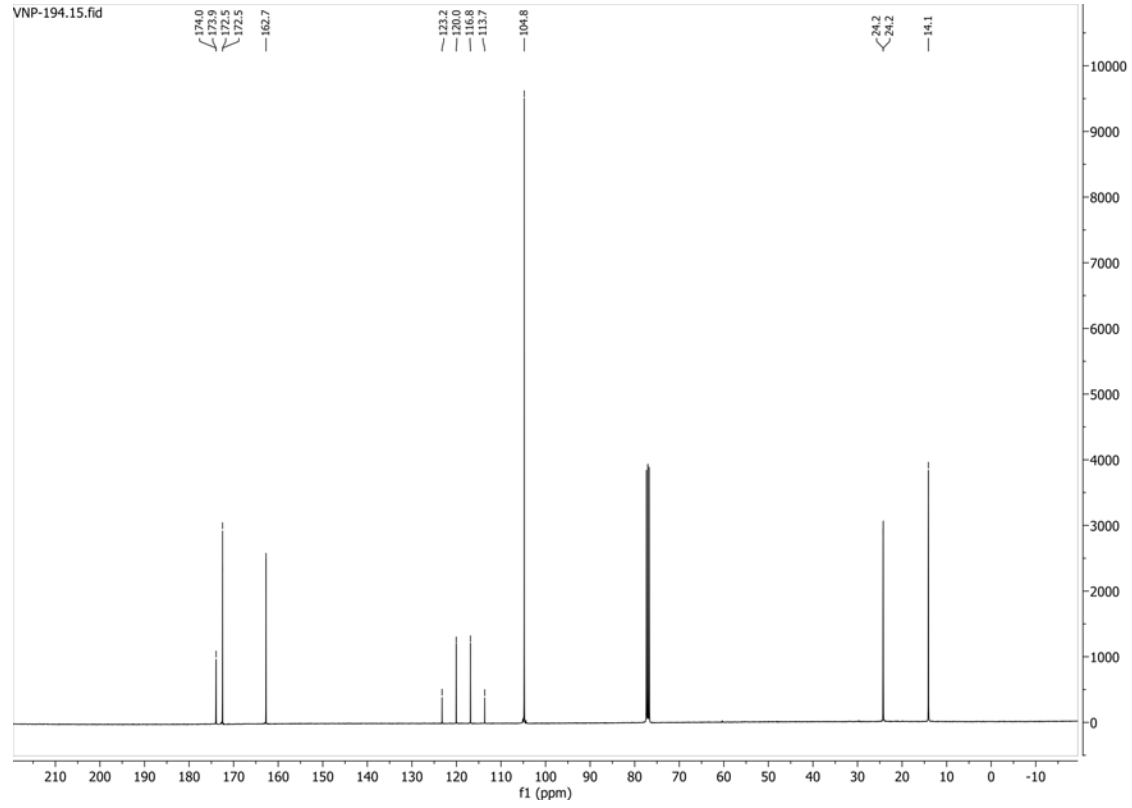

$^{19}F\{^1H\}$ spectrum of compound 30:

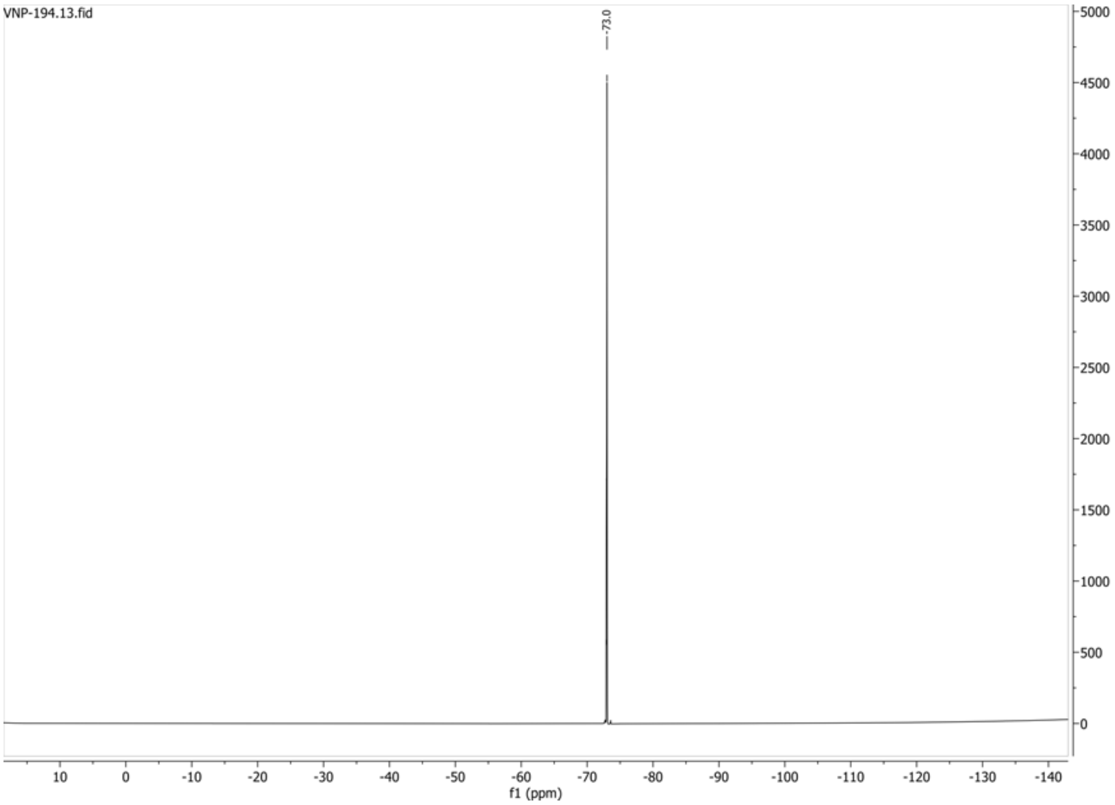


HSQC-DEPT135 spectrum of compound 30:

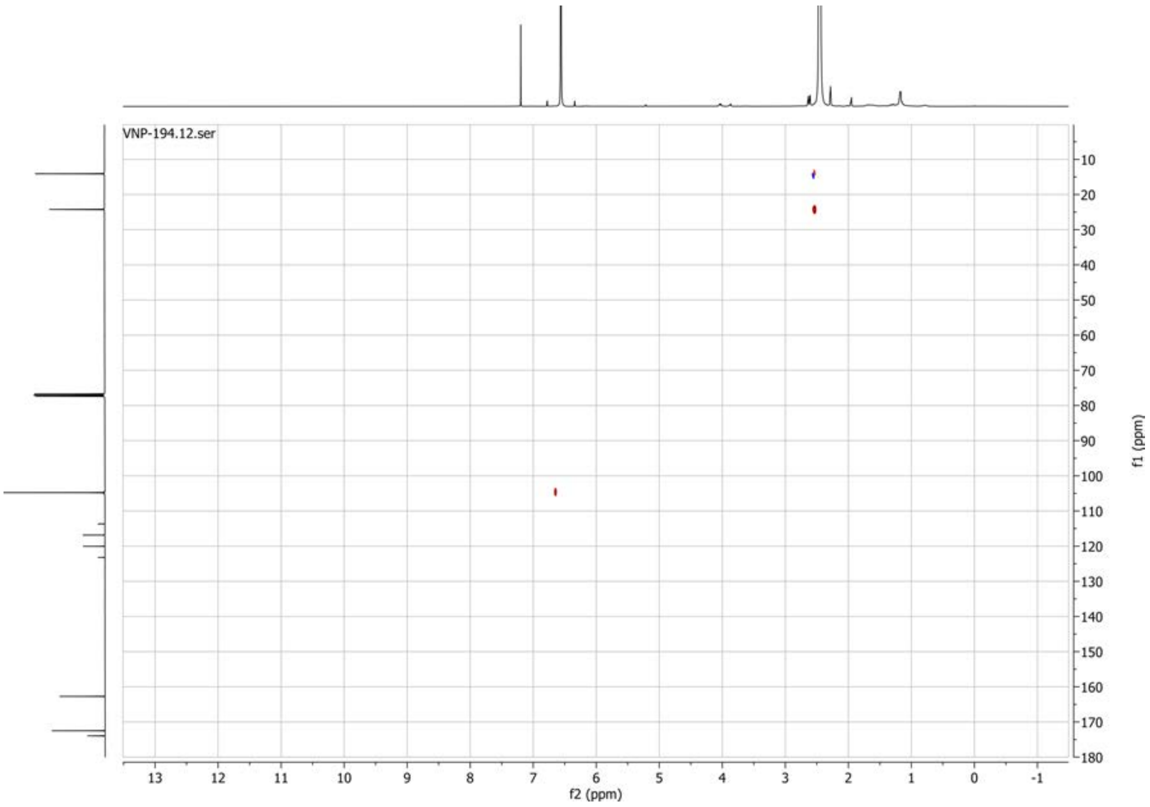

**1H spectrum of compound 31:**

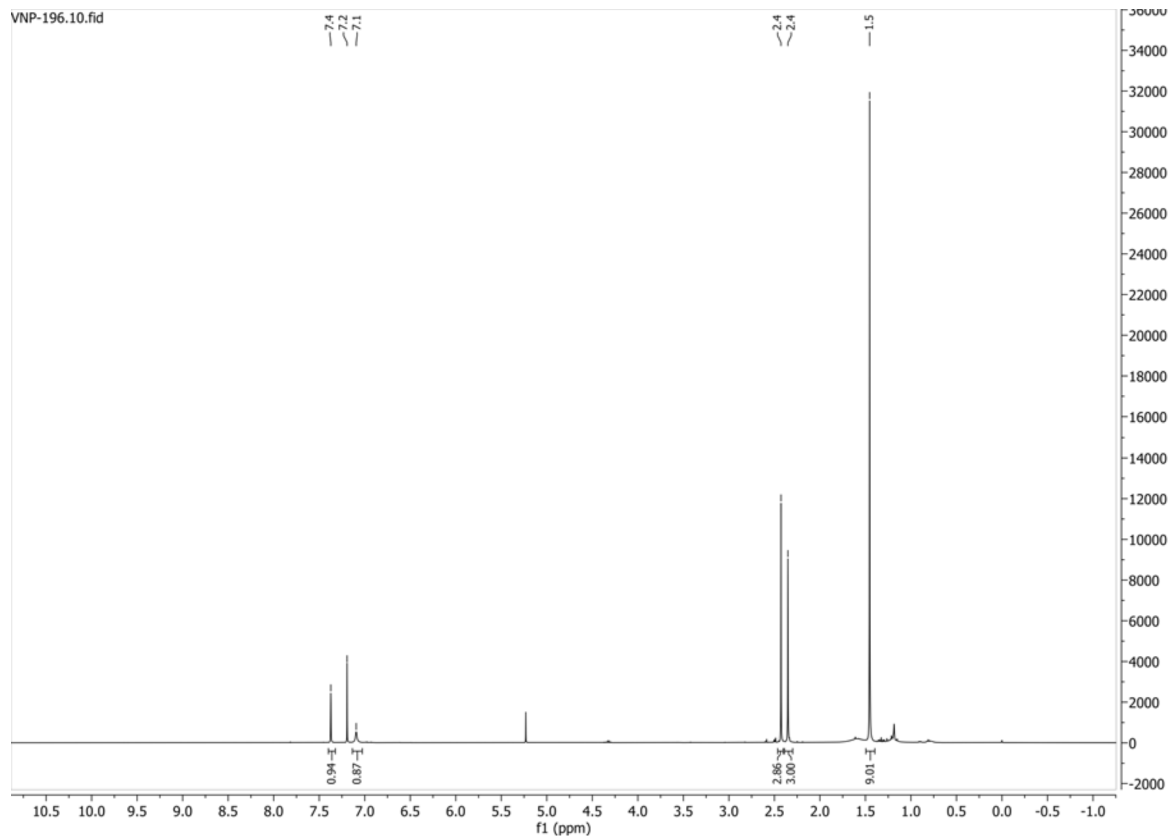


**13C{1H} spectrum of compound 31:**

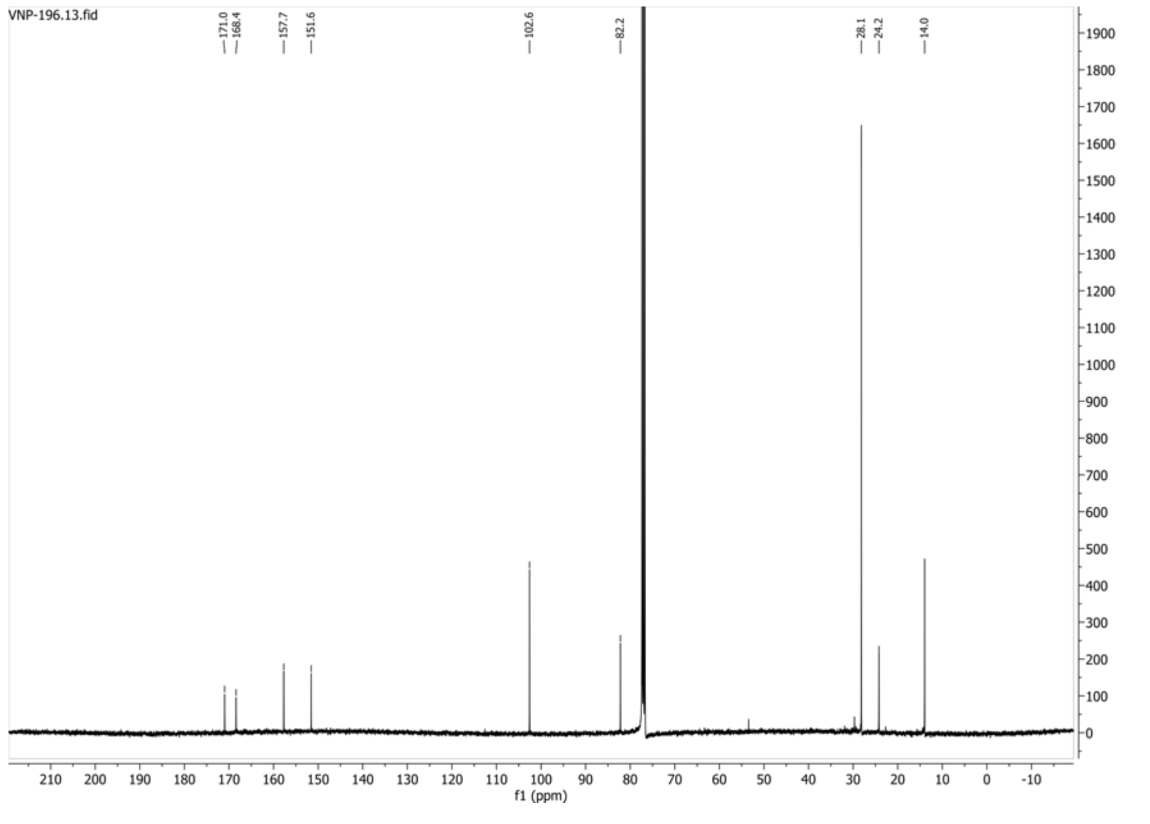

**HSQC-DEPT135 spectrum of compound 31:**

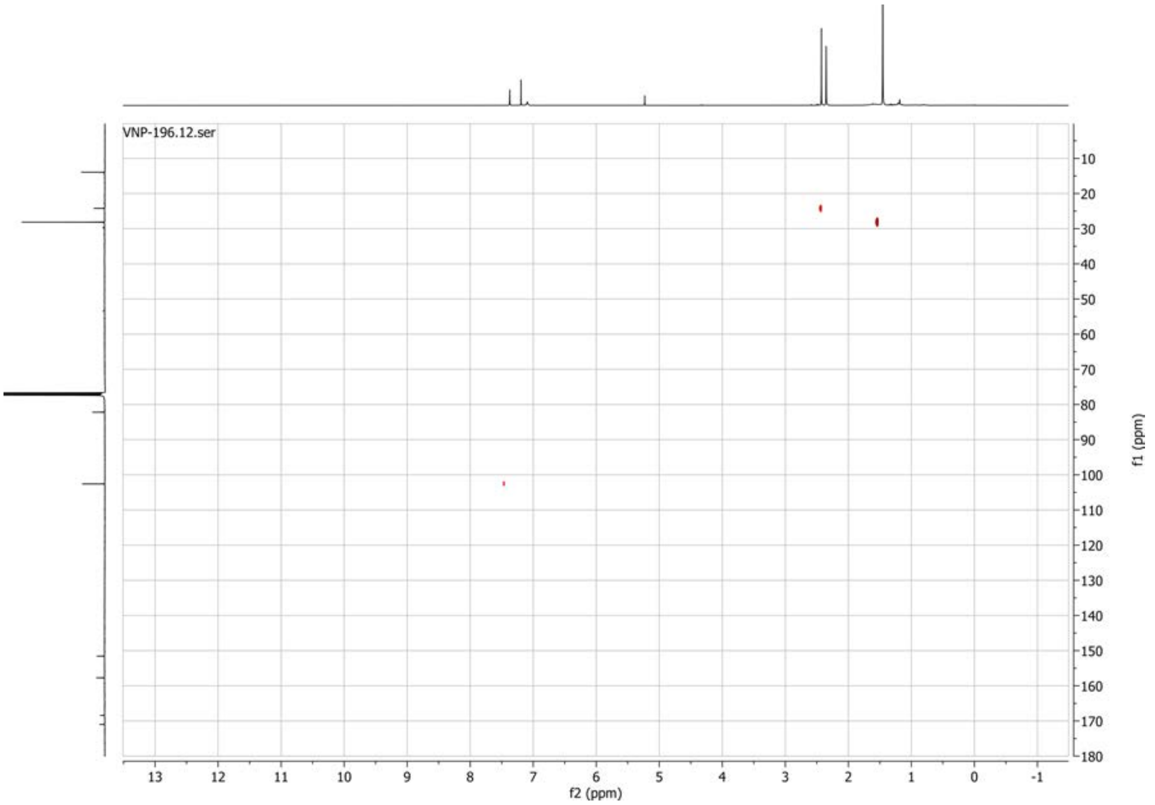

**1H spectrum of compound 32:**

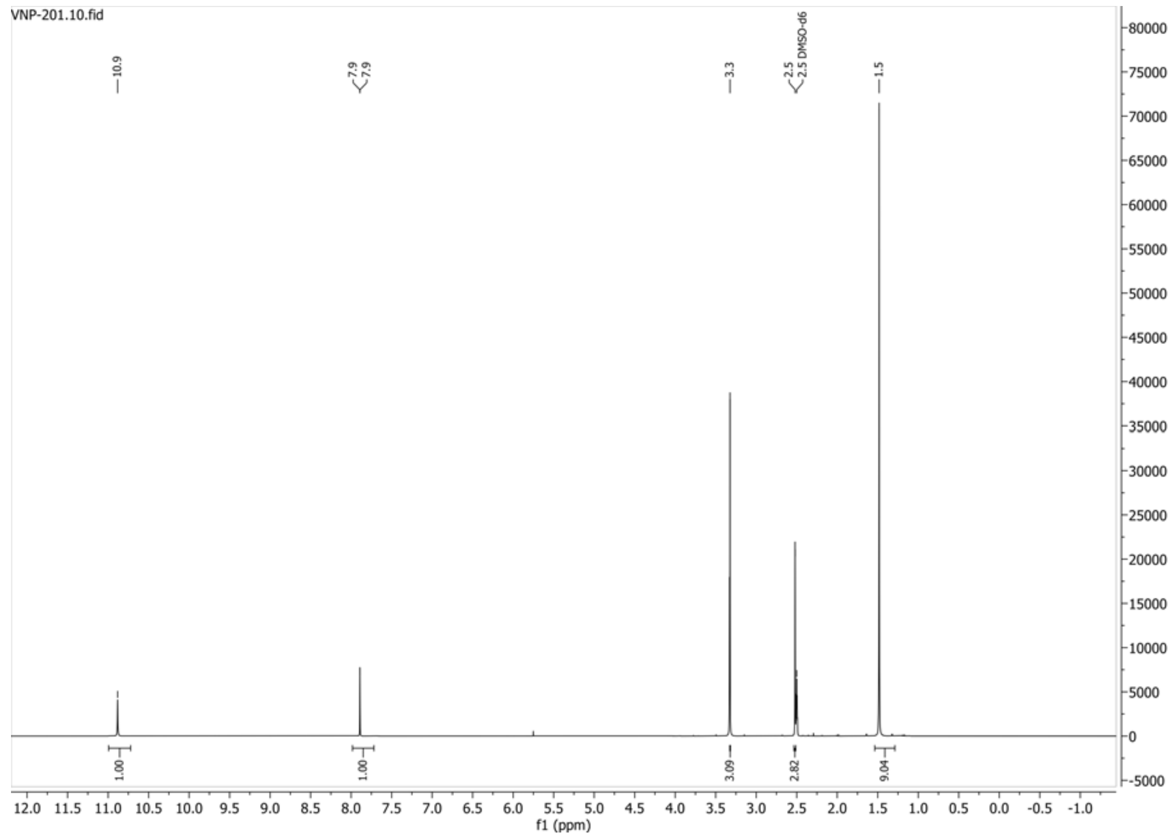


**13C{1H} spectrum of compound 32:**

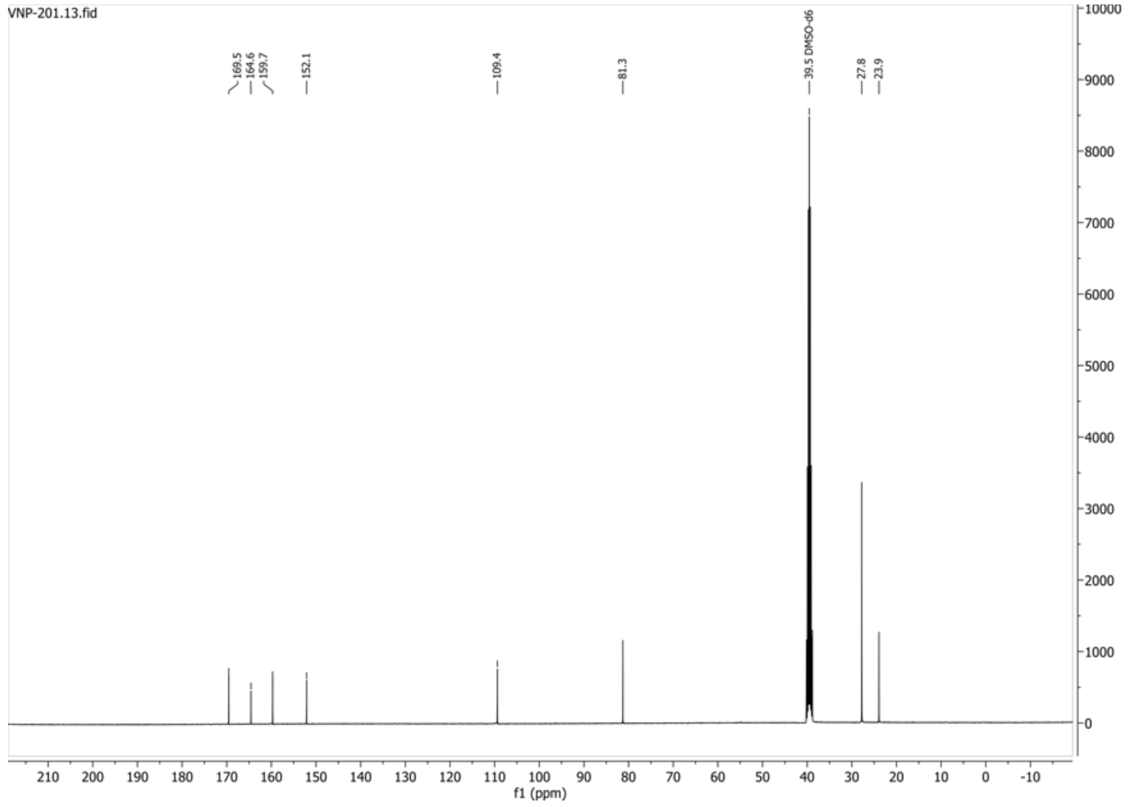

**HSQC-DEPT135 spectrum of compound 32:**

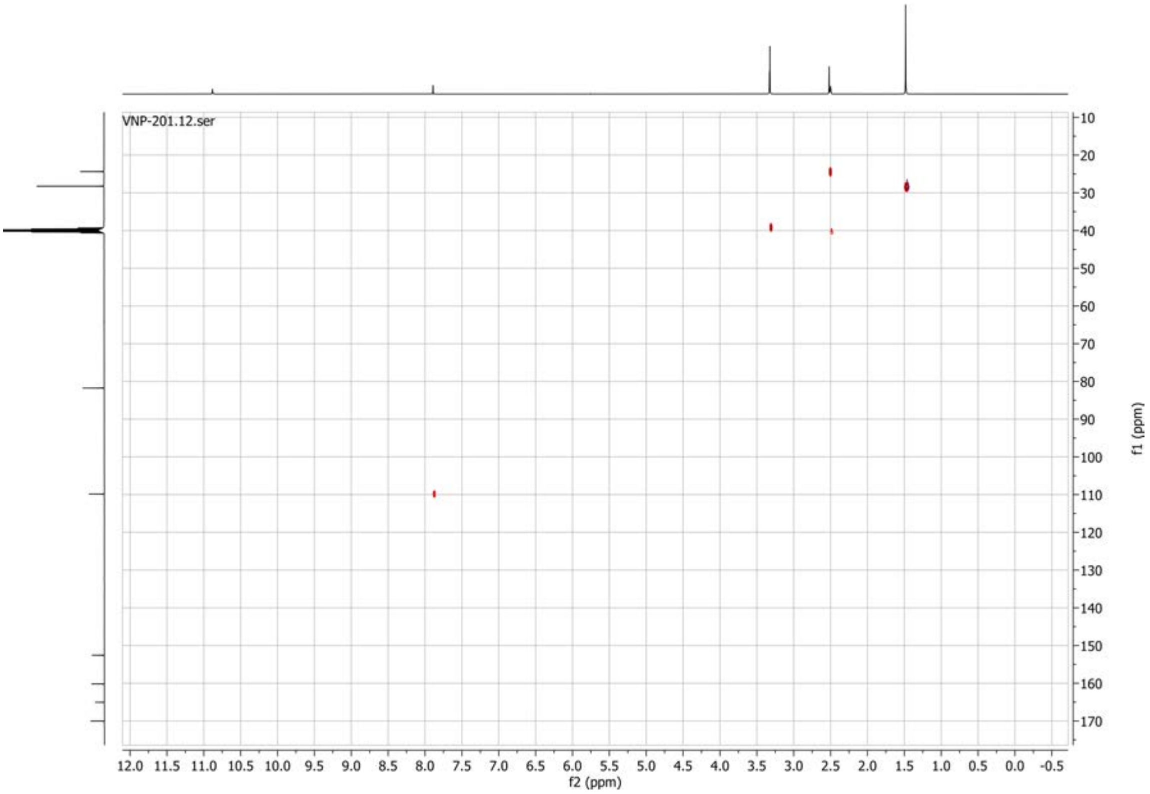

**1H spectrum of compound 34:**

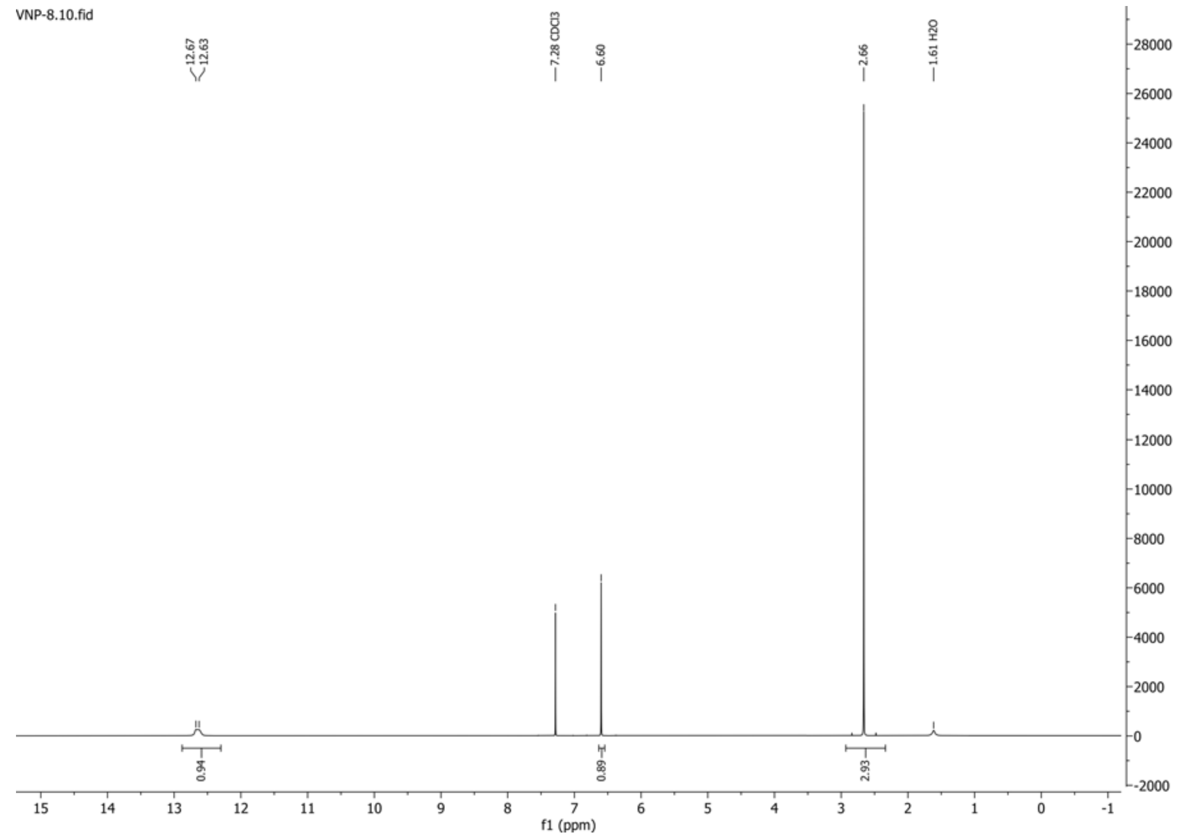


**13C{1H} spectrum of compound 34:**

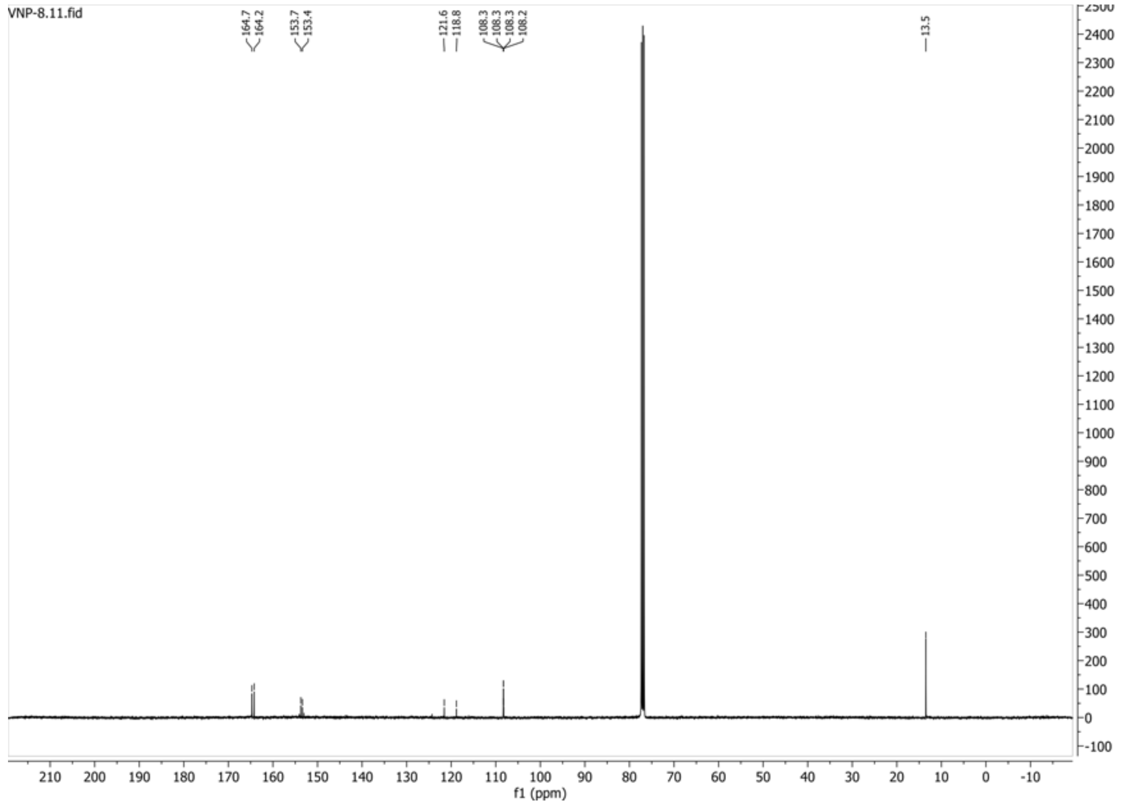

**1H spectrum of compound 35:**

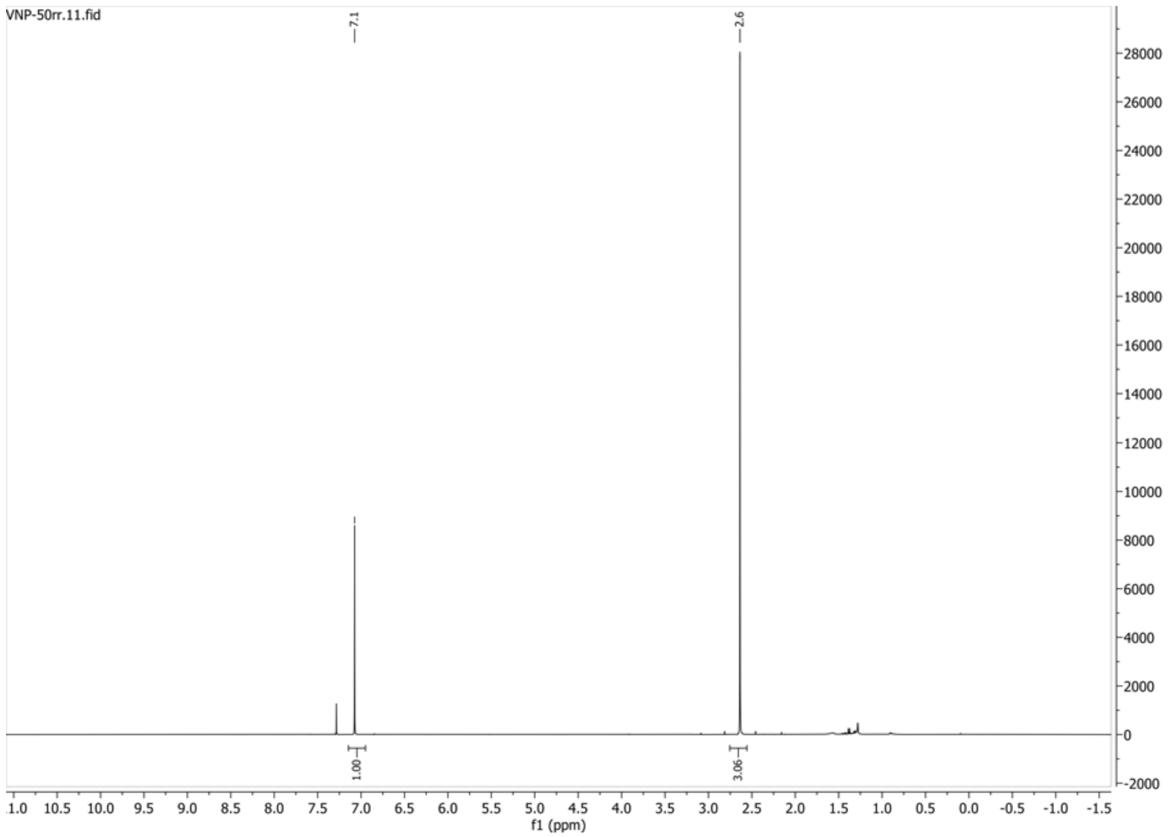


**13C{1H} spectrum of compound 35:**

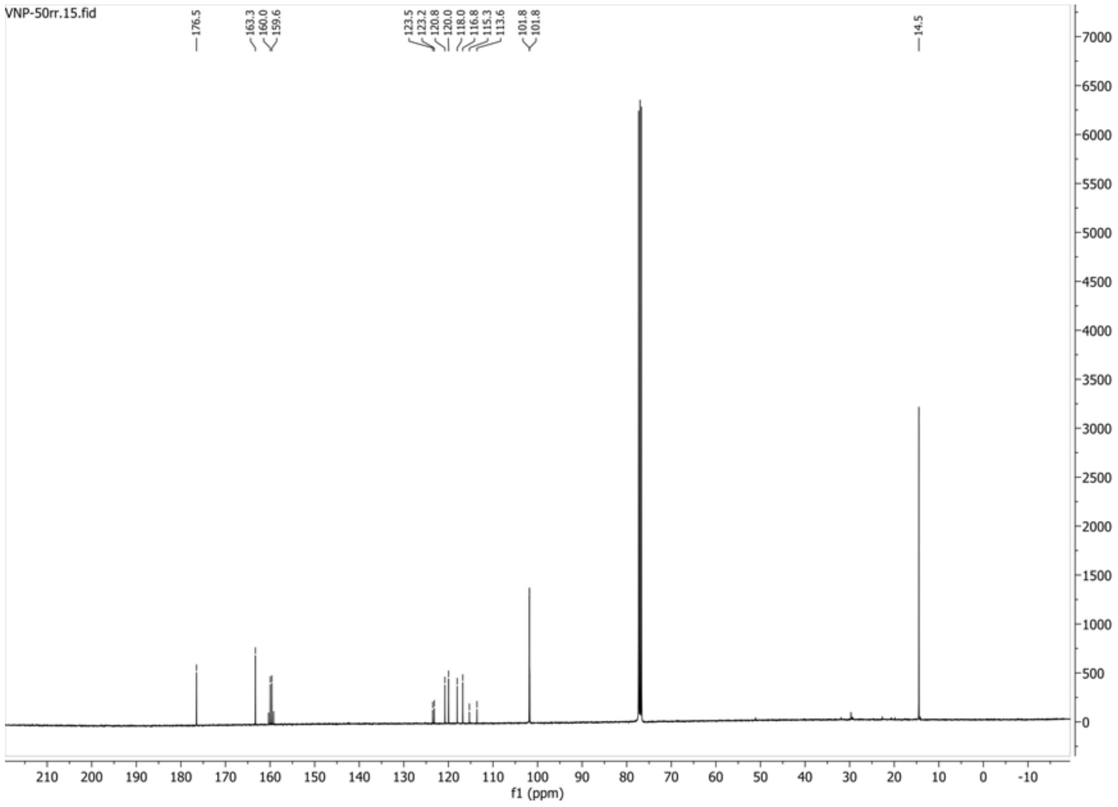

**19F{1H} spectrum of compound 35:**

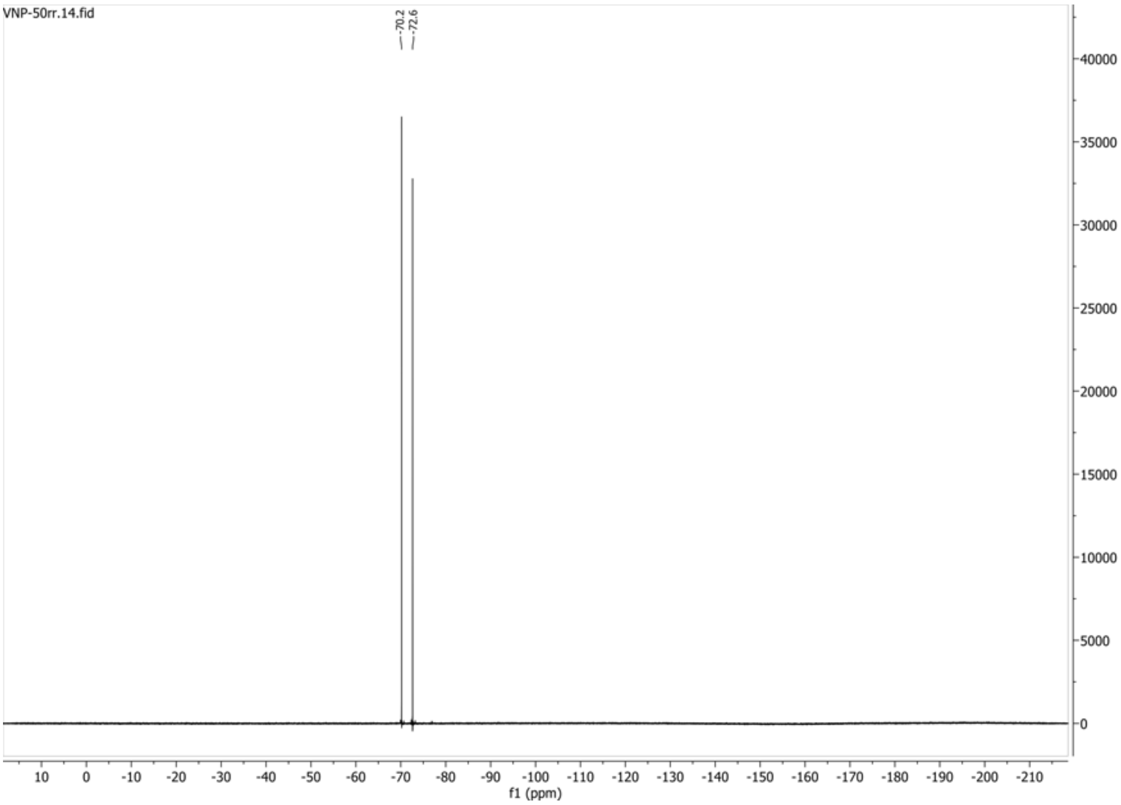


**HSQC-DEPT135 spectrum of compound 35:**

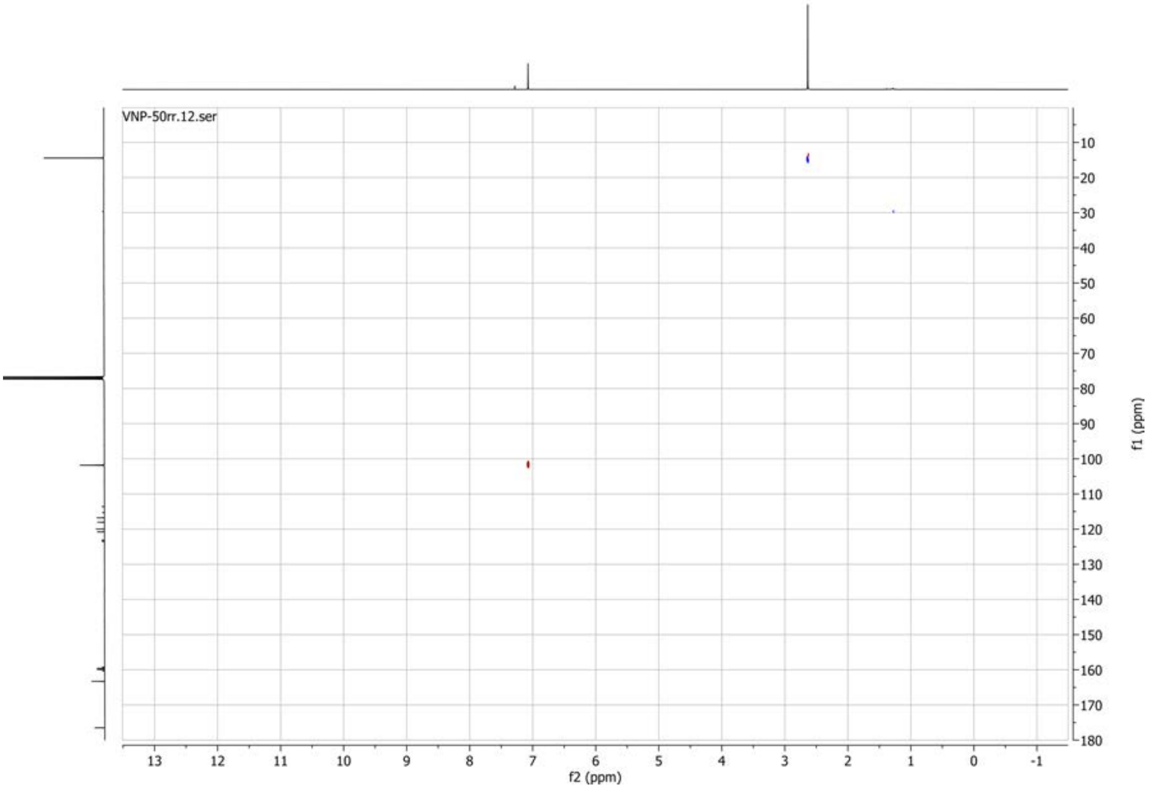

**1H spectrum of compound 39:**

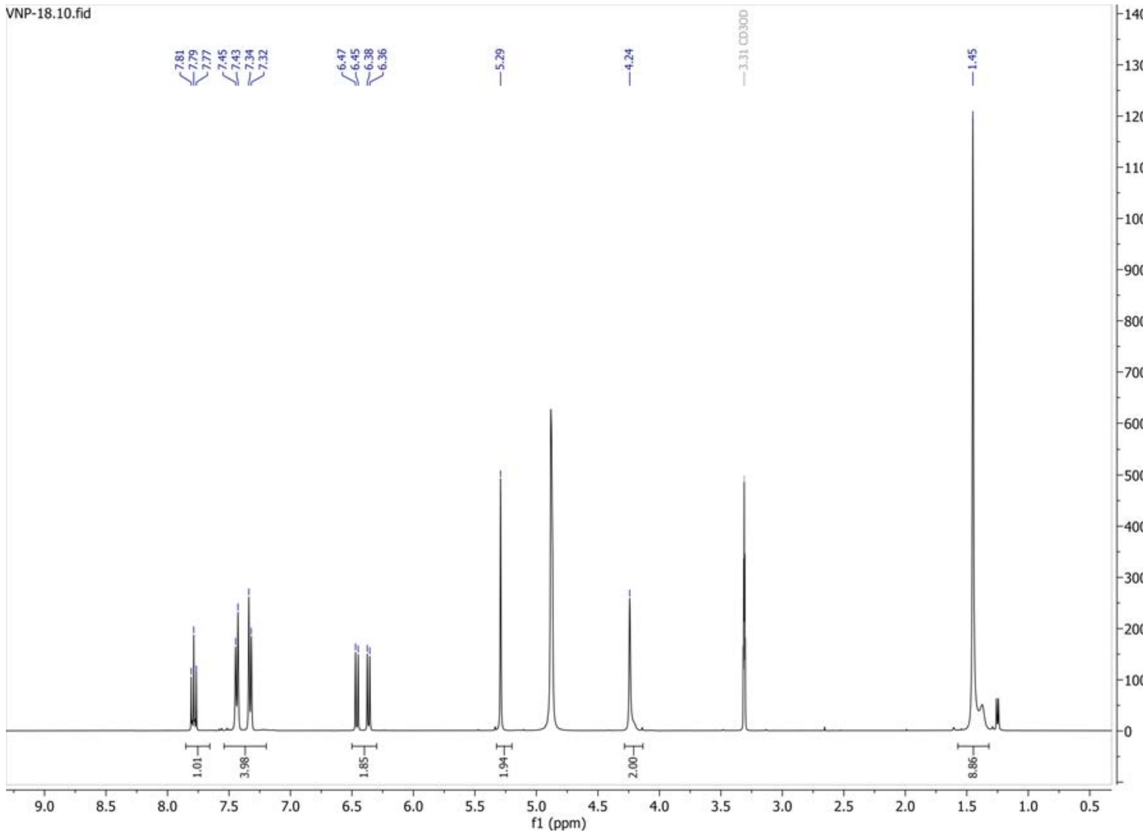


**13C{1H} spectrum of compound 39:**

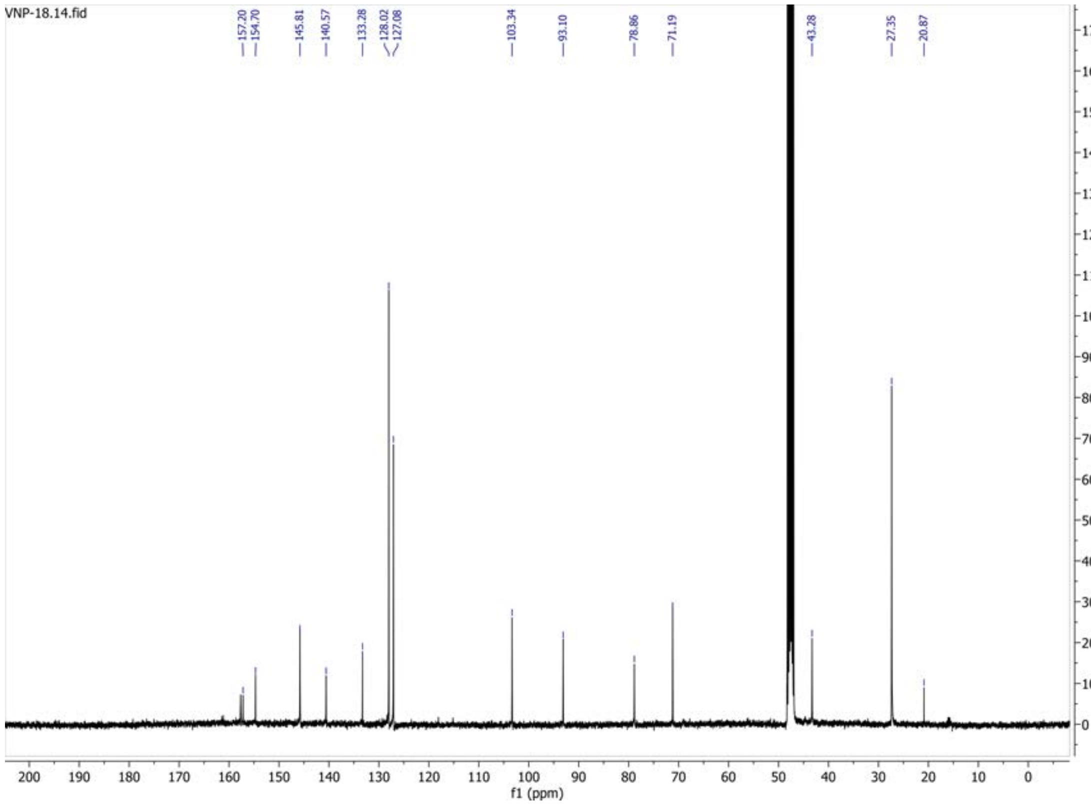

**DEPT135 spectrum of compound 39:**

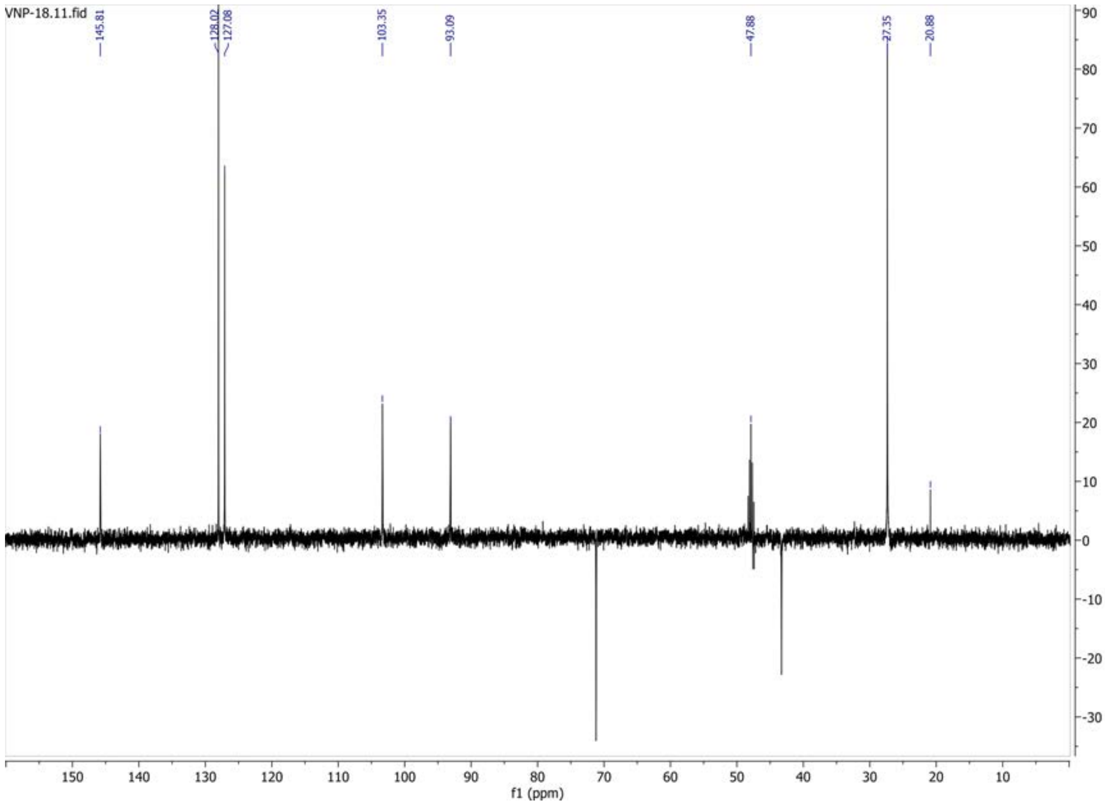

**1H spectrum of compound 41:** *small amount of EtOAc as impurity

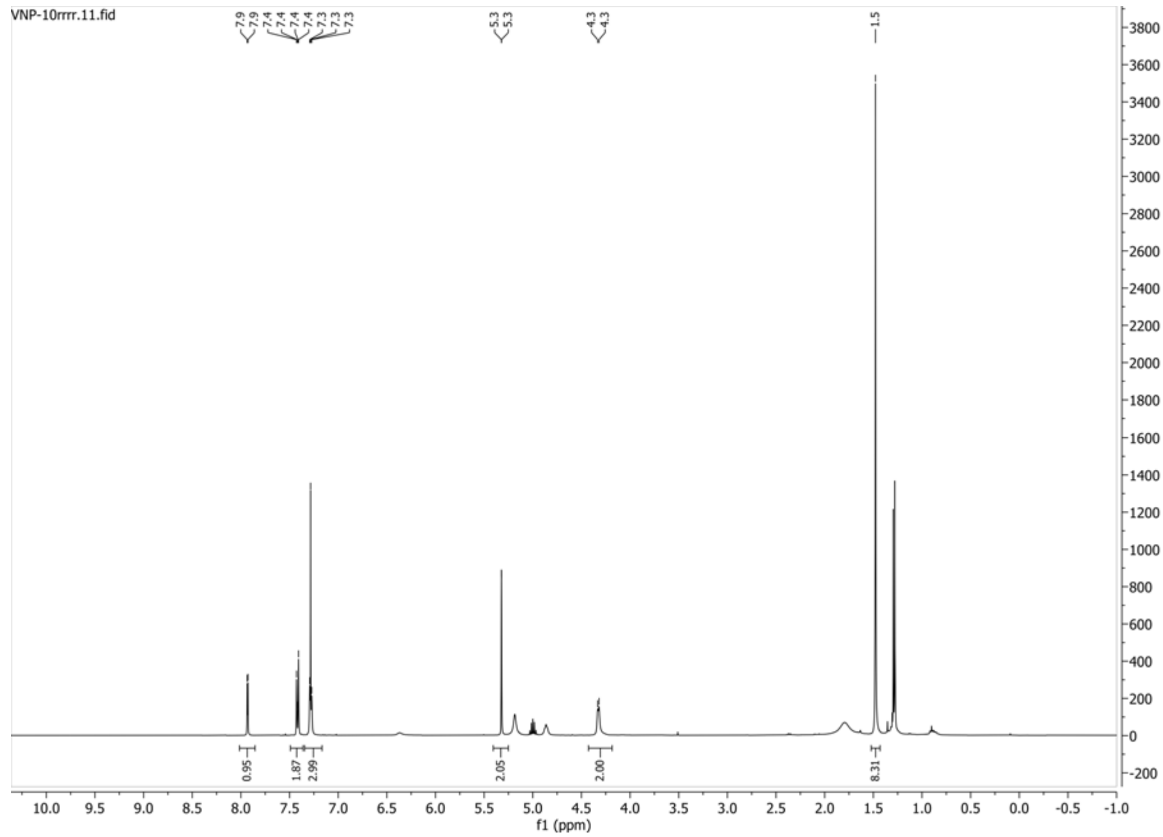


**13C{1H} spectrum of compound 41:** *small amount of EtOAc as impurity

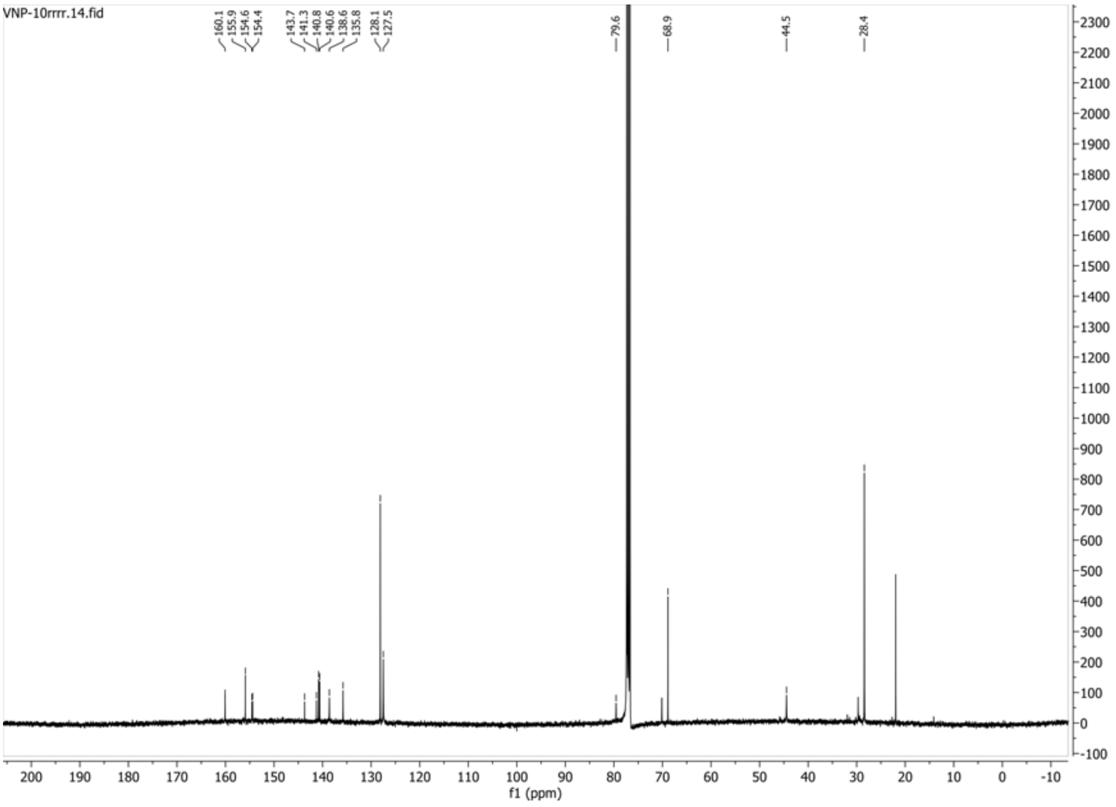

**HSQC-DEPT spectrum of compound 41:**

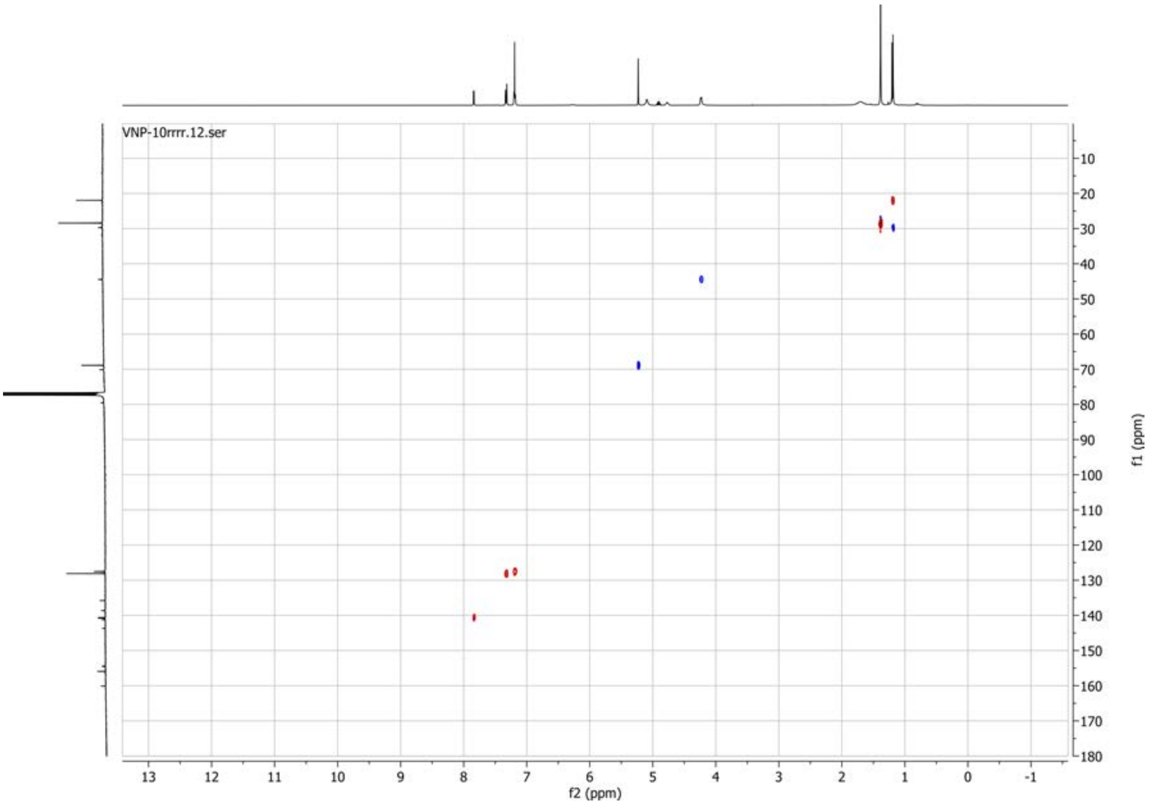


**19F spectrum of compound 41:**

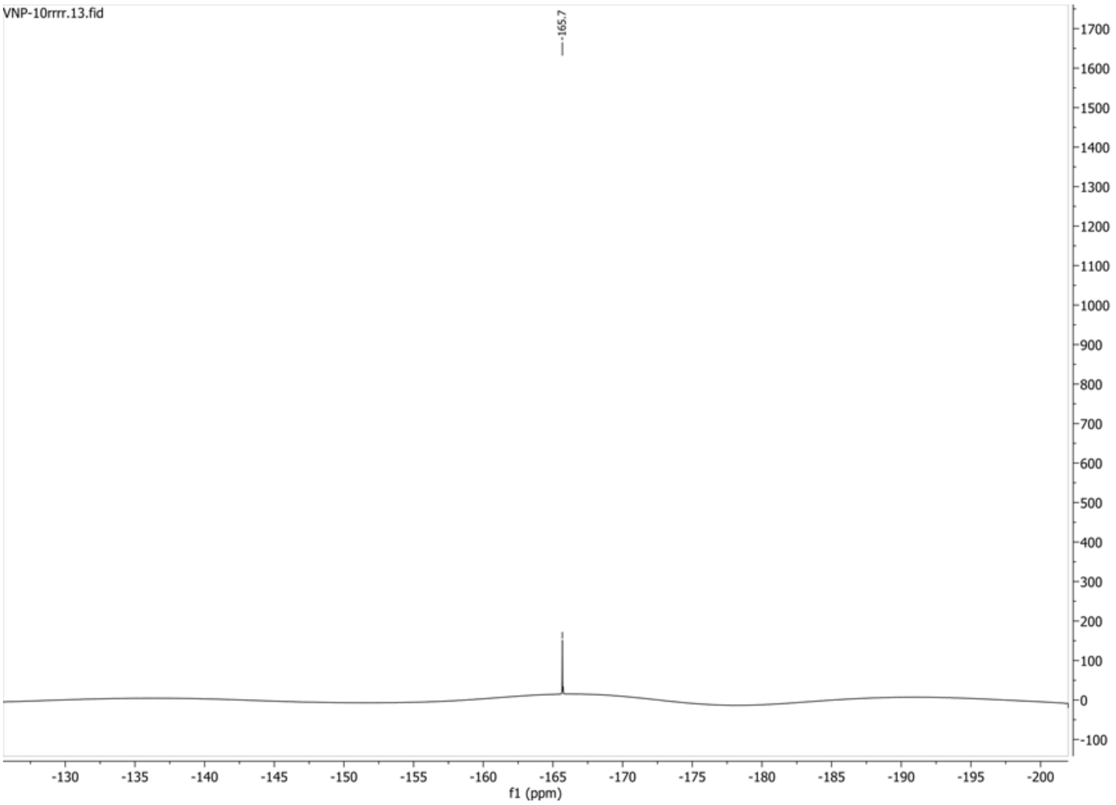

**1H spectrum of compound 36:**

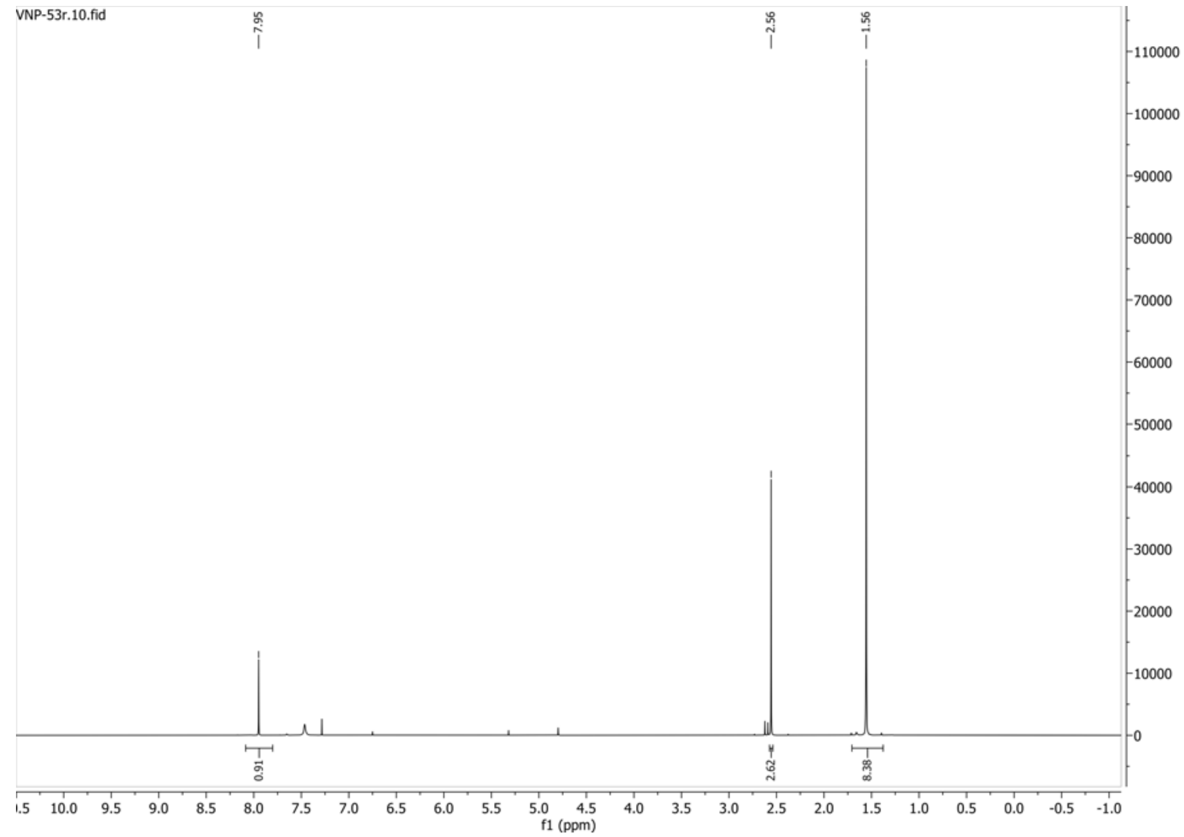


**13C{1H} spectrum of compound 36:**

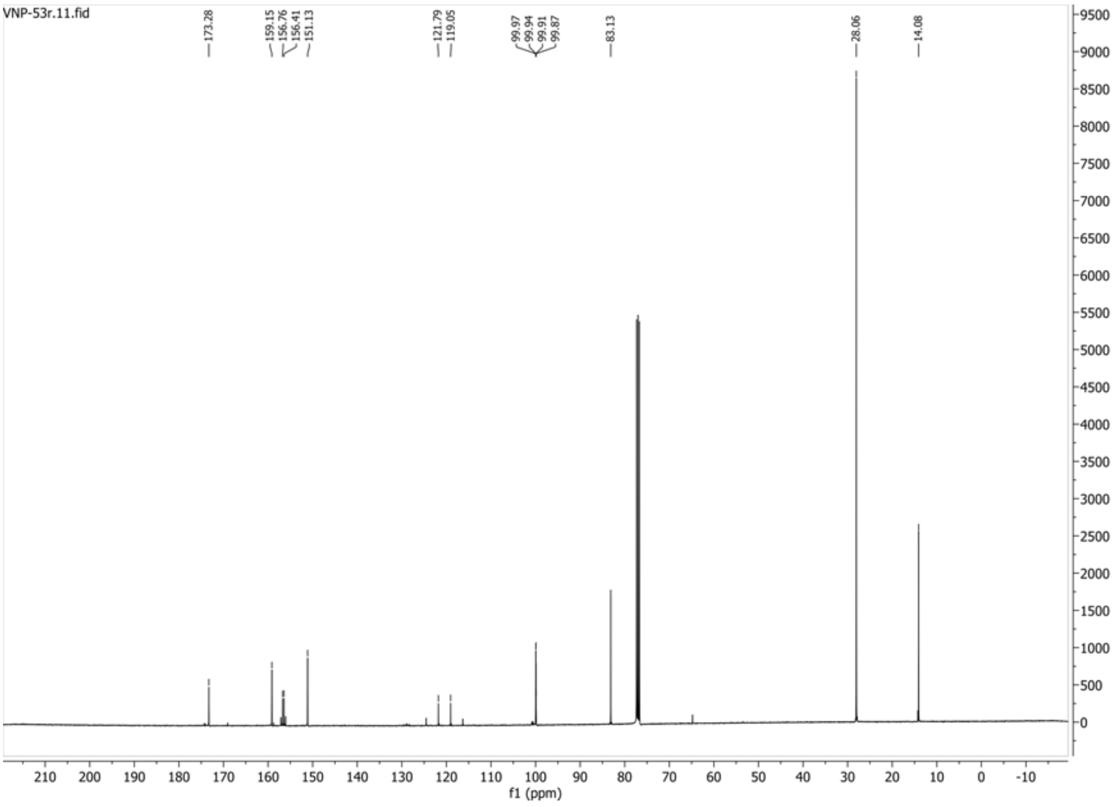

**19F spectrum of compound 36:**

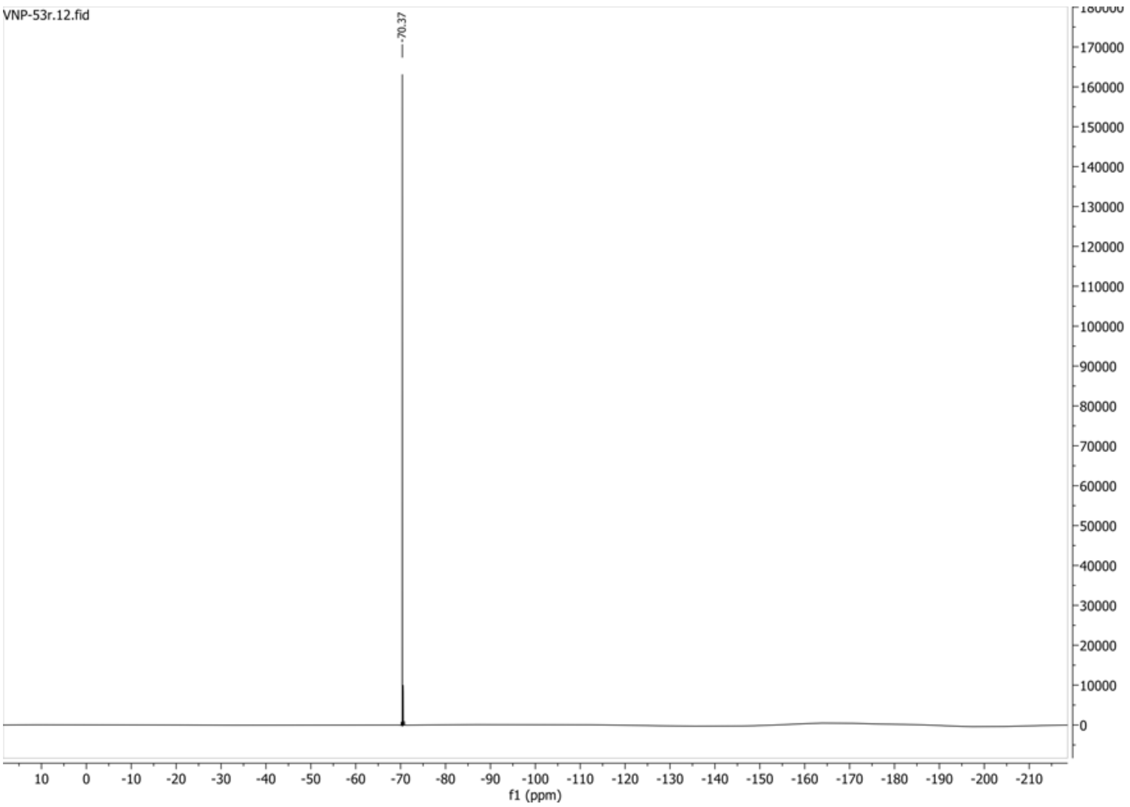

**1H spectrum of compound 37:**

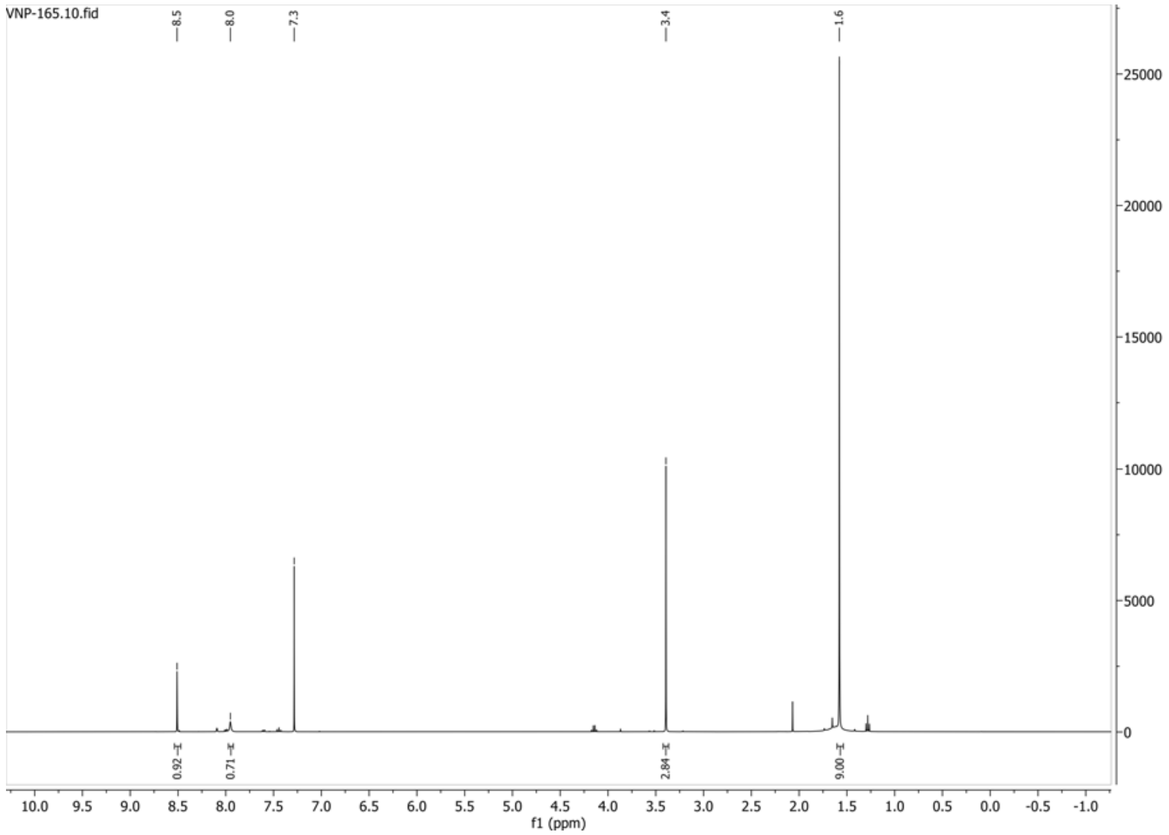


**13C{1H} spectrum of compound 37:**

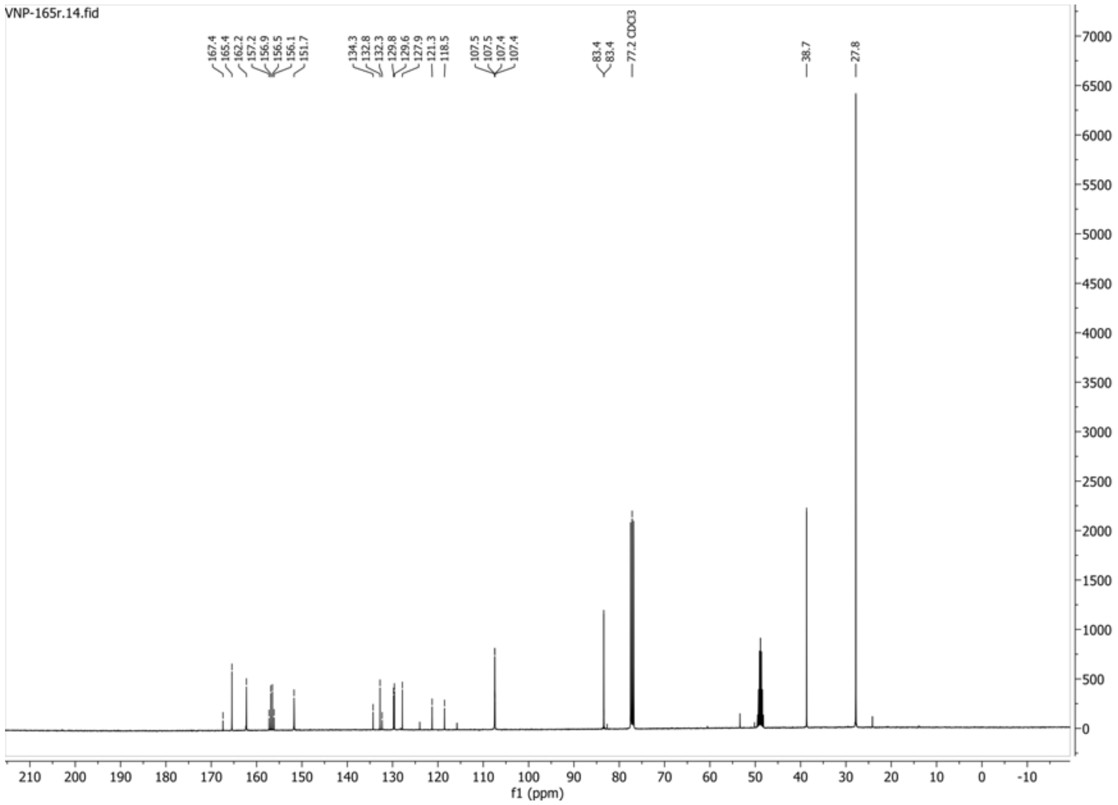

**19F spectrum of compound 37:**

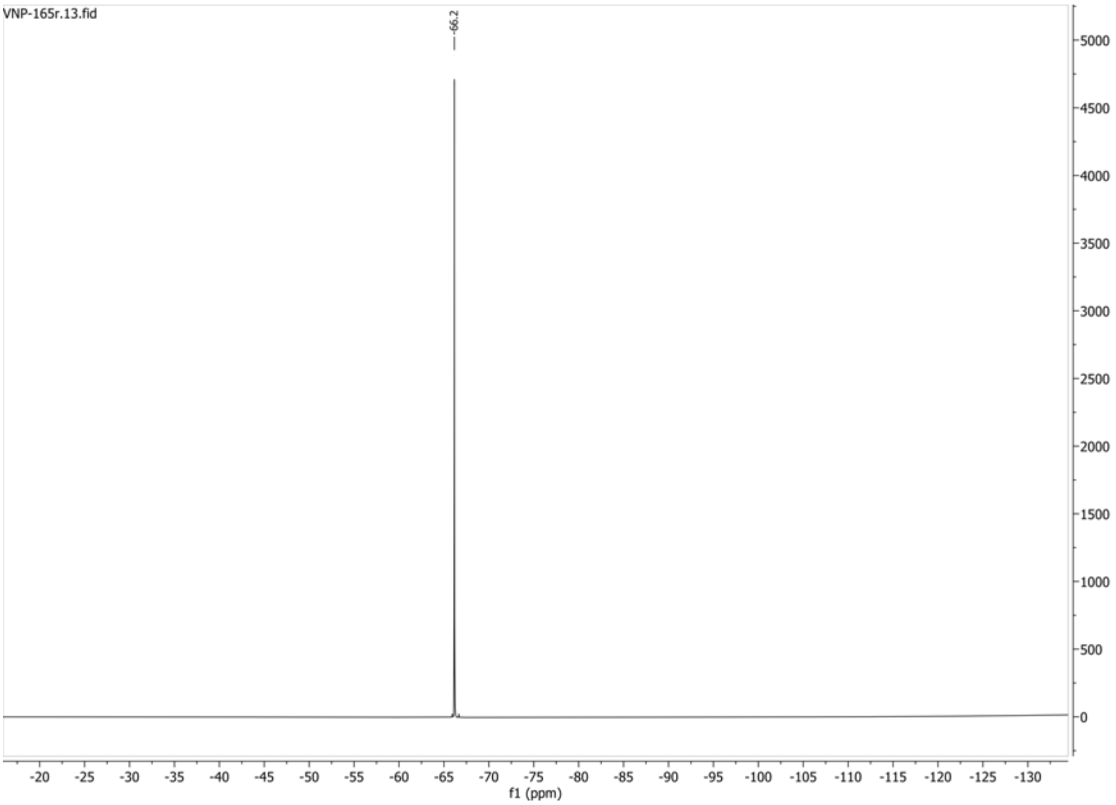

**1H spectrum of compound 48:**

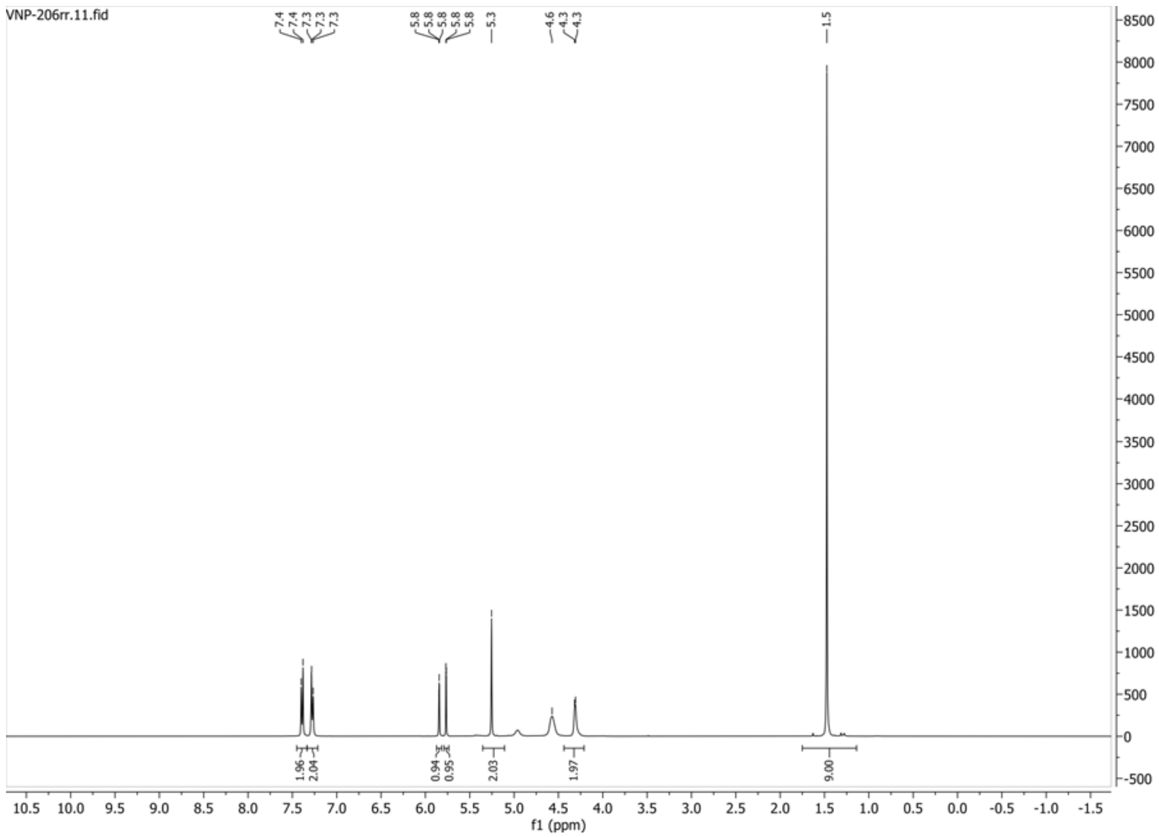


**13C{1H} spectrum of compound 48:**

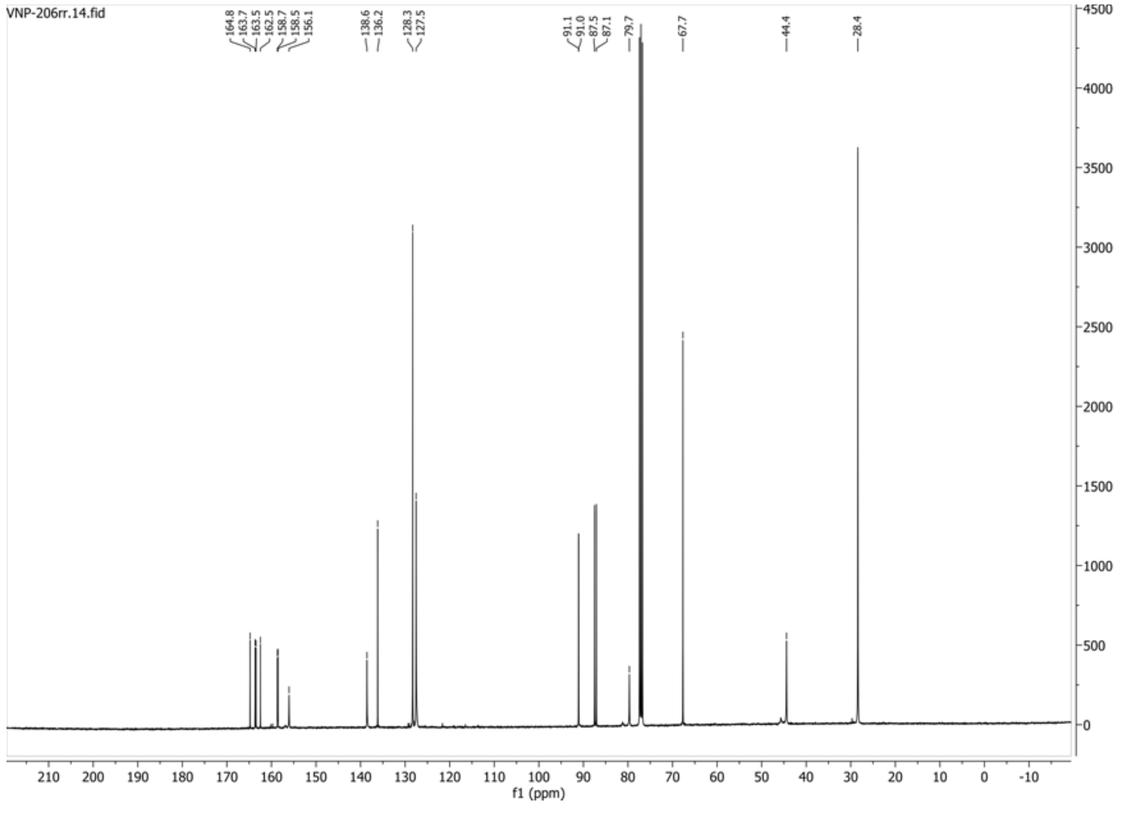

**HSQC-DEPT135 spectrum of compound 48:**

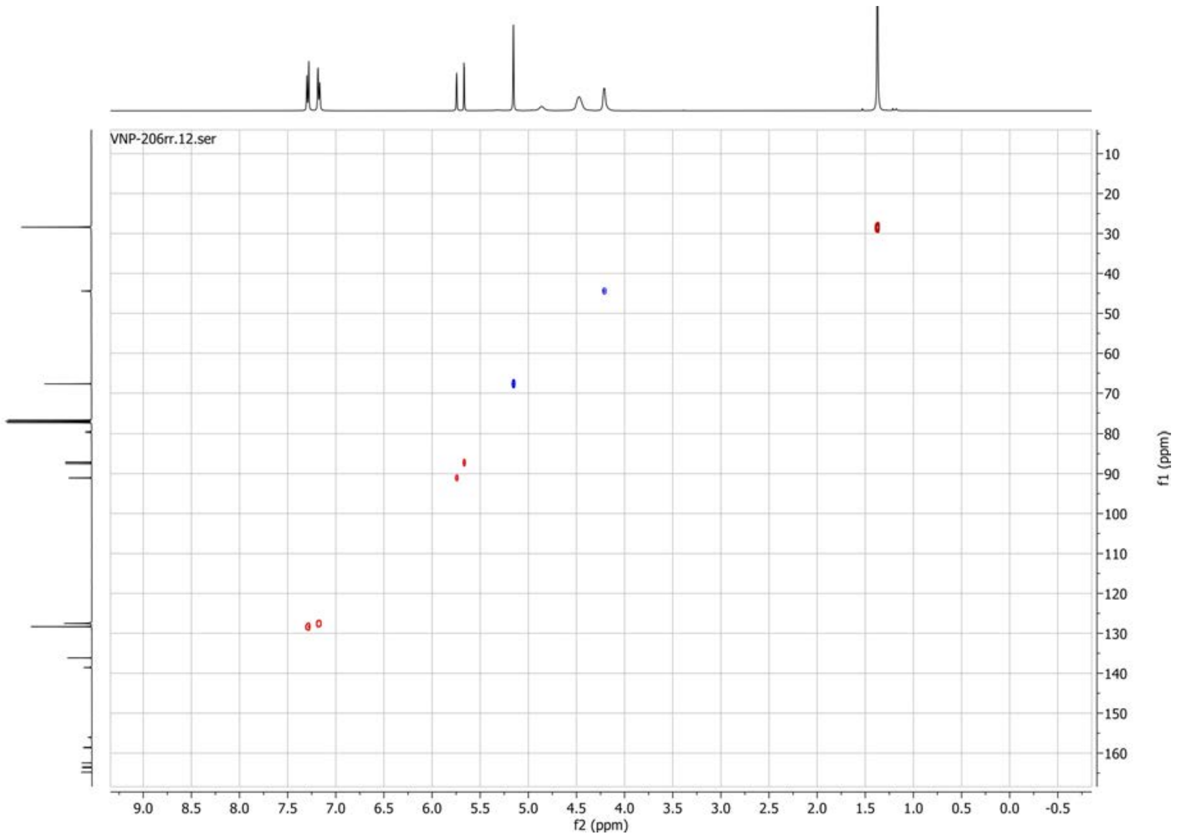


**19F spectrum of compound 48:**

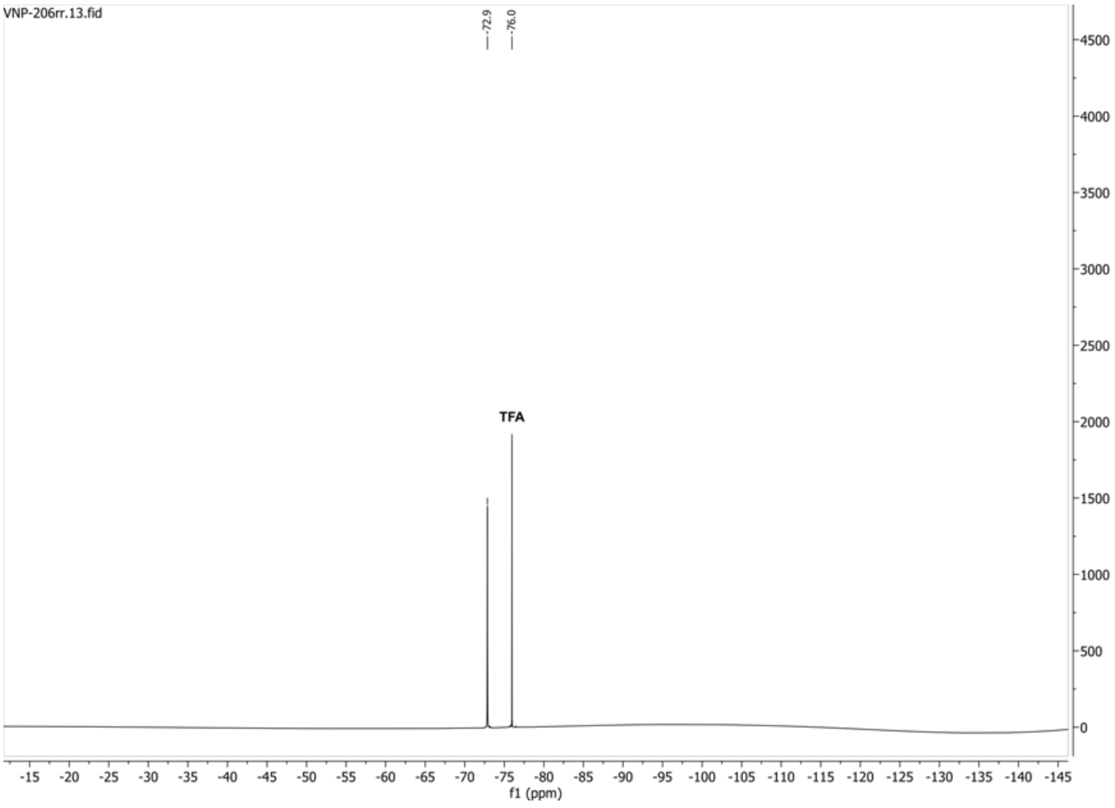

**1H spectrum of compound 52:**

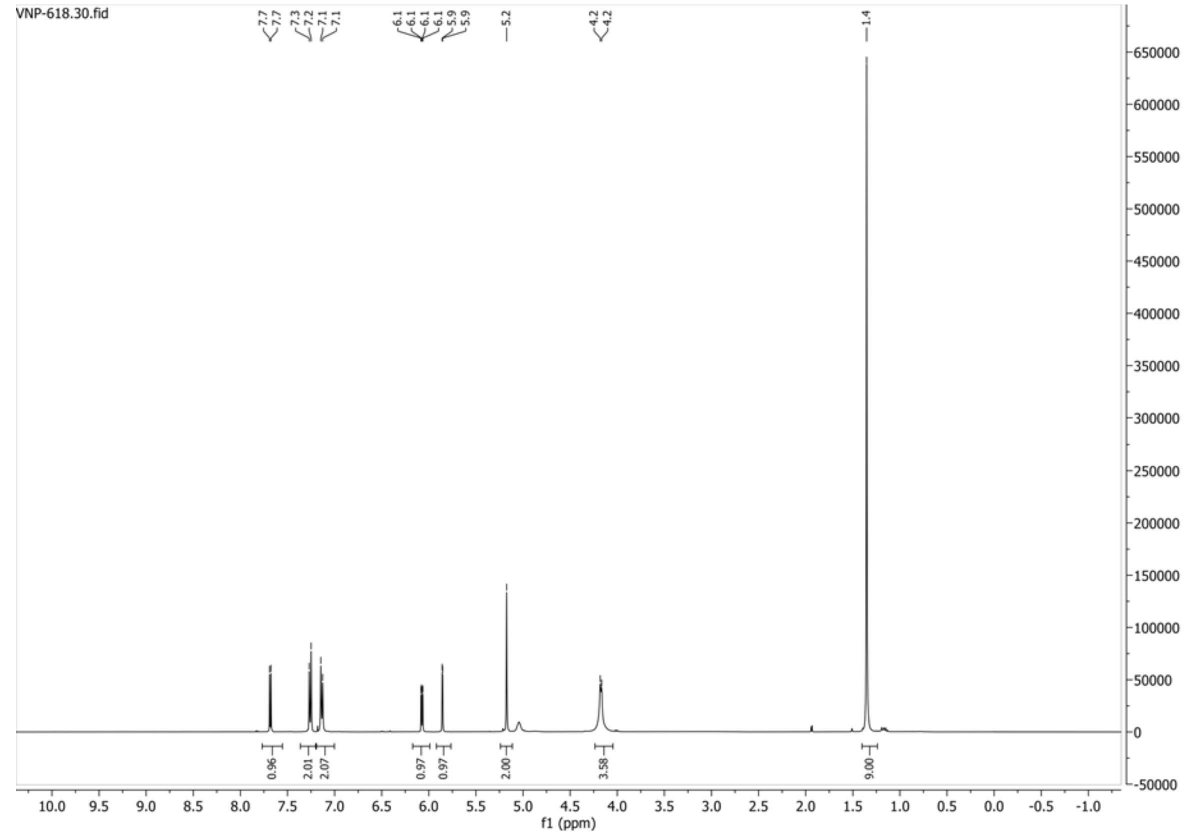


**13C{1H} spectrum of compound 52:**

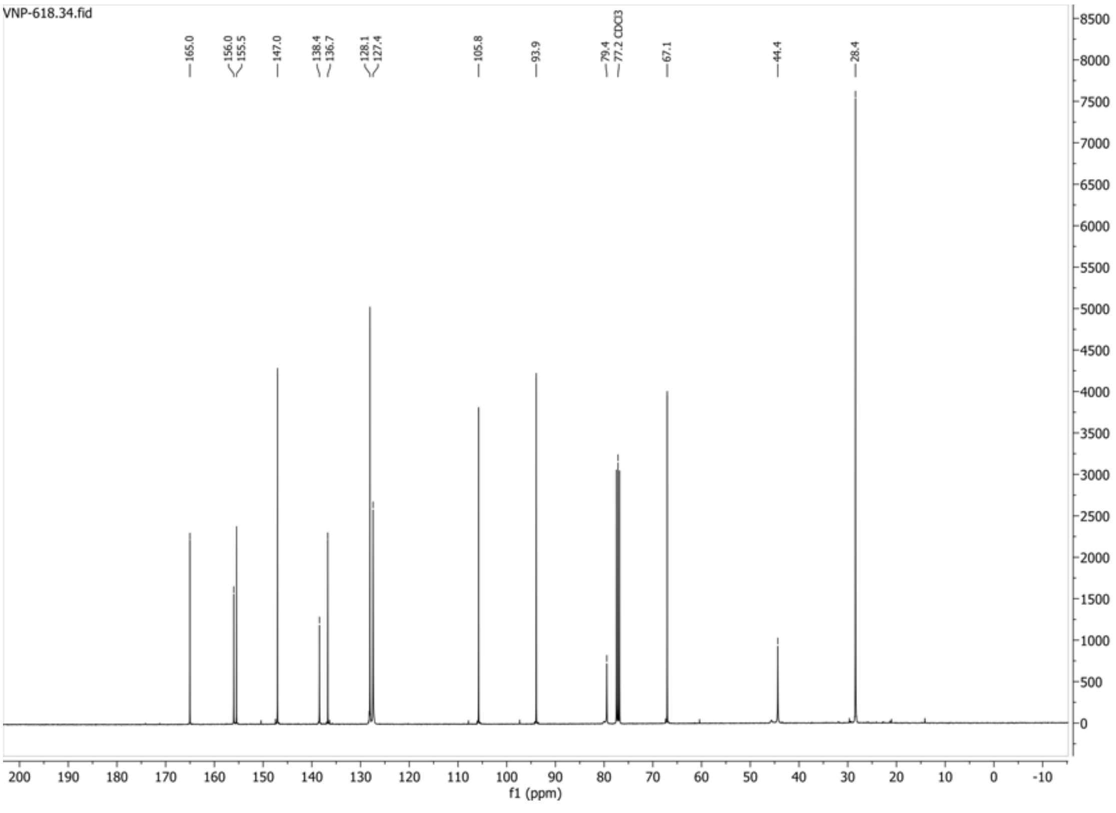

**HSQC-DEPT spectrum of compound 52:**

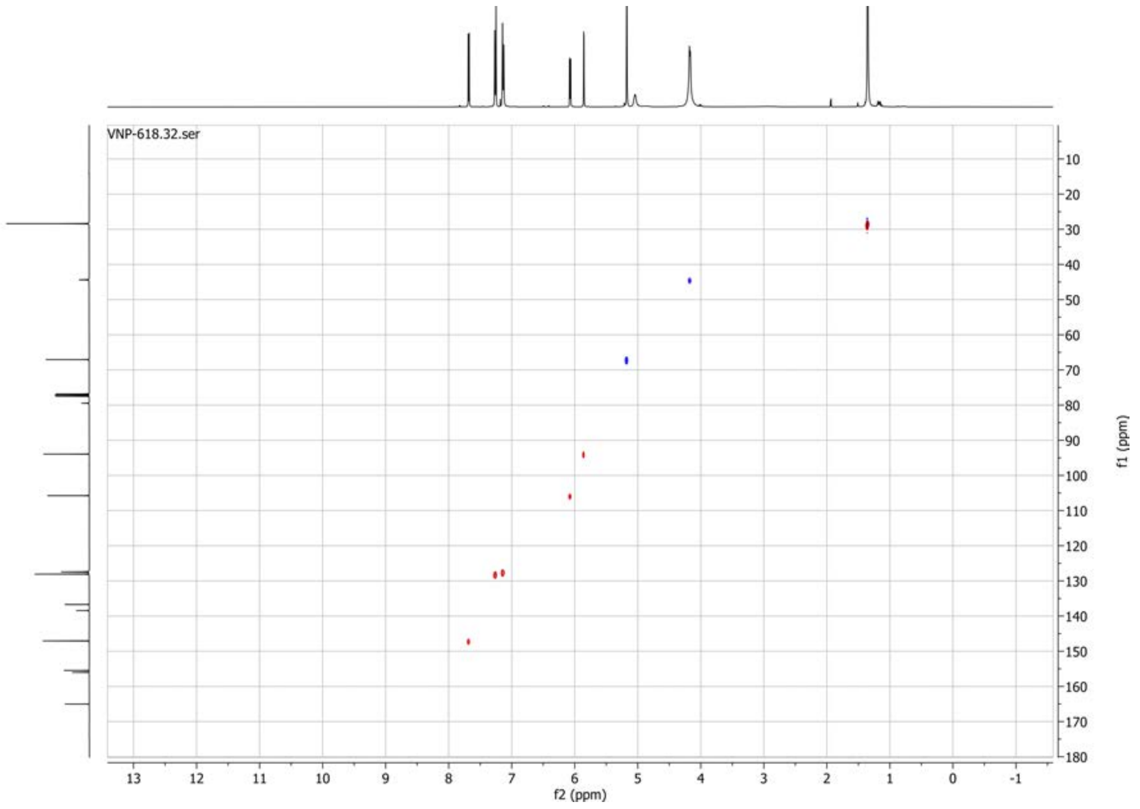

**1H spectrum of compound 56:**

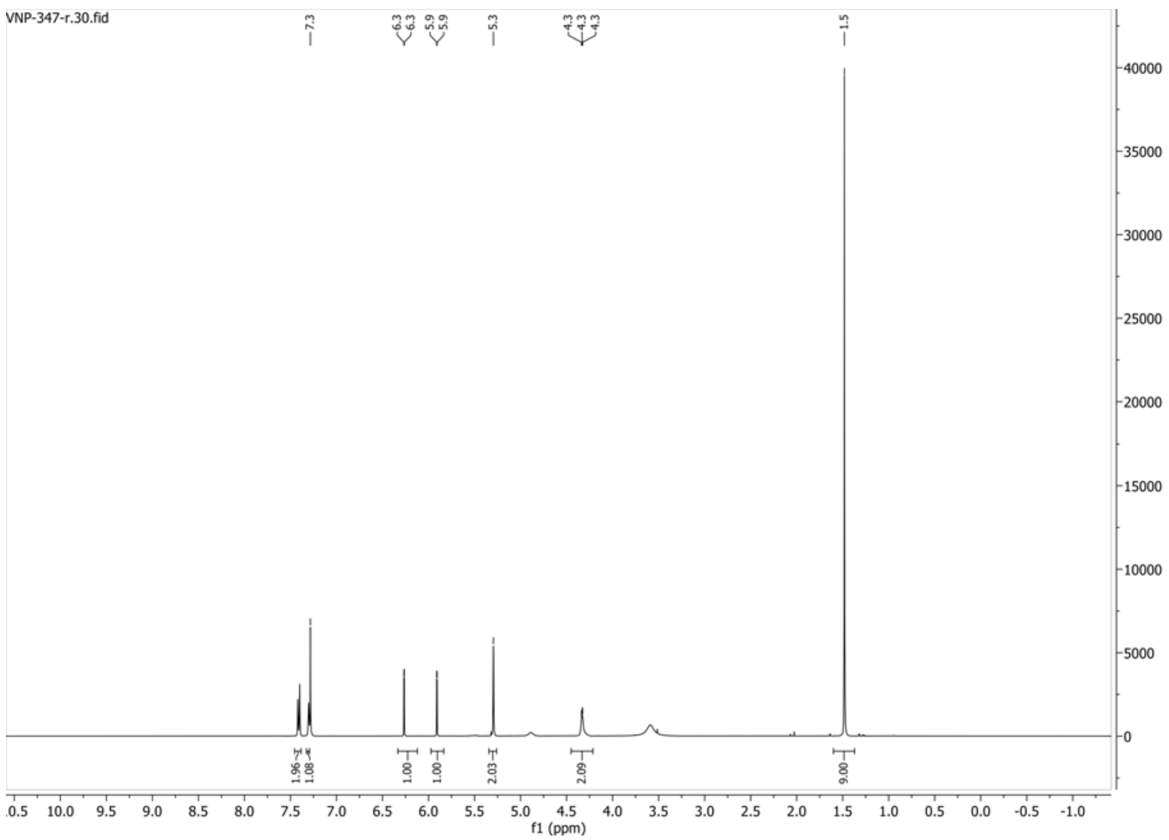


**13C{1H} spectrum of compound 56:**

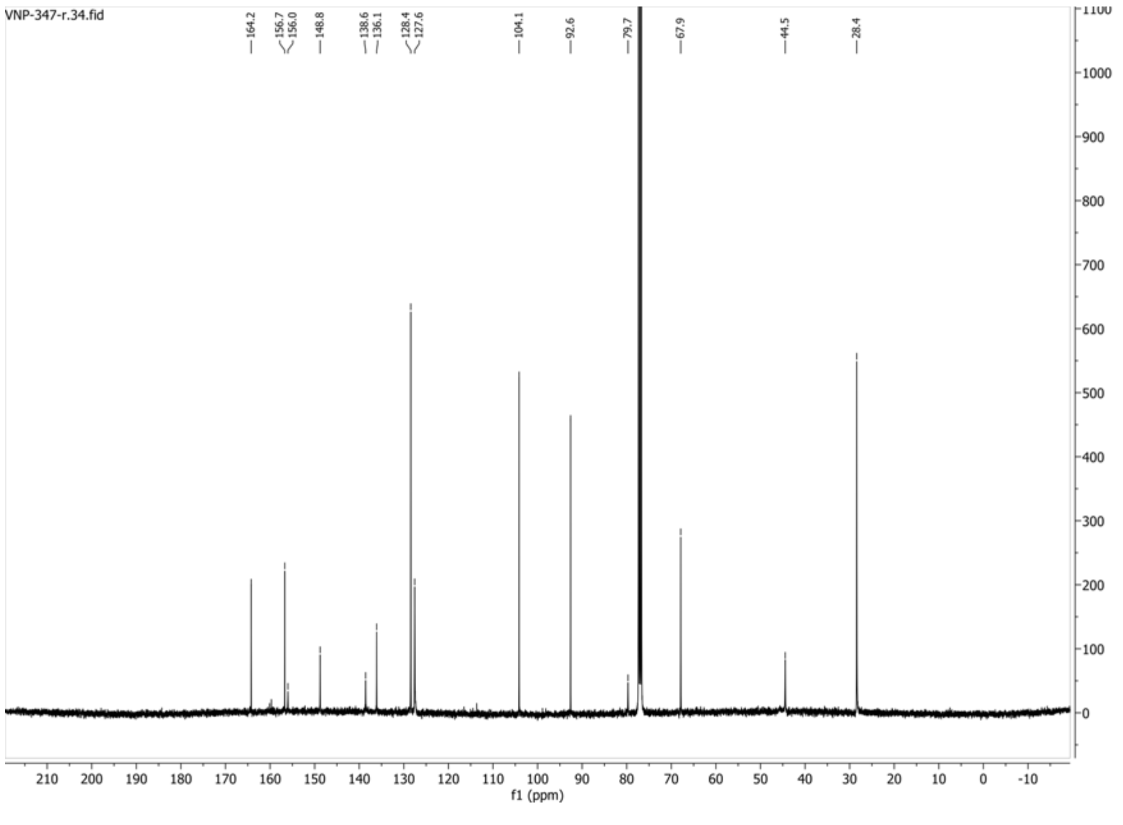

**HSQC-DEPT135 spectrum of compound 56:**

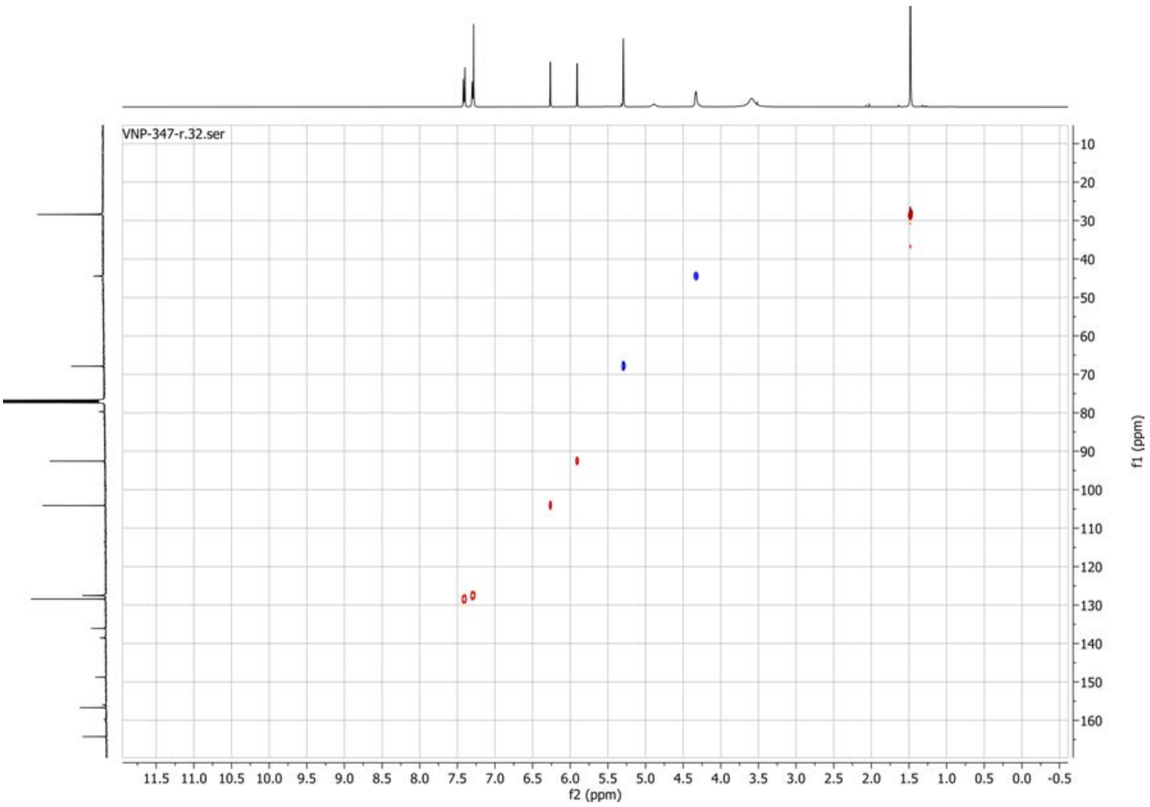

#### 1H spectrum of compound 57:

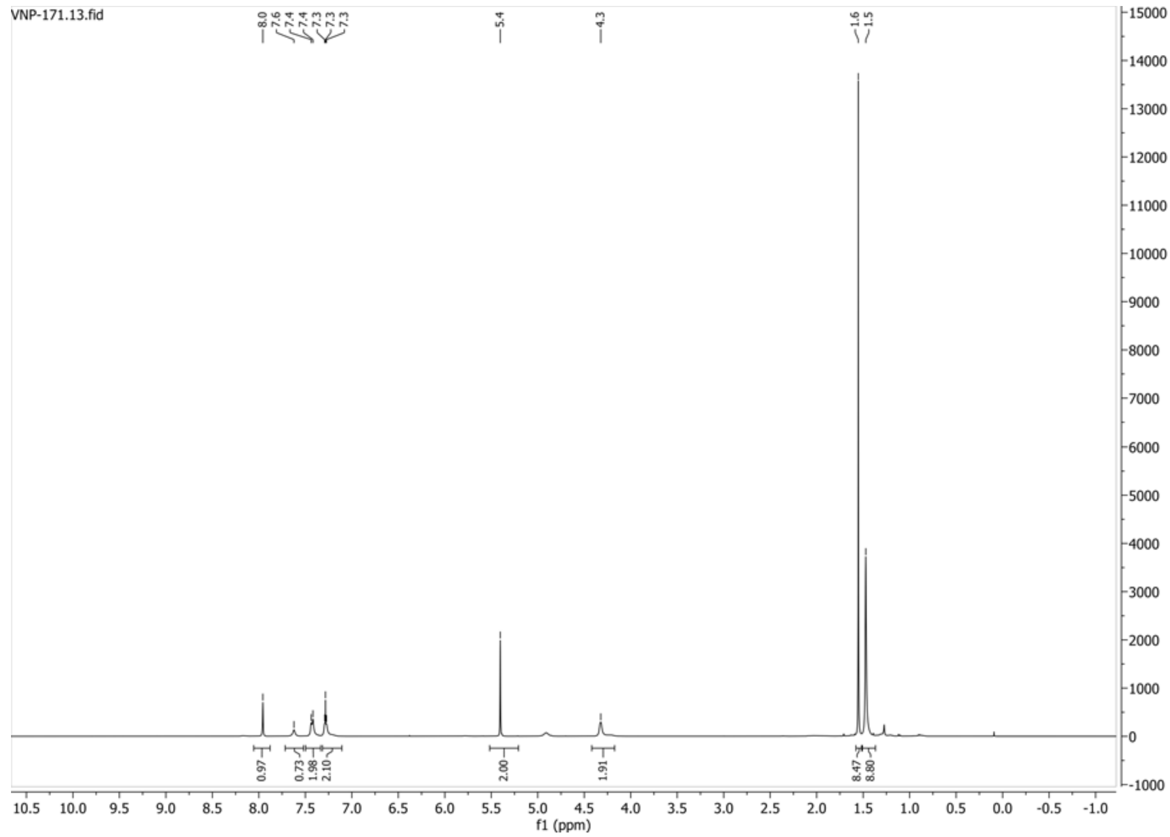


#### 13C{1H} spectrum of compound 57:

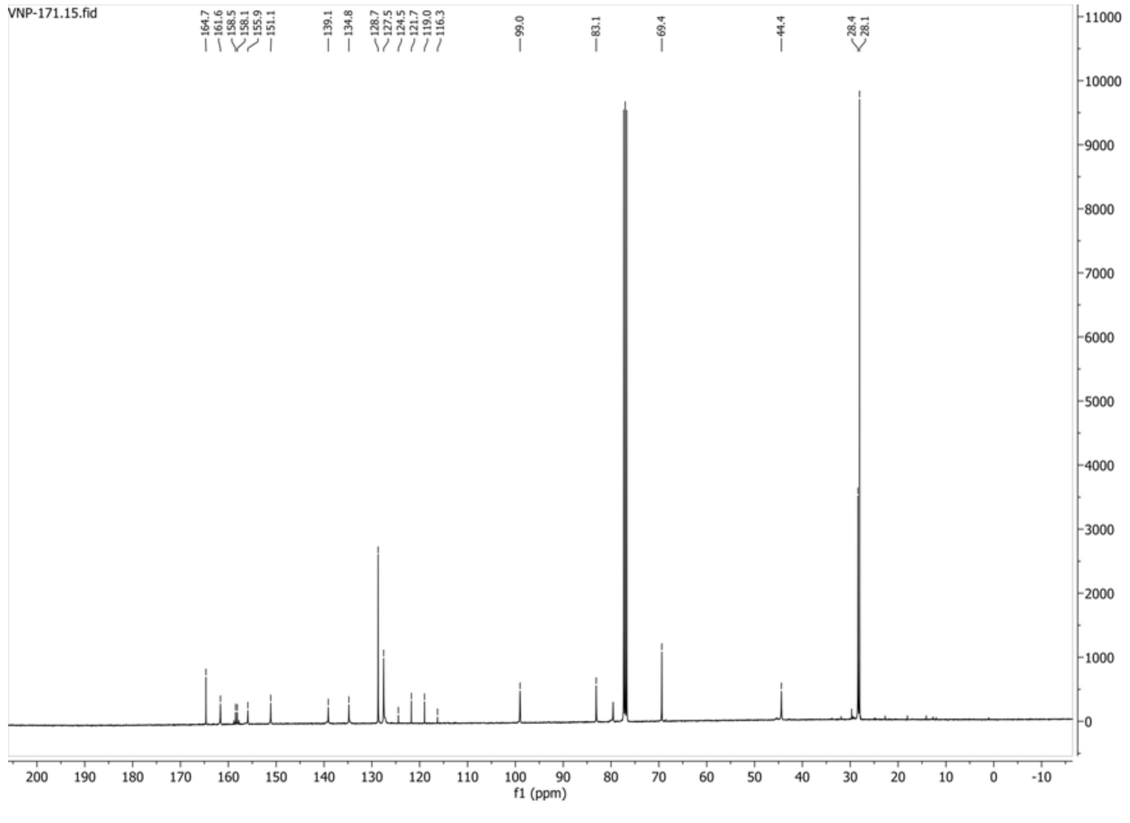

**HSQC-DEPT135 spectrum of compound 57:**

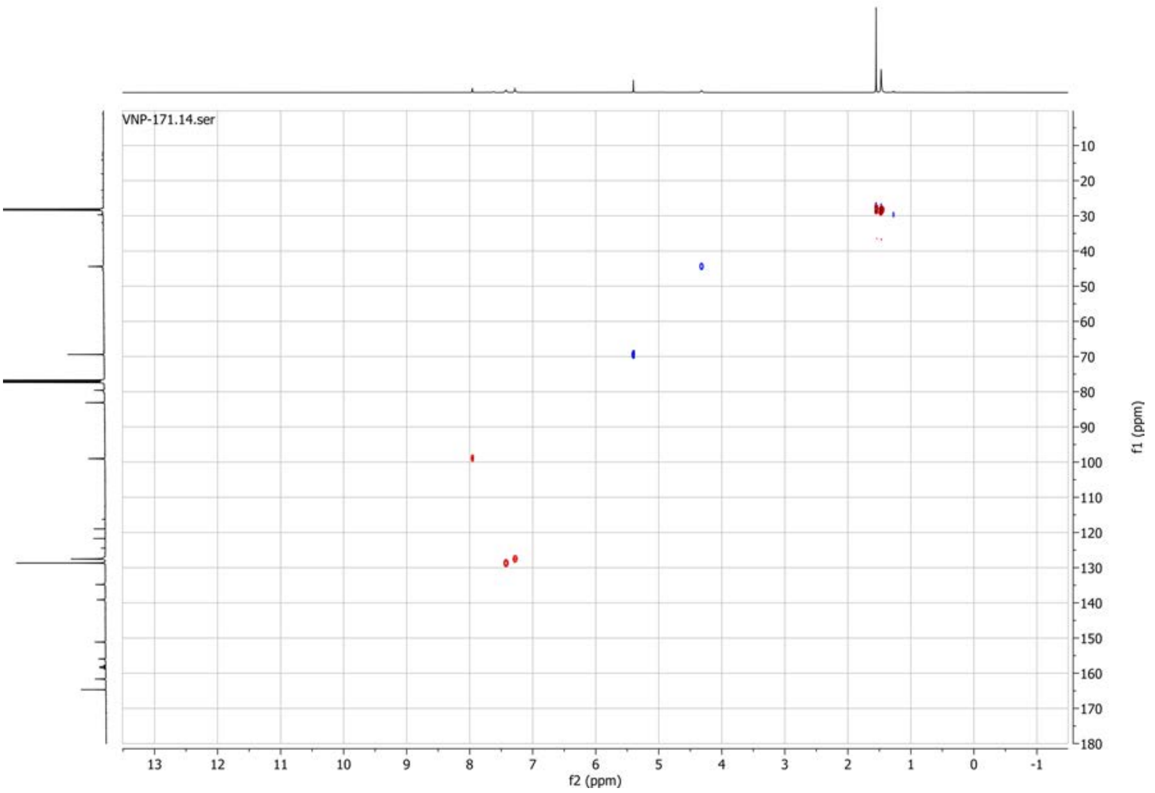


**19F spectrum of compound 57:**

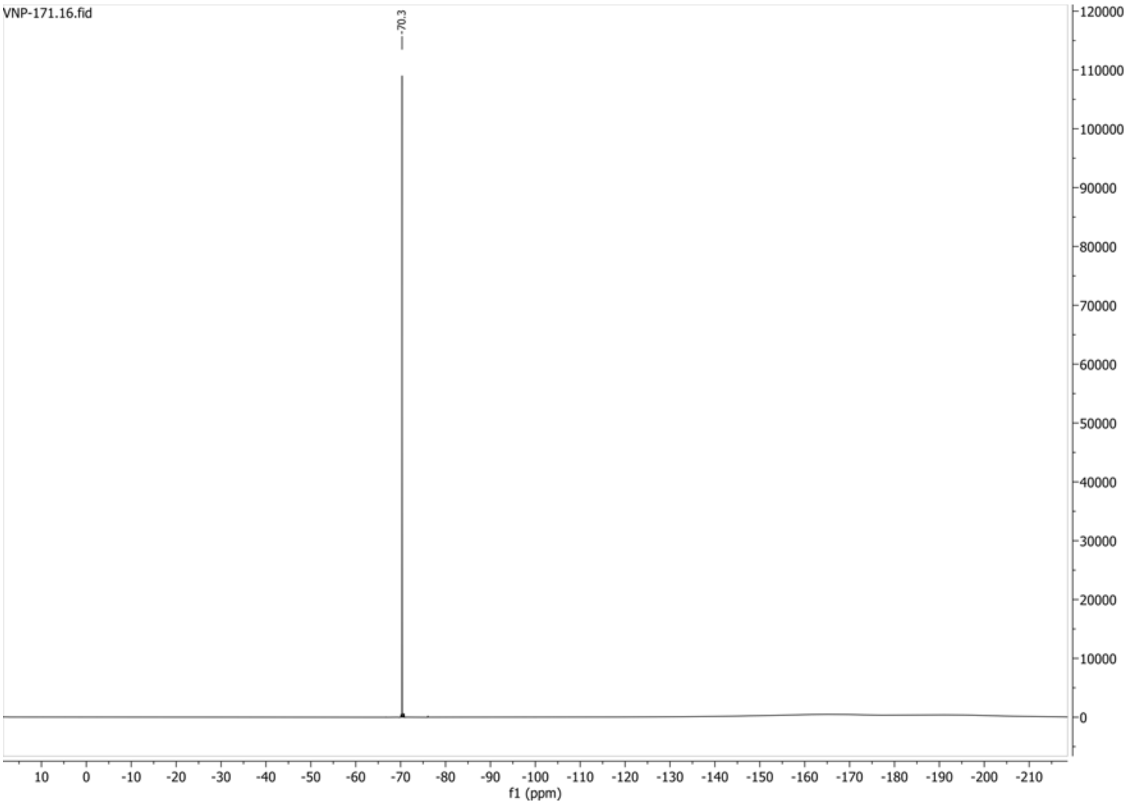

**1H spectrum of compound 58:**

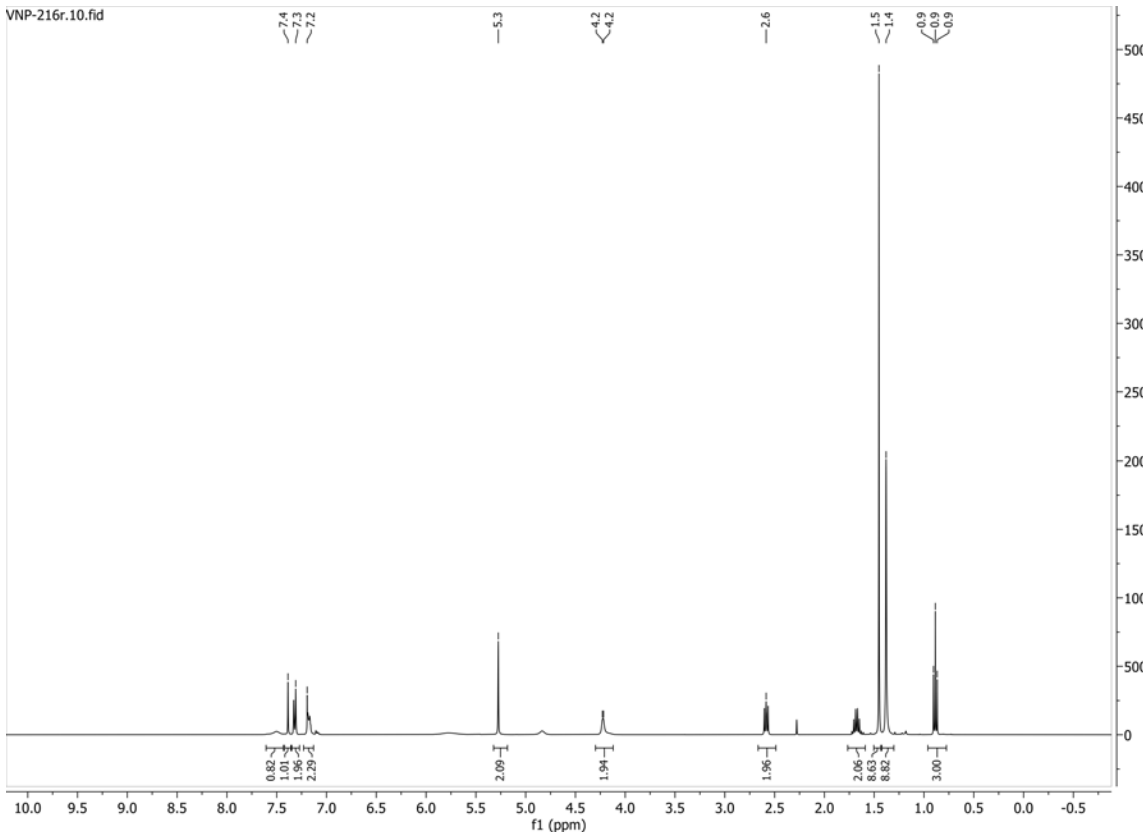


**13C{1H} spectrum of compound 58:**

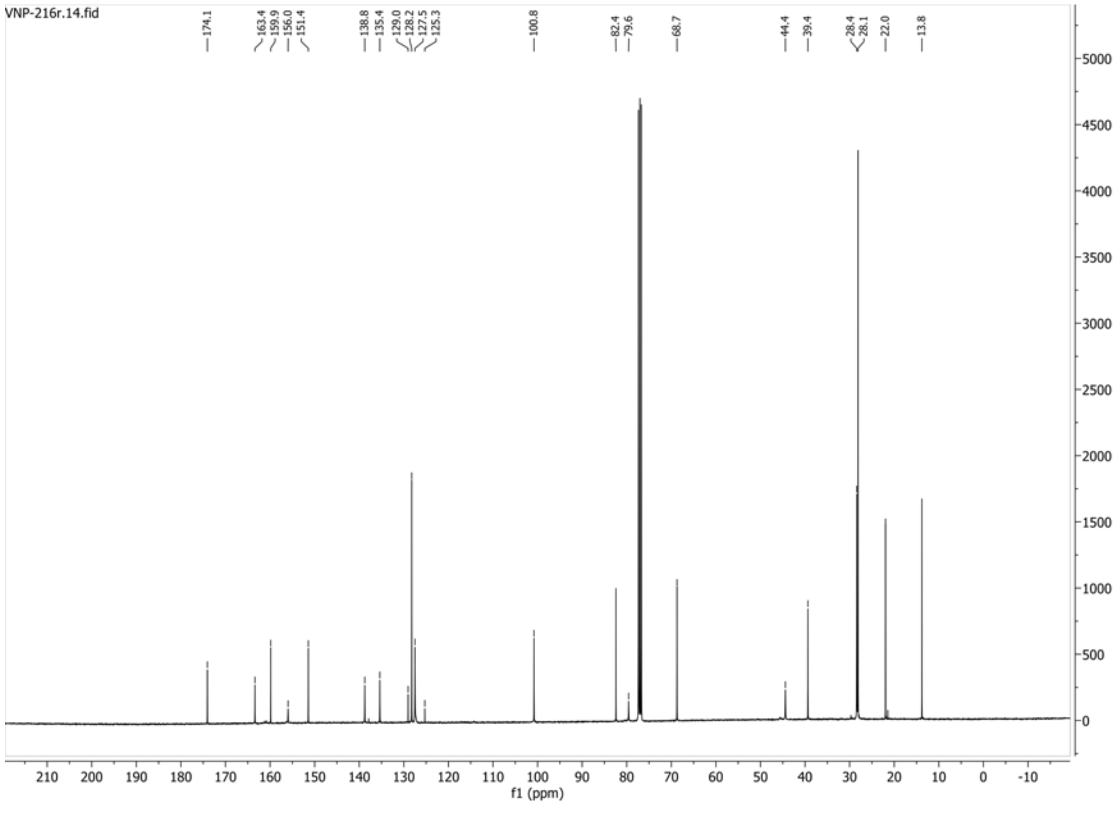

**HSQC-DEPT135 spectrum of compound 58:**

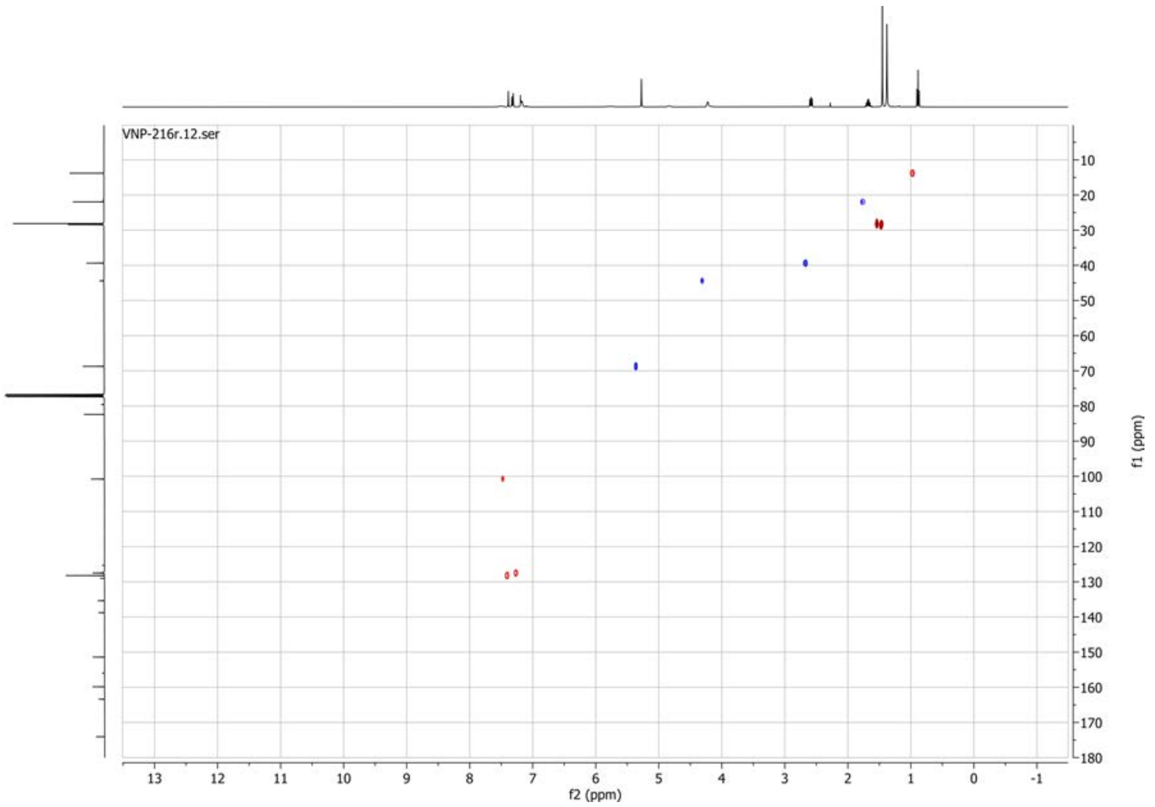

**1H spectrum of compound 59:**

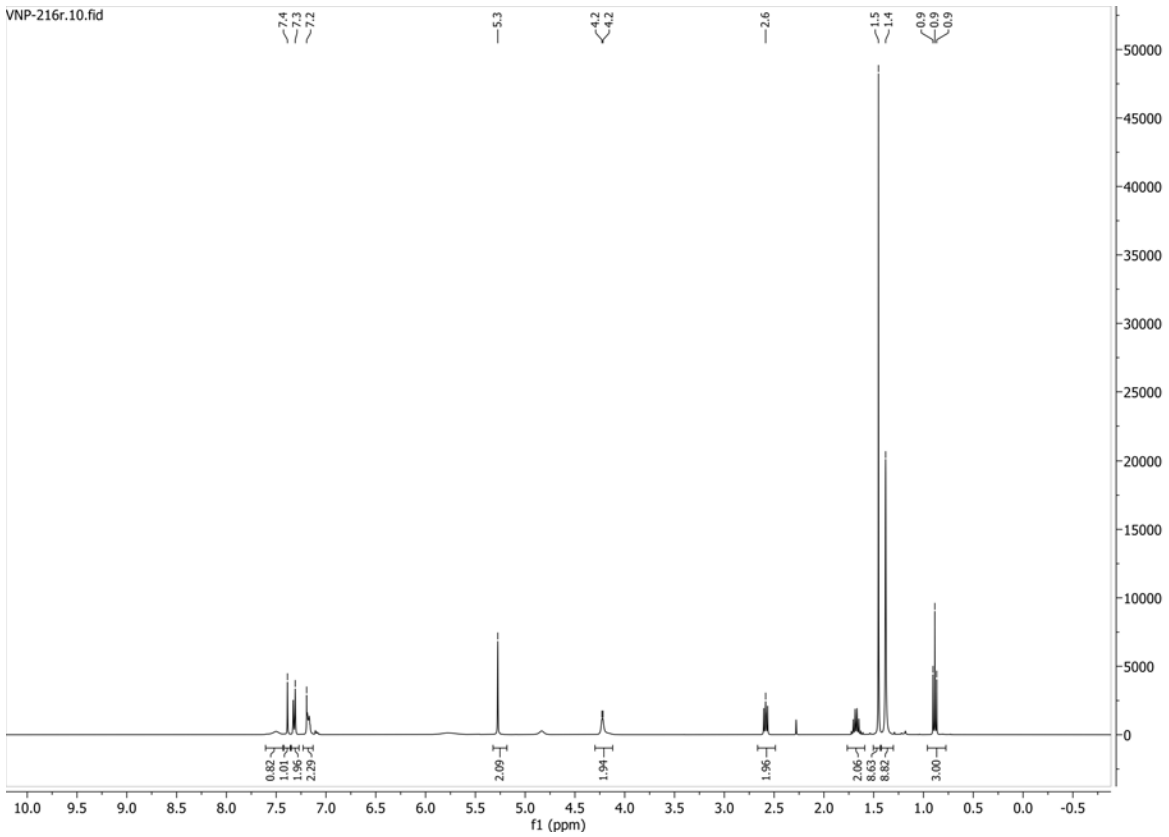


**13C{1H} spectrum of compound 59:**

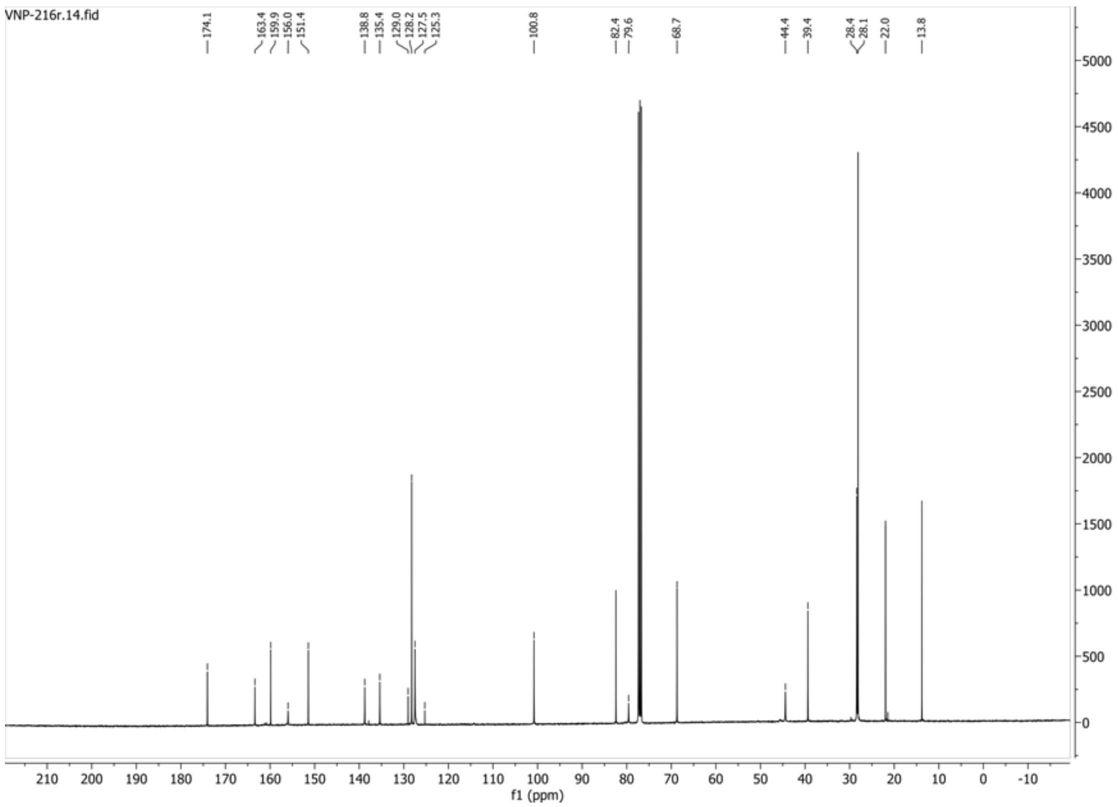

**13C{1H} spectrum of compound 59:**

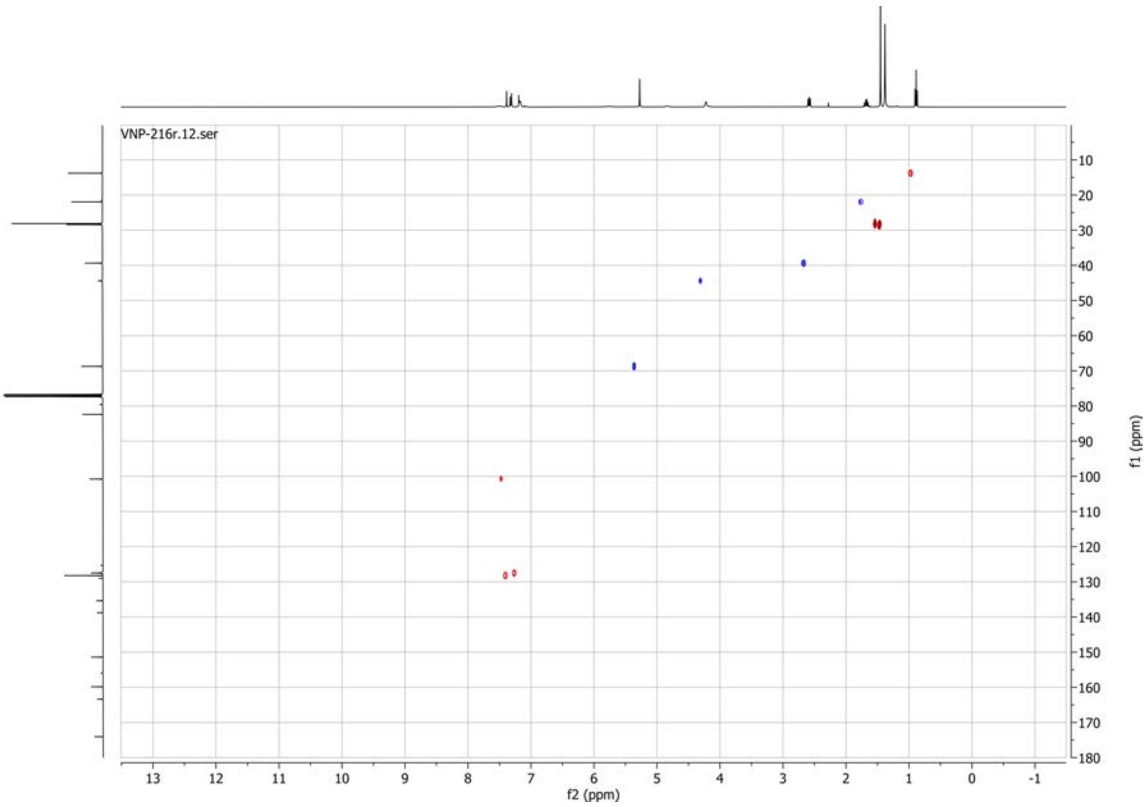

**1H spectrum of compound 60:**

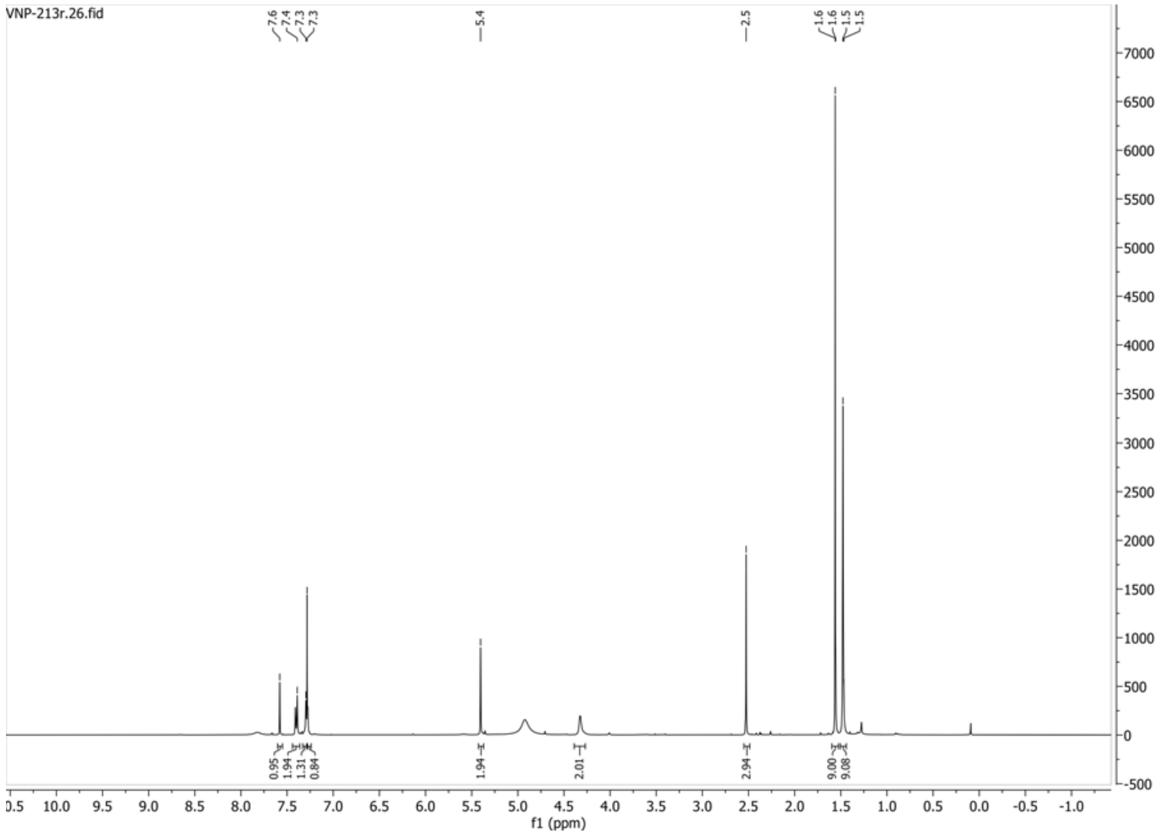


**13C{1H} spectrum of compound 60:**

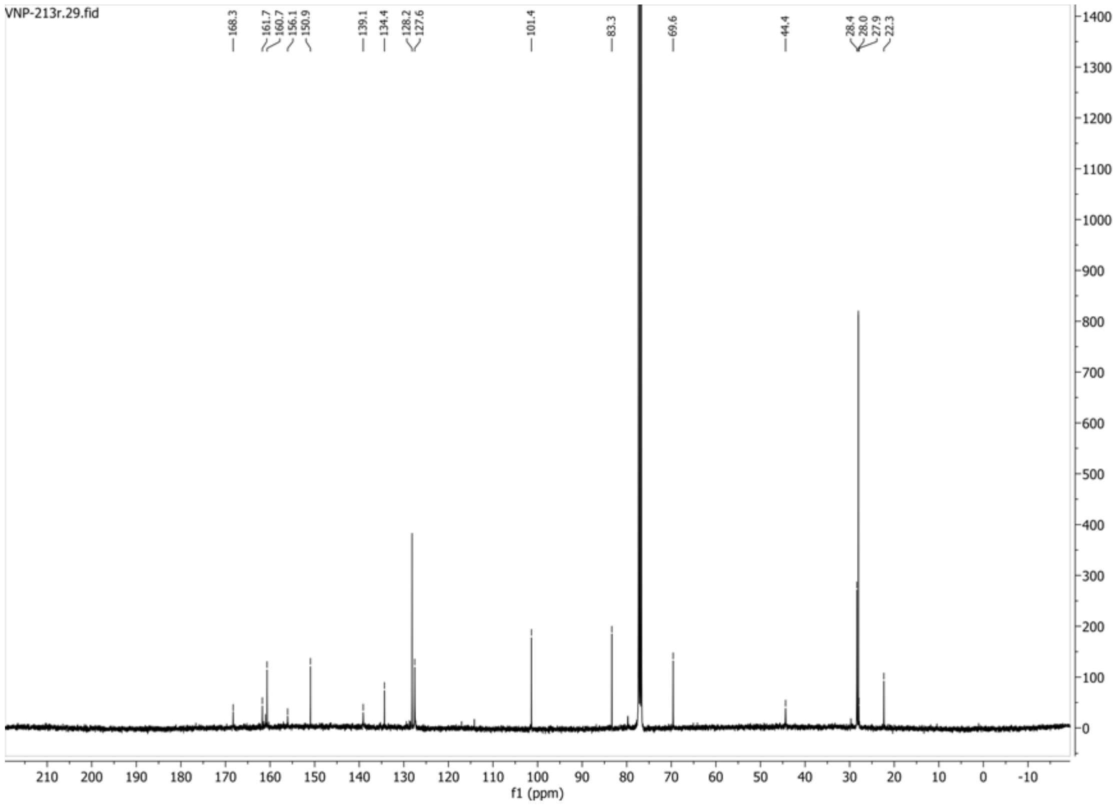

**HSQC-DEPT135 spectrum of compound 60:**

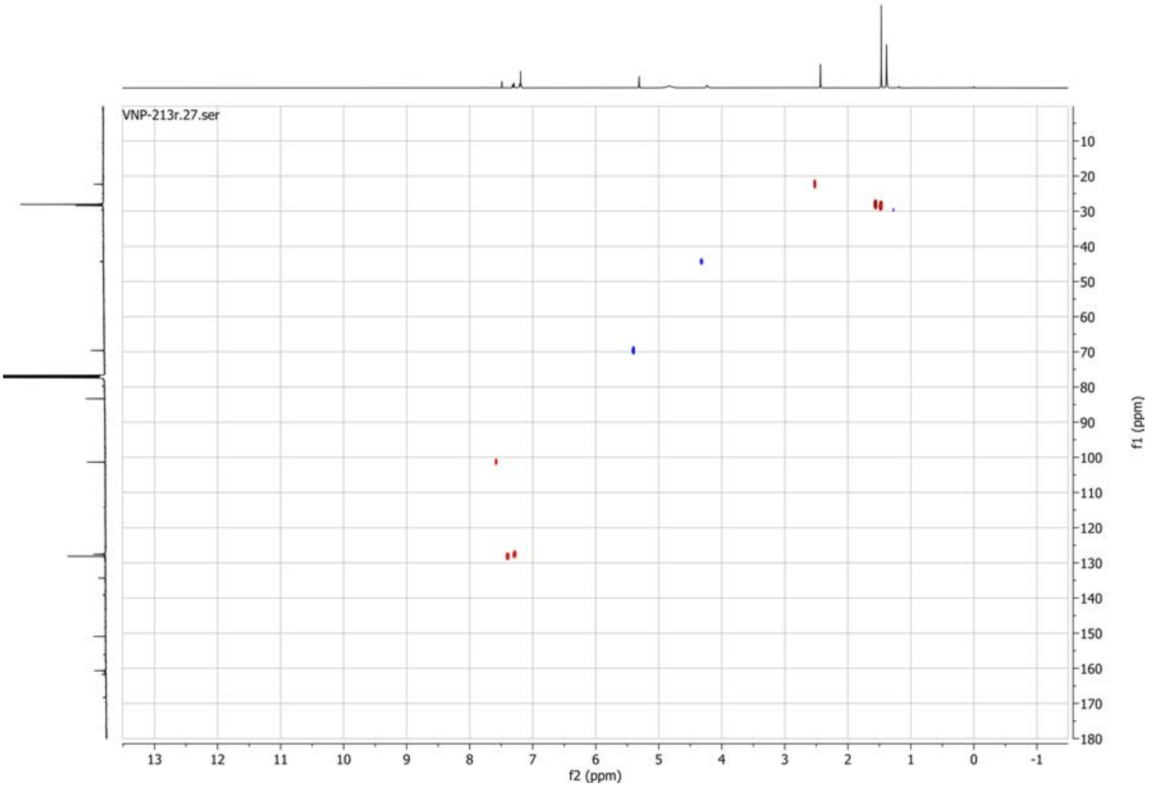

**1H spectrum of compound 62:**

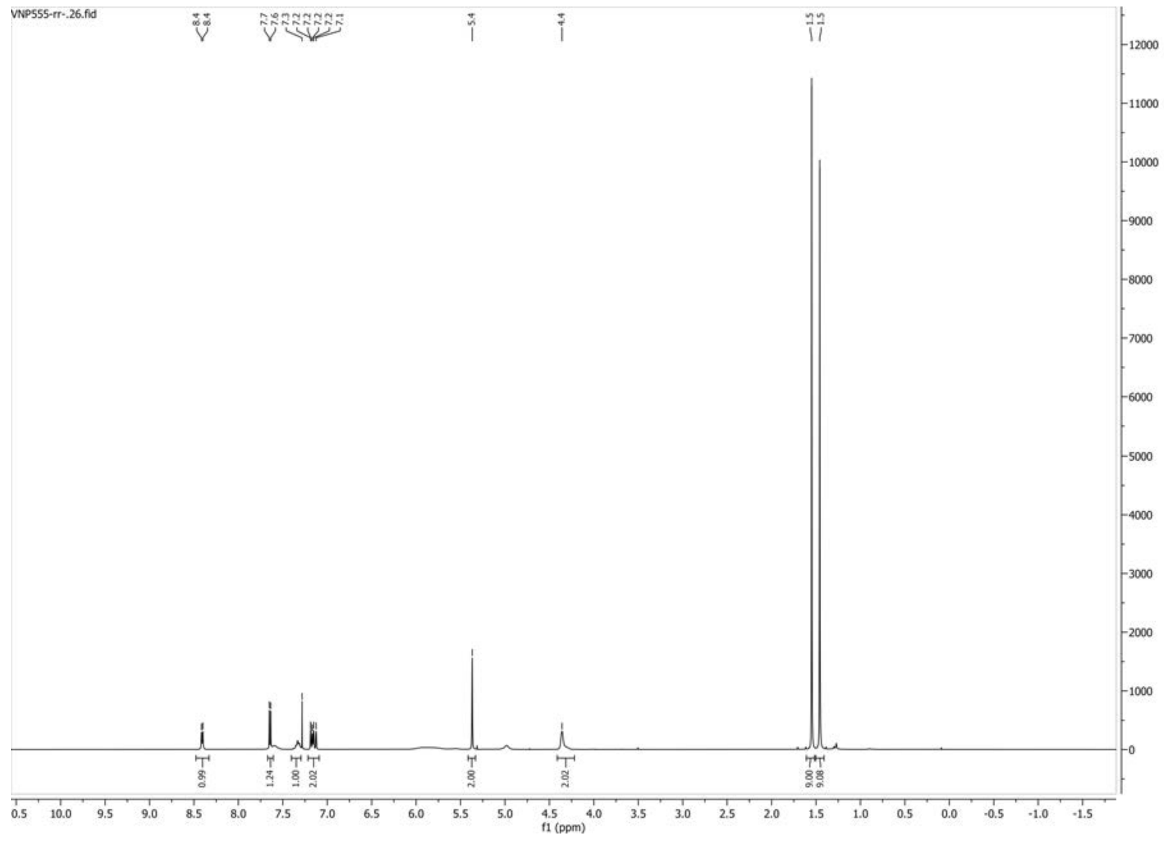

**13C{1H} spectrum of compound 62:**

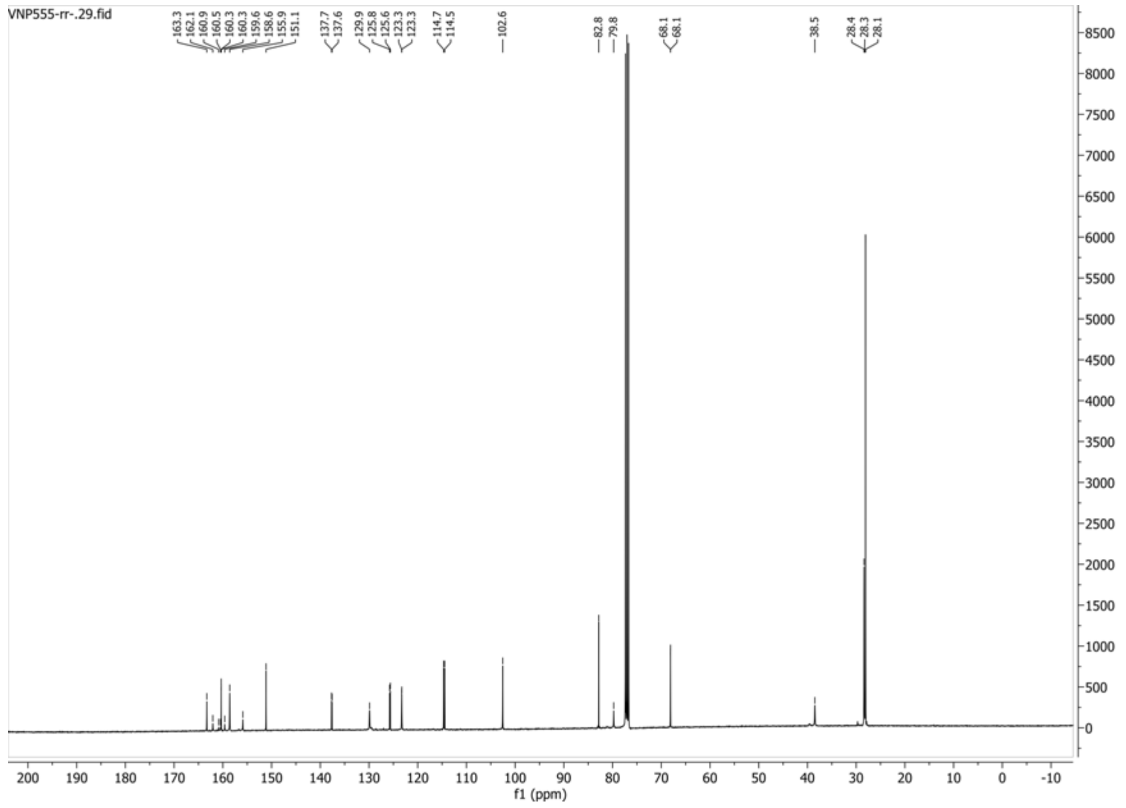

**HSQC-DEPT135 spectrum of compound 62:**

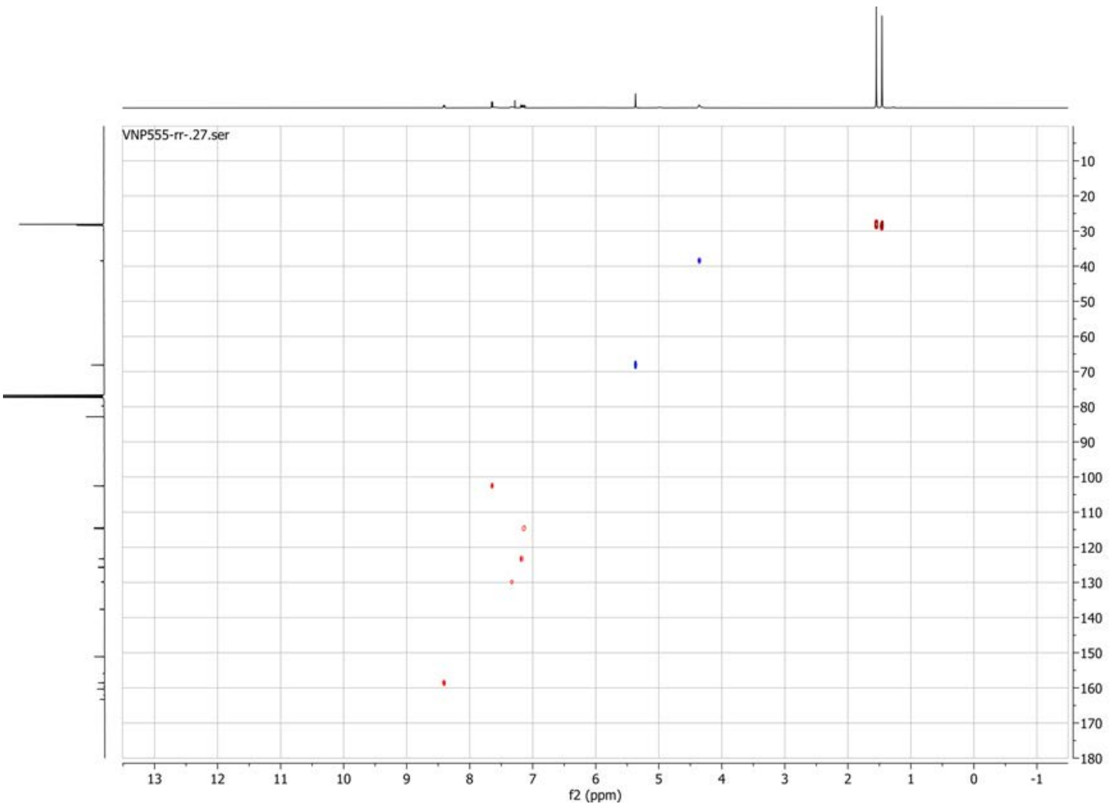


**19F spectrum of compound 62:**

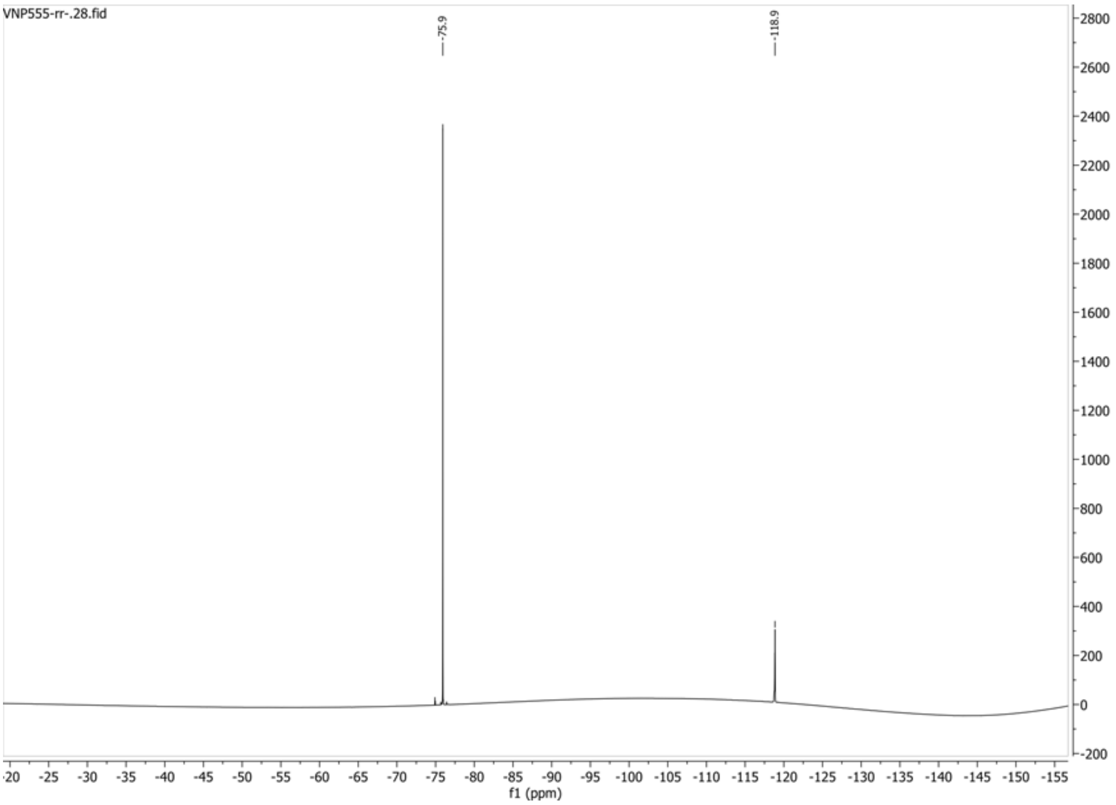

**1H spectrum of compound 64:**

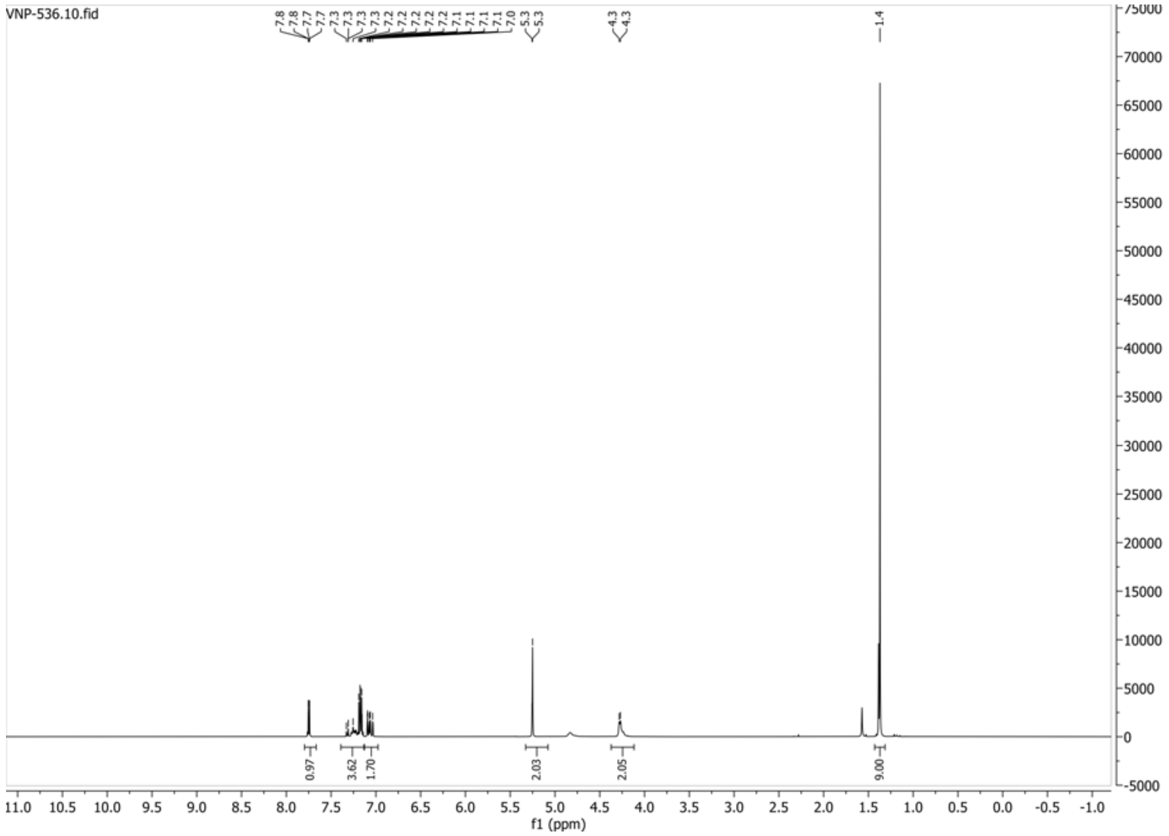


**13C{1H} spectrum of compound 64:**

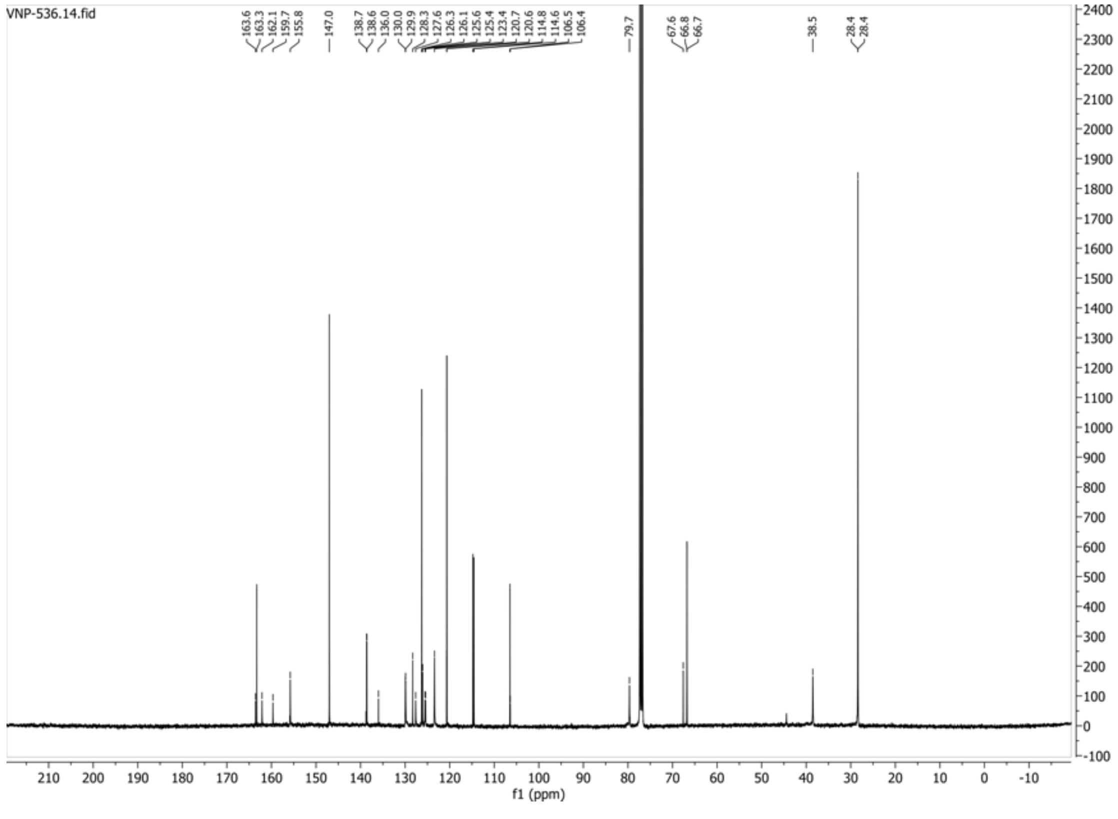

**HSQC-DEPT spectrum of compound 64:**

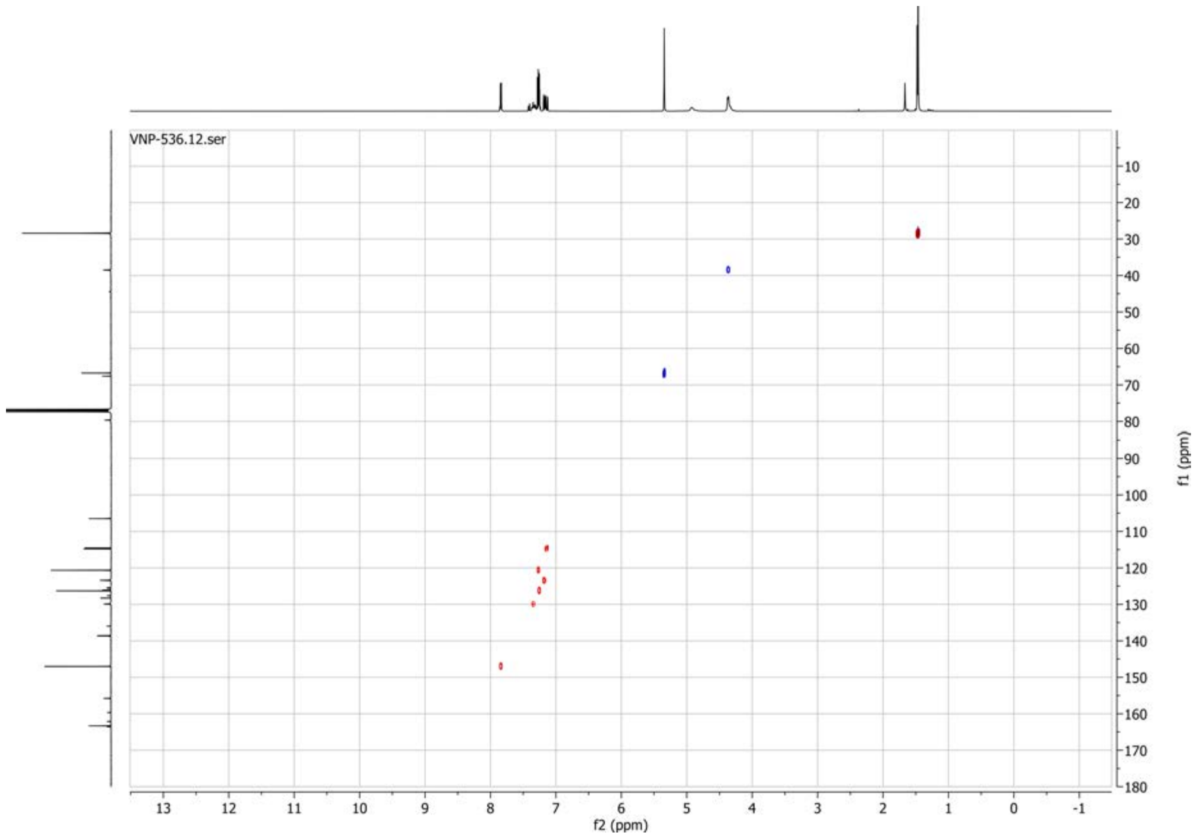


**19F spectrum of compound 64:**

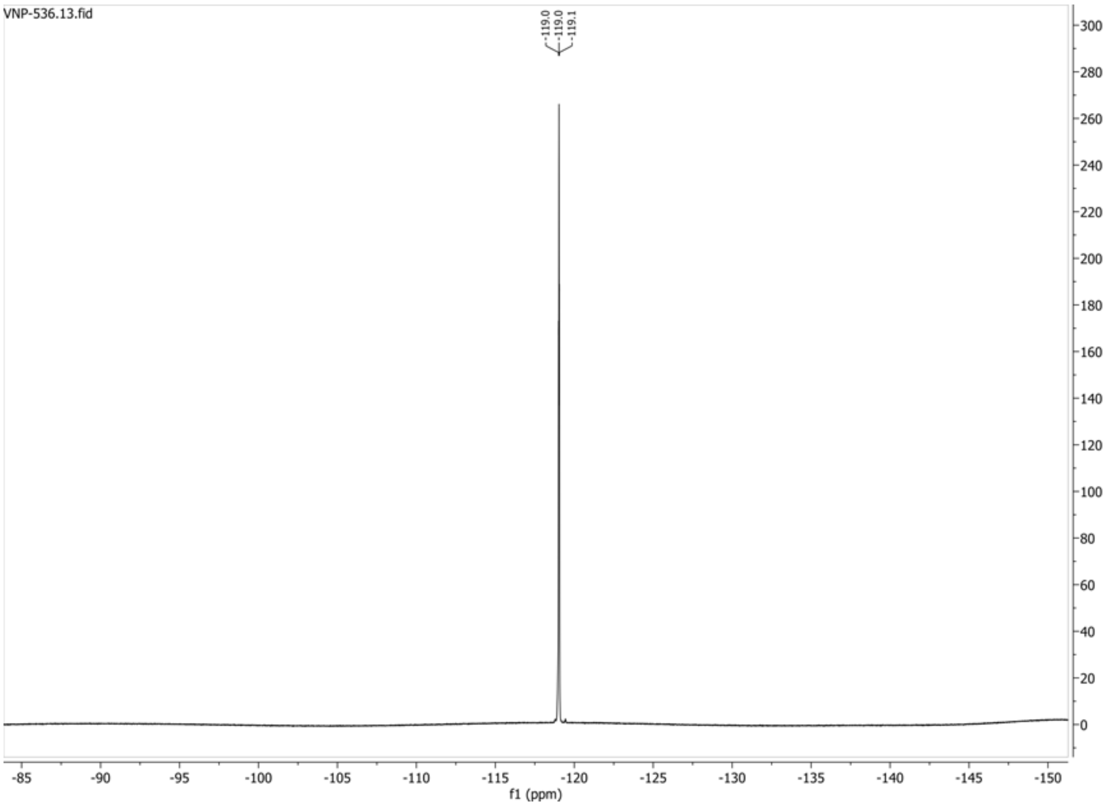

**1H spectrum of compound 65:**

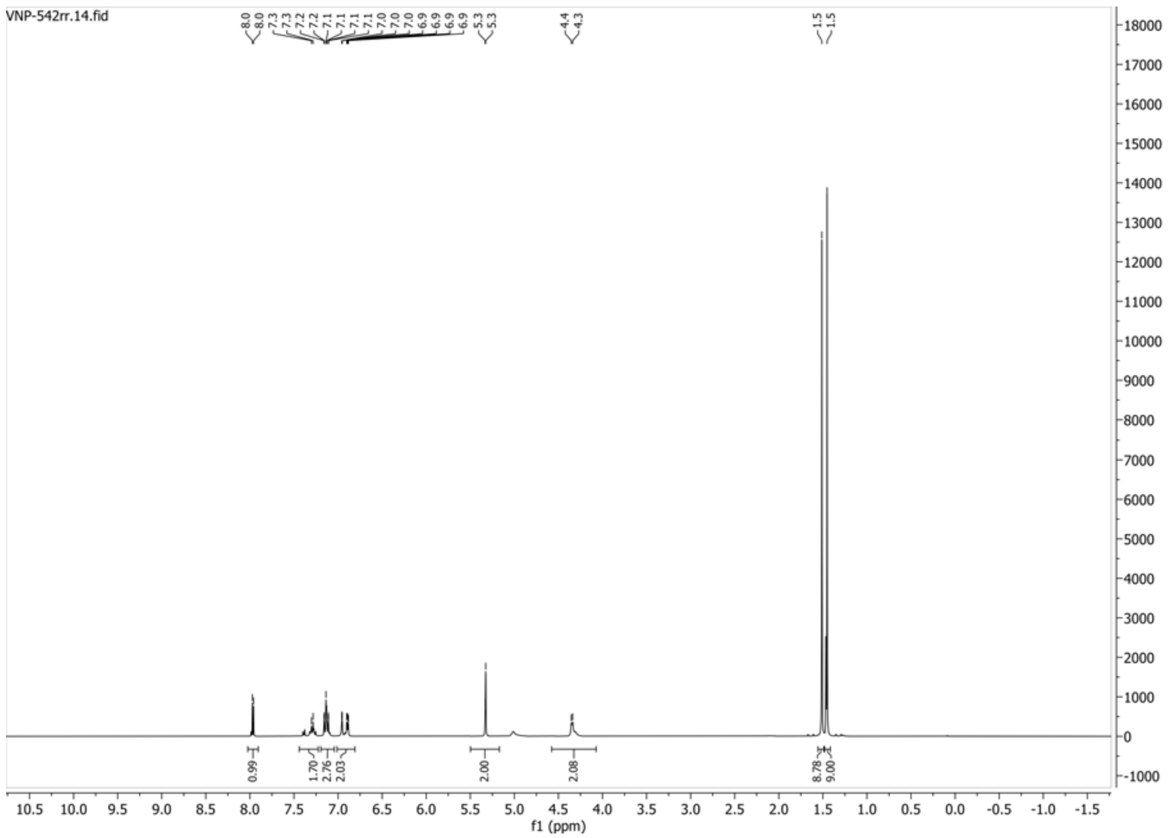

**13C{1H} spectrum of compound 65:**

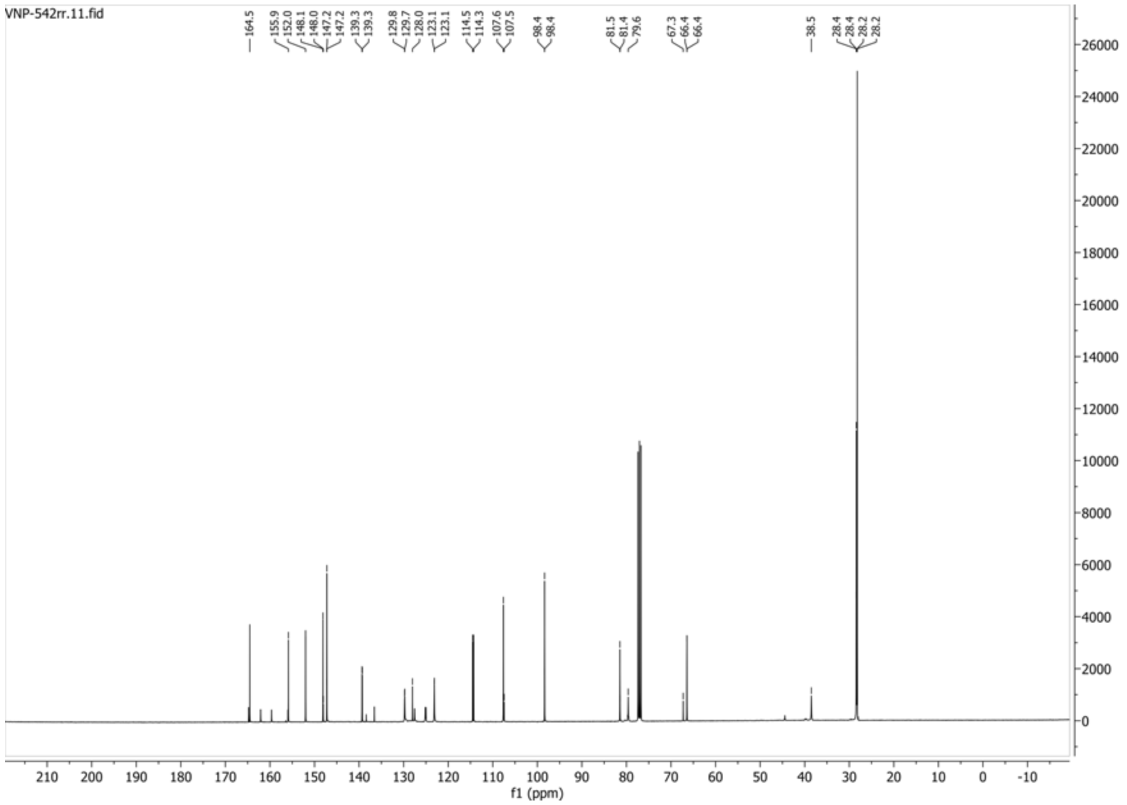

**HSQC-DEPT135 spectrum of compound 65:**

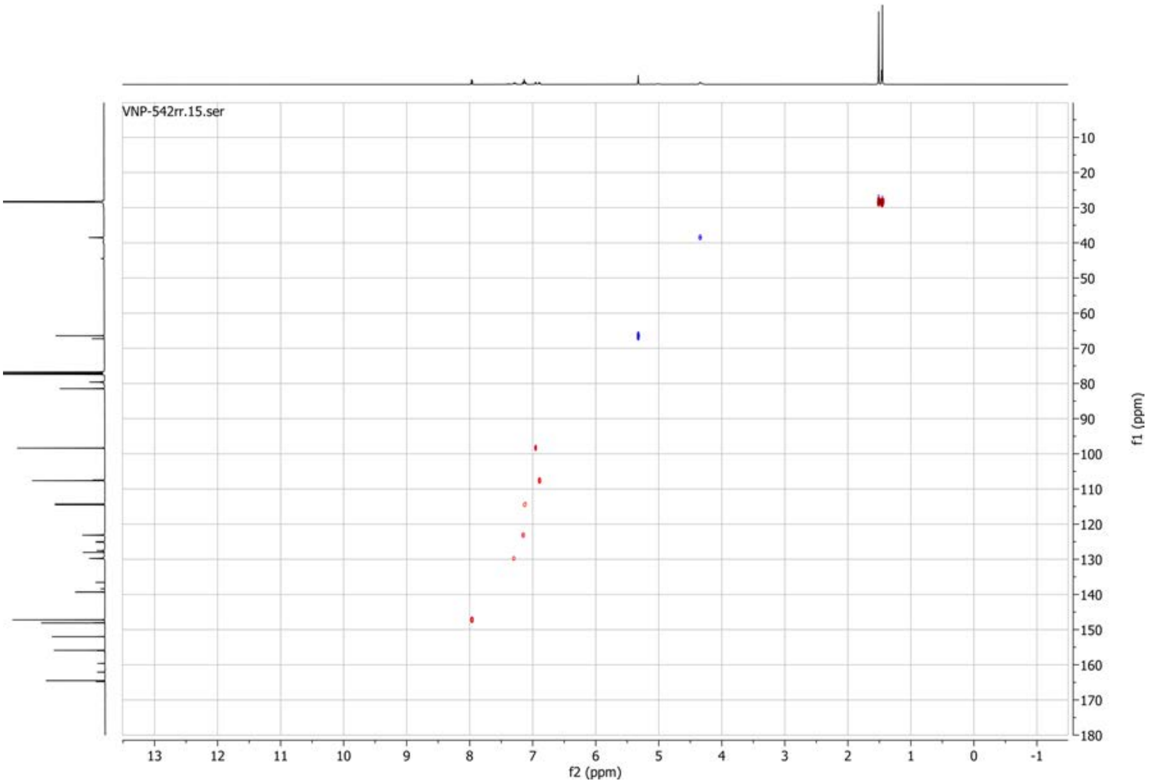


**19F spectrum of compound 65:**

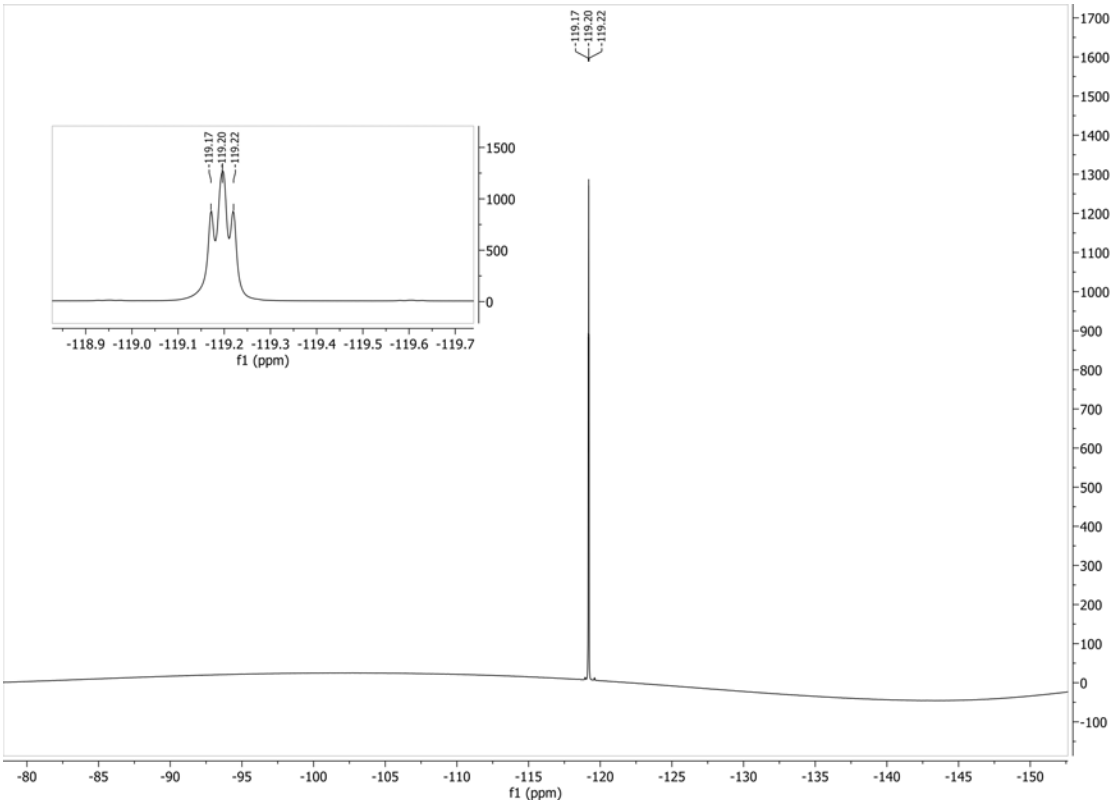

**1H spectrum of compound 61:**

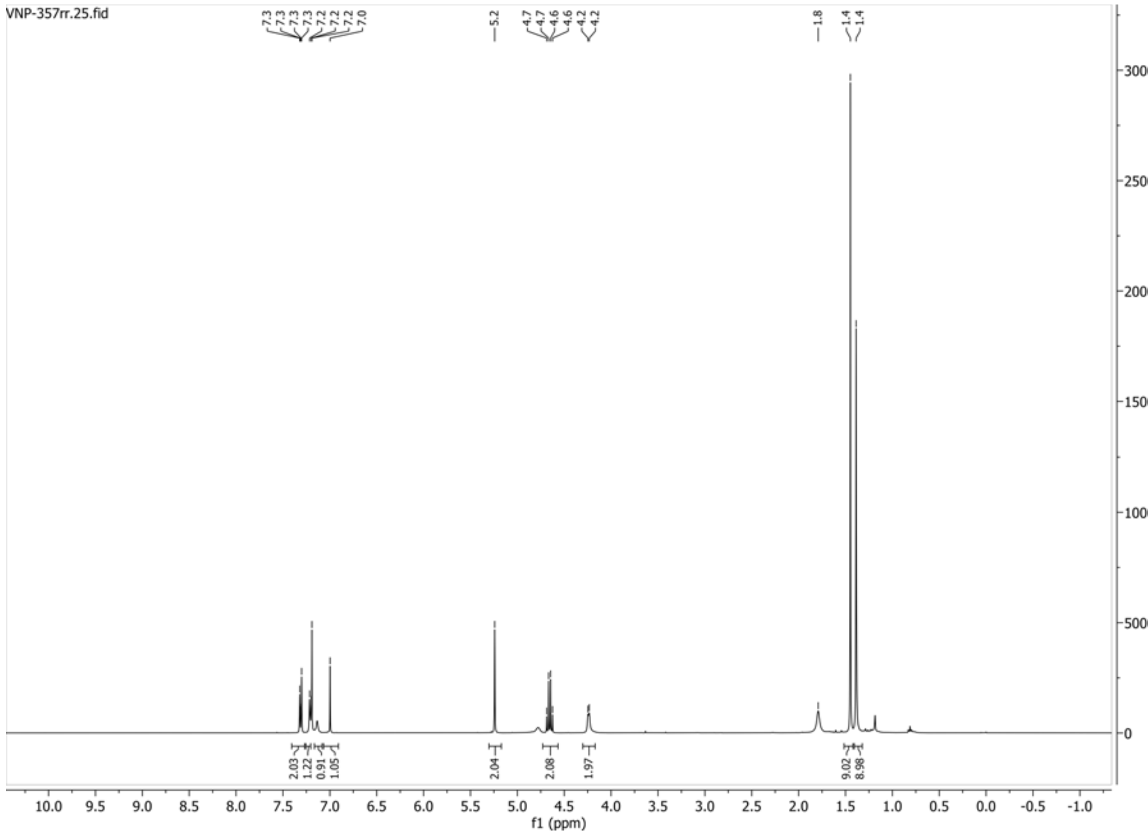


**13C{1H} spectrum of compound 61:**

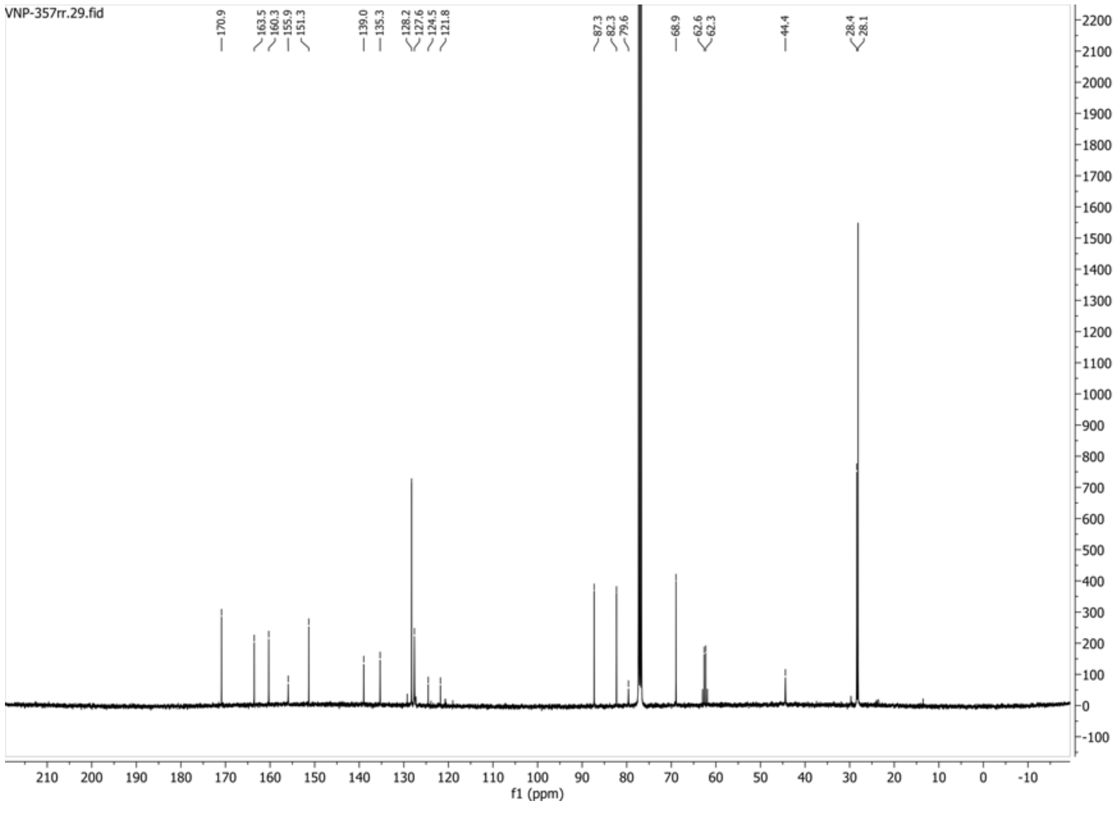

**HSQC-DEPT spectrum of compound 61:**

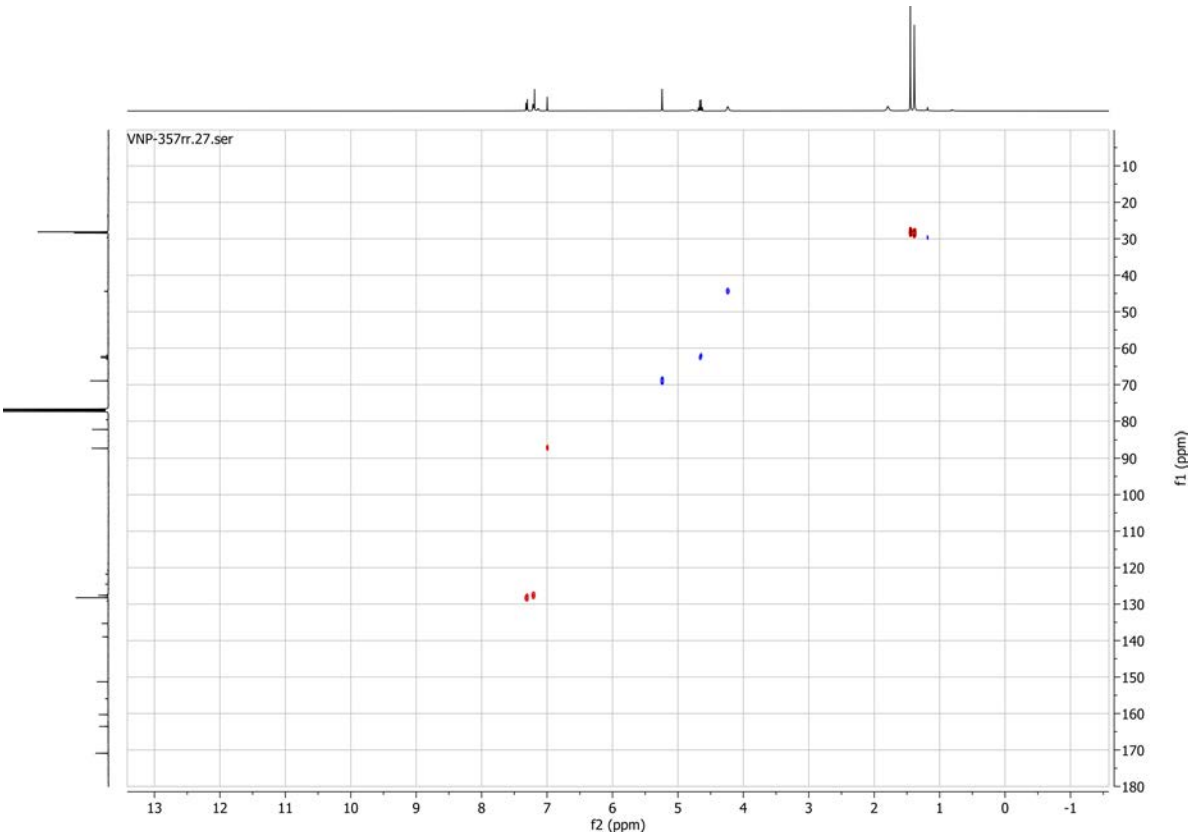


**19F spectrum of compound 61:**

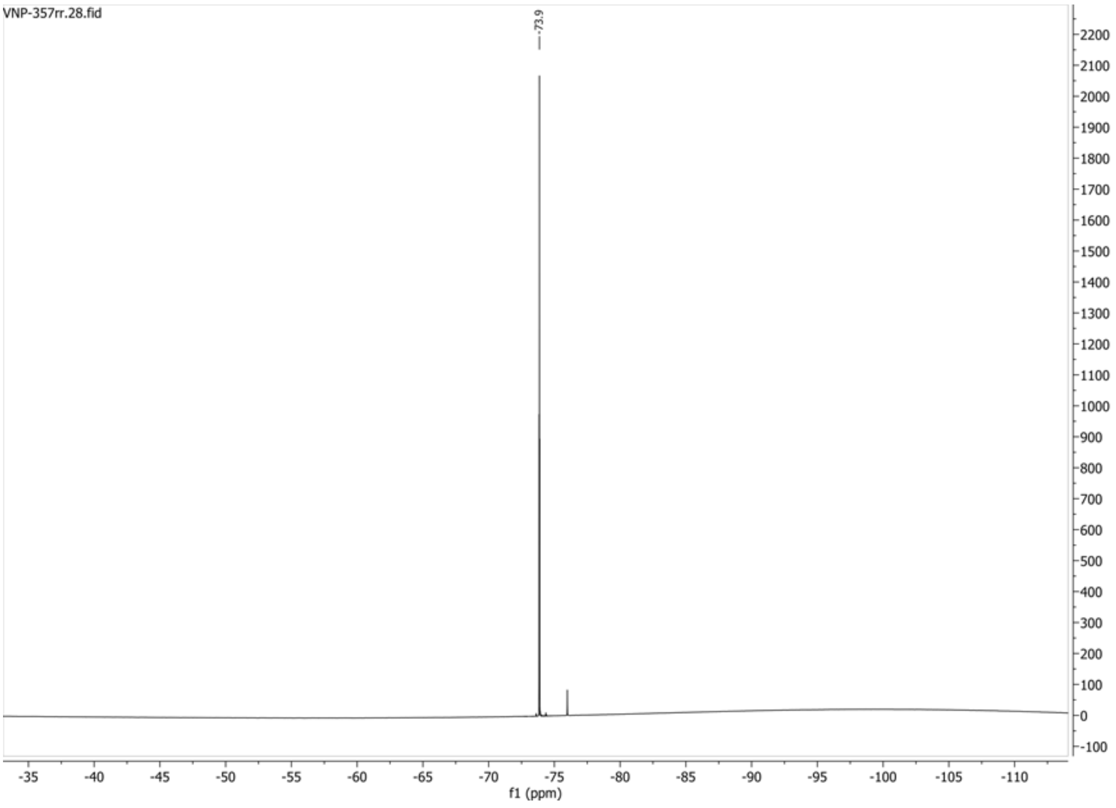
